# The Selection Landscape and Genetic Legacy of Ancient Eurasians

**DOI:** 10.1101/2022.09.22.509027

**Authors:** Evan K. Irving-Pease, Alba Refoyo-Martínez, William Barrie, Andrés Ingason, Alice Pearson, Anders Fischer, Karl-Göran Sjögren, Alma S. Halgren, Ruairidh Macleod, Fabrice Demeter, Rasmus A. Henriksen, Tharsika Vimala, Hugh McColl, Andrew H. Vaughn, Leo Speidel, Aaron J. Stern, Gabriele Scorrano, Abigail Ramsøe, Andrew J. Schork, Anders Rosengren, Lei Zhao, Kristian Kristiansen, Astrid K. N. Iversen, Lars Fugger, Peter H. Sudmant, Daniel J. Lawson, Richard Durbin, Thorfinn Korneliussen, Thomas Werge, Morten E. Allentoft, Martin Sikora, Rasmus Nielsen, Fernando Racimo, Eske Willerslev

**Affiliations:** Lundbeck Foundation GeoGenetics Centre, Globe Institute, University of Copenhagen, Copenhagen, Denmark; GeoGenetics Group, Department of Zoology, University of Cambridge, UK; Institute of Biological Psychiatry, Mental Health Services, Copenhagen University Hospital, Roskilde, Denmark; Department of Genetics, University of Cambridge, UK; Department of Zoology, University of Cambridge, UK; Department of Historical Studies, University of Gothenburg, 405 30 Gothenburg, Sweden; Sealand Archaeology, Gl. Roesnaesvej 27, 4400 Kalundborg, Denmark; Department of Integrative Biology, University of California Berkeley; UCL Genetics Institute, University College London, London, UK; National Museum of Natural History, Paris, France; Center for Computational Biology, University of California, Berkeley, USA; Ancient Genomics Laboratory, The Francis Crick Institute, London, UK; Neurogenomics Division, The Translational Genomics Research Institute (TGEN), Phoenix, AZ, USA; Oxford Centre for Neuroinflammation, Nuffield Department of Clinical Neurosciences, John Radcliffe Hospital, University of Oxford, Oxford, UK; Department of Clinical Medicine, Aarhus University Hospital, Aarhus, Denmark; MRC Human Immunology Unit, John Radcliffe Hospital, University of Oxford, Oxford, UK; Institute of Statistical Sciences, School of Mathematics, University of Bristol, Bristol, UK; Wellcome Sanger Institute, Cambridge, UK; Department of Clinical Medicine and Lundbeck Center for Geogenetics, GLOBE Institute, University of Copenhagen; Institute of Biological Psychiatry, Mental Health Center Sct Hans, Copenhagen University Hospital, Denmark; Trace and Environmental DNA (TrEnD) Laboratory, School of Molecular and Life Science, Curtin University,, Australia; Departments of Integrative Biology and Statistics, UC Berkeley, Berkeley 94720, USA; MARUM Center for Marine Environmental Sciences and Faculty of Geosciences, University of Bremen, Bremen, Germany

## Abstract

The Holocene (beginning ∼12,000 years ago) encompassed some of the most significant changes in human evolution, with far-reaching consequences for the dietary, physical, and mental health of present-day populations. Using a dataset of >1600 imputed ancient genomes ^1^, we modelled the selection landscape during the transition from hunting and gathering, to farming and pastoralism across West Eurasia. We identify major selection signals related to metabolism, including that selection at the FADS cluster began earlier than previously reported, and that selection near the LCT locus predates the emergence of the lactase persistence allele by thousands of years. We also find strong selection in the HLA region, possibly due to increased exposure to pathogens during the Bronze Age. Using ancient individuals to infer local ancestry tracts in >400,000 samples from the UK Biobank, we identify widespread differences in the distribution of Mesolithic, Neolithic, and Bronze Age ancestries across Eurasia. By calculating ancestry-specific polygenic risk scores, we show that height differences between Northern and Southern Europe are associated with differential Steppe ancestry, rather than selection, and that risk alleles for mood-related phenotypes are enriched for Neolithic farmer ancestry, while risk alleles for diabetes and Alzheimer’s disease are enriched for Western Hunter-gatherer ancestry. Our results suggest that ancient selection and migration were major contributors to the distribution of phenotypic diversity in present-day Europeans.

## Main

One of the central goals of human evolutionary genetics is to understand how natural selection has shaped the genomes of present-day people in response to changes in culture and environment. The transition from hunter-gatherers, to farmers, and subsequently pastoralists, during the Holocene in Eurasia, involved some of the most dramatic changes in diet, health and social organisation experienced during recent human evolution. These changes represent major shifts in environmental exposure, impacting the evolutionary forces acting on the human gene pool and imposing a series of heterogeneous selection pressures. As human lifestyles changed, close contact with domestic animals and higher population densities are likely to have increased exposure to infectious diseases, introducing new challenges to our immune system ^2,3^.

Our understanding of the genetic architecture of complex traits in humans has been substantially advanced by genome-wide association studies (GWAS), which have identified large numbers of genetic variants associated with phenotypes of interest ^4,5^. However, the extent to which these variants have been under directional selection during recent human evolution remains unclear. While signatures of selection can be identified from patterns of genetic diversity in extant populations ^6^, this can be challenging in humans, which have been exposed to highly diverse and dynamic local environments through time and space. In the complex mosaic of genetic affinities that constitute a present-day human genome, any putative signatures of selection may misrepresent the timing and magnitude of the selective process. For example, episodes of admixture between ancestral populations can result in present-day haplotypes which contain no evidence of selective processes occurring further back in time. Ancient DNA (aDNA) provides the potential to resolve these issues, by directly observing changes in trait-associated allele frequencies over time.

Whilst numerous prior studies have used ancient DNA to infer patterns of selection in Eurasia during the Holocene (e.g., ^7–9)^, many key questions remain unanswered. To what extent are present-day genetic differences due to natural selection or to differential patterns of admixture? What are the genetic legacies of Mesolithic, Neolithic, and Bronze Age populations in present-day complex traits? How has the complex admixture history of Holocene Eurasia affected our ability to detect natural selection in genetic data? To investigate these questions, we tested for traces of divergent selection in health and lifestyle-related genetic variants using three broad approaches. Firstly, we looked for evidence of selection by identifying strong differentiation in allele frequencies between ancient populations. Secondly, we reconstructed the allele frequency trajectories and selection coefficients of tens of thousands of trait-associated variants, using a novel chromosome painting technique to model ancestry-specific allele frequency trajectories through time. This allowed us to identify many trait-associated variants with novel evidence for directional selection, and to answer long-standing questions about the timing of selection for key health, dietary and pigmentation associated loci. Lastly, we used ancient genomes to infer local ancestry tracts in >400,000 present-day genomes from the UK Biobank ^5^, and calculated ancestry-specific polygenic risk scores for 35 complex traits. This allowed us to characterise the genetic legacy of Mesolithic, Neolithic, and Bronze Age populations in present-day phenotypes.

### Samples and data

Our analyses are undertaken on a large collection of shotgun-sequenced ancient genomes presented in the accompanying study ‘Population Genomics of Postglacial Western Eurasia’ ^1^. This dataset comprises 1,664 imputed diploid ancient genomes and more than 8.5 million SNPs, with an estimated imputation error rate of 1.9% and a phasing switch error rate of 2.0% for 1X genomes. Full details of the validation and benchmarking of the imputation and phasing of this dataset are provided in reference ^10^. These samples represent a considerable transect of Eurasia, ranging longitudinally from the Atlantic coast to Lake Baikal, and latitudinally from Scandinavia to the Middle East (Fig. 1). The included genomes constitute a thorough temporal sequence from 11,000 cal. BP to 1,000 cal. BP. This dataset allowed us to characterise in fine detail the changes in selective pressures exerted by major transitions in human culture and environment.

**Fig 1.**
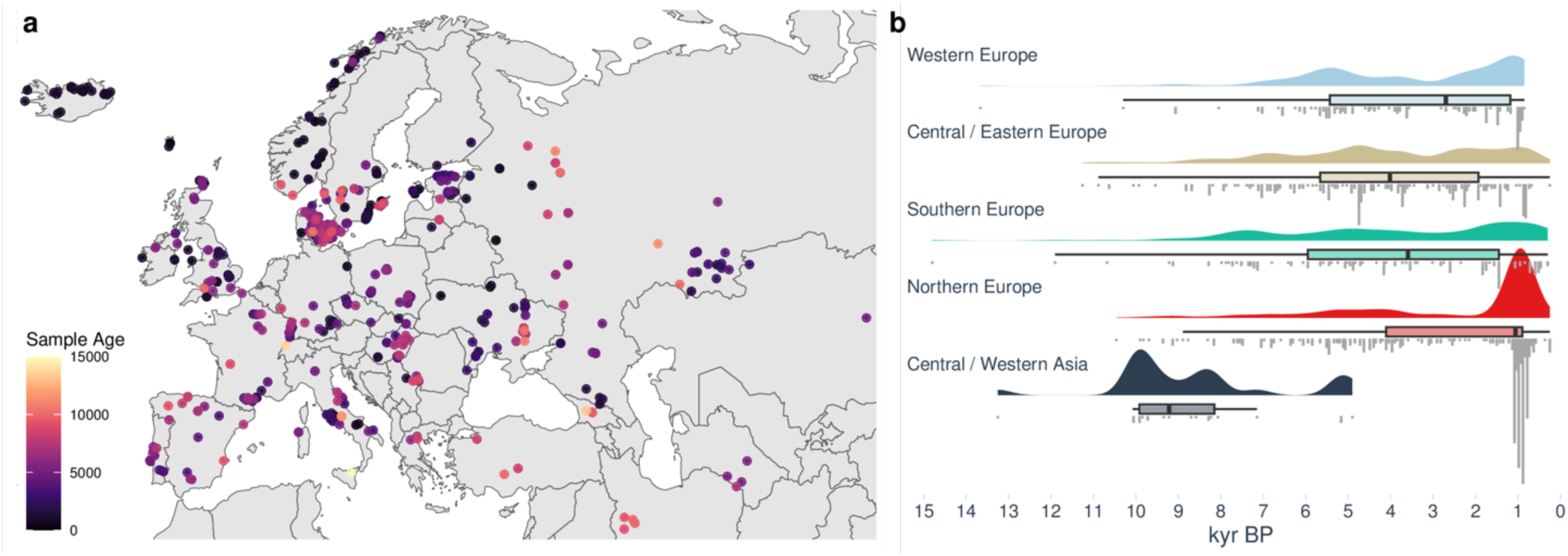
Geographic and temporal distribution of the 1,015 ancient genomes from West Eurasia. **a**, Map of West Eurasia showing sampling locations and ages of the ancient samples; **b**, Raincloud plot of the sample ages, grouped by sampling region: Western Europe (n=156), Central / Eastern Europe (n=268), Southern Europe (n=136), Northern Europe (n=432), and Central / Western Asia (n=23). Boxplot shows the median, and first and third quartiles of the sample ages, and whiskers extend to the largest value no further than 1.5 times the interquartile range.

### Genetic legacy of ancient Eurasians

We began our analysis by inferring local ancestry tracts in present-day populations by chromosome ‘painting’ ^11^ the UK Biobank (UKB) with Mesolithic, Neolithic, and Bronze Age individuals as tract sources. We used a pipeline adapted from GLOBETROTTER ^12^, and estimated admixture proportions via Non-Negative Least Squares (Supplementary Note 2). In total, we painted 433,395 present-day genomes, including 24,511 individuals born outside the UK, from 126 countries (Supplementary Note 1). Our results show that none of the Mesolithic, Neolithic or Bronze Age ancestries are homogeneously distributed among present-day Eurasian populations (Fig. 2). Western hunter-gatherer (WHG) related ancestries are highest in present-day individuals from the Baltic States, Belarus, Poland, and Russia; Eastern hunter-gatherer (EHG) related ancestries are highest in Mongolia, Finland, Estonia and Central Asia; and Caucasus hunter-gatherer (CHG) related ancestries are highest in countries east of the Caucasus, in Pakistan, India, Afghanistan and Iran, in accordance with previous results ^13^. The CHG-related ancestries likely reflect affinities to both Caucasus hunter-gatherer and Iranian Neolithic individuals, explaining the relatively high levels in south Asia ^14^. Consistent with expectations ^15^, Neolithic Anatolian-related farmer ancestries are concentrated around the Mediterranean basin, with high levels in southern Europe, the Near East, and North Africa, including the Horn of Africa, but are less frequent in Northern Europe. This is in direct contrast to the Steppe-related ancestries, which are found in high levels in northern Europe, peaking in Ireland, Iceland, Norway, and Sweden, and decreasing further south. There is also evidence for their spread into southern Asia. Overall, these results refine global patterns of spatial distributions of ancient ancestries amongst present-day individuals. We caution, however, that absolute admixture proportions should be interpreted with caution in regions where our ancient source populations are less directly related to present-day individuals, such as in Africa and East Asia. Whilst these values are dependent on the reference samples used, as well as the treatment of pre- or post-admixture drift, the relative geographical variation and associations should remain consistent.

**Fig 2.**
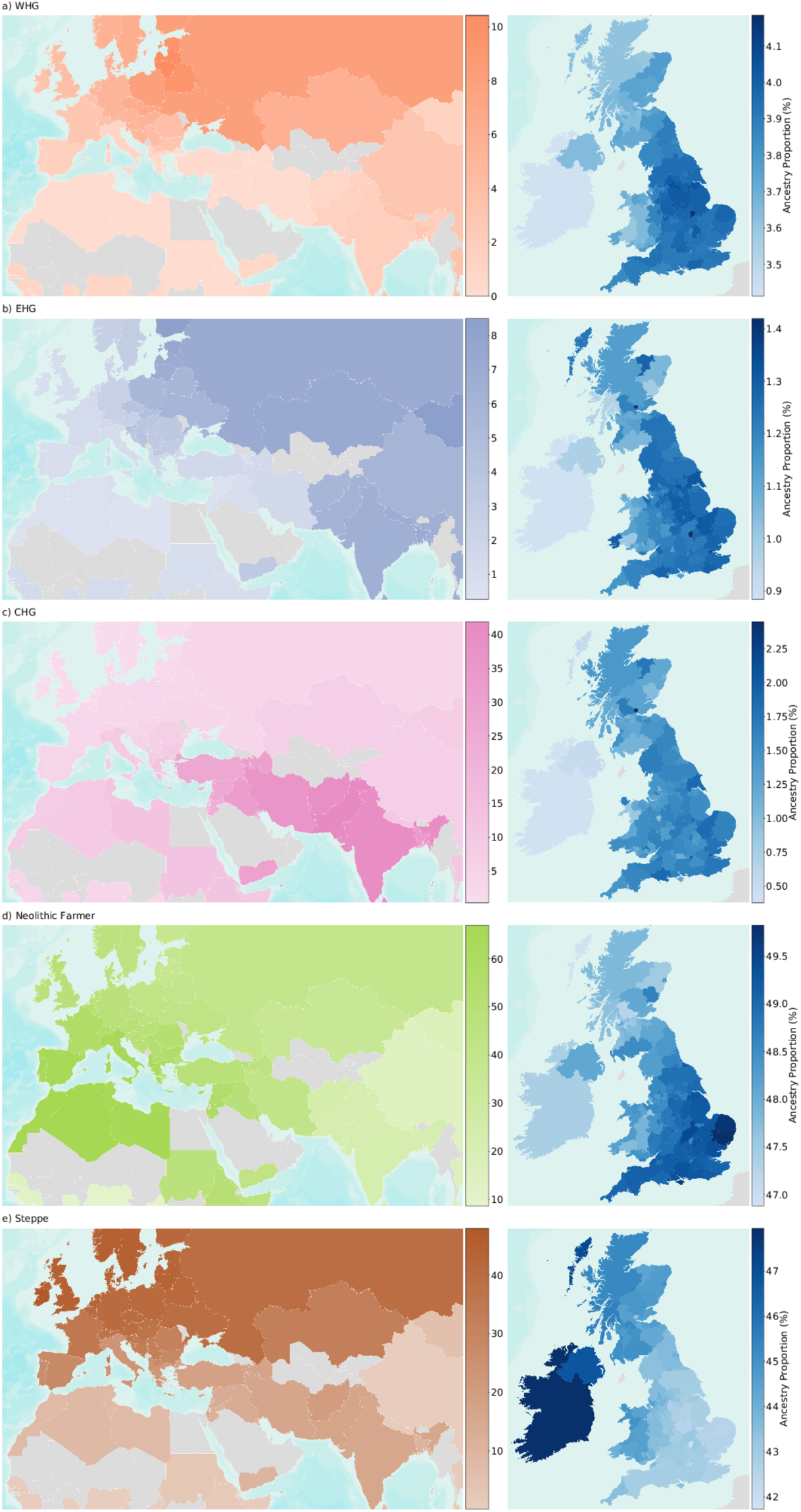
The genetic legacy of ancient Eurasian ancestries in present-day populations. Maps showing the average ancestry of **a**, Western hunter-gatherer (WHG); **b**, Eastern hunter-gatherer (EHG); **c**, Caucasus hunter-gatherer (CHG); **d**, Neolithic farmer; and **e**, Steppe pastoralist ancestry components per country (left) and per county or unitary authority within Great Britain and per-country for the Republic of Ireland and Northern Ireland (right). Estimation was performed using ChromoPainter and NNLS, on samples of a ‘typical ancestral background’ for each non-UK country (n=24,511) and Northern Ireland. For Great Britain, an average of self-identified ‘white British’ samples were used to represent each UK county and unitary authority, based on place of birth (n=408,884). Countries with <4 and counties with <15 samples are shown in grey. Map uses ArcGIS layers World Countries Generalized and World Terrain.

The availability of a large number of present-day genomes (n=408,884) from self-identified “white British” individuals who share similar positions on a PCA ^5^ allowed us to further examine the distribution of ancient ancestries at high resolution in present-day Britain (Supplementary Note 2). Although regional ancestry distributions differ by only a few percentage points, we find clear evidence of geographical heterogeneity across the United Kingdom. This can be visualised by averaging ancestry proportions per county, based on place of birth (Fig. 2, inset boxes). The proportion of Neolithic farmer ancestries is highest in southern and eastern England today and lower in Scotland, Wales, and Cornwall. Steppe-related ancestries are inversely distributed, peaking in the Outer Hebrides and Ireland, a pattern only previously described for Scotland ^16^. This regional pattern was already evident in the Pre-Roman Iron Age and persists to the present day even though immigrating Anglo-Saxons had relatively less affinities to Neolithic farmers than the Iron-Age individuals of southwest Briton. Although this Neolithic farmer/Steppe-related dichotomy mirrors the modern ‘Anglo-Saxon’/‘Celtic’ ethnic divide, its origins are older, resulting from continuous migration from a continental population relatively enriched in Neolithic farmer ancestries, starting as early as the Late Bronze Age ^17,18^. By measuring haplotypes from these ancestries in present-day individuals, we show that these patterns differentiate Wales and Cornwall as well as Scotland from England. We also find higher levels of WHG-related ancestries in central and Northern England. These results demonstrate clear ancestry differences within an ‘ethnic group’ (white British), highlighting the need to account for subtle population structure when using resources such as the UK Biobank ^19^.

### Ancestry-stratified selective sweeps

Having identified that significant differences in ancestries persist in seemingly homogeneous present-day populations, we sought to disentangle these effects by developing a novel chromosome painting technique that allows us to label haplotypes based on their genetic affinities to ancient individuals. To achieve this, we built a quantitative admixture graph model (Fig. 3; Supplementary Note 3) that represents the four major ancestry flows contributing to present-day European genomes over the last 50,000 years ^20^. We used this model to simulate genomes at time periods and in sample sizes equivalent to our empirical dataset, and inferred tree sequences using Relate ^21,22^. We trained a neural network classifier to estimate the path backwards in time through the population structure taken by each simulated individual, at each position in the genome. Our trained classifier was then used to infer the ancestral paths taken at each site, using 1,015 imputed ancient genomes from West Eurasia which passed quality filters. Using simulations, we show that our novel chromosome painting method has an average accuracy of 94.6% for the four ancestral paths leading to present-day Europeans and is robust to model misspecification.

**Fig 3.**
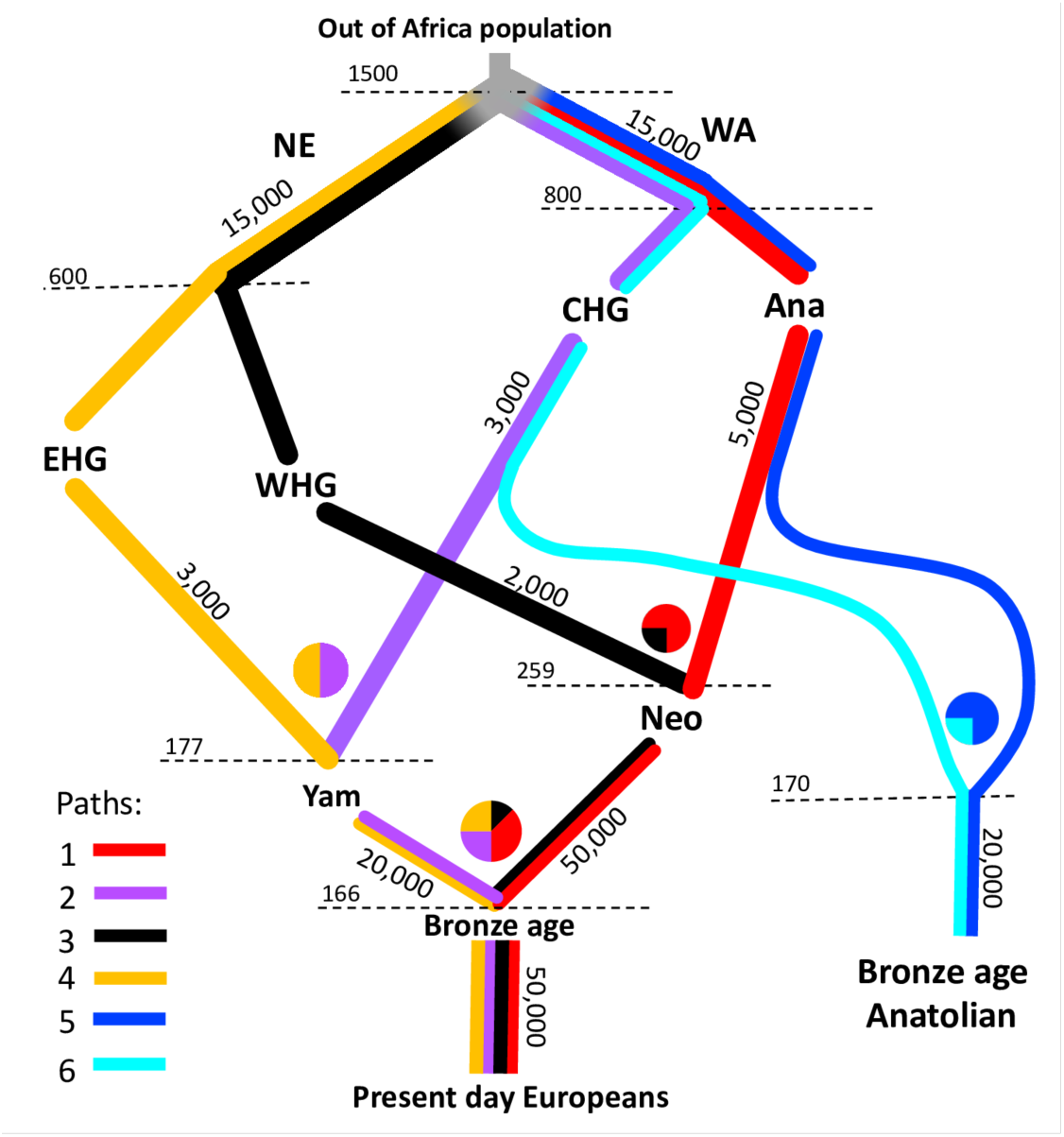
A schematic of the model of population structure in Europe. Quantitative admixture model used to simulate genomes to train the local ancestry neural network classifier. Moving down the figure is forwards in time and the population split times and admixture times are given in generations ago. Each branch is labelled with the effective population size of the population. Coloured lines represent the populations declared in the simulation that extend through time.

We then adapted CLUES ^23^ to model aDNA time-series data (Supplementary Notes 4 and 5) and used it to infer allele frequency trajectories and selection coefficients for 33,341 quality-controlled trait-associated variants from the GWAS Catalogue ^24^. An equal number of putatively neutral, frequency-paired variants were used as a control set (Supplementary Note 4). To control for possible confounders, we built a causal model to distinguish direct effects of age on allele frequency from indirect effects mediated by read depth, read length, and/or error rates (Supplementary Note 6), and developed a mapping bias test used to evaluate systematic differences between data from ancient and present-day populations (Supplementary Note 4). Because admixture between groups with differing allele frequencies can confound interpretation of allele frequency changes through time, we used the local ancestry paths from our novel chromosome painting model to stratify haplotypes in our selection tests. By conditioning on these path labels, we are able to infer selection trajectories while controlling for changes in admixture proportions through time.

Our analysis identified no genome-wide significant (p < 5e-8) selective sweeps when using genomes from present-day individuals alone (1000 Genomes Project populations GBR, FIN and TSI ^25^), although trait-associated variants were enriched for evidence of selection compared to the control group (p < 7.29e-35, Wilcoxon signed-rank test). In contrast, when using imputed aDNA genotype probabilities, we identified 11 genome-wide significant selective sweeps in the GWAS group (n=476 SNPs with p < 5e-8), and no sweeps in the control group, despite some SNPs exhibiting evidence of selection (n=51). These results are consistent with selection preferentially acting on trait-associated variants. We then conditioned our selection analysis on each of our four local ancestry pathways — i.e., local ancestry tracts passing through either Western hunter-gatherers (WHG), Eastern hunter-gatherers (EHG), Caucasus hunter-gatherers (CHG) or Anatolian farmers (ANA) — and identified 21 genome-wide significant selection peaks (Fig. 4 and Extended Data Figs. 1–10). This suggests that admixture between ancestral populations has masked evidence of selection at many trait-associated loci in Eurasian populations ^26^.

**Fig 4.**
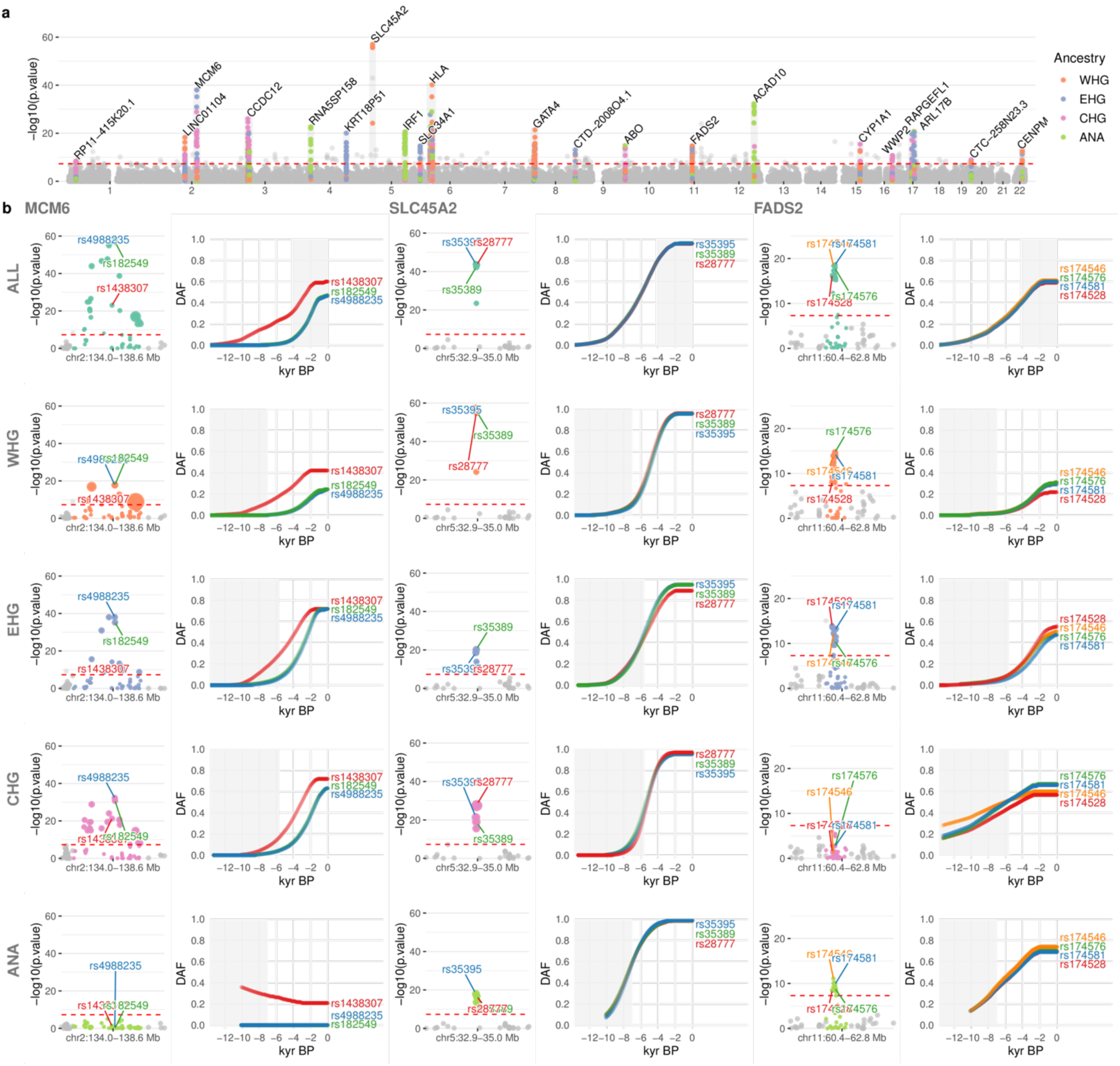
Genome-wide selection scan for trait-associated variants. **a**, Manhattan plot of p-values from selection scan with CLUES, based on a time-series of imputed aDNA genotype probabilities. Twenty-one genome-wide significant selection peaks highlighted in grey and labelled with the gene closest to the most significant SNP within each locus. Within each sweep, SNPs are positioned on the y-axis and coloured by their most significant marginal ancestry. Outside of the sweeps, SNPs show p-values from the pan-ancestry analysis and are coloured grey. Red dotted lines indicate genome-wide significance (p < 5e-8). **b**, Detailed plots for three genome-wide significant sweep loci: (i) MCM6, lactase persistence; (ii) SLC45A2, skin pigmentation; and (iii) FADS2, lipid metabolism. Rows show results for the pan-ancestry analysis (ALL) plus the four marginal ancestries: Western hunter-gatherers (WHG), Eastern hunter-gatherers (EHG), Caucasus hunter-gatherers (CHG) and Anatolian farmers (ANA). The first column of each loci shows zoomed Manhattan plots of the p-values for each ancestry, and column two shows allele frequency trajectories for the top SNPs across all ancestries (grey shading for the marginal ancestries indicates approximate temporal extent of the pre-admixture population).

### Selection on diet-associated loci

We find strong changes in selection associated with lactose digestion after the introduction of farming, but prior to the expansion of the Steppe pastoralists into Europe around 5,000 years ago ^27,28^, the timing of which is a long standing controversy ^29–32^. The strongest overall signal of selection in the pan-ancestry analysis is observed at the *MCM6 / LCT* locus (rs4988235:A; p=1.68e-59; s=0.0194), where the derived allele results in lactase persistence ^33^. The trajectory inferred from the pan-ancestry analysis indicates that the lactase persistence allele began increasing in frequency c. 6,000 years ago and has continued to increase up to present times (Fig. 4). In the ancestry-stratified analyses, this signal is driven primarily by sweeps in two of the ancestral backgrounds, associated with EHG and CHG. We also observed that many selected SNPs within this locus exhibited earlier evidence of selection than at rs4988235, suggesting that selection at the *MCM6/LCT* locus is more complex than previously thought. To investigate this further, we expanded our selection scan to include all SNPs within the ∼2.6 megabase (Mb) wide sweep locus (n=5,608) and checked for the earliest evidence of selection. We observed that the vast majority of genome-wide significant SNPs at this locus began rising in frequency earlier than rs4988235, indicating that strong positive selection at this locus predates the emergence of the lactase persistence allele by thousands of years. Among the alleles showing much earlier frequency rises was rs1438307:T (p=9.77e-24; s=0.0146), which began rising in frequency c. 12,000 years ago (Fig. 4). This allele has been shown to regulate energy expenditure and contribute to metabolic disease, and it has been hypothesised to be an ancient adaptation to famine ^34^. The high linkage disequilibrium between rs1438307 and rs4988235 in present-day individuals (R^2^ = 0.89 in 1000G GBR) may explain the recently observed correlation between frequency rises in the lactase persistence allele and archaeological proxies for famine and increased pathogen exposure ^35^. To control for potential bias introduced by imputation, we replicated these results using genotype likelihoods, called directly from the aDNA sequencing reads, and with publicly available 1240k capture array data from the Allen Ancient DNA Resource, version 52.2 ^36^ (Supplementary Note 4).

We also found strong selection in the FADS gene cluster — *FADS1* (rs174546:C; p=4.41e-19; s=0.0126) and *FADS2* (rs174581:G; p=2.21e-19; s=0.0138) — which are associated with fatty acid metabolism and known to respond to changes in diet from a more/less vegetarian to a more/less carnivorous diet ^37–41^. In contrast to previous results ^39–41^, we find that much of the selection associated with a more vegetarian diet occurred in Neolithic populations before they arrived in Europe, then continued during the Neolithic (Fig. 4). The strong signal of selection in this region in the pan-ancestry analysis is driven primarily by a sweep occurring across the EHG, WHG and ANA haplotypic backgrounds (Fig. 4). Interestingly, we do not find statistically significant evidence of selection at this locus in the CHG background, but most of the allele frequency rise in the EHG background occurs after their admixture with CHG (around 8 Kya ^42^), within whom the selected alleles were already close to present-day frequencies. This suggests that the selected alleles may already have existed at substantial frequencies in early farmer populations in the Middle East and among Caucasus Hunter-gatherers (associated with the ANA and CHG backgrounds, respectively) and were subject to continued selection as eastern groups moved northwards and westwards during the late Neolithic and Bronze Age periods.

When specifically comparing selection signatures differentiating ancient hunter-gatherer and farmer populations ^43^, we also observe a large number of regions associated with lipid and sugar metabolism, and various metabolic disorders (Supplementary Note 7). These include, for example, a region in chromosome 22 containing *PATZ1*, which regulates the expression of *FADS1*, and *MORC2*, which plays an important role in cellular lipid metabolism ^44^. Another region in chromosome 3 overlaps with *GPR15,* which is both related to immune tolerance and to intestinal homeostasis ^45,46^. Finally, in chromosome 18, we recover a selection candidate region spanning *SMAD7,* which is associated with inflammatory bowel diseases such as Crohn’s disease ^47^. Taken together these results suggest that the transition to agriculture imposed a substantial amount of selection for humans to adapt to a new diet and lifestyle, and that the prevalence of some diseases observed today may be a consequence of these selective processes.

### Selection on immunity-associated loci

We also observe evidence of strong selection in several loci associated with immunity and autoimmune disease (Supplementary Note 4). Some of these putative selection events occurred earlier than previously claimed and are likely associated with the transition to agriculture, which may help explain the high prevalence of autoimmune diseases today. Most notably, we detect an 8 Mb wide selection sweep signal in chromosome 6 (chr6:25.4-33.5 Mb), spanning the full length of the human leukocyte antigen (HLA) region. The selection trajectories of the variants within this locus support multiple independent sweeps, occurring at different times and with differing intensities. The strongest signal of selection at this locus in the pan-ancestry analysis is at an intergenic variant, located between *HLA-A* and *HLA-W* (rs7747253:A; p=7.56e-32; s=-0.0178), associated with protection against chickenpox (OR 0.888 ^5^), increased risk of intestinal infections (OR 1.08 ^48^) and decreased heel bone mineral density (OR 0.98 ^49^). This allele rapidly decreased in frequency, beginning c. 8,000 years ago (Extended Data Fig. 3), reducing risk of intestinal infections, at the cost of increasing risk of chickenpox. In contrast, the signal of selection at *C2* (rs9267677:C; p= 6.60e-26; s= 0.0441), also found within this sweep, shows a gradual increase in frequency beginning c. 4,000 years ago, before rising more rapidly c. 1,000 years ago. In this case, the favoured allele is associated with protection against some sexually transmitted diseases (STDs) (OR 0.786 ^48^), primarily those caused by human papillomavirus, and with increased psoriasis risk (OR 2.2 ^5^). This locus provides a good example of the hypothesis that the high prevalence of auto-immune diseases in present-day populations may, in part, be due to genetic trade-offs; by which selection increased protection against pathogens with the pleiotropic effect of increased susceptibility to auto-immune diseases ^50^.

These results also highlight the complex temporal dynamics of selection at the HLA locus, which not only plays a role in the regulation of the immune system but is also associated with many non-immune-related phenotypes. The high pleiotropy in this region makes it difficult to determine which selection pressures may have driven these increases in frequencies at different periods of time. However, profound shifts in lifestyle in Eurasian populations during the Holocene have been hypothesised to be drivers for strong selection on loci involved in immune response. These include a change in diet and closer contact with domestic animals, combined with higher mobility and increasing population density. We further explore the complex pattern of ancestry-specific selection at the HLA locus in our companion paper, “Elevated genetic risk for Multiple Sclerosis emerged in Steppe Pastoralist populations” ^51^.

We also identify selection signals at the *SLC22A4* (rs35260072:C; p=8.49e-20; s=0.0172) locus, associated with increased itch intensity from mosquito bites (OR 1.049 ^52^), protection against childhood and adult asthma (OR 0.902 and 0.909 ^48^) and asthma-related infections (OR 0.913 ^48^) and we find that the derived variant has been steadily rising in frequency since c. 9,000 years ago (Extended Data Fig. 9). However, in the same *SLC22A4* candidate region as rs35260072, we find that the frequency of the previously reported allele rs1050152:T (which also protects against asthma (OR 0.90 ^48^) and related infections) plateaued c. 1,500 years ago, contrary to previous reports suggesting a recent rise in frequency ^7^. Similarly, we detect selection at the *HECTD4* (rs11066188:A; p=9.51e-31 s=0.0198) and *ATXN2* (rs653178:C; p=3.73e-29; s=0.0189) loci, both of which have been rising in frequency for c. 9,000 years (Extended Data Fig. 4), also contrary to previous reports of a more recent rise in frequency ^7^. These SNPs are associated with protection against urethritis and urethral syndrome (OR 0.769 and 0.775 ^48^), which are often caused by STDs, or accumulated urethral damage from having more than 5 births. Both SNPs are also linked to increased risk of intestinal infectious diseases (OR 1.03 and 1.04), several non-specific parasitic diseases (OR 1.44 and 1.59 ^48^), schistosomiasis (OR 1.13 and 1.32 ^48^), helminthiases (OR 1.29 and 1.28 ^48^), spirochaetes (OR 1.14 and 1.12 ^48^), pneumonia (OR 1.03 and 1.03 ^48^) and viral hepatitis (OR 1.15 and 1.15 ^5^). These SNPs also increase the risk of celiac disease and rheumatoid arthritis ^53^. Thus, several highly pleiotropic disease-associated loci, which were previously thought to be the result of recent adaptation, may have been subject to selection for a much longer period of time.

### Selection on the 17q21.31 locus

We further detect signs of strong selection in a 2 Mb sweep on chromosome 17 (chr17:44.0-46.0 Mb), spanning a locus on 17q21.3, implicated in neurodegenerative and developmental disorders. The locus includes an inversion and other structural polymorphisms with indications of a recent positive selection sweep in some human populations ^54,55^. Specifically, partial duplications of the *KANSL1* gene likely occurred independently on the inverted (H2) and non-inverted (H1) haplotypes (Extended Data Fig. 11a) and both are found in high frequencies (15-25%) among current European and Middle Eastern populations but are much rarer in Sub-Saharan African and East Asian populations. We used both SNP genotypes and WGS read depth information to determine inversion (H1/H2) and *KANSL1* duplication (d) status in the ancient individuals studied here (Supplementary Note 8).

The H2 haplotype is observed in two of three previously published genomes ^56^ of Anatolian aceramic-associated Neolithic individuals (Bon001 and Bon004) from around 10,000 BP, but data were insufficient to identify *KANSL1* duplications. The oldest evidence for *KANSL1* duplications is observed in an early Neolithic individual (AH1 from 9,900 BP ^57^) from present-day Iran, followed by two Mesolithic individuals (NEO281 from 9,724 BP and KK1 ^58^ from 9,720 BP), from present-day Georgia, all of whom are heterozygous for the inversion and carry the inverted duplication. The *KANSL1* duplications are also detected in two Neolithic individuals, from present-day Russia (NEO560 from 7,919 BP (H1d) and NEO212 from 7,390 BP (H2d)). With both H1d and H2d having spread to large parts of Europe with Anatolian Neolithic farmer ancestries, their frequency seems unchanged in most of Europe as Steppe-related ancestries become dominant in large parts of the subcontinent (Extended Data Fig. 11c). The fact that both H1d and H2d are found in apparently high frequencies in both early Anatolian farmers and the earliest Steppe-related ancestry groups suggests that any selective sweep acting on the H1d and H2d variants would probably have occurred in populations ancestral to both.

We note that the strongest signal of selection observed in the pan-ancestry analysis at this locus is at *MAPT* (rs4792897:G; p=1.33e-18; s=0.0299 (Extended Data Fig. 8; Supplementary Note 4), which codes for the tau protein ^59^, and is associated with protection against mumps (OR 0.776 ^48^) and increased risk of snoring (OR 1.04 ^60^). More generally, polymorphisms in *MAPT* have been associated with increased risk of a number of neurodegenerative disorders, including Alzheimer’s disease and Parkinson’s disease ^61^. However, we caution that this region is also enriched for evidence of reference bias in our dataset—especially around the *KANSL1* gene—due to complex structural polymorphisms (Supplementary Note 10).

### Selection on pigmentation loci

Our results identify strong selection for lighter skin pigmentation in groups moving northwards and westwards, consistent with the hypothesis that selection is caused by reduced UV exposure and resulting vitamin D deficiency. We find that the most strongly selected alleles reached near-fixation several thousand years ago, suggesting that this process was not associated with recent sexual selection as previously proposed ^62^. In the pan-ancestry analysis we detect strong selection at the *SLC45A2* locus (rs35395:C; p=1.60e-44; s=0.0215) ^8,63^, with the selected allele (responsible for lighter skin), increasing in frequency from c. 13,000 years ago, until plateauing c. 2,000 years ago (Fig. 4). The predominant hypothesis is that high melanin levels in the skin are important in equatorial regions owing to its protection against UV radiation, whereas lighter skin has been selected for at higher latitudes (where UV radiation is less intense) because some UV penetration is required for cutaneous synthesis of vitamin D ^64,65^. Our findings confirm pigmentation alleles as major targets of selection during the Holocene ^7,66^ particularly on a small proportion of loci with large effect sizes ^8^.

Additionally, our results provide detailed information about the duration and geographic spread of these processes (Fig. 4) suggesting that an allele associated with lighter skin was selected for repeatedly, probably as a consequence of similar environmental pressures occurring at different times in different regions. In the ancestry-stratified analysis, all marginal ancestries show broad agreement at the *SLC45A2* locus (Fig. 4) but differ in the timing of their frequency shifts. The ANA-associated ancestry background shows the earliest evidence for selection at rs35395, followed by EHG and WHG around c. 10,000 years ago, and CHG c. 2,000 years later. In all ancestry backgrounds except ANA, the selected haplotypes plateau at high frequency by c. 2,000 years ago, whilst the ANA haplotype background reaches near fixation 1,000 years earlier. We also detect strong selection at the *SLC24A5* locus (rs1426654:A; p=2.28e-16; s=0.0185) which is also associated with skin pigmentation ^63,67^. At this locus, the selected allele increased in frequency even earlier than *SLC45A2* and reached near fixation c. 3,500 years ago. Selection on this locus thus seems to have occurred early on in groups that were moving northwards and westwards, and only later in the Western hunter-gatherer background after these groups encountered and admixed with the incoming populations.

### Selection among major axes of variation

Beyond patterns of genetic change at the Mesolithic-Neolithic transition, much genetic variability observed today reflects high genetic differentiation in the hunter-gatherer groups that eventually contributed to present-day European genetic diversity ^43^. Indeed, a substantial number of loci associated with cardiovascular disease, metabolism and lifestyle diseases trace their genetic variability prior to the Neolithic transition, to ancient differential selection in ancestry groups occupying different parts of the Eurasian continent (Supplementary Note 7). These may represent selection episodes that preceded the admixture events described above and led to differentiation between ancient hunter-gatherer groups in the late Pleistocene and early Holocene. One of these overlaps with the *SLC24A3* gene which is a salt sensitivity gene significantly expressed in obese individuals ^68^. Another spans *ROPN1* and *KALRN*, two genes involved in vascular disorders ^69^. A further region contains *SLC35F3*, which codes for a thiamine transport ^70^ and has been associated with hypertension in a Han Chinese cohort ^71^. Finally, there is a candidate region containing several genes (*CH25H, FAS*) associated with obesity and lipid metabolism ^72,73^ and another peak with several genes (*ASXL2, RAB10, HADHA, GPR113*) involved in glucose homeostasis and fatty acid metabolism ^74–77^. These loci predominantly reflect ancient patterns of extreme differentiation between Eastern and Western Eurasian genomes and may be candidates for selection after the separation of the Pleistocene populations that occupied different environments across the continent (roughly 45,000 years ago ^13^).

### Pathogenic structural variants

Rare, recurrent copy-number variants (CNVs) are known to cause neurodevelopmental disorders and are associated with a range of psychiatric and physical traits with variable expressivity and incomplete penetrance ^78,79^. To understand the prevalence of pathogenic structural variants over time we examined 50 genomic regions susceptible to recurrent CNVs, known to be the most prevalent drivers of human developmental pathologies ^80^. The analysis included 1442 ancient shotgun genomes passing quality control for CNV analysis (Supplementary Note 10) and 1093 present-day human genomes for comparison ^81,82^. We identified CNVs in ancient individuals at ten loci using a read-depth based approach and digital Comparative Genomic Hybridization ^83^. Although most of the observed CNVs (including duplications at 15q11.2 and *CHRNA7*, and CNVs spanning parts of the TAR locus and 22q11.2 distal) have not been unambiguously associated with disease in large studies, the identified CNVs include deletions and duplications that have been associated with developmental delay, dysmorphic features, and neuropsychiatric abnormalities such as autism (most notably at 1q21.1, 3q29, 16p12.1 and the DiGeorge/VCFS locus, but also deletions at 15q11.2 and duplications at 16p13.11). Overall, the carrier frequency in the ancient individuals is similar to that reported in the UK Biobank genomes (1.25% vs 1.6% at 15q11.2 and *CHRNA7* combined, and 0.8% vs 1.1% across the remaining loci combined) ^84^. These results suggest that large, recurrent CNVs that can lead to several pathologies were present at similar frequencies in the ancient and present-day populations included in this study.

### Phenotypic legacy of ancient Eurasians

In addition to identifying evidence of selection for trait-associated variants, we also estimated the contribution from different genetic ancestries (associated with EHG, CHG, WHG, Steppe pastoralists and Neolithic farmers) to variation in complex traits in present-day individuals. We calculated ancestry-specific polygenic risk scores — hereafter ancestral risk scores (ARS) — based on chromosome painting of >400,000 UKB genomes using ChromoPainter ^85^ (Fig. 5, Supplementary Note 9). This allowed us to identify which ancient ancestry components are over-represented in present-day UK populations at loci significantly associated with a given trait and is analogous to the genetic risk that a present-day individual would possess if they were composed entirely of one of the ancestry groupings defined in this study. This analysis avoids issues related to the portability of polygenic risk scores between populations ^86^, as our ancestral risk scores are calculated from the same individuals used to estimate the effect sizes. Working with large numbers of imputed ancient genomes provides high statistical power to use ancient populations as ancestral sources. We focused on 35 phenotypes whose polygenic scores were significantly over-dispersed among the ancient populations (Supplementary Note 9), as well as well as three large effect alleles at the *APOE* gene (ApoE2, ApoE3, and ApoE4) known to significantly mediate risk of developing Alzheimer’s disease ^87^. We emphasise that this approach makes no direct reference to ancient phenotypes, but instead describes how these genetic ancestry components contributed to the present-day phenotypic landscape.

**Fig. 5.**
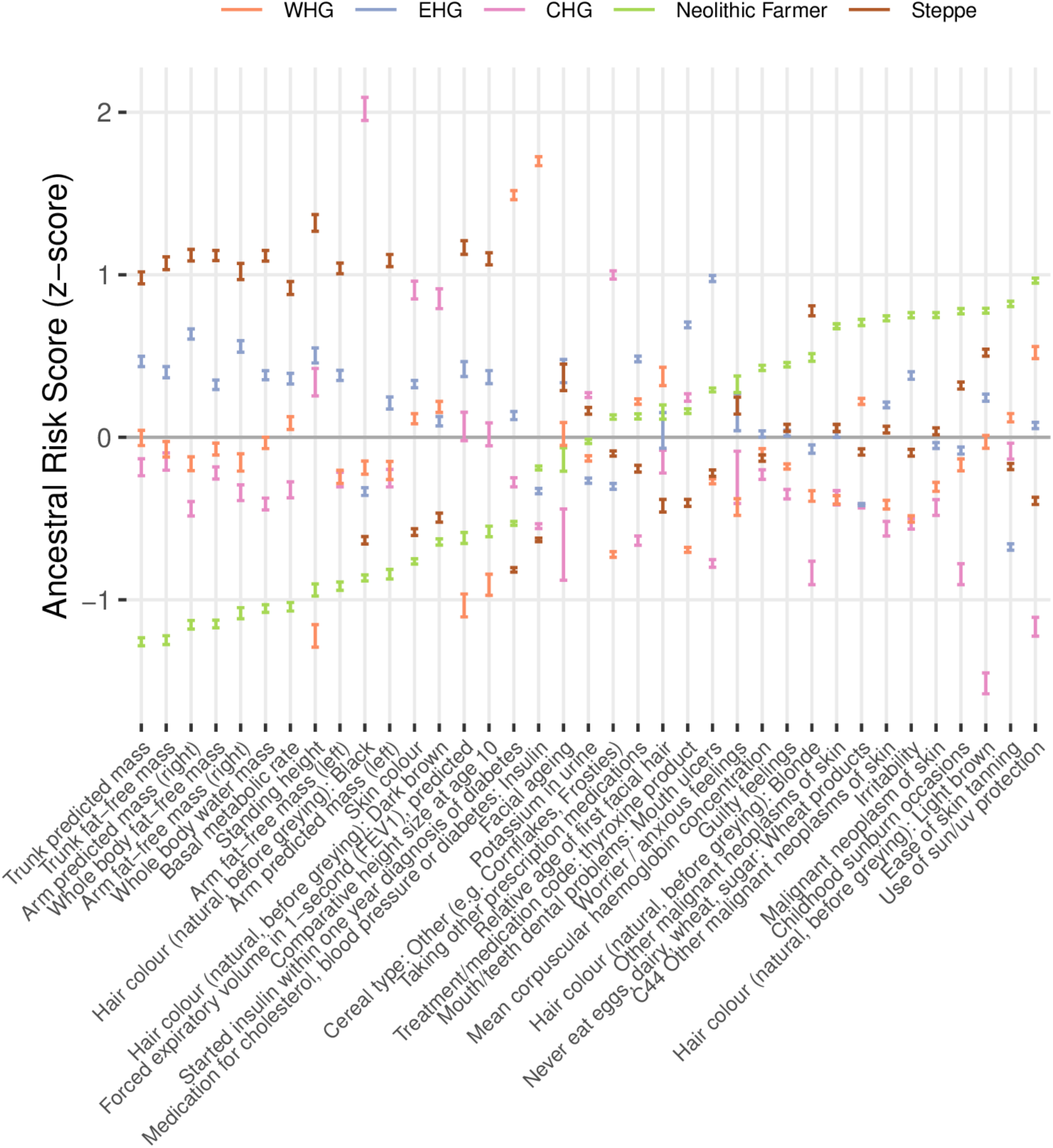
Ancestral risk scores (ARS) for 35 complex traits. Showing the genetic risk that a present-day individual would possess if they were composed entirely of one ancestry. Based on chromosome painting of the UK Biobank, for 35 complex traits found to be significantly over-dispersed in ancient populations. Confidence intervals (95%) are estimated by bootstrapping present-day samples (n=408,884) and centred on the mean estimate.

We find that for many anthropometric traits—like trunk predicted mass, forced expiratory volume in 1-second (FEV1), and basal metabolic rate—the ARS for Steppe ancestry was the highest, followed by EHG and CHG/WHG, whilst Neolithic farmer ancestry consistently scored the lowest for these measurements. Consistent with previous studies, hair and skin pigmentation also showed significant differences, with scores for skin colour for WHG, EHG and CHG higher (i.e. darker) than for Neolithic farmer and Steppe-associated ancestries ^8,9,27,28^; and scores for traits related to malignant neoplasms of the skin were elevated in Neolithic farmer-associated ancestries. Both Neolithic farmer and Steppe-associated ancestries have higher scores for blonde and light brown hair, while the Hunter-Gatherer-associated ancestries have higher scores for dark brown hair, and CHG-associated ancestries had the highest score for black hair.

In terms of genetic contributions to risk for diseases, the WHG ancestral component had strikingly high scores for traits related to cholesterol, blood pressure and diabetes. The Neolithic farmer component scored the highest for anxiety, guilty feelings, and irritability; CHG and WHG ancestry components consistently scored the lowest for these three traits. We found the ApoE4 allele (rs429358:C and rs7412:C, which increases risk of Alzheimer’s disease) preferentially painted with a WHG/EHG haplotypic background, suggesting it was likely brought into Western Eurasia by early hunter-gatherers (Supplementary Note 9). This result is in line with the present-day European distribution of this allele, which is highest in north-eastern Europe, where the proportion of these ancestries is larger than in other regions of the continent ^88^. In contrast, we found the ApoE2 allele (rs429358:T and rs7412:T, which decreases risk for Alzheimer’s disease) on a haplotypic background with affinities to Steppe pastoralists. Our pan-ancestry analysis identified positive selection favouring ApoE2 (p=6.99e-3; s=0.0130), beginning c. 7,000 years ago and plateauing c. 2,500 years ago (Supplementary Note 4). However, we did not identify evidence of selection for either ApoE3 (rs429358:T and rs7412:C) or ApoE4, contrary to a recent study with a smaller sample size and unphased genotypes ^89^. The selective forces likely favouring ApoE2 in Steppe pastoralists may be associated with protective immune responses against infectious challenges, such as protection against malaria or an unknown viral infection (Supplementary Note 9).

In light of the ancestry gradients within the United Kingdom and across Eurasia (Fig. 2), these results support the hypothesis that migration-mediated geographic variation in phenotypes and disease risk is commonplace, and points to a way forward for explaining geographically structured disease prevalence through differential admixture processes between present-day populations. These results also help to clarify the famous discussion of selection in Europe relating to height ^7,90^. Our finding that the Steppe and EHG associated ancestral components have elevated genetic values for height in the UK Biobank demonstrates that height differences between Northern and Southern Europe may be a consequence of differential ancestry, rather than selection, as claimed in many previous studies ^91^. However, our results do not preclude the possibility that height has been selected for in specific populations ^92,93^.

## Discussion

The fundamental changes in diet resulting from the transitions from hunting and gathering to farming, and subsequently to pastoralism, precipitated far-reaching consequences for the physical and mental health of present-day Eurasian populations. These dramatic cultural changes created a heterogeneous mix of selection pressures, likely related to changes in diet and increased population densities, including selection for resistance to novel infectious challenges. Due to the highly pleiotropic nature of each sweep region, it is difficult to ascribe causal factors to any of our selection signals, and we did not exhaustively test all non-trait associated variants. However, our results show that selection during the Holocene has had a substantial impact on present-day genetic disease risk, as well as the distribution of genetic factors affecting metabolic and anthropometric traits. Our analyses have also shown that the ability to detect signatures of natural selection in present-day human genomes is drastically limited by conflicting selection pressures in different ancestral populations masking the signals. Developing methods to trace selection while accounting for differential admixture allowed us to effectively double the number of genome-wide significant selection peaks and helped clarify the trajectories of a number of variants related to diet and lifestyle. Furthermore, we have shown that numerous complex traits thought to have been under local selection are better explained by differing genetic contributions of ancient individuals to present-day variation. Overall, our results emphasise how the interplay between ancient selection and major admixture events occurring in the Mesolithic, Neolithic and Bronze Age have profoundly shaped the patterns of genetic variation observed in present-day humans across Eurasia.

## Data availability

All ancient genomic data used in this study are already published and listed in Supplementary Table 1. Data was aligned to the human reference GRCh37. Modern human genomes were obtained from the 1000 Genomes Project (1KGP) ^25^, the Simons Genome Diversity Project (SGDP) ^81^, and the Human Genome Diversity Project (HGDP) ^82^. GWAS data was obtained from the GWAS Catalog ^24^, the FinnGen Study ^48^, and the UK Biobank (UKB)^5^.

## Code availability

The scripts used to run the chromosome painting (Supplementary Note 2) and calculate ARS in the UK Biobank (Supplementary Note 9) are available at https://github.com/will-camb/mesoneo_selection_paper (https://doi.org/10.5281/zenodo.8301166). The software to perform the ancestral path chromosome painting described in Supplementary Note 3 is available on GitHub at https://github.com/AliPearson/AncestralPaths (https://doi.org/10.5281/zenodo.8319452), and the demographic model is available in the *stdpopsim* library (see https://popsim-consortium.github.io/stdpopsim-docs/stable/catalog.html#sec_catalog_homsap_models_ancienteurope_4a21). The analysis pipeline and ‘conda’ environment necessary to replicate the analysis of allele frequency trajectories of trait-associated variants in Supplementary Note 4 are available at https://github.com/ekirving/mesoneo_paper (https://doi.org/10.5281/zenodo.8289755). The modified version of CLUES used in this study is available from https://github.com/standard-aaron/clues (https://doi.org/10.5281/zenodo.8228252). The pipeline to replicate the analyses for Supplementary Note 7 can be found at https://github.com/albarema/neo (https://doi.org/10.5281/zenodo.8301253). All other analyses relied upon available software which has been fully referenced in the manuscript and detailed in the relevant supplementary notes.

## Acknowledgements

We thank all the former and current staff at the Lundbeck Foundation GeoGenetics Centre and the GeoGenetics Sequencing Core and colleagues across the many institutions detailed below. We are particularly grateful to L. Olsen as project manager for the Lundbeck Foundation GeoGenetics Centre project. We thank UKB for access to the UKB genomic resource. We want to acknowledge the participants and investigators of the FinnGen study. We are thankful to Illumina for collaboration. E.W. thanks St. John’s College, Cambridge, for providing a stimulating environment of discussion and learning.

The Lundbeck Foundation GeoGenetics Centre is supported by the Lundbeck Foundation (R302-2018-2155 and R155-2013-16338), the Novo Nordisk Foundation (NNF18SA0035006), the Wellcome Trust (214300), Carlsberg Foundation (CF18-0024), the Danish National Research Foundation (DNRF94 and DNRF174), the University of Copenhagen (KU2016 programme), Ferring Pharmaceuticals A/S, and a COREX ERC Synergy grant (ID 951385). This research has been conducted using the UKB Resource and the iPSYCH Initiative, funded by the Lundbeck Foundation (R102-A9118 and R155-2014-1724). This work was further supported by the Swedish Foundation for Humanities and Social Sciences grant (Riksbankens Jubileumsfond M16-0455:1) to K.K.. E.K.I.-P. and A.R.-M. were supported by the Lundbeck Foundation (R302-2018-2155) and the Novo Nordisk Foundation (NNF18SA0035006). A.P., R.D. and E.W. were supported by the Wellcome Trust (214300). R.M. was supported by a SSHRC doctoral studentship (G101449). R.A.H. and T.K. were supported by the Carlsberg Foundation (CF19-0712). L.S. was supported by a Sir Henry Wellcome fellowship (220457/Z/20/Z). L.F. was supported by the OAK Foundation (OCAY-15-520). P.H.S. was supported by the Institute of General Medical Sciences (R35GM142916) and a Vallee Scholars Award. R.N. was supported by the National Institutes of Health (R01GM138634). F.R. was supported by a Villum Young Investigator Grant (project no. 00025300), a Novo Nordisk Fonden Data Science Ascending Investigator Award (NNF22OC0076816)), and by the European Research Council (ERC) under the European Union’s Horizon Europe programme (grant agreements No. 101077592 and 951385).

## Author Information

These authors contributed equally: Evan K. Irving-Pease, Alba Refoyo-Martínez, William Barrie, Andrés Ingason, Alice Pearson, and Anders Fischer.

These authors equally supervised research: Peter H. Sudmant, Daniel J. Lawson, Richard Durbin, Thorfinn Korneliussen, Thomas Werge, Morten E. Allentoft, Martin Sikora, Rasmus Nielsen, Fernando Racimo, Eske Willerslev

## Contributions

E.K.I-P, A.R-M, W.B., A.I., A.P.., and A.F. contributed equally to this work. P.H.S., D.J.L., R.D., T.S.K., T.W., M.E.A., M.S., R.N., F.R., and E.W. led the study. A.F., T.W., M.E.A., M.S., and E.W. conceptualised the study. P.H.S., D.J.L., R.D., T.S.K., T.W., M.E.A., M.S., R.N., F.R., and E.W. supervised the research. M.E.A., K.K., R.D., T.W., R.N. and E.W. acquired funding for research. E.K.I-P, A.R-M, A.I., A.P., W.B., A.V., L.S., A.J. Stern, K.K., D.J.L., R.D., T.S.K.. M.E.A., M.S., R.N., and F.R. were involved in developing and applying methodology. E.K.I-P, A.R-M, A.I., A.P., W.B., A.S.H., R.A.H, T.V., H.M., A.V., L.S., A. Ramsøe, A.J. Schork, A. Rosengren, L.Z., P.H.S., T.S.K., M.E.A., M.S., and F.R undertook formal analyses of data. E.K.I-P, A.R-M, A.I., A.P., A.F., W.B., K.G.S., A.S.H., R.A.H, T.V., A.J. Stern, A. Ramsøe, A. Rosengren, L.Z., A.K.N.I., L.F., P.H.S., D.J.L., T.S.K., M.S., F.R. and E.W. drafted the main text. E.K.I-P, A.R-M, A.I., A.P., A.F., W.B., K.G.S., A.S.H., R.A.H, T.V., A.J. Stern, G.S., A. Ramsøe, A. Rosengren, L.Z., A.K.N.I., L.F., P.H.S., D.J.L., M.S., and E.W. drafted supplementary notes and materials. E.K.I-P, A.R-M, A.I., A.P., A.F., W.B., K.G.S., A.S.H., R.M., F.D., R.A.H, T.V., H.M., A. Ramsøe, A.J. Schork, L.Z., K.K., A.K.N.I., L.F., P.H.S., D.J.L., R.D., T.S.K., T.W., M.E.A., M.S., R.N., F.R., and E.W. were involved in reviewing drafts and editing. All co-authors read, commented on, and agreed upon the submitted manuscript.

## Extended data figures

**Extended Data Fig. 1.**
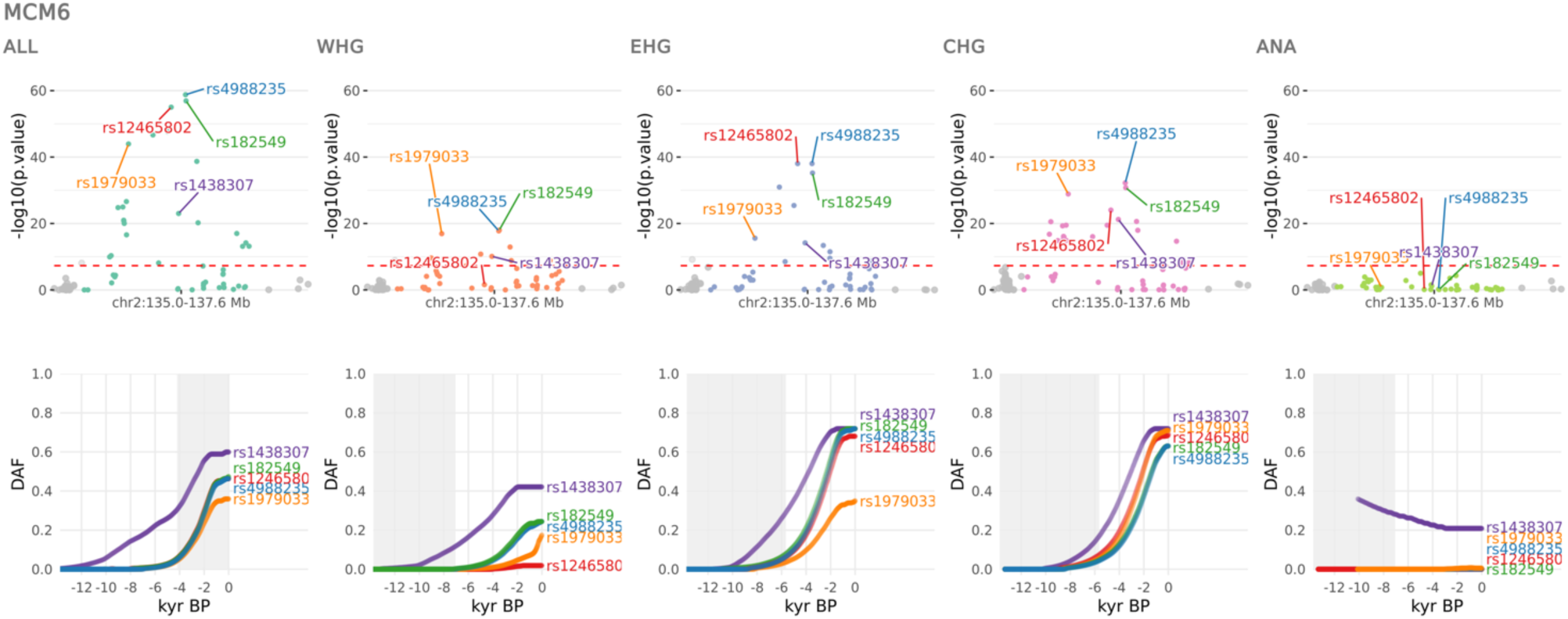
Selection at the MCM6 locus. CLUES selection results for the most significant sweep locus, showing the pan-ancestry analysis (ALL) plus the four marginal ancestries: Western hunter-gatherers (WHG), Eastern hunter-gatherers (EHG), Caucasus hunter-gatherers (CHG) and Anatolian farmers (ANA). Row one shows zoomed Manhattan plots of the p-values for each ancestry, and row two shows allele trajectories for the top SNPs across all ancestries (grey shading for the marginal ancestries indicates approximate temporal extent of the pre-admixture population).

**Extended Data Fig. 2.**
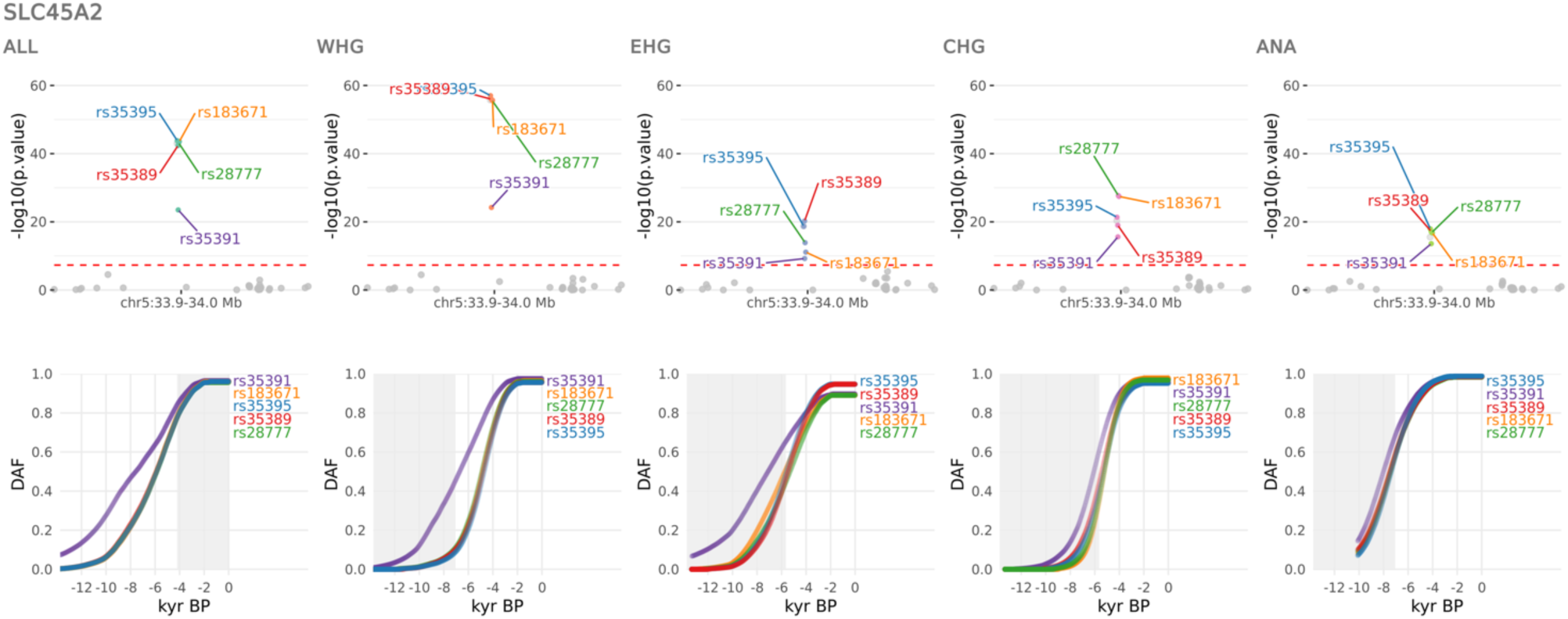
Selection at the SLC45A2 locus. CLUES selection results for the second most significant sweep locus, showing the pan-ancestry analysis (ALL) plus the four marginal ancestries: Western hunter-gatherers (WHG), Eastern hunter-gatherers (EHG), Caucasus hunter-gatherers (CHG) and Anatolian farmers (ANA). Row one shows zoomed Manhattan plots of the p-values for each ancestry, and row two shows allele trajectories for the top SNPs across all ancestries (grey shading for the marginal ancestries indicates approximate temporal extent of the pre-admixture population).

**Extended Data Fig. 3.**
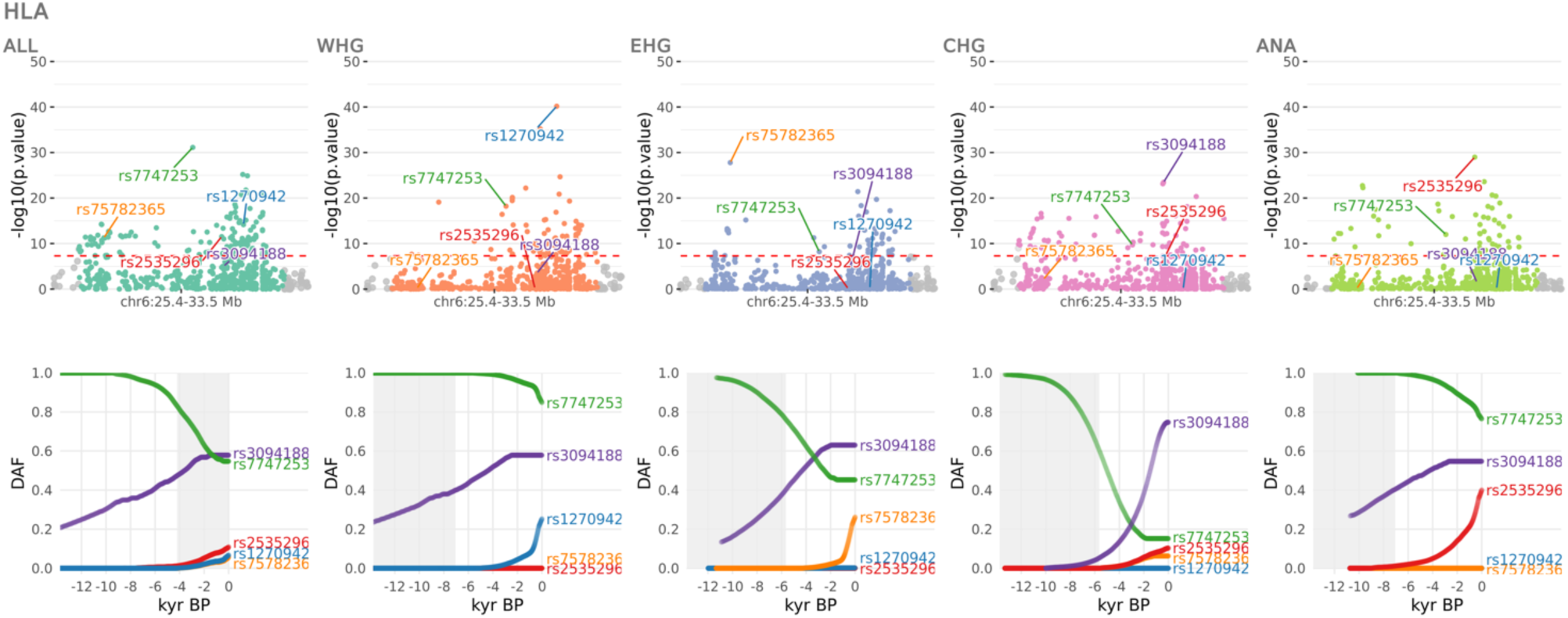
Selection at the HLA locus. CLUES selection results for the third most significant sweep locus, showing the pan-ancestry analysis (ALL) plus the four marginal ancestries: Western hunter-gatherers (WHG), Eastern hunter-gatherers (EHG), Caucasus hunter-gatherers (CHG) and Anatolian farmers (ANA). Row one shows zoomed Manhattan plots of the p-values for each ancestry, and row two shows allele trajectories for the top SNPs across all ancestries (grey shading for the marginal ancestries indicates approximate temporal extent of the pre-admixture population).

**Extended Data Fig. 4.**
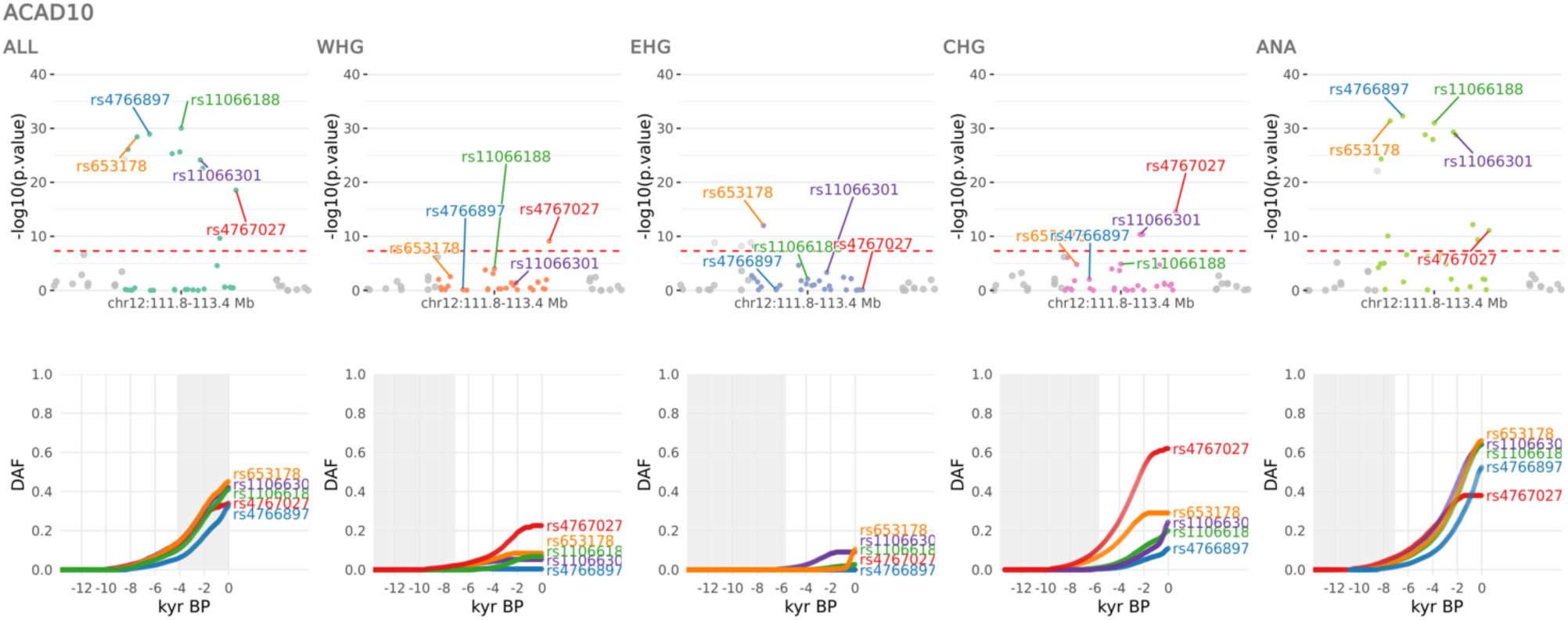
Selection at the ACAD10 locus. CLUES selection results for the fourth most significant sweep locus, showing the pan-ancestry analysis (ALL) plus the four marginal ancestries: Western hunter-gatherers (WHG), Eastern hunter-gatherers (EHG), Caucasus hunter-gatherers (CHG) and Anatolian farmers (ANA). Row one shows zoomed Manhattan plots of the p-values for each ancestry, and row two shows allele trajectories for the top SNPs across all ancestries (grey shading for the marginal ancestries indicates approximate temporal extent of the pre-admixture population).

**Extended Data Fig. 5.**
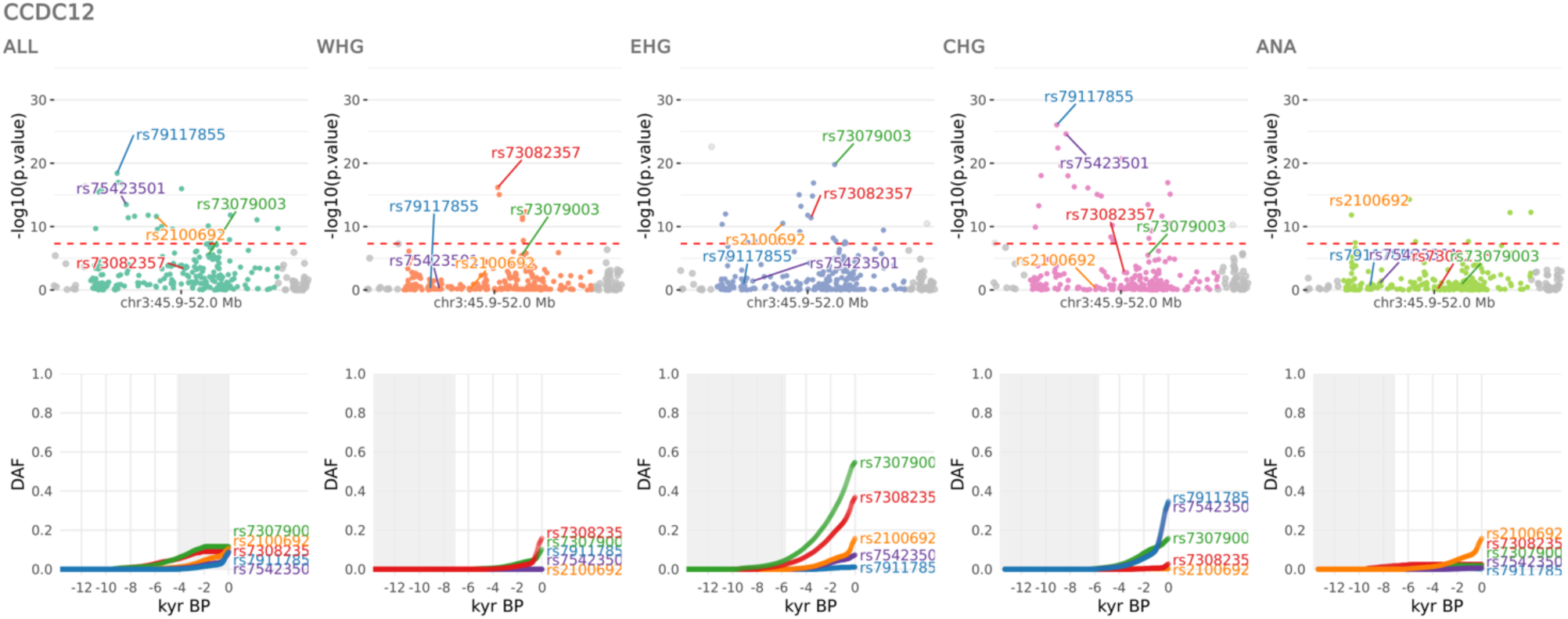
Selection at the CCDC12 locus. CLUES selection results for the fifth most significant sweep locus, showing the pan-ancestry analysis (ALL) plus the four marginal ancestries: Western hunter-gatherers (WHG), Eastern hunter-gatherers (EHG), Caucasus hunter-gatherers (CHG) and Anatolian farmers (ANA). Row one shows zoomed Manhattan plots of the p-values for each ancestry, and row two shows allele trajectories for the top SNPs across all ancestries (grey shading for the marginal ancestries indicates approximate temporal extent of the pre-admixture population).

**Extended Data Fig. 6.**
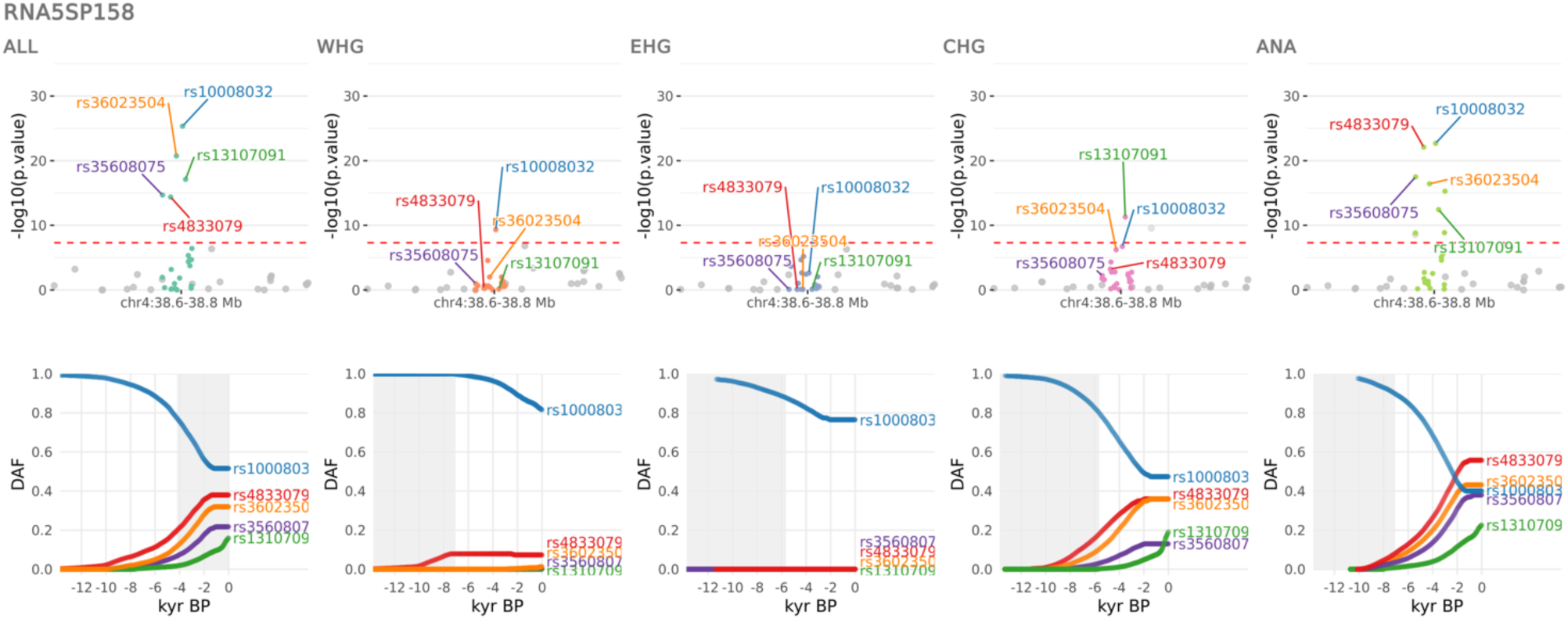
Selection at the RNA5SP158 locus. CLUES selection results for the sixth most significant sweep locus, showing the pan-ancestry analysis (ALL) plus the four marginal ancestries: Western hunter-gatherers (WHG), Eastern hunter-gatherers (EHG), Caucasus hunter-gatherers (CHG) and Anatolian farmers (ANA). Row one shows zoomed Manhattan plots of the p-values for each ancestry, and row two shows allele trajectories for the top SNPs across all ancestries (grey shading for the marginal ancestries indicates approximate temporal extent of the pre-admixture population).

**Extended Data Fig. 7.**
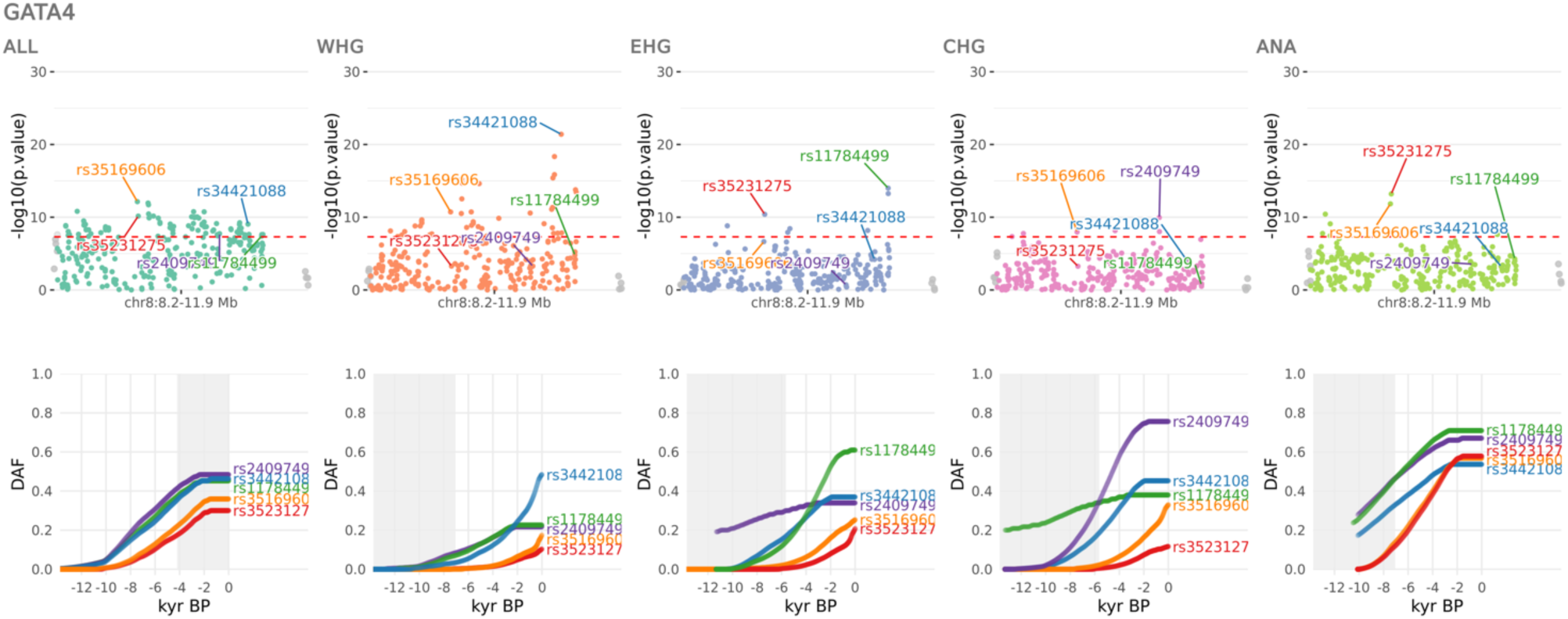
Selection at the GATA4 locus. CLUES selection results for the seventh most significant sweep locus, showing the pan-ancestry analysis (ALL) plus the four marginal ancestries: Western hunter-gatherers (WHG), Eastern hunter-gatherers (EHG), Caucasus hunter-gatherers (CHG) and Anatolian farmers (ANA). Row one shows zoomed Manhattan plots of the p-values for each ancestry, and row two shows allele trajectories for the top SNPs across all ancestries (grey shading for the marginal ancestries indicates approximate temporal extent of the pre-admixture population).

**Extended Data Fig. 8.**
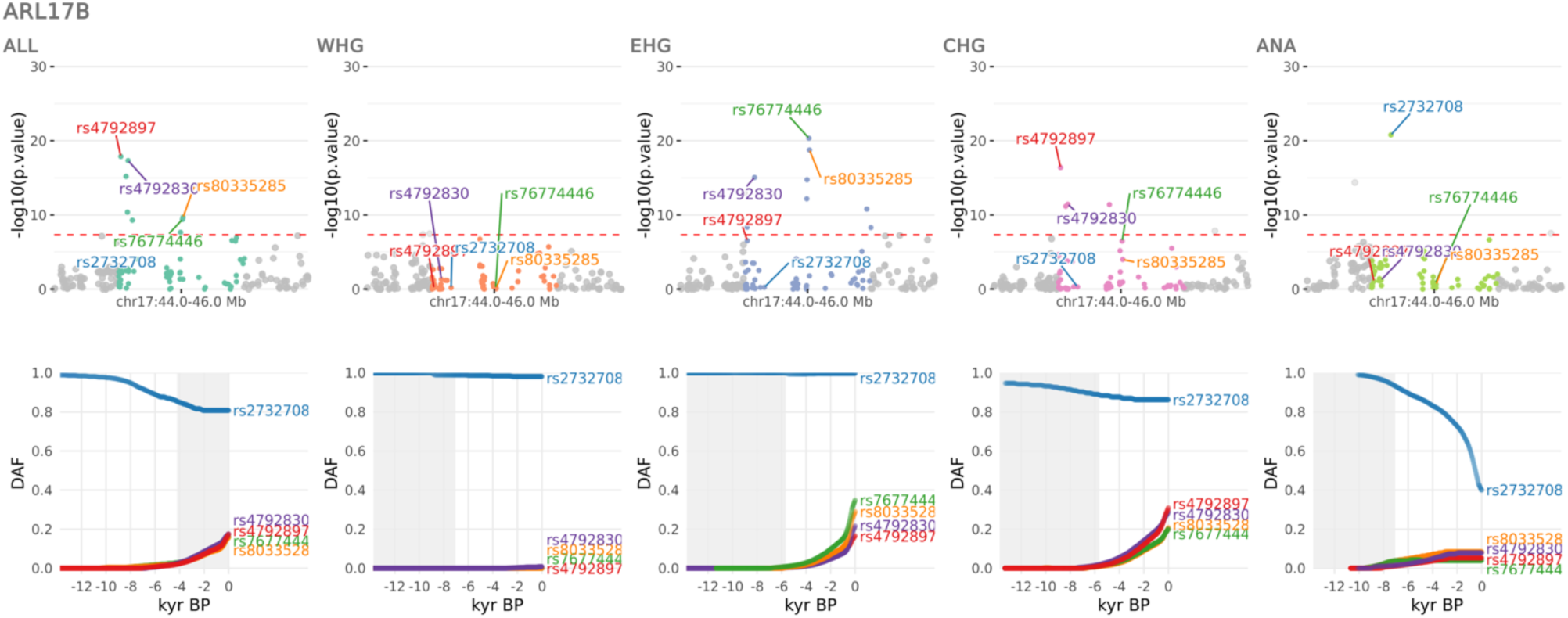
Selection at the ARL17B locus. CLUES selection results for the eight most significant sweep locus, showing the pan-ancestry analysis (ALL) plus the four marginal ancestries: Western hunter-gatherers (WHG), Eastern hunter-gatherers (EHG), Caucasus hunter-gatherers (CHG) and Anatolian farmers (ANA). Row one shows zoomed Manhattan plots of the p-values for each ancestry, and row two shows allele trajectories for the top SNPs across all ancestries (grey shading for the marginal ancestries indicates approximate temporal extent of the pre-admixture population).

**Extended Data Fig. 9.**
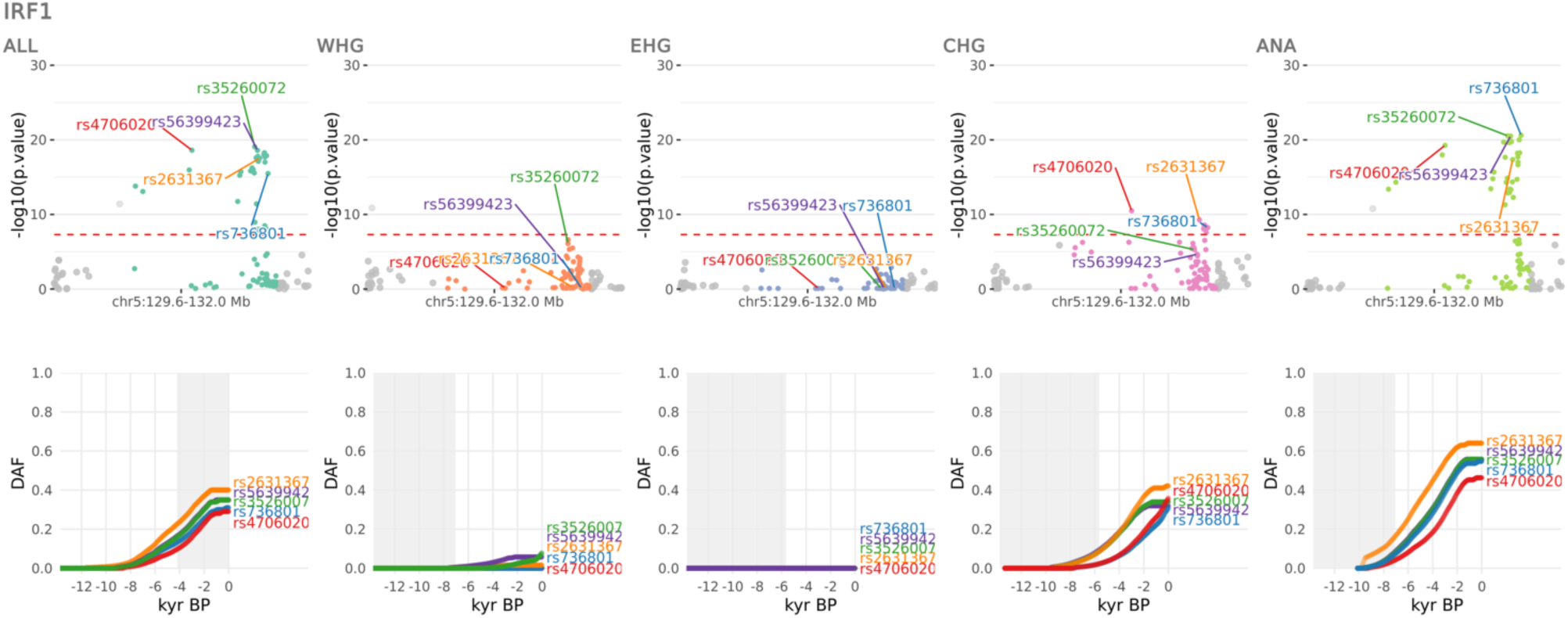
Selection at the IRF1 locus. CLUES selection results for the ninth most significant sweep locus, showing the pan-ancestry analysis (ALL) plus the four marginal ancestries: Western hunter-gatherers (WHG), Eastern hunter-gatherers (EHG), Caucasus hunter-gatherers (CHG) and Anatolian farmers (ANA). Row one shows zoomed Manhattan plots of the p-values for each ancestry, and row two shows allele trajectories for the top SNPs across all ancestries (grey shading for the marginal ancestries indicates approximate temporal extent of the pre-admixture population).

**Extended Data Fig. 10.**
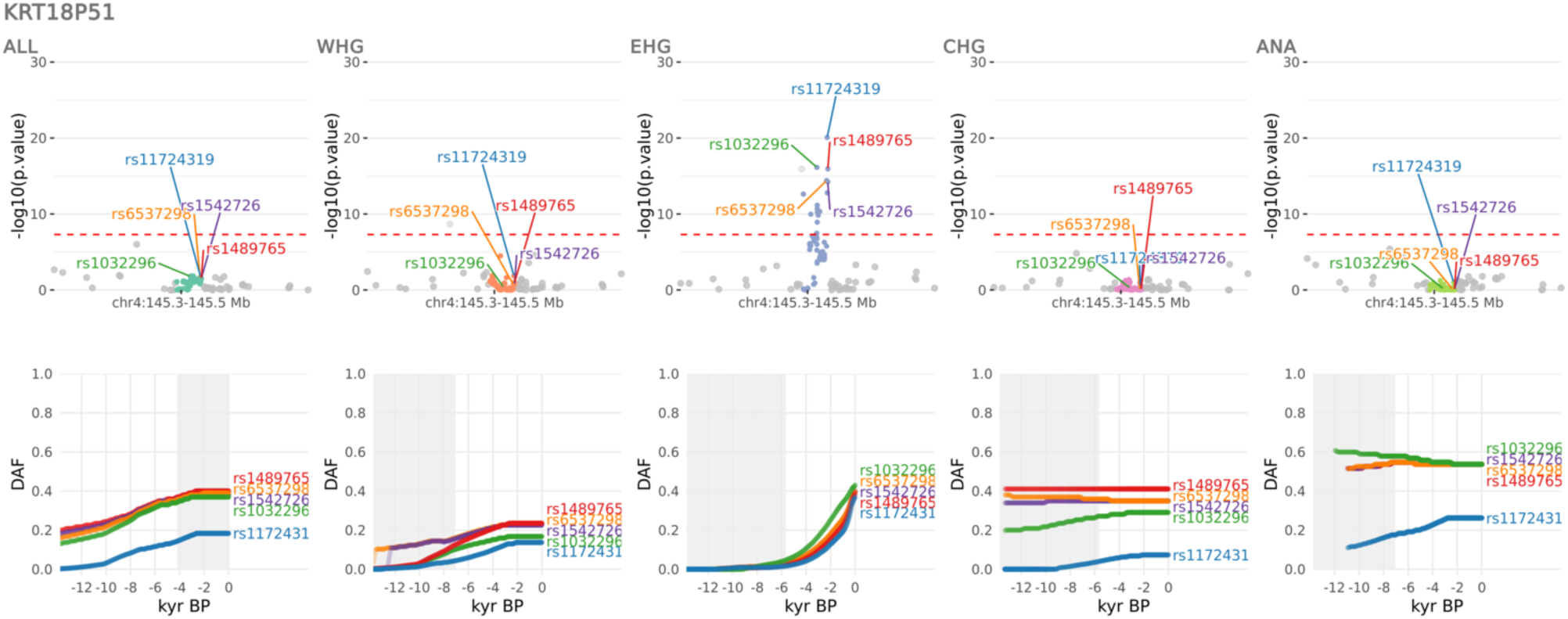
Selection at the KRT18P51 locus. CLUES selection results for the tenth most significant sweep locus, showing the pan-ancestry analysis (ALL) plus the four marginal ancestries: Western hunter-gatherers (WHG), Eastern hunter-gatherers (EHG), Caucasus hunter-gatherers (CHG) and Anatolian farmers (ANA). Row one shows zoomed Manhattan plots of the p-values for each ancestry, and row two shows allele trajectories for the top SNPs across all ancestries (grey shading for the marginal ancestries indicates approximate temporal extent of the pre-admixture population).

**Extended Data Fig. 11.**
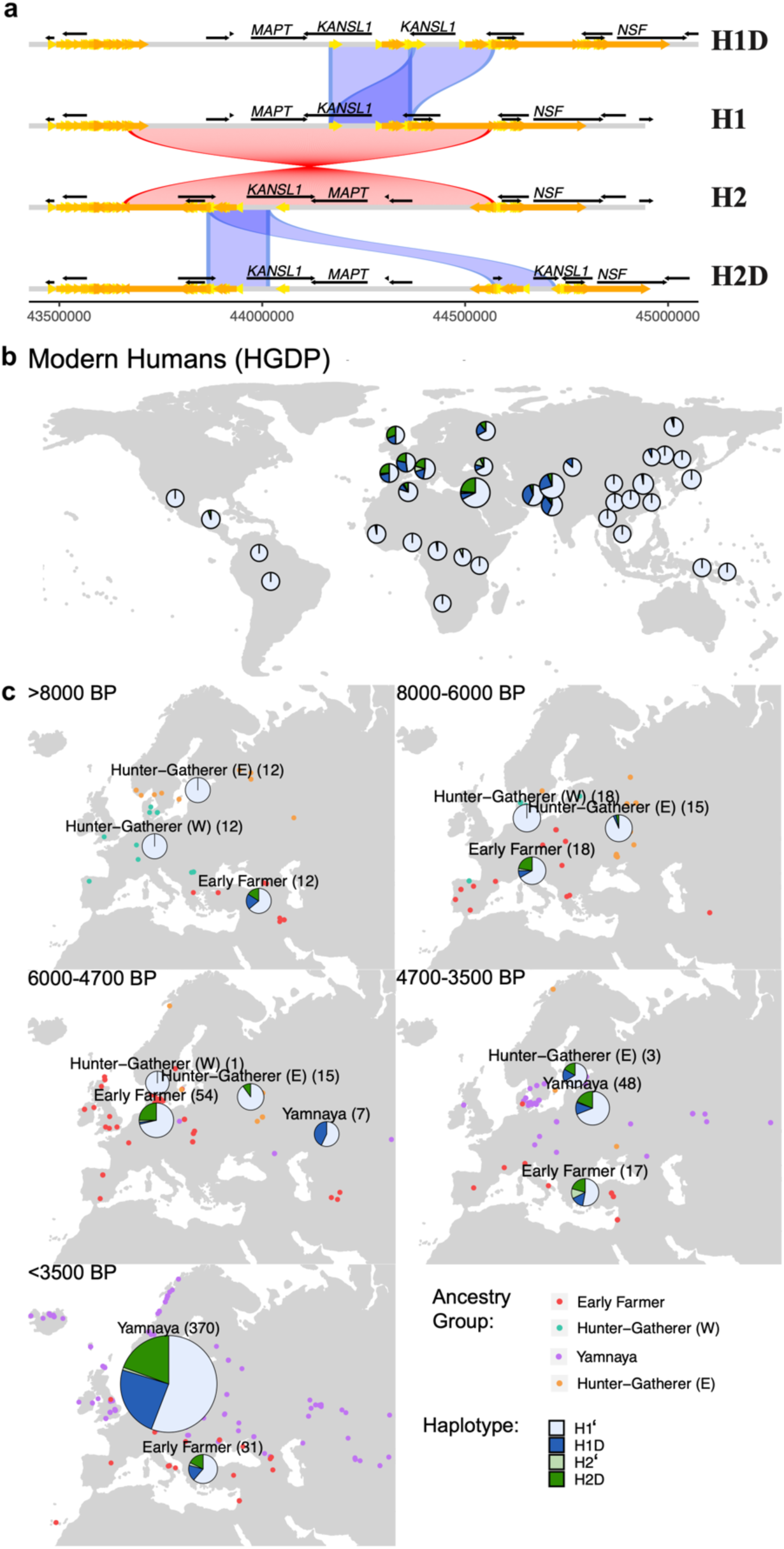
The 17q21.31 inversion locus. A) Haplotypes of the 17q21.31 locus: the ancestral (non-inverted) H1 17q21.31 and the inverted H2 haplotype. Duplications of the *KANSL1* gene have occurred independently on both lineages yielding H1D and H2D haplotypes. B) Frequency of the 17q21.31 inversion and duplication haplotypes across present-day global populations (Human Genome Diversity Project ^82^). D) Change in the frequency of the 17q21.31 inversion haplotype through time.

## References

1. Allentoft, M. E. et al. Population Genomics of Stone Age Eurasia. bioRxiv 2022.05.04.490594 (2022) doi:10.1101/2022.05.04.490594.

2. Page, A. E. et al. Reproductive trade-offs in extant hunter-gatherers suggest adaptive mechanism for the Neolithic expansion. Proc. Natl. Acad. Sci. U. S. A. 113, 4694–4699 (2016).

3. Marciniak, S., Bergey, C., Silva, A. M. & Hałuszko, A. An integrative skeletal and paleogenomic analysis of prehistoric stature variation suggests relatively reduced health for early European farmers. bioRxiv (2021).

4. Visscher, P. M. et al. 10 Years of GWAS Discovery: Biology, Function, and Translation. Am. J. Hum. Genet. 101, 5–22 (2017).

5. Bycroft, C. et al. The UK Biobank resource with deep phenotyping and genomic data. Nature 562, 203–209 (2018).

6. Vitti, J. J., Grossman, S. R. & Sabeti, P. C. Detecting natural selection in genomic data. Annu. Rev. Genet. 47, 97–120 (2013).

7. Mathieson, I. et al. Genome-wide patterns of selection in 230 ancient Eurasians. Nature 528, 499– 503 (2015).

8. Ju, D. & Mathieson, I. The evolution of skin pigmentation-associated variation in West Eurasia. Proc. Natl. Acad. Sci. U. S. A. 118, (2021).

9. Wilde, S. et al. Direct evidence for positive selection of skin, hair, and eye pigmentation in Europeans during the last 5,000 y. Proc. Natl. Acad. Sci. U. S. A. 111, 4832–4837 (2014).

10. Sousa da Mota, B., et al. Imputation of ancient human genomes. Nat. Commun. 14, 3660 (2023).

11. Lawson, D. J., Hellenthal, G., Myers, S. & Falush, D. Inference of population structure using dense haplotype data. PLoS Genet. 8, e1002453 (2012).

12. Hellenthal, G. et al. A genetic atlas of human admixture history. Science 343, 747–751 (2014).

13. Sikora, M. et al. The population history of northeastern Siberia since the Pleistocene. Nature 570, 182–188 (2019).

14. Shinde, V. et al. An Ancient Harappan Genome Lacks Ancestry from Steppe Pastoralists or Iranian Farmers. Cell 179, 729–735.e10 (2019).

15. Hofmanová, Z. et al. Early farmers from across Europe directly descended from Neolithic Aegeans. Proc. Natl. Acad. Sci. U. S. A. 113, 6886–6891 (2016).

16. Galinsky, K. J., Loh, P.-R., Mallick, S., Patterson, N. J. & Price, A. L. Population Structure of UK Biobank and Ancient Eurasians Reveals Adaptation at Genes Influencing Blood Pressure. Am. J. Hum. Genet. 99, 1130–1139 (2016).

17. Patterson, N. et al. Large-scale migration into Britain during the Middle to Late Bronze Age. Nature (2022) doi:10.1038/s41586-021-04287-4.

18. Olalde, I. et al. The Beaker phenomenon and the genomic transformation of northwest Europe. Nature 555, 190–196 (2018).

19. Zaidi, A. A. & Mathieson, I. Demographic history mediates the effect of stratification on polygenic scores. Elife 9, e61548 (2020).

20. Jones, E. R. et al. Upper Palaeolithic genomes reveal deep roots of modern Eurasians. Nat. Commun. 6, 8912 (2015).

21. Speidel, L., Forest, M., Shi, S. & Myers, S. R. A method for genome-wide genealogy estimation for thousands of samples. Nat. Genet. 51, 1321–1329 (2019).

22. Speidel, L. et al. Inferring Population Histories for Ancient Genomes Using Genome-Wide Genealogies. Mol. Biol. Evol. 38, 3497–3511 (2021).

23. Stern, A. J., Wilton, P. R. & Nielsen, R. An approximate full-likelihood method for inferring selection and allele frequency trajectories from DNA sequence data. PLoS Genet. 15, e1008384 (2019).

24. Buniello, A. et al. The NHGRI-EBI GWAS Catalog of published genome-wide association studies, targeted arrays and summary statistics 2019. Nucleic Acids Res. 47, D1005–D1012 (2019).

25. 1000 Genomes Project Consortium et al. A global reference for human genetic variation. Nature 526, 68–74 (2015).

26. Souilmi, Y. et al. Admixture has obscured signals of historical hard sweeps in humans. Nat Ecol Evol 6, 2003–2015 (2022).

27. Allentoft, M. E. et al. Population genomics of Bronze Age Eurasia. Nature 522, 167–172 (2015).

28. Haak, W. et al. Massive migration from the steppe was a source for Indo-European languages in Europe. Nature 522, 207–211 (2015).

29. Enattah, N. S. et al. Independent introduction of two lactase-persistence alleles into human populations reflects different history of adaptation to milk culture. Am. J. Hum. Genet. 82, 57–72 (2008).

30. Itan, Y., Powell, A., Beaumont, M. A., Burger, J. & Thomas, M. G. The origins of lactase persistence in Europe. PLoS Comput. Biol. 5, e1000491 (2009).

31. Ségurel, L. & Bon, C. On the Evolution of Lactase Persistence in Humans. Annu. Rev. Genomics Hum. Genet. 18, 297–319 (2017).

32. Segurel, L. et al. Why and when was lactase persistence selected for? Insights from Central Asian herders and ancient DNA. PLoS Biol. 18, e3000742 (2020).

33. Enattah, N. S. et al. Identification of a variant associated with adult-type hypolactasia. Nat. Genet. 30, 233–237 (2002).

34. Wang, L. et al. A MicroRNA Linking Human Positive Selection and Metabolic Disorders. Cell 183, 684–701.e14 (2020).

35. Evershed, R. P. et al. Dairying, diseases and the evolution of lactase persistence in Europe. Nature 608, 336–345 (2022).

36. Mallick, S., et al. The Allen Ancient DNA Resource (AADR): A curated compendium of ancient human genomes. bioRxiv 2023.04.06.535797 (2023).

37. Willer, C. J. et al. Discovery and refinement of loci associated with lipid levels. Nat. Genet. 45, 1274–1283 (2013).

38. Gallois, A. et al. A comprehensive study of metabolite genetics reveals strong pleiotropy and heterogeneity across time and context. Nat. Commun. 10, 4788 (2019).

39. Buckley, M. T. et al. Selection in Europeans on Fatty Acid Desaturases Associated with Dietary Changes. Mol. Biol. Evol. 34, 1307–1318 (2017).

40. Ye, K., Gao, F., Wang, D., Bar-Yosef, O. & Keinan, A. Dietary adaptation of FADS genes in Europe varied across time and geography. Nat Ecol Evol 1, 167 (2017).

41. Mathieson, S. & Mathieson, I. FADS1 and the Timing of Human Adaptation to Agriculture. Mol. Biol. Evol. 35, 2957–2970 (2018).

42. Lazaridis, I. The evolutionary history of human populations in Europe. Curr. Opin. Genet. Dev. 53, 21–27 (2018).

43. Luu, K., Bazin, E. & Blum, M. G. B. pcadapt: an R package to perform genome scans for selection based on principal component analysis. Mol. Ecol. Resour. 17, 67–77 (2017).

44. Sánchez-Solana, B., Li, D.-Q. & Kumar, R. Cytosolic functions of MORC2 in lipogenesis and adipogenesis. Biochim. Biophys. Acta 1843, 316–326 (2014).

45. Kim, S. V. et al. GPR15-mediated homing controls immune homeostasis in the large intestine mucosa. Science 340, 1456–1459 (2013).

46. Nguyen, L. P., et al. Role and species-specific expression of colon T cell homing receptor GPR15 in colitis. Nature Immunology vol. 16 207–213 Preprint at 10.1038/ni.3079 (2015).

47. Monteleone, G. et al. Mongersen, an oral SMAD7 antisense oligonucleotide, and Crohn’s disease. N. Engl. J. Med. 372, 1104–1113 (2015).

48. Kurki, M. I. et al. FinnGen provides genetic insights from a well-phenotyped isolated population. Nature 613, 508–518 (2023).

49. Morris, J. A. et al. An atlas of genetic influences on osteoporosis in humans and mice. Nat. Genet. 51, 258–266 (2018).

50. Brinkworth, J. F. & Barreiro, L. B. The contribution of natural selection to present-day susceptibility to chronic inflammatory and autoimmune disease. Curr. Opin. Immunol. 31, 66–78 (2014).

51. Barrie, W. et al. Genetic risk for Multiple Sclerosis originated in Pastoralist Steppe populations. bioRxiv 2022.09.23.509097 (2022) doi:10.1101/2022.09.23.509097.

52. Jones, A. V. et al. GWAS of self-reported mosquito bite size, itch intensity and attractiveness to mosquitoes implicates immune-related predisposition loci. Hum. Mol. Genet. 26, 1391–1406 (2017).

53. Gutierrez-Achury, J. et al. Functional implications of disease-specific variants in loci jointly associated with coeliac disease and rheumatoid arthritis. Hum. Mol. Genet. 25, 180–190 (2016).

54. Stefansson, H. et al. A common inversion under selection in Europeans. Nat. Genet. 37, 129–137 (2005).

55. Steinberg, K. M. et al. Structural diversity and African origin of the 17q21.31 inversion polymorphism. Nat. Genet. 44, 872–880 (2012).

56. Kılınç, G. M. et al. The Demographic Development of the First Farmers in Anatolia. Curr. Biol. 26, 2659–2666 (2016).

57. Broushaki, F. et al. Early Neolithic genomes from the eastern Fertile Crescent. Science 353, 499– 503 (2016).

58. Jones, E. R. et al. The Neolithic Transition in the Baltic Was Not Driven by Admixture with Early European Farmers. Curr. Biol. 27, 576–582 (2017).

59. Andreadis, A., Brown, W. M. & Kosik, K. S. Structure and novel exons of the human tau gene. Biochemistry 31, 10626–10633 (1992).

60. Jansen, P. R. et al. Genome-wide analysis of insomnia in 1,331,010 individuals identifies new risk loci and functional pathways. Nat. Genet. 51, 394–403 (2019).

61. Desikan, R. S. et al. Genetic overlap between Alzheimer’s disease and Parkinson’s disease at the MAPT locus. Mol. Psychiatry 20, 1588–1595 (2015).

62. Aoki, K. Sexual selection as a cause of human skin colour variation: Darwin’s hypothesis revisited. Ann. Hum. Biol. 29, 589–608 (2002).

63. Lona-Durazo, F. et al. Meta-analysis of GWA studies provides new insights on the genetic architecture of skin pigmentation in recently admixed populations. BMC Genet. 20, 59 (2019).

64. Jablonski, N. G. & Chaplin, G. The evolution of human skin coloration. J. Hum. Evol. 39, 57–106 (2000).

65. Engelsen, O. The relationship between ultraviolet radiation exposure and vitamin D status. Nutrients 2, 482–495 (2010).

66. Voight, B. F., Kudaravalli, S., Wen, X. & Pritchard, J. K. A Map of Recent Positive Selection in the Human Genome. PLoS Biol. 4, e72 (2006).

67. Martin, A. R. et al. An Unexpectedly Complex Architecture for Skin Pigmentation in Africans. Cell 171, 1340–1353.e14 (2017).

68. Wu, H. et al. Transcriptome Sequencing to Detect the Potential Role of Long Noncoding RNAs in Salt-Sensitive Hypertensive Rats. Biomed Res. Int. 2019, 2816959 (2019).

69. Wang, L. et al. Peakwide Mapping on Chromosome 3q13 Identifies the Kalirin Gene as a Novel Candidate Gene for Coronary Artery Disease. The American Journal of Human Genetics vol. 80 650– 663 Preprint at 10.1086/512981 (2007).

70. Zhang, K. et al. Genetic implication of a novel thiamine transporter in human hypertension. J. Am. Coll. Cardiol. 63, 1542–1555 (2014).

71. Zang, X.-L. et al. Association of a SNP in SLC35F3 Gene with the Risk of Hypertension in a Chinese Han Population. Frontiers in Genetics vol. 7 Preprint at 10.3389/fgene.2016.00108 (2016).

72. Russo, L. et al. Cholesterol 25-hydroxylase (CH25H) as a promoter of adipose tissue inflammation in obesity and diabetes. Mol Metab 39, 100983 (2020).

73. Demir, A., Kahraman, R., Candan, G. & Ergen, A. The role of FAS gene variants in inflammatory bowel disease. Turk. J. Gastroenterol. 31, 356–361 (2020).

74. Izawa, T. et al. ASXL2 Regulates Glucose, Lipid, and Skeletal Homeostasis. Cell Rep. 11, 1625– 1637 (2015).

75. Vazirani, R. P. et al. Disruption of Adipose Rab10-Dependent Insulin Signaling Causes Hepatic Insulin Resistance. Diabetes 65, 1577–1589 (2016).

76. Thapa, D. et al. The protein acetylase GCN5L1 modulates hepatic fatty acid oxidation activity via acetylation of the mitochondrial β-oxidation enzyme HADHA. J. Biol. Chem. 293, 17676–17684 (2018).

77. Ong, H. S. & Yim, H. C. H. Microbial Factors in Inflammatory Diseases and Cancers. Regulation of Inflammatory Signaling in Health and Disease 153–174 Preprint at 10.1007/978-981-10-5987-2_7 (2017).

78. Girirajan, S., Campbell, C. D. & Eichler, E. E. Human copy number variation and complex genetic disease. Annu. Rev. Genet. 45, 203–226 (2011).

79. Weise, A. et al. Microdeletion and microduplication syndromes. J. Histochem. Cytochem. 60, 346– 358 (2012).

80. Girirajan, S. et al. Phenotypic heterogeneity of genomic disorders and rare copy-number variants. N. Engl. J. Med. 367, 1321–1331 (2012).

81. Mallick, S. et al. The Simons Genome Diversity Project: 300 genomes from 142 diverse populations. Nature 538, 201–206 (2016).

82. Bergström, A. et al. Insights into human genetic variation and population history from 929 diverse genomes. Science 367, (2020).

83. Sudmant, P. H. et al. Diversity of human copy number variation and multicopy genes. Science 330, 641–646 (2010).

84. Crawford, K. et al. Medical consequences of pathogenic CNVs in adults: analysis of the UK Biobank. J. Med. Genet. 56, 131–138 (2019).

85. Lawson, D. J., Hellenthal, G., Myers, S. & Falush, D. Inference of population structure using dense haplotype data. PLoS Genet. 8, e1002453 (2012).

86. Martin, A. R. et al. Human Demographic History Impacts Genetic Risk Prediction across Diverse Populations. Am. J. Hum. Genet. 100, 635–649 (2017).

87. Corder, E. H. et al. Gene dose of apolipoprotein E type 4 allele and the risk of Alzheimer’s disease in late onset families. Science 261, 921–923 (1993).

88. Belloy, M. E., Napolioni, V. & Greicius, M. D. A Quarter Century of APOE and Alzheimer’s Disease: Progress to Date and the Path Forward. Neuron 101, 820–838 (2019).

89. Kolbe, D. et al. Current allele distribution of the human longevity gene APOE in Europe can mainly be explained by ancient admixture. Aging Cell e13819 (2023).

90. Rosenstock, E. et al. Human stature in the Near East and Europe ca. 10,000–1000 BC: its spatiotemporal development in a Bayesian errors-in-variables model. Archaeol. Anthropol. Sci. 11, 5657–5690 (2019).

91. Field, Y. et al. Detection of human adaptation during the past 2000 years. Science 354, 760–764 (2016).

92. Chen, M. et al. Evidence of Polygenic Adaptation in Sardinia at Height-Associated Loci Ascertained from the Biobank Japan. Am. J. Hum. Genet. 107, 60–71 (2020).

93. Howe, L. J. et al. Within-sibship genome-wide association analyses decrease bias in estimates of direct genetic effects. Nat. Genet. 54, 581–592 (2022).

