## Supplementary Information for "The Selection Landscape and Genetic Legacy of Ancient Eurasians"


1) Selecting non-British individuals from the UK Biobank 5
- Introduction 5
- Methods 5
- Results 6
- Discussion 6
- References 6

2) Painting the UK Biobank 8
- Introduction 8
- Methods 8
  - Painting pipeline introduction 8
  - Reference/donor panel formation 8
  - Target/recipient panel formation 9
  - SNP selection and merging of the panels 9
  - Painting process 10
  - Painting at biobank scale 11
- Results 11
  - Ancestry-PCs relationship 11
  - Ancestry-geographic variation 12
- Discussion 13
- Figures 14
- References 18

3) Local Ancestry Inference for Complex Population Histories 20

Abstract 20
Introduction 20
Method and Materials 23
Concept of path ancestry 23
Constructing a path model of European population history 25
Comparison of simulated data to MesoNeo genomes 27
A method for estimating local path ancestry 30
Training and testing a neural network on simulated European genomes 32
Comparison to an advanced LAI tool 34
Results 35
Estimation of time since admixture in Europe 35
Discussion 41
Supplementary Information 43
3.1 Excluding Bronze Age Anatolians 43
3.2 Conditions affecting classifier performance 44
3.3 Testing model misspecification 52
3.4 A method to estimate time since admixture 58
References 68
4) Estimating allele frequency trajectories of trait-associated variants 72
Introduction 72
Methods 73
SNP Ascertainment 73
1000G ARG 74
Modifications to CLUES 74
Selection Analysis 75
Results 80
Selection in 1000G EUR 80
Selection in aDNA Time Series 81
Selection in simulations with Ancestral Paintings 103
Selection in aDNA with Ancestral Paintings 105

Selection at the LCT/MCM6 locus 138
Selection at the APOE locus 143
Discussion 145
Pan-ancestry selection 146
Ancestry stratified selection trajectories 148
Robustness of analysis 149
References 152
5) Validating CLUES on Ancient Genotypes 180
Introduction 180
Validation of $\chi^2_1$ Test Statistic 180
Estimation of Selection Coefficients 181
Effect of Demography and Sampling Density on Selection Coefficient Estimation 182
Estimation of Allele Frequencies 184
6) Detangling Direct and Indirect impacts of sample age from the Mesolithic-Neolithic data on genotype imputation 187
Introduction 187
Methods 187
Discussion 193
References 194
7) Identifying candidates for positive selection using patterns of ancient population differentiation 195
Introduction 195
Methods 195
Results 196
Eurasian scan 196
West Eurasian scan 197
Neolithic vs. hunter-gatherer scan 199
Figures 201
References 209
8) Calling chr17q21.31 KANSL1 duplications in ancient genomes 220
Introduction 220
Methods 220

Results 222
References 222
9) Calculating ancestral contributions to modern complex phenotypes 224
Introduction 224
Methods 225
Trait ascertainment 225
Calculating ancestral risk scores 226
Results 227
Discussion 228
ApoE2 and its association with protection against severe malaria 229
ApoE isoforms and hepatitis C infection 232
ApoE isoforms and herpes simplex type 1 infection 233
ApoE isoforms and coronavirus infection 233
Figures 235
References 240
10) Pathogenic structural variants in ancient vs. modern-day humans 246
Introduction 246
Methods 246
Results 247
References 271

### 1) Selecting non-British individuals from the UK Biobank

William Barrie[1] and Dan Lawson[2]

[1] Zoology Department, University of Cambridge, UK.
[2] School of Mathematics and Integrative Epidemiology Unit, University of Bristol, UK.

#### Introduction

The UK Biobank (UKB) contains approximately 40,000 individuals not born in the UK. Because many of these individuals are recently admixed or of British ancestry, we set up a pipeline to (1) exclude genetically British-like individuals and (2) select individuals of a typical genetic ancestral background for each country, in order to investigate the genetic contribution of each ancient ancestry to modern European, Asian and African populations.

#### Methods

For individuals from each country in Europe, Asia and Africa but not the UK, (Data-Field 1647: Country of birth (UK/elsewhere) and Data-Field 20115: Country of Birth (non-UK origin)), we ran two density-based scans using Scikit-learn (0.21.2)'s[1] DBSCAN method (Density-Based Spatial Clustering of Applications with Noise) on a distance matrix constructed using the first 18 PCs[2], weighted by their Eigenvalues. This algorithm finds cores of high density within a distance matrix, which can be any shape, and can include nearby non-core points. The eps parameter can be adjusted to determine how strict the clustering is. This is preferable to using visual PC cut-offs, for which it is difficult to include higher PCs; and k-means clustering, which assumes clusters are convex and all points must be clustered.

In the first scan, designed to remove individuals with British-like ancestry who were born abroad, individuals from a given country were combined with 8,000 random white British individuals, and the clustering algorithm was run on the combined data. Any individuals born abroad who clustered with the white British were excluded (eps=60). For countries that are very similar to Britain in ancestry (e.g., Germany, Denmark) this is a balance between excluding individuals who are genuinely British (very common in the 'German' samples) but not biasing the samples away from British-like ancestry.

In the second scan, the remaining individuals were clustered, and the largest cluster was chosen to represent a typical ancestry for that country. The appropriate eps value (i.e. how strict the clustering should be) is a reflection of the genetic diversity of a country, and so was adjusted manually to reflect this (Supplementary Table 2). In a minority of cases, the major cluster was not the indigenous ancestral background, and so the second-largest was chosen (for example, in Kenya the largest cluster was individuals of Indian origin). All selections were visually verified. Countries that had no obvious main cluster (usually due to low sample numbers) were excluded; any country with 3 or fewer individuals was also excluded.

In order to select Irish individuals (Republic of Ireland and Northern Ireland), step 1 was skipped but step 2 was run with relatively tight parameters, in both cases excluding approximately 20% of individuals.

In order to test the effectiveness of the pipeline in selecting individuals of a similar ancestral background, we looked at the variance in the genome-wide painting proportions for each country. Countries with high variance would indicate recent admixture.

#### Results

This pipeline selected 24,511 individuals from 126 countries. These selected samples were painted using a reference/donor panel of ancient individuals (Supplementary Note 2).

The countries that had high variance in ancestry proportions among individuals (and therefore likely that the DBSCAN was not effective in choosing individuals of a similar ancestral background) were Kazakhstan, Yemen, Egypt, Seychelles. Results for these countries should be interpreted with caution.

#### Discussion

The UKB represents an important source of data for white British people but also for people from other countries globally. Usually, researchers restrict themselves to the white British cohort, but here we develop a method to select individuals from other countries. This transforms the UKB from a resource that is informative about British ancestry to one that can be used to make inferences about populations worldwide.

#### References

1. Pedregosa, F. *et al.* Scikit-learn: Machine learning in Python. *The Journal of Machine*

*Learning Research* **12**, 2825–2830 (2011).

2. Bycroft, C. *et al.* The UK Biobank resource with deep phenotyping and genomic data. *Nature* **562**, 203–209 (2018).

### 2) Painting the UK Biobank

William Barrie[1] and Dan Lawson[2]

[1] Zoology Department, University of Cambridge, UK.
[2] School of Mathematics and Integrative Epidemiology Unit, University of Bristol, UK.

#### Introduction

Here, we develop new methods to use ChromoPainter [1] on a biobank scale to 'paint' modern genomes from the UK Biobank (UKB) using ancient genomes, grouped into reference populations, as donors. Painting was done following the pipeline of Margaryan et al. [2] based on GLOBETROTTER [3], and admixture proportions were estimated using Non-Negative Least squares. These results (technically most recent coalescences) are used as a proxy for ancestry. We store both genome-wide and local ancestry (i.e., per variant per individual) results.

#### Methods

##### Painting pipeline introduction

The process of painting consists of forming a reference/donor panel consisting of ancient individuals of as pure ancestry as possible, having undergone QC and clustering using fineSTRUCTURE. The target/recipient panel and reference/donor panel are then filtered for variants, merged, and the target panel is painted using the reference panel as donors.

##### Reference/donor panel formation

We used imputed best guess haplotypes filtered for imputation information score (FORMAT/INFO) above 0.5. Samples were selected based on IBD-sharing, visual PCA inspection, and fineSTRUCTURE analysis (unsupervised clustering based on the co-ancestry matrix output of ChromoPainter; Figure S1); low coverage, contaminated, and related individuals were excluded. The aim was to group samples into as pure 'source' populations as possible, while maintaining reasonable numbers in each population. We do not expect our filters for white British/non-British individuals to be perfect; furthermore, modelling modern Eurasians as a mixture of hunter-gatherer/Steppe/farmer is overly simplistic. Therefore, we also include ancient African and East Asian reference populations to account for possible ‘non-European’ ancestry.

Ultimately, 318 individuals split into ten reference populations were used (Figure S2, Figure S3, Supplementary Table 3): western hunter-gatherer (WHG), eastern hunter-gatherer (EHG), Caucasus hunter-gatherer (CHG), Farmer Anatolian, Farmer Early, Farmer Middle, Farmer Late, Yamnaya, African and East Asian. Populations are characterised by preferentially copying from individuals within the population, as well as being biologically and historically meaningful. This dataset is henceforth called the “present aDNA dataset”.

The farmers are split into four separate populations due to their differing behaviour as donors (columns) in the fineSTRUCTURE analysis (Figure S1). There is a cline in their degree of WHG admixture that roughly correlates with age, while some samples also show Steppe admixture. Given the nature of the splits, the differences between these groups should be interpreted with caution, and for most downstream analysis these groups are merged into a ‘Farmer’ ancestry.

#### Target/recipient panel formation

We used white British individuals from the UKB as reported in Bycroft et al.[4]; these are individuals who self-reported as white British and have British-like ancestry according to PCA. We also used individuals from the UKB of a typical ancestral background selected by country of origin (Supplementary Note 1). We used phased haplotype data, downloaded from https://www.ukbiobank.ac.uk. This totalled 408,884 white British individuals, and 24,511 non-British individuals. This dataset is henceforth called the “UKB dataset”.

#### SNP selection and merging of the panels

Due to computational considerations, the number of SNPs used in the painting was limited to those in the UKB Axiom Array; these SNPs were chosen to capture genome-wide variation, rare and coding variants, and variants relevant to specific phenotypes or regions of interest [4]. The present aDNA dataset and UKB datasets were merged and filtered for these variants using QCTOOL v2 (https://www.well.ox.ac.uk/~gav/qctool_v2/), and then filtered to exclude variants with a minor allele frequency below 1% using bcftools (http://samtools.github.io/bcftools/), leaving a total of 549,323 SNPs across chromosomes 1-22.

#### Painting process

ChromoPainter [1] uses an approach premised on the observation that markers on the same chromosome are inherited together unless separated by recombination; at the population level, this results in linkage disequilibrium (LD) between close markers that reflect a shared history of descent. The haplotype-based algorithm of ChromoPainter aims to harness this information, detecting shared haplotypes to reconstruct phased recipient genomes as chunks 'copied' from donors.

Considering the genealogy of a single locus, we can identify one or more closest relatives to that locus, henceforth called 'nearest neighbours'; if viewed as a genealogy, these are the other leaves of the tree underneath the first coalescence. Therefore, at each locus of each haplotype, there exists one or more nearest neighbours. ChromoPainter aims to identify these using an approximate method based on that introduced by Li and Stephens [5]: the Hidden Markov Model (HMM), which explicitly reconstructs the haplotype of a recipient/target individual as a series of chunks of genetic material donated by the other donor/reference individuals, using information on the types of the recipient and potential donor at each SNP. This approach is probabilistic, calculating the expectations of which haplotype acts as donor to a recipient as a function of position over an infinite number of paintings [1]. Although ChromoPainter was originally intended to use this information, in the form of a 'co-ancestry matrix', to ascertain fine-scale population structure and clustering (in the fineSTRUCTURE software package), the software can be used with pre-defined donor and recipient populations.

If the donor panel is formed of ancient individuals and the recipient individual is modern, the nearest neighbour should reflect some history of that locus. The ability of chromosome painting to accurately infer ancestry is expected to depend on the diversity between donor populations: more genetically similar populations and the algorithm will find it difficult to correctly identify the nearest neighbour(s). There is also the issue of 'masking' whereby haplotypes from older populations would have travelled through more recent populations before arriving in the modern population; this causes a genome-wide bias towards the more recent ancient populations, the effects of which are discussed below. Here, we use nearest neighbour as a proxy for local ancestry - i.e., which population that haplotype came from (which may not be a single unique population from our panel).

Chromosome painting cannot include the target of painting. Therefore, painting was done (following the pipeline of Margaryan et al. [2] based on GLOBETROTTER [3] by leaving out one individual at random (chosen independently for each chromosome) from each other donor population for all donor individuals. Target individuals from the UK Biobank were painted by similarly removing one individual at random from all donor populations. This ensures that individuals from the reference and UK Biobank are exchangeable.

Once we had a well-chosen set of ancient populations from the present aDNA panel, each individual was repainted twice leaving out themselves as a possible donor: first to learn the painting parameters Ne and μ, and then to learn a genome-wide individual-specific donor-prior. For each of the reference populations, the average amount of genome received from each donor individual was learnt. We then painted the modern individuals in the UKB panel using the reference populations and the learnt parameters and priors.

The probability that each recipient copied each donor population at every SNP was recorded. The genome-wide information for each recipient was also stored, in the form of (i) chunkcounts, the number of chunks copied from each donor population and (ii) chunk lengths, the sum of the lengths of the chunks copied from each population, weighted by their copying probability. Admixture proportions were then estimated using Non-Negative Least Squares (NNLS).

##### Painting at biobank scale

We used custom scripts to speed up this process (specifically reading from large phase files), to enable running for large numbers of recipients in parallel across multiple nodes, and to store the local copying probabilities in a memory-efficient format in real time (all scripts available at https://github.com/will-camb/Nero/tree/master/scripts/cp_panel_scripts). The total CPU time for painting the UKB panel was approximately 550,000 CPU hours.

#### Results

##### Ancestry-PCs relationship

PCA is a dimensionality reduction technique that can be applied to genetic data, the results of which are useful as a means to visualise variation between individuals/groups, and are expected to reflect historical events that cause differences in ancestry due to drift, admixture etc. It is well established that PC1 vs PC2 vs PC3 generally separate African, European, and East Asian populations. We ran multivariate linear regressions using ancestry components to predict UKB PCs [4]. Previous work has shown that the main UKB PCs that reflect British population structure are PCs 5 and 9, describing variation between English, Scottish and Welsh ancestry, and PCs 11 and 14 which further separate structure within Wales and England [6].

We found significant correlations between ancestry components and PC4 (R-squared=0.553) and PC5 (R-squared=0.376), as well as PC1 (R-squared=0.165) and PC7 (R-squared=0.130). We found that the high PC4 correlation with ancestry component was largely driven by a Steppe (Yamnaya/EHG) vs Farmer divide, both within Britain and internationally: high PC4 values are associated with high Steppe/low Farmer ancestry, while low PC4 values are associated with low Steppe/high Farmer ancestry.

#### Ancestry-geographic variation

Within the British Isles, all individuals were painted with similar proportions from each reference population, as expected when measuring coalescence tracts rather than direct admixture tracts and after a long time since admixture events; but the differences in copying proportions showed significant geographic heterogeneity. We ran multivariate linear regressions, using longitude and latitude of place of birth ("Place of birth in UK – east co-ordinate" and "Place of birth in UK – north co-ordinate") to predict NNLS ancestry fractions. We found significant correlations for Yamnaya ancestry (R-squared=0.081), Farmer ancestry (R-squared=0.066), CHG ancestry (R-squared=0.015), WHG ancestry (R-squared=0.007), African ancestry (R-squared=0.011) and EHG ancestry (R-squared=0.01, longitude only). To visualise this, we assigned individuals to a county based on their UKB place of birth data and plotted the average admixture proportion per county for each ancestry, binned in ten equal interval quantiles using ArcGIS Online (www.arcgis.com; Figure 2, main text).

We found that Neolithic farmer ancestry was highest in southern and eastern England and lower in populations in Scotland, Wales, and Cornwall. We found the opposite pattern in Yamnaya ancestry, representing the Steppe component, which has previously been shown to be higher in Scotland but not Wales [7]; we found this was highest in the Outer Hebrides. This Farmer/Yamnaya dichotomy broadly reflects an 'Anglo-Saxon'/'Celtic' distribution. We are unable to date when these subtle population structures arose but note that the Neolithic Anatolian-related farmer ancestry is already present in the British and Roman Iron Age but lower in Saxon individuals, meaning these patterns cannot be explained just by Saxon-related ancestry. They are likely a result of pre-Roman migration between 1000 and 875 BC which resulted in a slight increase in Early Farmer ancestry in England and Wales but not

Scotland [8], although we note our results show a marked difference between Wales/Cornwall and England too. We also found higher levels of WHG-related ancestry in central and Northern England.

Looking at a continent-wide level, the hunter-gatherer ancestries display distinct structure in modern populations (Figure 2, main text). WHG-related ancestry is highest in present-day individuals from the Baltic States, Belarus, Poland and Russia; EHG-related ancestry is highest in Mongolia, Finland, Estonia and Central Asia; and CHG-related ancestry is maximised in countries east of the Caucasus, in Pakistan, India, Afghanistan and Iran, in accordance with previous results [9]. The CHG-related ancestry likely picks up both Caucasus hunter-gatherer and Iranian Neolithic signals, explaining the relatively high levels in south Asia [10]. Consistent with expectations [11,12], Neolithic Anatolian-related farmer ancestry is concentrated around the Mediterranean basin, with high levels in southern Europe, the Near East and North Africa, including the Horn of Africa but less in Northern Europe. A contrasting pattern was observed in Yamnaya-related ancestry decreasing from high levels in northern Europe, peaking in Ireland, Iceland, Norway and Sweden, but decreasing further south where Neolithic farmer ancestry still dominates. There is also evidence for its spread into southern Asia. These results provide a new level of detail on the modern distribution of ancient ancestries.

To better understand how countries varied in their ancestry proportions, we ran Scikit-learn's PCA [13] on the average admixture proportions per country. We then ran a hierarchical clustering algorithm on the first 4 PCs (explained variance=0.244), and built a dendrogram (Figure S4) [14]. For further analysis, we excluded countries in clusters dominated by African or East Asian ancestry, leaving 80 countries.

We used Sklearn's StandardScaler utility class to standardise each feature (zero mean and unit variance), then ran a PCA using the standardised average admixture proportions for each of the 80 remaining countries (https://github.com/erdogant/distfit), and plotted a biplot (PC1 vs PC2 with loadings for each feature plotted), which shows the correlations between ancestries (Figure S5). This gives a more visual representation of how countries group by ancestry, and broadly reflects actual geography.

#### Discussion

The large ancient DNA panel established here combined with the UKB allows us to trace for the first time the fine-scale distribution of Mesolithic/Neolithic/Bronze Age ancestry components in modern British individuals, using DNA directly from ancient individuals. It also demonstrates the ancestry differences within an ‘ethnic group’ (white British) traditionally regarded as being relatively homogenous, highlighting the need for care over ancestry considerations when using resources like the UK Biobank.

#### Figures

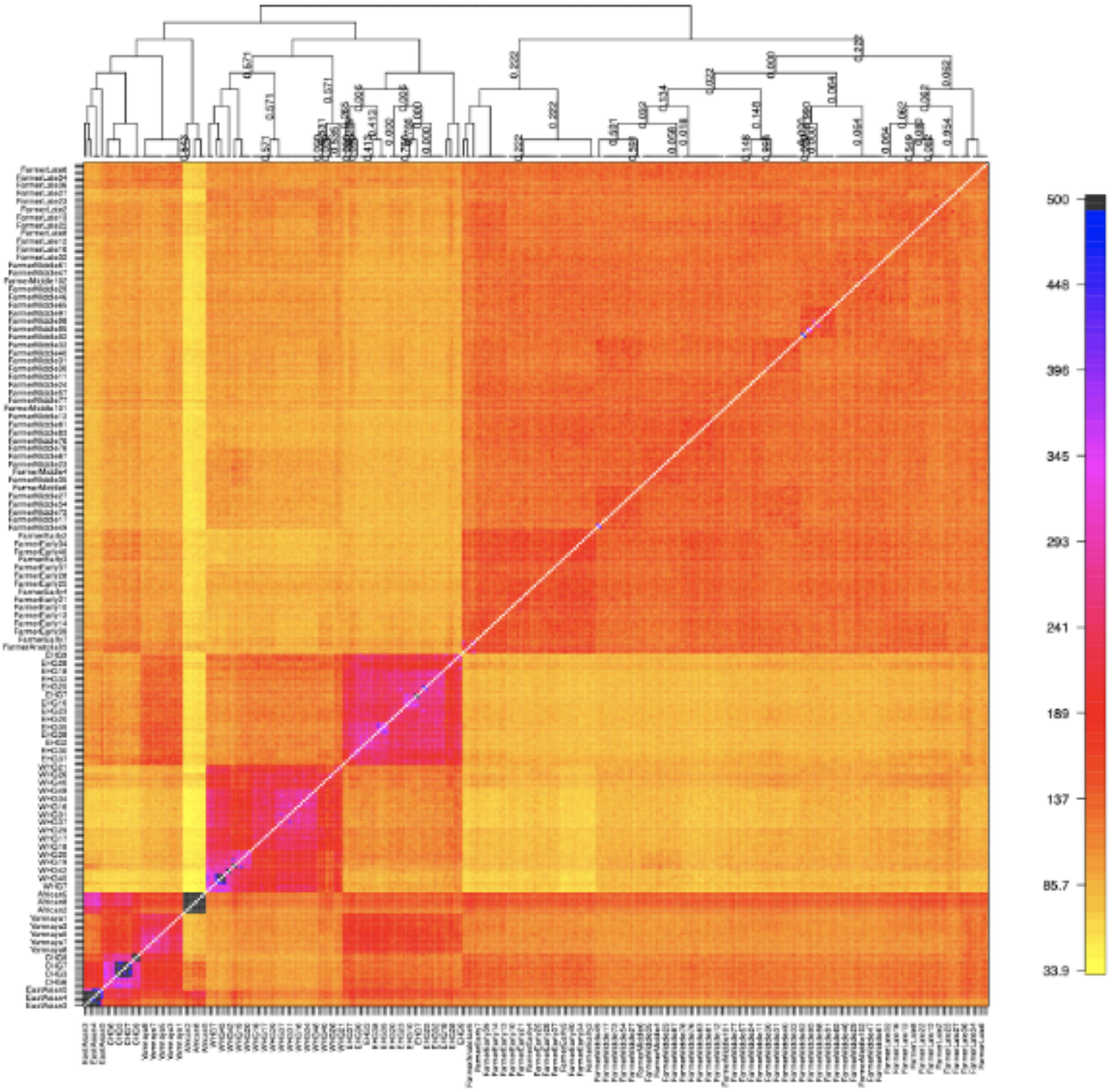


**Figure S1.** Co-ancestry heatmap of selected ancient samples. The output of fineSTRUCTURE analysis of the ancient reference panel, showing copying proportions between ancient populations (columns=donors, rows=recipients). There is a cline in Hunter-Gatherer admixture in the Farmers, roughly correlating with age. For most downstream analyses, the Farmer populations were merged.

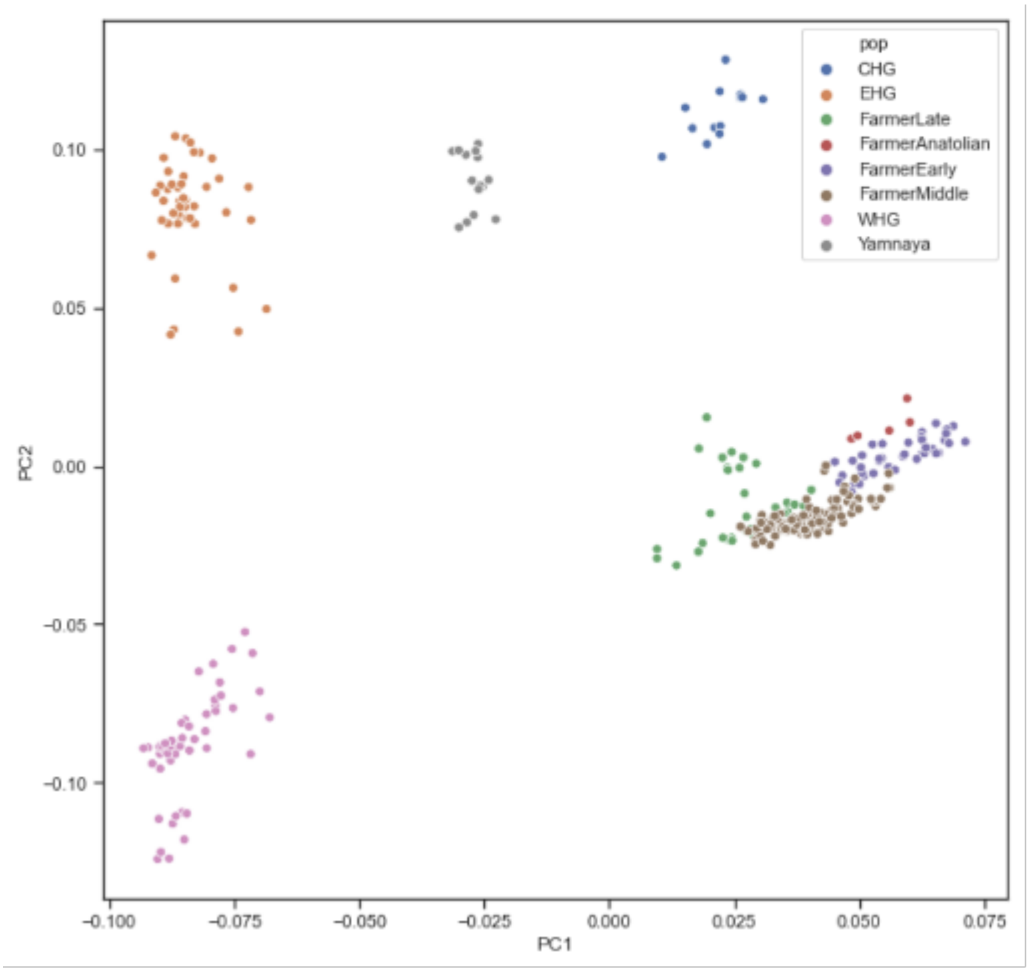


**Figure S2.** PCA of ancient reference samples, coloured by assigned population. PC1 vs PC2 of a PCA of the ancient western Eurasian samples (excluding African and East Asian), coloured by their assigned population used in the painting. As can be seen, populations are fairly distinct, with intermediate admixed individuals having been excluded. Some Farmers are admixed with Steppe and Hunter-Gatherer populations to differing degrees, but particularly among later individuals.

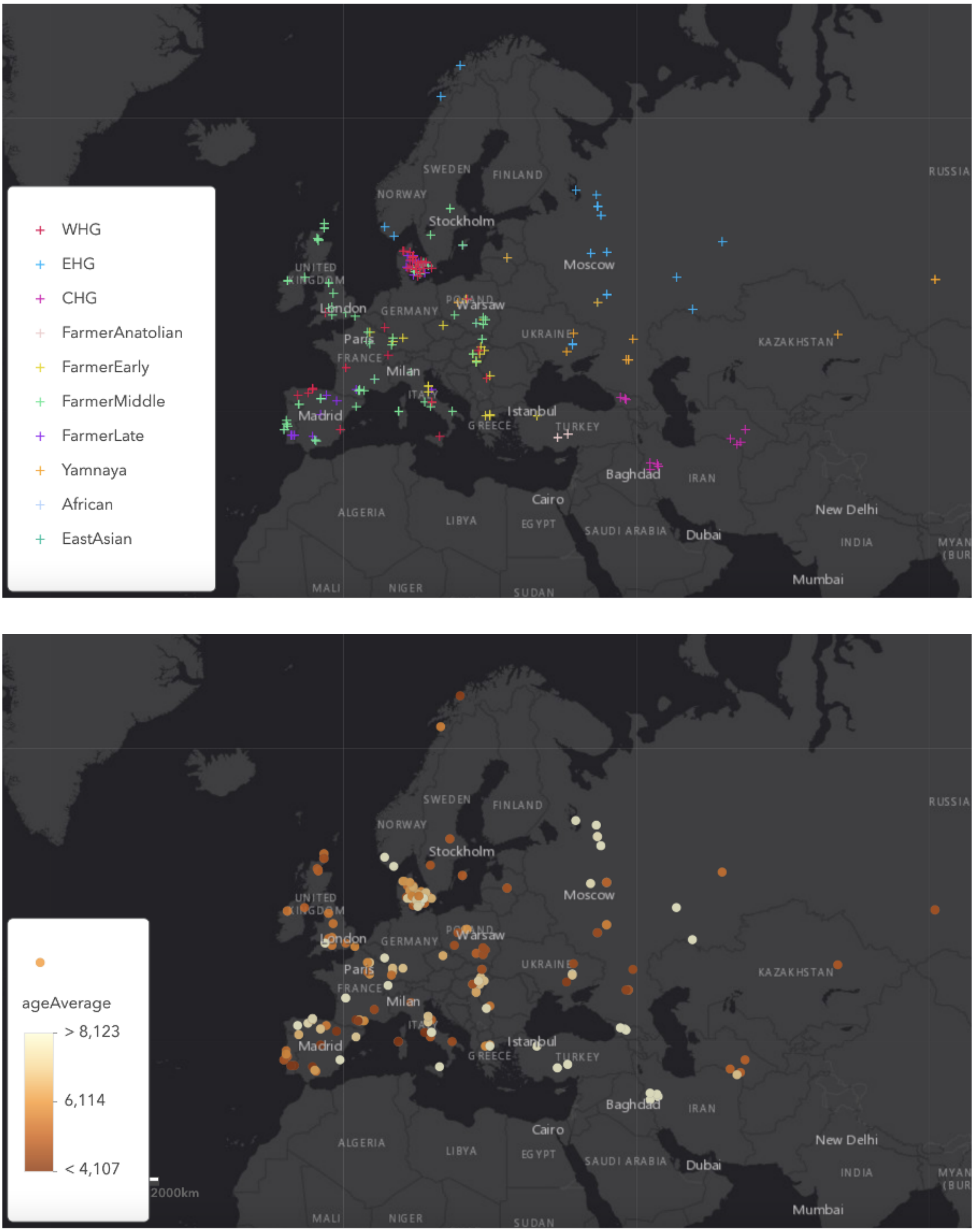


**Figure S3.** Maps of ancient sample locations coloured by assigned reference population (above) and age (below). Not showing African and East Asian samples.

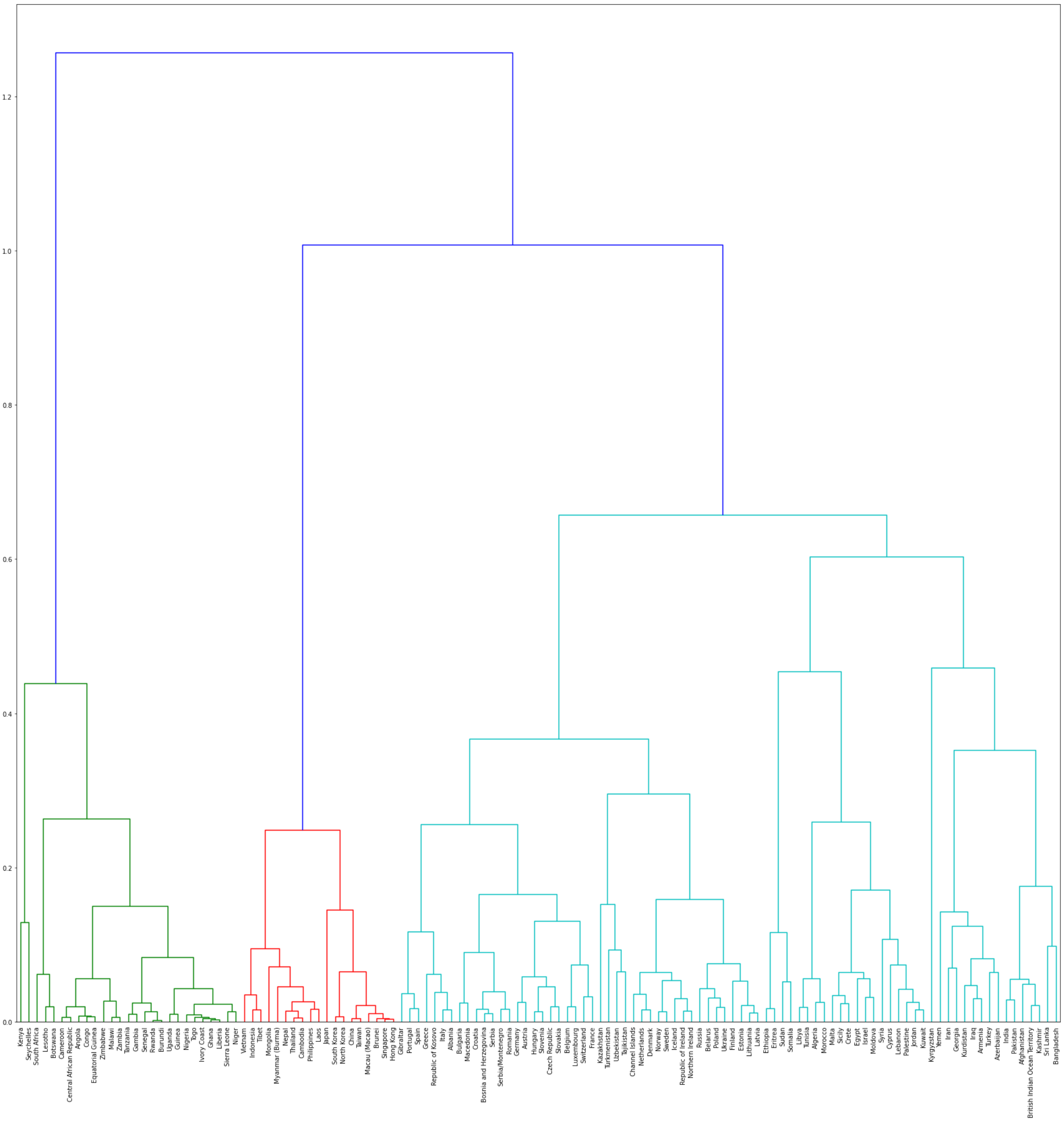


**Figure S4.** Dendrogram based on hierarchical clustering of first 4 PCs of average admixture proportions per country. For further analysis, countries in clusters dominated by African (green) or East Asian (red) ancestry were dropped.

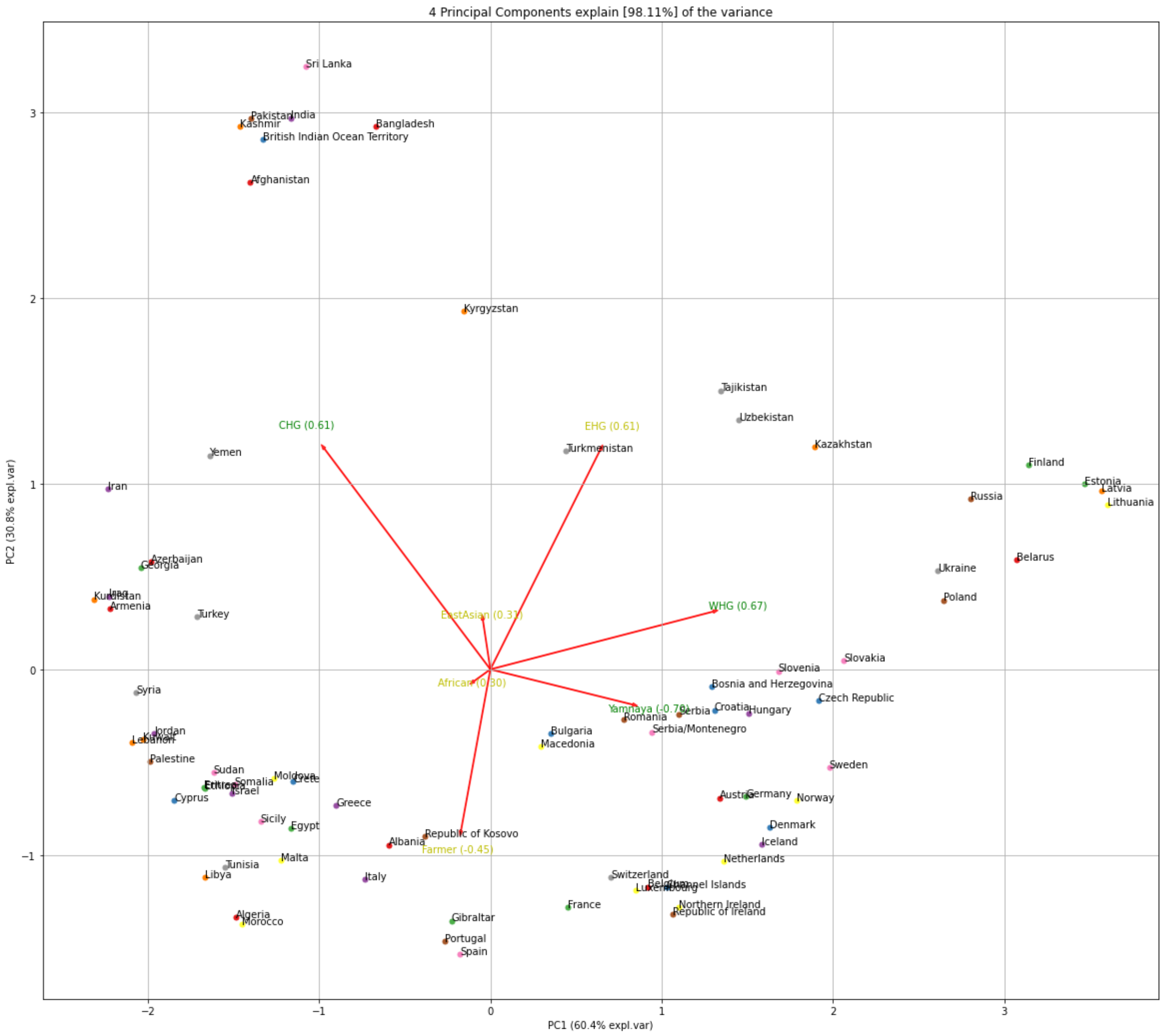


**Figure S5.** PCA biplot of standardised average NNLS admixture proportion per country, based on 80 countries in Europe, West/Southern Asia, the Middle East, and North Africa.

#### References


1. Lawson, D. J., Hellenthal, G., Myers, S. & Falush, D. Inference of population structure using dense haplotype data. *PLoS Genet.* **8**, e1002453 (2012).

2. Margaryan, A. *et al.* Population genomics of the Viking world. *Nature* **585**, 390–396 (2020).

3. Hellenthal, G. *et al.* A genetic atlas of human admixture history. *Science* **343**, 747–751 (2014).

4. Bycroft, C. *et al.* The UK Biobank resource with deep phenotyping and genomic data. *Nature* **562**, 203–209 (2018).

5. Li, N. & Stephens, M. Modeling linkage disequilibrium and identifying recombination hotspots using single-nucleotide polymorphism data. *Genetics* **165**, 2213–2233 (2003).

6. Sarmanova, A., Morris, T. & Lawson, D. J. Population stratification in GWAS meta-analysis should be standardized to the best available reference datasets. *bioRxiv* 2020.09.03.281568 (2020) doi:10.1101/2020.09.03.281568.

7. Galinsky, K. J., Loh, P.-R., Mallick, S., Patterson, N. J. & Price, A. L. Population Structure of UK Biobank and Ancient Eurasians Reveals Adaptation at Genes Influencing Blood Pressure. *Am. J. Hum. Genet.* **99**, 1130–1139 (2016).

8. Patterson, N. *et al.* Large-scale migration into Britain during the Middle to Late Bronze Age. *Nature* (2021) doi:10.1038/s41586-021-04287-4.

9. Sikora, M. *et al.* The population history of northeastern Siberia since the Pleistocene. *Nature* **570**, 182–188 (2019).

10. Shinde, V. *et al.* An Ancient Harappan Genome Lacks Ancestry from Steppe Pastoralists or Iranian Farmers. *Cell* **179**, 729–735.e10 (2019).

11. Hofmanová, Z. *et al.* Early farmers from across Europe directly descended from Neolithic Aegeans. *Proc. Natl. Acad. Sci. U. S. A.* **113**, 6886–6891 (2016).

12. Feldman, M. *et al.* Late Pleistocene human genome suggests a local origin for the first farmers of central Anatolia. *Nat. Commun.* **10**, 1218 (2019).

13. Pedregosa, F. *et al.* Scikit-learn: Machine learning in Python. *J. Mach. Learn. Res.* **12**, 2825–2830 (2011).

14. Virtanen, P. *et al.* SciPy 1.0: fundamental algorithms for scientific computing in Python. *Nat. Methods* **17**, 261–272 (2020).

### 3) Local Ancestry Inference for Complex Population Histories

Alice Pearson[1], Richard Durbin[1,2]

[1] Department of Genetics, University of Cambridge, UK.

[2] Wellcome Sanger Institute, Wellcome Genome Campus, UK.

#### Abstract

It has become apparent from ancient DNA analysis, that the history of many human populations from across the globe are often complex, involving multiple population split, admixture, migration and isolation events. Local ancestry inference (LAI) aims to identify from which ancestral population chromosomal segments in admixed individuals are inherited. However, ancestry in existing LAI tools is characterised by a discrete population identity, a definition which is limited in the context of a complex demographic history involving multiple admixture events at different times. Moreover, many LAI tools rely on a reference panel of present day genomes that act as proxies for the ancestral populations. For ancient admixture events, these proxy genomes are likely only distantly related to the true ancestral populations. Here we present a new method that leverages advances in ancient DNA sequencing and genealogical inference to address these issues in LAI. The method applies machine learning to tree sequences inferred for ancient and present day genomes and is based on a deterministic model of population structure, within which we introduce the concept of path ancestry. We show that the method is robust to a variety of demographic scenarios, generalises over model misspecification and that it outperforms a leading local ancestry inference tool. We further describe a downstream method to estimate the time since admixture for individuals with painted chromosomes. We apply the method to a large ancient DNA dataset covering Europe and West Eurasia and show that the inferred admixture ages are a better metric than sample ages alone for understanding movements of people across Europe in the past.

#### Introduction

Despite a wealth of data, patterns in modern genomes alone are difficult to interpret as they are an indirect measure of past events. Moreover, much of the genetic variation which was present in past populations does not exist in the modern gene pool due to ancient demographic events that exacerbate drift, such as bottlenecks and isolation. Ancient DNA provides a genetic snapshot of a time before these processes have taken place, meaning we regain some of the lost information. As a result, the use of aDNA has transformed our understanding of human origins and evolution in recent years [1] showing that the histories of human populations are often complex, involving periods of isolation and migrations with admixture events [2].

Admixture events occur when there is migration between divergent populations, and they interbreed. Chromosomes of the resulting admixed individuals will originate in one of the two ancestral populations. With each generation there is recombination, meaning that over time chromosomes in the admixed population will be a mosaic of chunks originating in the two ancestral populations. Local ancestry inference (LAI) is the process of decomposing admixed chromosomes into these ancestral chunks and assigning each chunk an ancestry label. Many tools are available that perform LAI due to its importance for understanding population structure, migration history and disease risks [3, 4].

Early LAI methods [5, 6] use unlinked markers that are characteristic of populations, so called ancestry informative markers (AIMs). The hidden Markov model (HMM) structure employed by these methods relied on the assumption that markers are independent and so do not model background LD but admixture LD alone. With the decrease in cost of genotyping in recent years, the ability to deconvolve local ancestry with greater resolution using denser SNP data became possible but violated the assumption of independence between SNPs. Methods that leverage this denser data emerged [7, 8] that employed a pruning step which makes sure SNPs are unlinked in the ancestral populations. However, not explicitly modelling background LD within ancestral populations prevents the use of many informative SNPs that are linked. Later methods now both utilise denser SNP data and model background LD together with admixture LD by using extensions of the basic HMM approach and leveraging haplotype frequencies, which vary more between populations than SNP frequencies [9–11]. More recently machine learning has been used for local ancestry inference and has shown to be computational efficient and accurate. Furthermore, the ever increasing availability of genomic data provides training data for supervised methods [12, 13].

Complex population histories pose a problem for LAI as most of the existing methods require sequences that represent discrete ancestral populations to form a reference panel [7, 9, 10, 12, 13]. In reality, haplotypes are inherited from generation to generation, often through a complex demographic history, possibly involving multiple admixture events between many populations. An ancestral population may itself be an admixed population from an earlier event in history. In other words, populations are more like a braided river than a sequence of well-defined discrete population identities. Thus, assignment of a haplotype in an admixed chromosome to a single ancestral population in a reference panel does not inform us of ancestry further back in time or take into account the genealogical relationship of that ancestral population to other reference populations. Some LAI tools allow multiple reference populations that could represent many populations involved in a focal population history from different time periods [7, 9, 12, 13], but haplotypes in focal admixed samples can only be assigned to one of these ‘ancestries’ from the reference panel when in fact two or more assignments may be true. Moreover, the reference panel is often made up of present day samples that are closely related proxies of the true ancestral populations and thus the accuracy of ancestry assignment varies with how well the reference populations represent the true ancestry populations. The relationship of present day genomes to ancestral populations that existed deeper in time becomes more distant making LAI less accurate for more ancient admixture events.

Here we develop a local ancestry inference method that takes the genealogical relationships of ancestral populations into account and leverages the power of ancient samples. For this we redefine ‘ancestry’ as no longer a discrete population identity, but a complete path back in time through the population history. The path that a haplotype takes backwards in time from a focal individual is fully informative about its local ancestry. Ancient samples are used to represent ancestral populations along these paths as they are more closely related to the true ancestral groups involved in the admixture events. The method involves building tree sequences, a representation of the ancestral recombination graph, between a large set of sample sequences using RELATE [14]. RELATE enables sample sequences to be placed in the past [15] and so we can incorporate ancient samples into the tree sequences [15]. We then train a neural network, using a simulated training set, that classifies the ancestry path for each sample haplotype in a genomic region given the local tree and a known population history.

There has been extensive research on the population history of Europeans [2, 16, 17] and the broad picture is well established. Therefore, we used a large dataset of present day and high quality shotgun sequenced, imputed ancient genomes (MesoNeo dataset) to test the method. We constructed a model in msprime [18] that represents the major ancestry flows contributing to modern European genomes over the last 50,000 years, from which we simulated tree sequence and variation data. Local ancestry inference performed by a neural network, trained on simulated data from this model, is as good if not better than inference using GNOMix [13], a leading LAI tool. Additionally, we showed the method is robust to a range of simulated demographic scenarios and model misspecification.

Lastly, we applied a technique to infer the time since admixture of admixed MesoNeo individuals using the rate of switching between painted segments and thus gain insight into the patterns of admixture events that occurred in history between major ancient European groups.

#### Method and Materials

##### Concept of path ancestry

We explain in more detail the issue current LAI tools face when using a reference panel of proxy ancestral population from which only one population can represent the ancestry of each admixed haplotype. Figure S6 shows a diagrammatic example of a population structure with two consecutive admixture events and two population split events; population C is an ancestral population to population E. A haplotype in E can be assigned to both C and A; both are true at the same time. Existing LAI methods could include both C and A as discrete ancestral reference populations, producing confusing results or choose either C or A for the reference panel and only use half the available data. In other words, all LAI methods treat ancestral populations as discrete entities with no genealogical relationships with each other, requiring them to effectively draw a line in the past and take the populations existing at that time as ‘pure’ ancestral populations.

The concept of ‘path’ ancestry takes time and genealogical relationships of ancestral populations into account. We redefine ‘ancestry’, not as a discrete population identity, but a complete path back in time through the population history. The path that a haplotype takes backwards in time from a focal individual is fully informative about its local ancestry: By determining what populations a haplotype has been carried by inheritance through a structured population history, the relationship of the haplotype to all relevant historical and admixing populations is established.

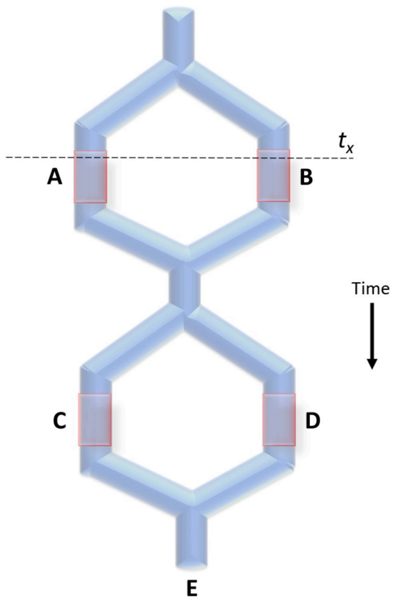


**Figure S6.** Population structure with paths. Schematic of a structured meta-population that goes through two population split events and two admixture events. Populations marked A, B,C and D are 'ancestral populations' and population E is the admixed population whose local ancestry is of interest.

Returning to the example in Figure S6; take population E as the population whose haplotypes we wish to perform LAI on. We have representative samples from populations A, B, C, and D from within this structure which are termed 'ancestral populations'. Between these four ancestral populations, there are four paths that haplotypes could have taken backwards in time from population E, through multiple populations:

$$
\begin{aligned}
E \to C \to A \\
E \to D \to A \\
E \to C \to B \\
E \to D \to B
\end{aligned}
$$

By using paths as local ancestry labels instead of discrete population identities, we are able to use all available data from all four ancestral populations and assign four different labels that convey meaningful information about the history of a haplotype.

Ancient samples within this framework are a huge advantage: The further back in time a population existed, the more difficult it is to find a closely related proxy population that exists today and from whom genomic data is available. Ancient samples are likely to be less diverged from the true ancestral populations and so act as better representatives that existed before one or many admixture and split events.

#### Constructing a path model of European population history

Using the extensive amount of previous work elucidating the genetic history of Europeans, we put forward a standardised model of European population structure. Figure S7 shows a schematic of this demographic model that describes the population structure in Europe during the last 50,000 years. Shortly after the expansion of anatomically modern humans into Eurasia, there is a population split 15,000 years ago (1500 generations ago) between the Northern Europeans (NE), who continued moving northwest into Europe, and West Asians (WA) who stayed more locally in the Levant and South Caucasus area. The WA population then splits to form the Caucasus hunter gatherers/early Iranians (CHG) and the Anatolians (Ana) ~24,000 years ago (800 generations ago). Within the NE, ~18,000 years ago (600 generations ago), the Western hunter gatherers (WHG) and Eastern hunter gatherers (EHG) diverged. At that point, the four separate populations that make up present day European ancestry are distinct in the model. The timing of divergence between the four distinct groups appears to correspond to the onset of the Last Glacial Maximum [17, 19], during which time these populations became isolated from each other in refugia.

The model subsequently describes how admixture between these populations leads to the modern European gene pool. At the start of the Neolithic around 9,000 years ago, farmers from Anatolian moved into Europe. These incoming Anatolian farmers admixed with the local WHG ~7,800 years ago (259 generations ago) to form the Neolithic farmer population [20, 21] and their ancestry reaches Britain by 6,000 years ago [22].

During the late Neolithic, steppe populations, characterised by the Yamnaya (Yam) culture, appear as a mix of EHG and CHG/Iranian ancestries [17, 19, 23]“-5,300 years ago (177 generations ago). At a similar time, the Bronze Age Anatolian group (BAA) represents a later Anatolian population formed from an admixture of CHGs with Ana −5,100 years ago (170 generations ago). Around the start of the Bronze Age −4,900 years ago (166 generations ago), migrants with Yam ancestry moved west into Europe and had a profound impact on the genetic landscape by admixing with the Neolithic farmers to form the European Bronze Age population, leading to present day Europeans [16, 20, 23–25].

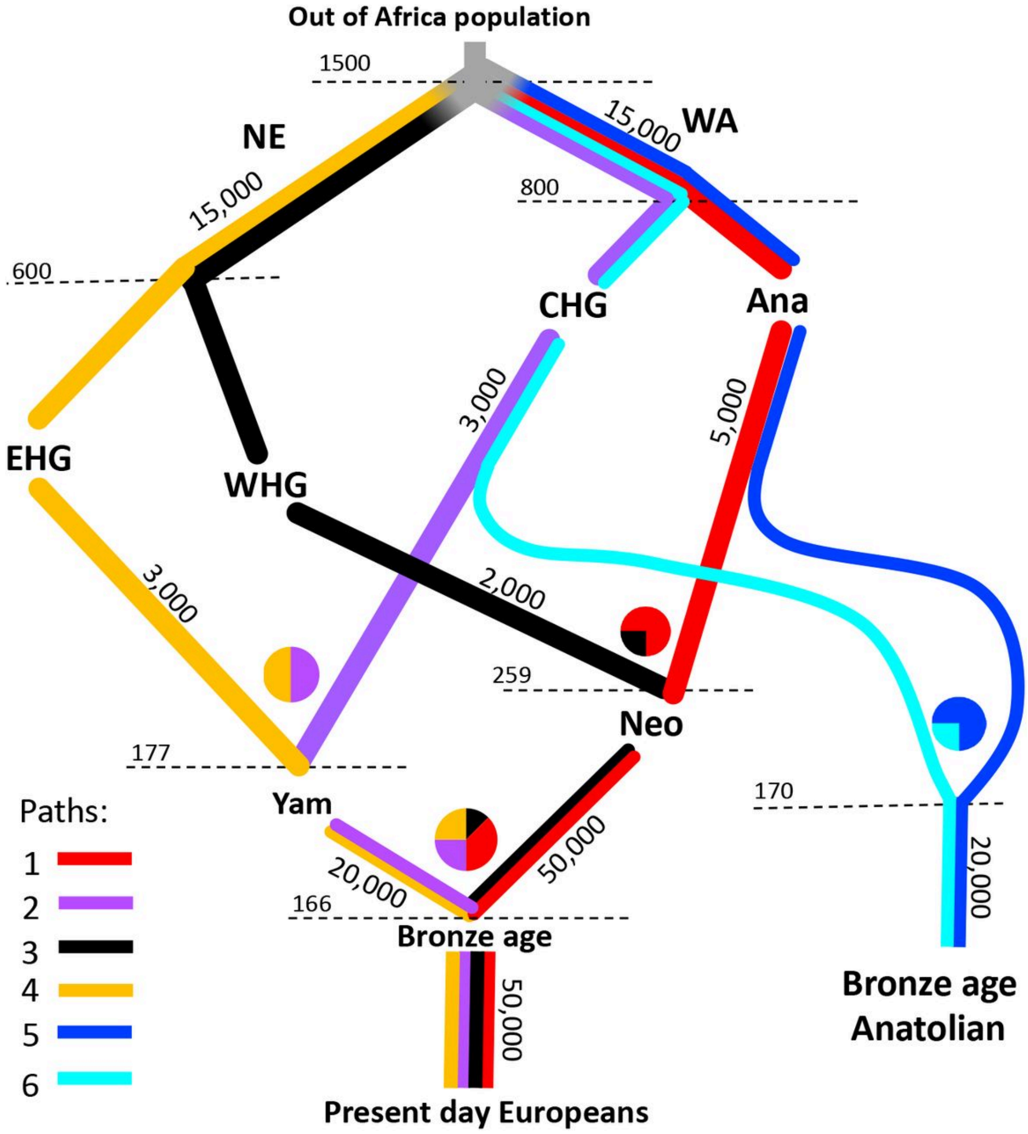


**Figure S7.** Schematic diagram of the model of European population structure. Population labels are abbreviated. Bronze Age = Bronze Age European population, Yam = Yamnaya steppe, Neo = Neolithic farmers, WHG = Western hunter gatherers, CHG = Caucasus hunter gatherers, EHG = Eastern hunter gatherers, Ana = Anatolian farmers, NE = Ancient Northern Europeans, WA = Ancient West Asians. Effective population sizes are shown along edges and admixture fractions are displayed as pie charts. The timing of population splits and admixture events are shown at the dotted lines in units of generations ago.

Present day Europeans are often described as a three-way mix of WHG, Anatolian Farmer and steppe Yamnaya [2]. However, the Yamnaya ancestry itself resolves into EHG and CHG ancestry [17, 26] and these two groups have different deep origins as seen in Figure S7 [17].

Therefore, we propose a model of four ancestral streams that lead to present-day Europeans.

Six different paths that haplotypes can take from any sampled individual are shown in different colours in Figure S7. Path 1 = red, starts from the present day Europeans, going back through Neolithic farmers, Anatolian farmers, West Asians to the root. Path 2 = purple, starts at present day Europeans, going back through the Yamnaya, Caucasus hunter gatherers, then West Asians to the root. Path 3 = black, starts with present day Europeans, going back through Neolithic farmers to Western hunter gatherers and then through Northern Europeans to the root. Path 4 = orange, starts at present day Europeans, going back through the Yamnaya to Eastern hunter gatherers and then through Northern Europeans to the root. Paths 5 = blue, starting in the Bronze Age Anatolians and joins path 1 part. Path 6 = cyan, starts in Bronze Age Anatolians and joins path 2. When paths overlap, lineages from all overlapping paths can coalesce.

We constructed our model of European population history in msprime [18], a coalescent simulator. The population sizes and admixture fractions are shown in the schematic in Figure S7. This model has been submitted to the stdpopsim catalogue (https://github.com/popsim-consortium/stdpopsim/blob/main/stdpopsim/catalog/HomSap/demographic_models.py) and is publicly available for users to simulate data from.

#### Comparison of simulated data to MesoNeo genomes

The MesoNeo dataset is a large collection of both ancient and present day samples. While the dataset contains samples from all over the world, most are from people that lived in the last 10,000 years in Eurasia. The dataset is composed of 1,490 genomes that have been imputed to 3.7 million SNPs once filtered for coverage (>0.1X), low imputation quality, close relatives and >0.5 imputation INFO score (See [27] for data availability).

To test how well our model represents the true history of Europeans, we compared data simulated from the model to the MesoNeo genomes. Based on PCA (Figure S26) we subset the 1,492 genomes from the MesoNeo dataset to those that were tightly clustered in the groups relevant to the genetic history of Europeans, so that only samples diagnostic of each population in our model were kept.

For our analysis of the time since admixture (see below), we chose 476 diploid individuals (952 haploid) including 91 GBR 1000 Genomes samples. During simulation, samples were taken from populations and times to match each real diploid sample, using their radiocarbon or context dates. We aimed to create a simulated dataset that closely matched our MesoNeo subset genetically.

For our time-series selection analysis (Supplementary Note 4), we used an expanded set of 1,015 ancient genomes relevant to modern European genetic history. This included samples that have a West Eurasian archaeological location and lying on EHG-WHG-CHG-Farmer clines in Principle Component Analysis and excluded very old West Eurasian samples and archaics. These were merged with all 503 present-day European 1000 Genomes Project samples to make up our dataset, totalling 1518 diploid samples.

Upon simulation, data for sample individuals is produced, in the form of both VCF files and tree sequence files. We simulated chromosomes of length 200 Mbp with a recombination rate of 1e-8 per bp per generation and neutral mutations dropped onto the branches at a rate of 1.25e-8 per bp per generation. It is also possible to simulate using a real human chromosome recombination map in which case the sequence length is given by the file and a variable recombination rate across the sequence is applied. To manipulate and examine trees in tree sequence files we use the tskit python API.

We performed Principal Component Analysis on the simulated genotype data to compare the structure to that of the real data. Both the MesoNeo chromosomes and nine simulated chromosomes were filtered for minor allele frequency 5%. We used EIGENSOFT smartpca [28] with the outlier removal option disabled to perform PCA with no projection on both the imputed MesoNeo samples together with 1000 Genomes GBR samples and the simulated samples.

In Figure S8 the cluster for each simulated ancient population falls in the same vicinity of PCA space as the real MesoNeo samples and the variation explained by PC1 and PC2 is comparable. Overall, the similarity of the PCA suggests that a lot of the underlying structure that determines how these groups relate to each other is captured by the demographic model.

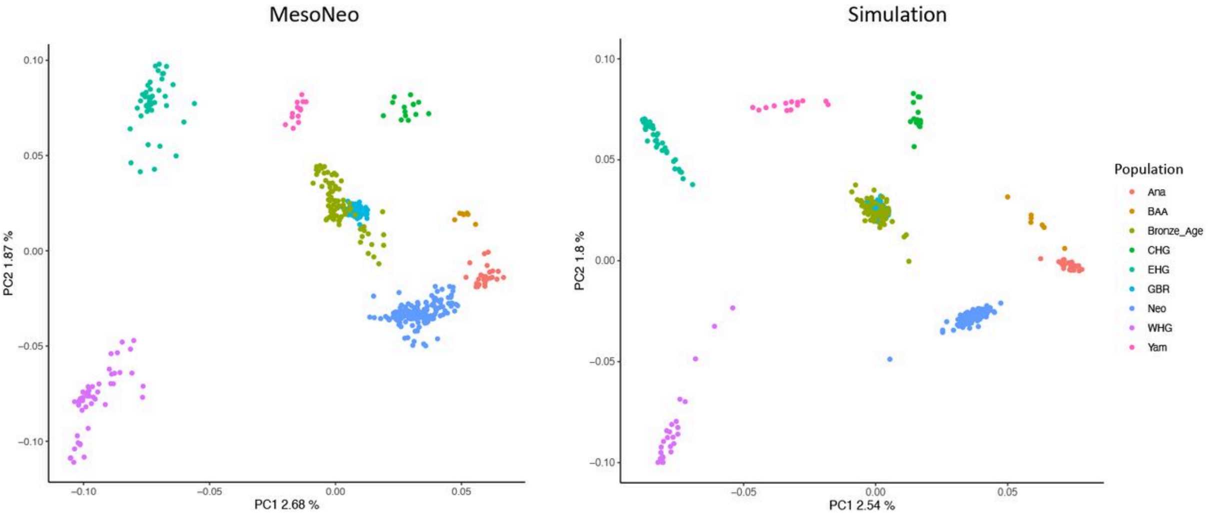


**Figure S8.** Comparison of simulated and MesoNeo PCA.

Weir and Cockerham weighted Fst statistics were calculated over all chromosomes using VCFtools between all pairwise populations. The same was done for the nine simulated sequences. The results shown in Figure S9 display the Fst pairwise values in the simulated data plotted against those in the real data. The correlation of Fst values between the real and simulated data is appreciable at 0.96. This means the demographic model produces data with relative population divergences that are very similar to that in the real data, suggesting the model represents the real population structure well.

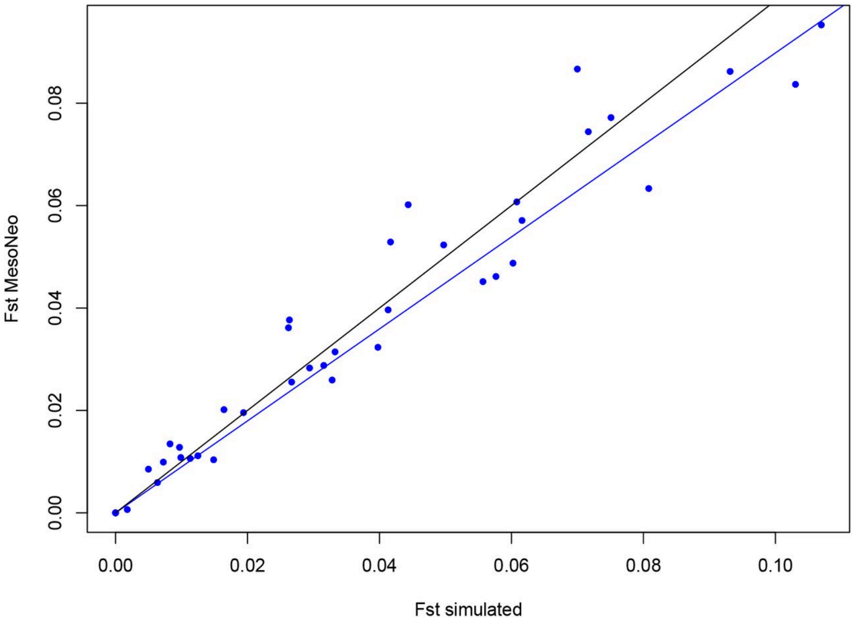


**Figure S9.** Pairwise Fst correlation.

The relationships of these ancient groups and present day Europeans to each other and the split/admixture times are well established and needed no adjustments in the model. However, to produce simulated PCA and Fst results that matched the real data, the population sizes shown in Figure S7 needed some tuning. Both the Fst analysis and PCA final results suggest that, while the model is undoubtedly not entirely correct, it produces simulated data that looks close enough to the true data, by these measurements, that we assume the model represents the real population history of Europeans.

##### A method for estimating local path ancestry

Our method infers the path that segments of a chromosome have taken through a population history. Each segment is covered by a single marginal tree in a tree sequence. Figure S10 depicts how the path ancestry of focal haplotype can be determined from the tree relating that haplotype to all other sample haplotypes: We traverse up the tree from the focal haplotype, jumping to successive parent nodes towards the root i.e parent, grandparent, great-grandparent etc. If we know the population identity of each of the internal nodes traversed to along the way, then we know the path. This information is recorded in tree sequences simulated by msprime and so the path ancestry of haplotypes in simulated tree sequences is straight-forward to find with tskit (https://tskit.dev/tskit/docs/stable/introduction.html).

However in RELATE tree sequences, inferred from simulated or real variant data, the population label of internal nodes is not known. We therefore adapted the concept of Genealogical Nearest Neighbours (GNNs) [29] to identify the population identity of each internal node encountered during traversal from a focal admixed haplotype. This involves recording the proportion of each ancestral group that makes up the sample leaves below each internal node, not including leaves seen at the previous nodes in traversal. The ordered collection of all x GNN distributions of all x nodes examined during a tree traversal reflects the path that the focal haplotype has taken to the root. Figure S10 demonstrates how these ordered GNNs can be used to determine the path.

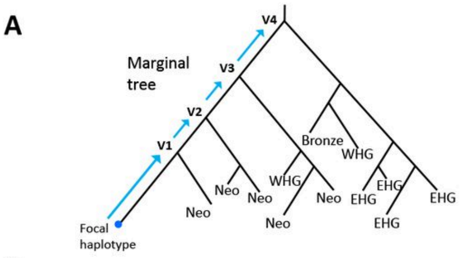


B

| Node | GNN proportions | | | | | | | | |
|---|---|---|---|---|---|---|---|---|---|
| | Bronze age | Yam | Neo | WHG | EHG | CHG | Ana | BAA | Age (normalised) |
| V1 | 0 | 0 | 1 | 0 | 0 | 0 | 0 | 0 | 172/ 1500 |
| V2 | 0 | 0 | 1 | 0 | 0 | 0 | 0 | 0 | 319/ 1500 |
| V3 | 0 | 0 | 2/3 | 1/3 | 0 | 0 | 0 | 0 | 382/ 1500 |
| V4 | 1/6 | 0 | 0 | 1/6 | 4/6 | 0 | 0 | 0 | 603/ 1500 |

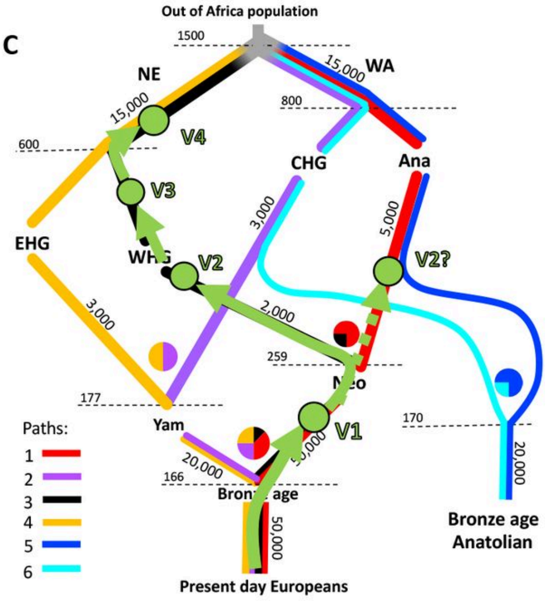


**Figure S10.** Overview of GNN extraction for a European population. (A) shows an example marginal tree that relates the focal haplotype to all other haplotypes in the dataset. Samples are shown by their population labels. (B) shows the GNN matrix determined from the marginal tree, traversing up four nodes towards the root V1-V4. (C) shows how the path can be determined within the model of population history given the GNN matrix by mapping the nodes to the paths.

We need to be able to assign paths to GNNs from millions of sample haplotypes, so we implemented supervised machine learning, specifically a convolutional neural network. First we simulate variant and tree sequence data from a demographic model of the history of the populations under analysis, such as our model of European population structure. Next we infer RELATE tree sequences from the simulated genotype data. We then extract training GNNs from the RELATE trees and train a neural network to predict the path given the true path label extracted from the corresponding simulated tree sequences. By training on GNNs extracted from RELATE trees, we aim to ‘train in’ any bias from the RELATE inference so that when we apply the neural network to tree sequences inferred by RELATE from the real data, these biases are somewhat accounted for.

Training data for the network is the set of GNN distributions for the first five informative nodes traversed towards the root extracted from the RELATE inferred trees. The GNN distributions are configured as a 5 x n matrix, one row per 5 informative nodes examined.

Columns 1–n contain the proportions, between 0 and 1, of leaves belonging to each of n ancient sampled groups and column n +1 contains the age of the node. Informative nodes are those that have at least one leaf from the set of ancient sampled groups. If the root of the tree is reached in less than five informative nodes, then the remaining rows of the matrix are filled with −15 as ‘padding’. The training labels signify the path extracted from the corresponding simulated tree sequences (determined simply using tskit).

The method also works when traversing up to the root and finding the path for ancient sample haplotypes. Therefore, we also train over GNNs extracted from admixed samples, while samples that are diagnostic of one path (e.g., Figure S7 CHG, WHG, EHG and Ana) can be assigned by their population identity alone.

The method performs well for a range of demographic scenarios including variable number of paths and extent of population differentiation, as well as being robust to misspecification in the underlying model (Supplementary Notes 3.2, 3.3).

##### Training and testing a neural network on simulated European genomes

We simulated three 200Mbp tree sequences from our model of European population structure, using different random seeds. RELATE tree sequences were inferred from the corresponding simulated VCF files. Default parameters for RELATE were used; 1.25e-8 mutation rate and starting population size estimate of haplotypes of 30,000.

We extracted 10,000 training pairs of GNNs and true labels from each of the five admixed sample populations (GBR, Bronze Age, Yamnaya, Neolithic farmers and Bronze Age anatolians), 50,000 pairs in total. GNNs were taken from the trees covering evenly spaced sites across three tree sequences output from RELATE, avoiding most of the correlation between trees. True path labels were taken from the trees covering the same sites in the corresponding simulated tree sequences. We trained the classifier to predict the path labels from the GNNs.

For testing, we generated five more simulated and RELATE inferred pairs of tree sequences and tested the classifier on 10,000 GNNs from each to obtain a mean accuracy of 93.12% with a standard deviation of 0.29% across the five tree sequences.

To test the precision of the classifier at identifying each path in each population, we pooled the testing GNNs from all five sequences and applied the classifier to each admixed population separately. Figure S11 shows the confusion matrices for each population, normalised by the sum of the rows to show the precision i.e of the labels assigned a class, what proportion are true positives. The GBR obtains the lowest accuracy. GBR samples have the greatest number of generations between sampling time and admixture time. More generations since the admixture event means more recombination events have broken down admixture LD, resulting in shorter tracts of ancestry. RELATE uses flanking SNPs to calculate the distance matrix and allows some relaxation on the mapping of SNPs when constructing a new tree along the chromosome. These properties make it more difficult to determine the switch of ancestry at the edges of tracts as there is some inertia in the changing of tree topologies compared to the true tree sequences. Shorter tracts produce more edges and therefore less accuracy in the GNN assignment. However, high accuracy is still maintained for all populations, even the GBR.

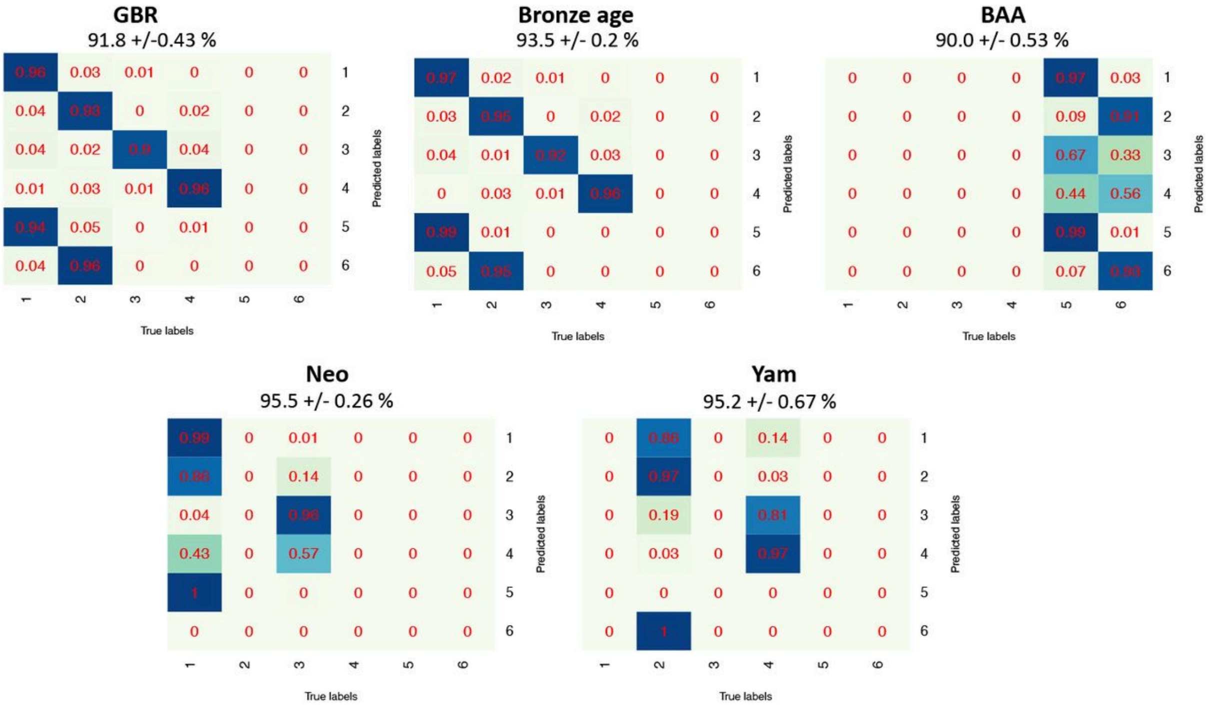


**Figure S11.** Confusion matrices per population when testing on simulated data. Values are normalised by the sum of rows to show the precision values, when testing a classifier trained on the model of European population structure.

Figure S27 shows a classified painted chromosome alongside the true simulated chromosome. Noise appears as short tracts of correlated trees, within larger chunks of ancestry covering many sites in admixture LD.

#### Comparison to an advanced LAI tool

GNOMix is a recent LAI tool that has been shown to outperform previous methods on whole genome data [13]. Like other LAI methods, GNOMix views each reference population as a discrete ancestral population with no awareness of the relationships between reference populations. It therefore is a good tool to compare our method against simulated data of Europeans.

For the GNOMix reference panel we use samples from the four 'path' populations (EHG, WHG, CHG and Ana) as ancestral populations that correspond to our paths 1 to 4 from the model. These are the populations that lie on one path only (Figure S7). For brevity we only test the four admixed populations as query sequences that are characterised by these four paths, GBR, Bronze Age, Neo, and Yam, and do not test the BAA. It should be noted however that the BAA could be tested with GNOMix using the Ana and CHG as ancestral populations which would correspond to our paths 5 and 6.

We simulated five 200Mbp length sequences from the model of European population structure. We extracted ancestry predictions, produced by our method and by GNOMix, from evenly spaced sites across the five sequences. Inference with GNOMix was done using the default logistic regression base and xgboost smoother modules. All other parameters were default including a window size of 0.2cM for Bronze Age, Neo and Yamnaya samples. Because the GBR are 166 generations since admixture we decreased the window size to 0.02cM to account for the smaller admixture tracts. The mean and standard deviation for each admixed population, across the five sequences, obtained by each method were used to perform two-sample T-tests (two-sided) to test for a significant difference in accuracy (Table S1).

**Table 15.** Comparison of accuracy to GNOMix for each of four admixed populations.

| Population | GNOMix accuracy | Ancestral Paths accuracy | T-test (two-sided) p-value |
|---|---|---|---|
| GBR | 74.7 | 91.8 | 5.96E-12 |
| Bronze Age | 87.1 | 93.5 | 1.17E-07 |
| Neolithic farmers | 94.4 | 95.5 | 1.49E-05 |
| Yamnaya | 98.6 | 95.2 | 1.59E-05 |

For the GBR, Bronze Age and Neo our method is significantly more accurate. GNOMix was significantly more accurate for the Yam population. While GNOMix has high accuracies for populations closer to the time of admixture, classification of the present-day GBR samples is much less accurate than our method. Despite reducing the window size for GBR classification, it appears GNOMix struggles with smaller sized ancestry tracts. Our method displays high accuracy for all populations, demonstrating its versatility compared to GNOMix.

#### Results

We inferred a RELATE tree sequence for each autosome of our subset of 476 MesoNeo genomes, using the fine scale recombination map for each chromosome, a mutation rate of 1.25e-8, and starting population size estimate of 30,000. All chromosomes for all samples were painted by the classifier trained on the model of European population structure.

##### Estimation of time since admixture in Europe

We devised a technique to infer the time since admixture of individuals using the painted chromosomes. The method involves fitting exponential decay curves to the probability of being in the same path at two positions separated by varying genetic distances along a chromosome. The time since admixture in generations is extracted from a parameter of the exponential decay function (Supplementary Note 3.4). This process is performed on all autosomes, treating homologous chromosomes independently. We obtain two estimates for each diploid sample, one estimate per ancestry, which are then combined as a weighted average to get an estimate of time since admixture per individual. For samples which consist of multiple paths, we combine the paths that make up each parent population involved in an admixture event, creating larger admixture chunks. For example, the Bronze Age admixture event is estimated from Bronze Age samples by combining paths 2 with 4 and paths 1 with 3 to represent the Yamnaya and Neolithic farmers admixing respectively.

We estimated the time of admixture for each sample in the three admixed MesoNeo populations; Neolithic farmers, Steppe Yamnaya, and the Bronze Age population. The number of genomes over which this was performed was increased from our original 476 to include samples that fall in between groups in the PCA (Figure S26), to explore whether these samples were the result of more recent admixture. The new total number of samples was 963, including all European 1000 Genomes samples. We showed that there was little decrease in accuracy when painting extra samples if we use the same set of 476 diagnostic samples in the GNNs (Supplementary Note 3.3.5).

We calculated a date of admixture as the sample age plus the estimated time since admixture for each individual, combining the standard error of the radiocarbon age estimate and the time since admixture estimate. Given the coordinates of the archaeological sites, we fit spatiotemporal models of how time of admixture depends on latitude and longitude and compared these to the same models constructed using sample age alone.

##### Neolithic farmers

Of the 176 Neolithic farmers individuals in our subset dataset, we were able to estimate the time of admixture for 173. Figure S12 shows the inferred time of admixture in years against sample age. A linear regression to the points fits a model coefficient where every year younger in sample age increases the time since admixture in that sample by 0.28 years (p-value = 2.37e-06). This best fit line has a shallow gradient compared to the theoretical best fit line in which the Neolithic farmers are formed by one instantaneous admixture event happening 7,800 years ago. This suggests the migration of Neolithic farmers and admixture events between Neolithic farmers and WHG was less punctuated and more of a continuous process, ranging between 8,000 and 4,000 years ago.

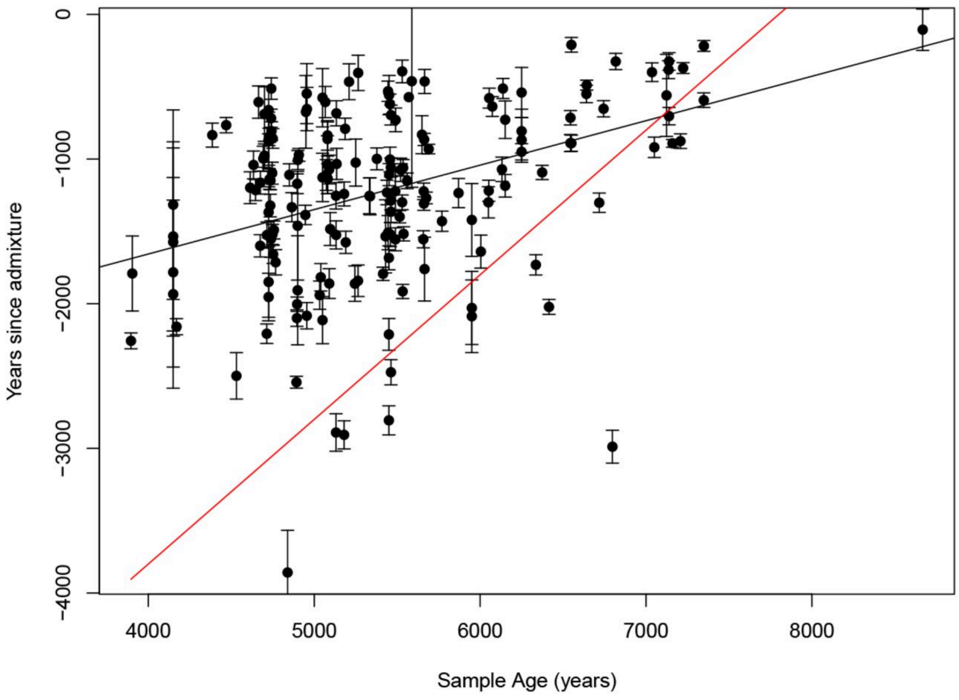

**Figure S12.** Correlation of years since admixture and sample age in Neolithic farmer samples (n=173). The best fit line is coloured black. The theoretical best fit line, in the scenario of instantaneous admixture happening at 7,800 years ago, is coloured red. Error bars show the combined standard error of the radiocarbon age estimate and the time since admixture estimate.

Two theories exist for the route that Anatolian farmer ancestry took from its origin in Anatolian into the European continent. One is a path along the northern Mediterranean coast to Iberia first before expanding north and the second is an inland route following the Danube River, north and west.

We fit two linear models of Inferred admixture time ~ longitude + latitude and Sample age ~ longitude + latitude (Table 23) which showed longitude and latitude are both highly significant predictors of admixture date. The model has a significant overall p-value of 5.947e-11 and explains 23.4% of the variance in the data. The predicted coefficients of longitude and latitude suggest that the admixture events occur 25 years later with every degree east and 42 years later with every degree north. Similar results are shown in Figure S13 for which we performed linear interpolation of admixture time between the points of inferred admixture time of individual samples by kriging across Europe. The pattern of admixture of Anatolian farmers with WHG with time starts in Iberia and moves north east across Europe, implying the coastal route.

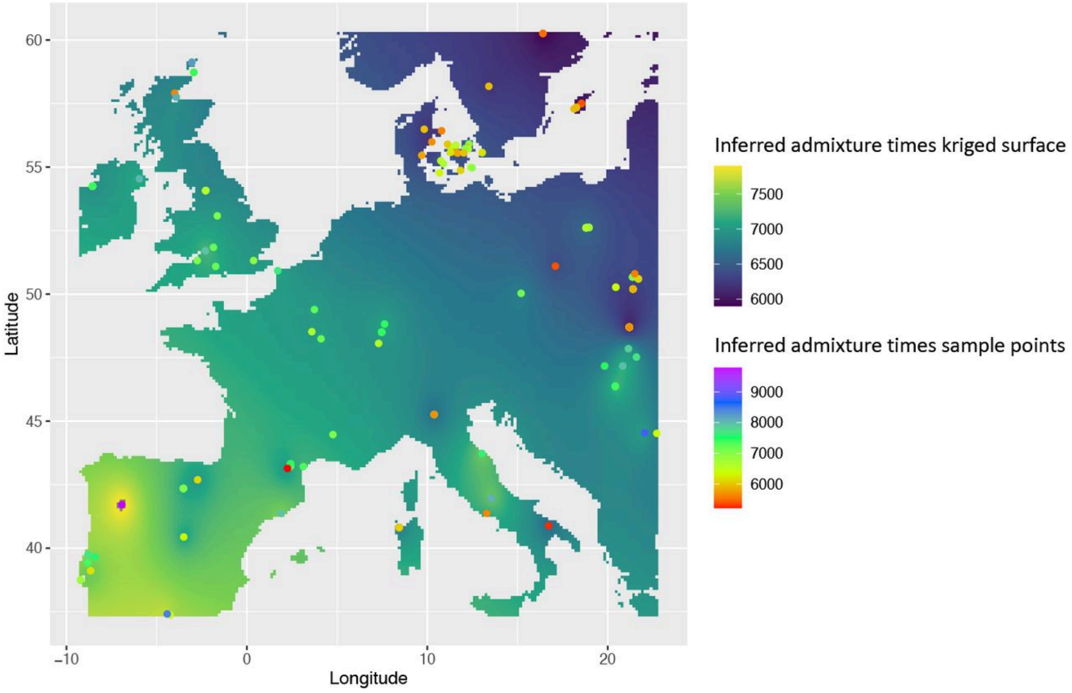

**Figure S13.** Linear interpolation of admixture time of Neolithic farmers across Europe. Map of Europe with points to show the archaeological position of Neolithic farmer samples coloured by the inferred admixture time in years before present. The surface colour is admixture time in years before present created by linear interpolation between inferred sample points.

The MesoNeo dataset contains few samples of Neolithic farmers from along the Mediterranean coast east of Spain, due to low sample availability, or perhaps poor preservation, and so we may not be capturing the earliest admixed samples. Alternatively, the first migrants moving along the coast may not have admixed with local hunter gatherer populations until they reached Iberia. Both scenarios would be consistent with our results and in the future, more samples from critical regions could help provide clarity.

In contrast, while longitude and latitude are also significant predictors of sample ages (Table 23), the variance explained by such a model is only 8.4%. The model suggests a more gentle southeast to northwest gradient of older to younger sample ages. Overall, a model involving longitude and latitude to predict inferred admixture time is more informative than one predicting sample age, in terms of understanding the impact of Anatolian farmers in Europe.

##### Bronze Age population

The migration of the Yamnaya associated ancestry into Europe is typically thought to have been a fast migration, possibly accompanied by violence [30] and substantial alteration of the environment [31].

Figure S14 shows the inferred time since admixture in years of the Yamnaya (path 2 and 4) and Neolithic farmers (paths 1 and 3) plotted against sample age. Of the 105 Bronze Age individuals in our MesoNeo subset, exponential curves could be fit to 97. A linear regression to these points implies that every year younger in sample archaeological age, the time since admixture in that sample increases by 0.78 years (p-value = 1.536e-11). A coefficient value of 1 would mean contemporaneous admixture over the whole continent, as shown by the theoretical best fit line in red. Such a high coefficient found for the Bronze Age admixture is consistent with a rapid migration of the Yamnaya across the continent between 4000 and 5500 years ago, with admixture events quickly following.

Moreover, linear models incorporating longitude and latitude do not find either as a significant coefficient (Table S10). This is supportive evidence of rapid movement of Yamnaya into Europe from the steppe with admixture following soon after the migration started, and then the admixture continuing with admixed individuals, producing no detectable variance in inferred admixture time with geography. The variance in the data explained by a model of longitude and latitude alone is small at only 4.9%. While this value for the inferred admixture times is small, models using sample ages alone explain the data variance even less well at only 2.1%.

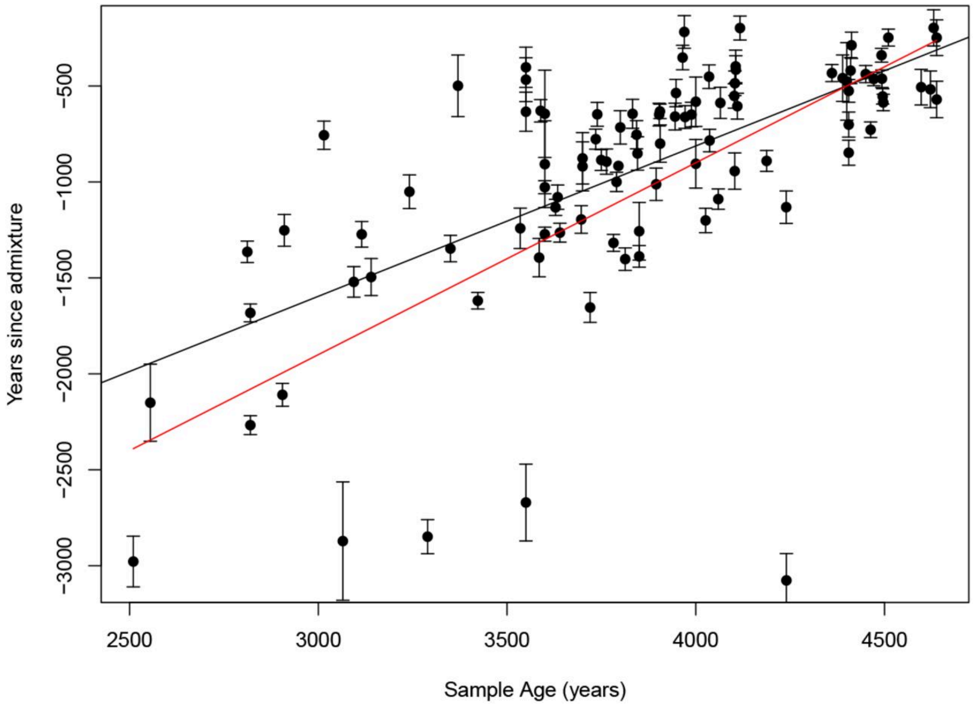


**Figure S14.** Correlation of years since admixture and sample age in Bronze Age samples (n=97). The best fit line is coloured black. The theoretical best fit line, in the scenario of instantaneous admixture happening at 4,900 years ago, is coloured red. Error bars show the combined standard error of the radiocarbon age estimate and the time since admixture estimate.

#### Steppe Yamnaya

The MesoNeo dataset provides more Yamnaya related individuals than were previously available, adding more data points for dating the admixture event between EHG and CHG groups. While this is a substantial boost in sample size, there are still relatively few samples and therefore not enough power to detect any trends in admixture time with sample age or geography. This is compounded by the samples being from disparate geographic locations in Ukraine, Poland, Kazakhstan and across Russia.

Despite this, we obtained admixture time estimates from 16 of the 17 Yamnaya samples present in our MesoNeo subset (Figure S15). The genetic formation of the Yamnaya population is not well understood. Culturally they appear in the archaeological records 3300-2600 BC [32]. Most of our estimated admixture dates are 5000-6000 years before present, a millennium or so before their believed cultural formation. Three outlier samples have inferred admixture times of more than 7000 years ago. These all have large standard errors on the estimates and so are less reliable. However, even allowing for larger confidence intervals based on higher standard errors, their admixture times are placed well before 6,000 years before present.

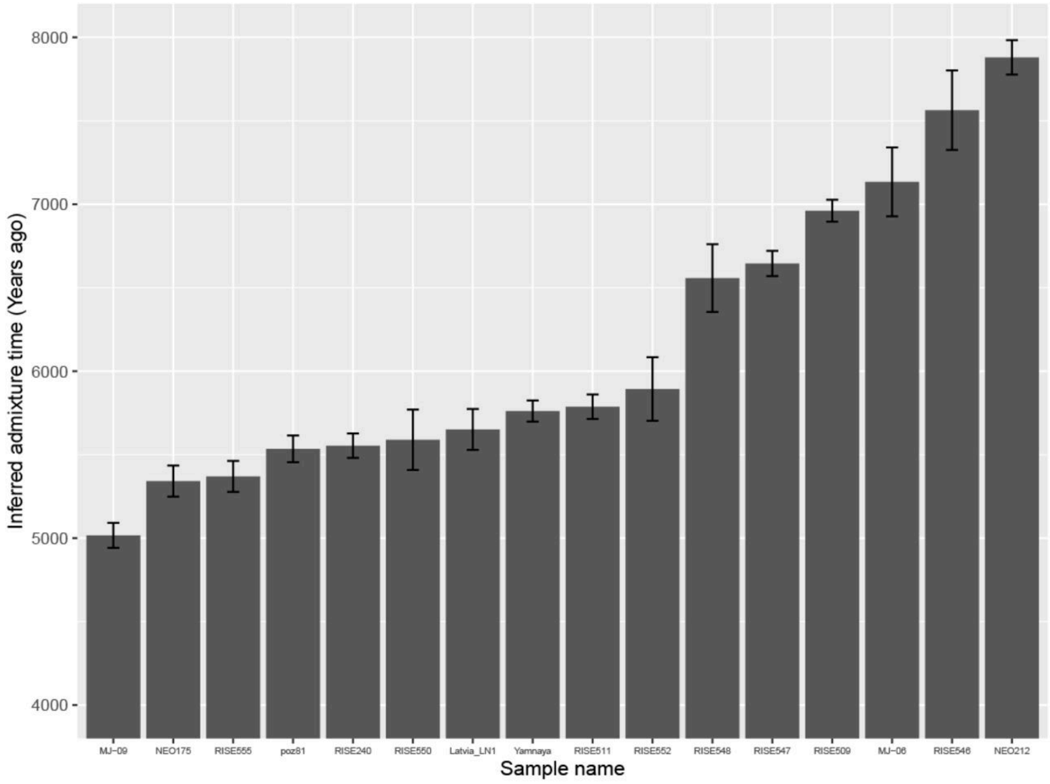


**Figure S15.** Inferred admixture times for Yamnaya samples (n=16). Bar plot of the inferred admixture times for Yamnaya samples, with error bars showing the standard error around weighted mean estimates.

These admixture dates are much older than the archaeological appearance of the Yamnaya and so, subject to technical artefacts they suggest extended prior genetic contact between

EHG and CHG well before the emergence of the Yamnaya culture, potentially as early as 7000 years ago.

#### Discussion

Here we have presented a method for inferring local ancestry in ancient and modern genomes given an explicit model population history. The redefinition of local ancestry as a path, rather than a static identity, back in time through various populations is more appropriate to complex population histories. It is becoming more apparent, especially from ancient DNA analysis, that the histories of human populations across the globe are characterised by multiple population split, admixture, migration and isolation events and an appropriate analogy is a braided river rather than a tree. Hence, with the growing amount of ancient DNA data available, the path concept of ancestry is becoming more viable with and applicable to populations other than Europeans. We suggest that it is an important step to begin thinking of ancestry in this way rather than in terms of static identity. It is this reframing of the meaning of ancestry which separates our method from pre-existing local ancestry inference tools.

We have used a highly interpretable machine learning framework to perform a process that could be done manually. This helps to avoid the pitfalls of a 'black box' where it would be difficult to know which basic features are driving classification. Our method performs as well as a leading local ancestry inference tool, GNOMix, for populations that are close to the time of admixture and performs better for populations with more time since admixture.

The method has its foundations the ability of RELATE to correctly construct a tree sequence. In essence we are embedding a tree sequence within a population structure and thereby assuming that the trees fit within the model constraints and reflect the underlying population structure. There are two reasons why this may not be the case: Firstly, any model of the past demographic structure of a population will be inaccurate. In most cases the structure will be simplified to represent only major population events and ignore smaller scale migrations. For some populations, there may be no pre-existing knowledge of the population structure and results from PCA and ADMIXTURE analysis may be unclear. While we have shown in Supplementary Note 3.3 that the neural network can generalise over misspecification in the model, but with too little understanding of the history we may not obtain results of much confidence or meaning. Secondly, RELATE may not infer the tree sequences correctly, despite the model representing the true structure well. We have attempted to compensate for biases in the RELATE inference by training the classifier with GNNs extracted from RELATE inferred trees paired with labels from the true simulated trees. Results on simulations indicate this is successful.

We applied our method to a large imputed dataset of ancient European and West Asian genomes and used a subsequent technique to infer the time since admixture of ancient individuals. Imputation borrows information from closely related, higher coverage individuals, making samples that coalesce most recently with each other, more similar to each other than their true sequences actually are. Intuitively this effect should help the topology building process of RELATE by helping identification of nearest neighbours. Conversely, there is no way for imputation to recover variants that are private to ancient groups and so not present in the reference panel which will hamper neighbour identification in ancient individuals. Overall, we suspect the effect of imputation on the accuracy of inferring tree topologies is minimal, however pervasive imputation errors are a topic of research [33] and further analysis is needed to evaluate how much these errors might be affecting RELATE inference.

Additionally, in the local ancestry painting phase errors will create switches of ancestry across homologous chromosomes. As a result, phase errors will cause ancestral tracts to appear smaller than in truth and therefore produce admixture times that are older than in truth. It is possible that our calculated admixtures times have overestimated absolute times as we have not accounted for phase error. A future strategy to implement that accounts for phase errors is to use both haplotypes from an admixed sample when recording the changes in ancestry at increasing genetic distance and not treat the two chromosomes from the same sample independently. Switches in ancestry across the homologous chromosomes will be accounted for by measuring over both haplotypes.

However, our conclusions drawn from the relative admixture times between samples are still valid, revealing spatiotemporal patterns of admixture of different populations in Europe. We showed how the inferred admixture date is superior for identifying these patterns than using the archaeological sample age alone. Results from Neolithic farmer genomes suggest that the movement of Anatolian farmers into Europe and their subsequent admixture with WHG was slow and can be explained by geography to a large extent. Admixture appears to occur first in Iberia and moves north west over time, potentially supporting a route of Anatolian farmers along the north Mediterranean coast into Europe. In contrast, results from Bronze Age genomes suggest that the movement of steppe Yamnaya into Europe was fast with admixture occurring soon after migration. We found that the genetic formation of the

Yamnaya is 5000-6000 years before present, approximately a millennium before their believed cultural formation determined from archaeology which is consistent with other recent results [34]. We also show evidence of potential contact between the EHG and CHG in the eighth millennium before present.

It is important to note that admixture times are not always a proxy for movement of people. It is possible that the Anatolian Farmers moved at a faster pace into Europe, but the subsequent integration with the local hunter gatherer populations was a slower process that continued long after migrants arrived in an area. There is evidence of persistent un-admixed hunter gatherer pockets existing long after the arrival of Anatolian migrants to the same area which is seen in parts of the World today, such as the Hadza in East Africa.

For populations where the historical structure is well understood, the method can be adapted to these populations. For those where a demographic model from which to simulate is not available, one must be custom built, although the demographies of many species and human populations are now available in the stdpopsim catalogue (https://github.com/popsim-consortium/stdpopsim/blob/main/stdpopsim/catalog/HomSap/demographic_models.py).

#### Supplementary Information

Code is available at the following link: https://github.com/AliPearson/AncestralPaths

##### 3.1 Excluding Bronze Age Anatolians

Our model of European population structure includes the Bronze Age Anatolians (BAA) from which paths 5 and 6 lead. Given these paths are not directly relevant to the history of present day Europeans, we tested how the accuracy of classification is altered when the BAA are removed from the GNN distributions and paths 5 and 6 are removed as labels. This leaves a four path model and a reduced GNN matrix size over which to train a neural network. We tested the accuracy of this classifier in each of the 4 path classes, averaged over 5 testing tree sequences, in the GBR population.

The overall accuracy across the four paths in the model with no BAA is 94.3% +/- 0.27%. This is significantly greater than the overall accuracy in the model containing BAA when taken over all six paths in that model (p-value = 1.473e-05). However, when averaging over just the four non-BAA paths in the model containing BAA we obtain an accuracy of 94.6% +

0.46%, which is not significantly different to the classifier trained on the model excluding BAA (p-value = 0.0918).

This result can be seen in Figure S16 where the precision in each of the four non-BAA paths in the model including BAA is very similar to those in the model excluding BAA.

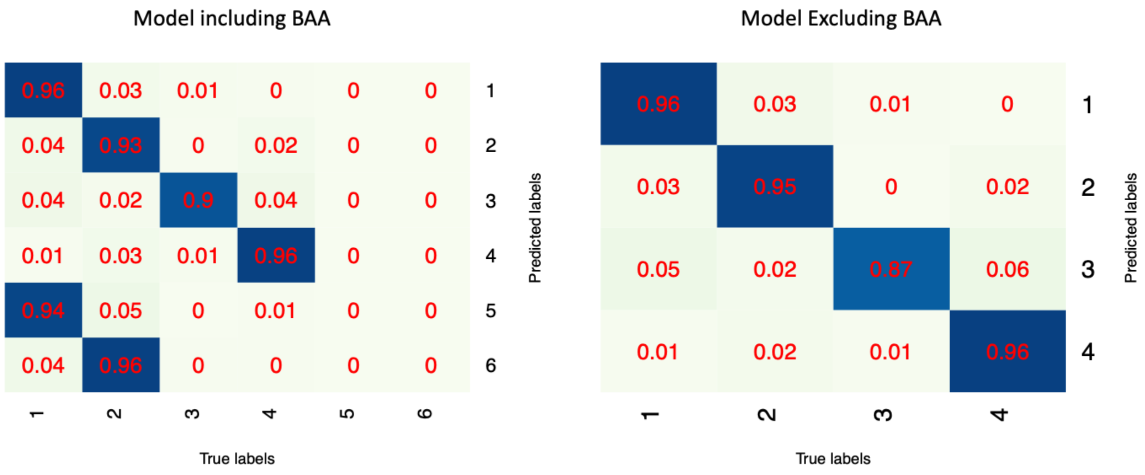


**Figure S16.** Confusion matrices showing the precision in each path tested over GBR samples in 5 testing tree sequences. The classifier trained on the model including BAA can predict one of six paths, while the classifier trained on the model excluding BAA can predict one of four paths.

This result demonstrates that accessory paths such as those leading from the BAA in our model of European population structure can be added with no detriment to precision of other paths. Although the overall accuracy across all paths is lower when the BAA are included, the accuracy is still over 90%.

#### 3.2 Conditions affecting classifier performance

The ”path ancestry” inference approach outlined above is applicable to many populations of humans and other species. To help decide what types of populations and time resolutions this method would be appropriate for we assessed its performance under a variety of demographic scenarios. We simulated data from a range of demography models, systematically varying features and parameters and we measured the overall classifier accuracy and precision in each class when tested on simulated data.

Unless otherwise specified, we simulated tree sequences of 20 Mbp in length with 1.25e-8 mutation rate and constant recombination rate of 1e-8. All admixture events involve equal proportions contributed by each participating population. The ordering of population split and admixture events is determined randomly by sampling from the active populations. The timing of split events is three generations after the previous split event while the timing of admixture events is fifteen generations following the previous admixture event. All populations have an effective population size of 10,000. Twenty five diploid samples were taken from all the 'path' populations that are present after all split events have occurred and also from all admixed populations. The sampling time for each population was sampled uniformly from within the time each population was active and all samples per population were taken at the same time. An example is shown in Figure S17.

Training GNNs and label pairs were extracted from all admixed populations. To gather the same number of pairs for each path in each admixed population from sufficiently spaced sites, we simulated five training tree sequences and five testing sequences. The largest source of variation is across tree sequences so testing was performed on each sequence separately and a mean and standard deviation across sequences was calculated. Two sample t-tests were then performed to assess if there was a significant difference in classifier accuracy when a parameter was changed. We then pooled the testing GNNs and applied the classifiers to produce confusion matrices and calculate precision values in each class.

##### 3.2.1 Inference with varying path number and divergence time

We tested how the number of paths in the model and how much differentiation between the 'path' populations affected the classifier's ability. This was to explore 1. how complex the demographic history could be for the classifier to tease apart separate paths and 2. how applicable the method is to more finely structured populations with recent admixture compared to populations with deep structure.

We simulated demographic models that contained two, four, six and eight paths. All demographic models started with a single trunk population that, through binary population splits, divides into several populations corresponding to the number of paths. These populations remain separate for around a specified number of generations (10, 50, 100, 500, 1500 or 3500) before admixing successively with each other until one population remains (Figure S17). The same random seed was used to simulate models of the same path number, so all models with the same path number but different divergence times had the same ordering of population splits and admixtures.

Figure S18 shows the accuracy and standard deviation for classifiers trained on all demographic combinations of path number and separation times. The accuracy decreases as the number of paths in the model increases and as the number of generations that all the paths are diverged decreases. All path numbers show a rapid increase in accuracy from 100 to 500 generations of divergence. The more generations that the ‘path’ populations are separate, the larger the allele frequency differences become between paths for pre-existing variants due to drift acting for a longer period of time. Additionally, more mutations accumulate that differentiate paths while paths are separated. More mutations and greater allele frequency differences means that RELATE is better able to resolve the correct topologies, which results in the GNNs looking more consistent within each class.

More paths increases the number of classes that the classifier must differentiate between and therefore the opportunity to confuse between those classes. Models with more paths will require more time of divergence to achieve the same accuracy as models with fewer paths. Models with only two paths maintain accuracies of above 50% with as few as ten generations of separation time. By 3500 generations (~100,000 years) of divergence, models containing two, four and six paths have accuracies above 90% and overlapping error bars, showing that with enough time for divergence the decrease in accuracy due to path number can be mitigated.

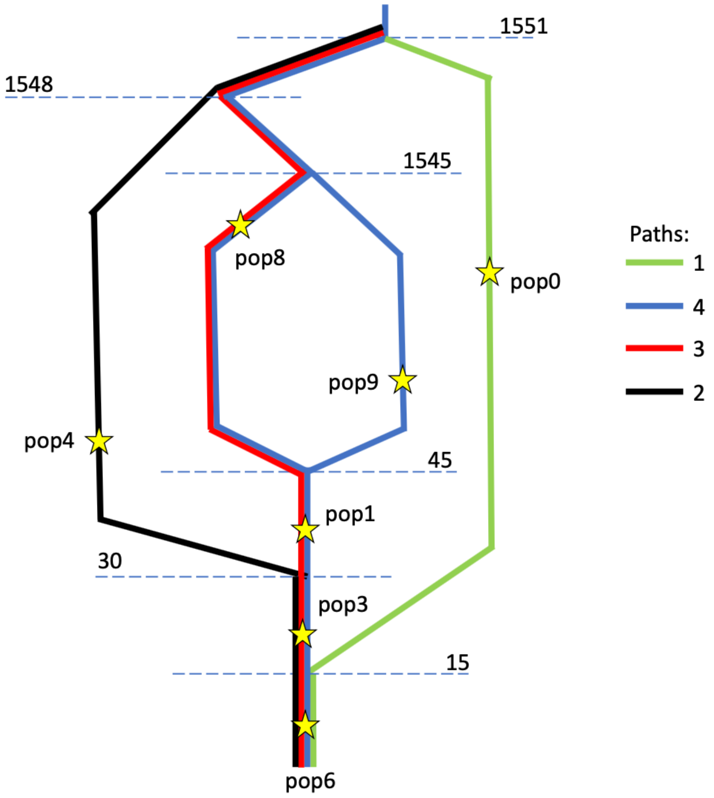


**Figure S17.** An example of a four path demography that was simulated for testing. Stars indicate approximately where in time 25 diploid samples are taken. The populations and paths are arbitrarily numbered. Population split and admixture times are shown on the dotted lines in units of generations ago. This example has a ‘path’ divergence time of 1500 generations, the time period between the last population split and the first admixture event.

Overall, models with smaller path numbers are better suited when there is very fine scale, recent structure involving closely related populations. Likewise, when the populations are very diverged and deep structure is present, models containing more paths are viable.

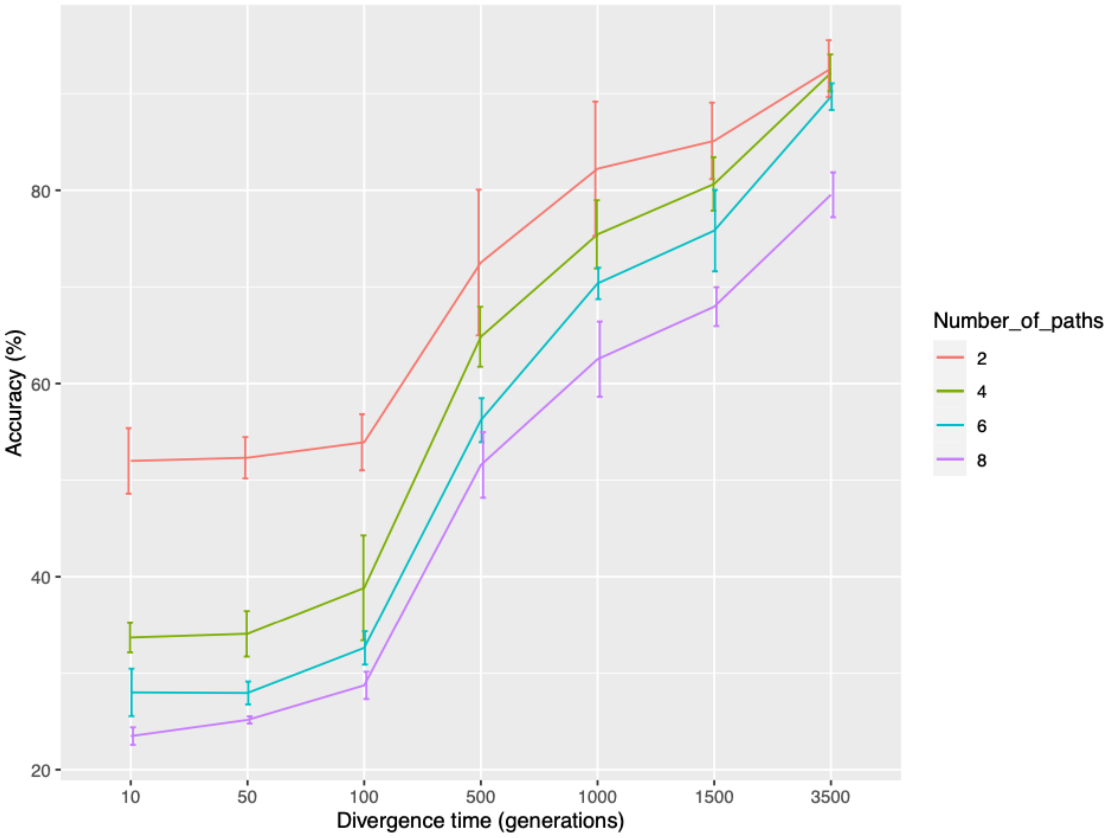


**Figure S18.** Plot showing the mean and standard deviation of accuracy of classifiers trained on models with different path numbers and 'path' population divergence times.

##### 3.2.2 Inference with imbalanced population size

Next, we tested how imbalance in population size on paths affects classification. We used a demographic model of four paths and 1500 generations of separation between 'path' populations. One 'path' population was chosen to have a population size of 50,000 from its emergence after a split to its disappearance after an admixture. All other population sizes were 10,000. A control set of tree sequences were simulated with the same population split and admixture events but with all population sizes set at 10,000.

Compared to the control demographic with an accuracy of 80.66% +/- 2.48%, the classifier trained on the imbalanced population size demographic demonstrates a significantly lower accuracy of 73.45% +/- 0.71% (two sample t-test p=0.0033). Comparing the confusion matrices (Figure S19), path 3, containing the differently sized population, exhibits the drop in precision and is mostly confused with path 4. This behaviour is not unexpected given paths 3 and 4 are sister paths, descending from a common population that split (Figure S17). A larger population size will reduce the number of coalescence events occurring around the time of higher population size compared to the other paths, pushing coalescences into older time periods before paths 3 and 4 separated. Many GNNs for path 3 will not have a coalescence event falling in the period of higher population size, making them look like path 4 or other GNNs and so are misclassified.

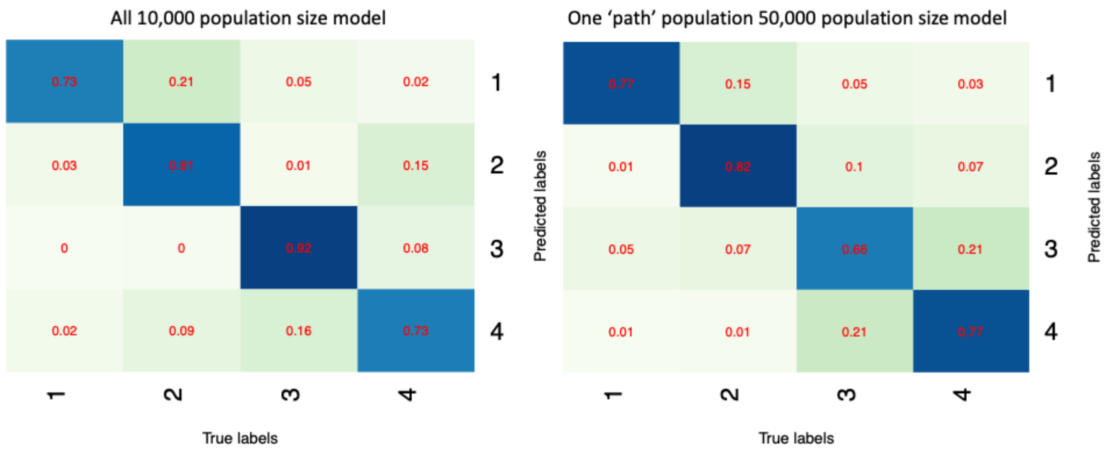


**Figure S19.** Confusion matrices of the model with all populations with size 10,000 compared to the model where one population has size 50,000.

##### 3.2.3 Inference with imbalanced sampling

It is characteristic of ancient DNA datasets that some populations may only be represented by a few samples, while others many. Rather than subsetting some groups to match the sample size of the groups with the fewest samples, we tested if an imbalanced number of samples taken from each group alters the classification of certain paths. We simulated from three demographies, with a divergence time of 1,500 and four paths, and trained a classifier on each. In the control demographic model, the sample size was 25 diploids for all 'path' and admixed populations. In the other demographic model, one 'path' population sample size was either reduced to 5 diploids or increased to 50 diploids. The same order and timing of split and admixture times were used, and all population sizes were 10,000. Path 1 contained the population with variable samples.

**Table 16.** Table displaying the mean and standard deviation of accuracy of classifiers trained on models with variable number of samples taken from a population along path 1.

| Number of samples from path 1 population | Mean accuracy (%) | Accuracy standard deviation (%) |
|---|---|---|
| 5 | 80.22 | 4.48 |
| 25 | 80.66 | 2.77 |
| 50 | 82.49 | 2.65 |

There was no significant difference in overall accuracy of the classifiers when tested pairwise with two sample t-tests (p-values = 0.318, 0.856, 0.364). Table 16 shows the accuracies for each model and the standard deviations. The model with 5 sampled diploids on path 1 has the highest standard deviation. This is likely because there are fewer examples of path 1 labels from those samples in the training data and so the training has not captured as much of the variance of path 1 GNNs as other classes when training the classifier. Testing that classifier, this translates to greater variance in the accuracy.

In the confusion matrices (Figure S20) path 1 has greater precision in the 50 diploid model than the other two models, which have comparable precision values in path 1. However, the overall mean accuracies are not significantly different suggesting there is no systematic bias due to imbalanced sample sizes, rather an increase in precision due to greater sample size on one path.

##### 3.2.4 Inference and overall sample size

Lastly, we tested the effect of overall sample size on classification ability. We simulated from seven different demographic models with a divergence time of 1500 and four paths and trained a classifier to each. Each had a different number of diploids sampled evenly from the admixed and 'path' populations: 5, 10, 15, 20, 25, 30 or 50 diploids. The same order and timing of split and admixture times were used, and all population sizes were 10,000. The same number of training GNNs were used to train classifiers for each demography so as not to confound results with different amounts of training data.

Figure S21 shows how the mean accuracy increases as the number of diploids taken from all sampled populations increases. There is a rapid increase in accuracy between 5 and 20 diploids, after which the increase in accuracy with more samples is less. At 50 diploids the accuracy is around 90% and the standard deviation is low. Even with only 10 samples, with a 1500 generation diverged four path demography, the accuracies can be over 70%.

More samples means more variation in the training GNNs, given that we used the same number of training GNNs from each demography. This prevents overfitting of the neural network allowing for greater flexibility to novel testing GNNs. This results in greater accuracy and smaller standard deviations upon testing.

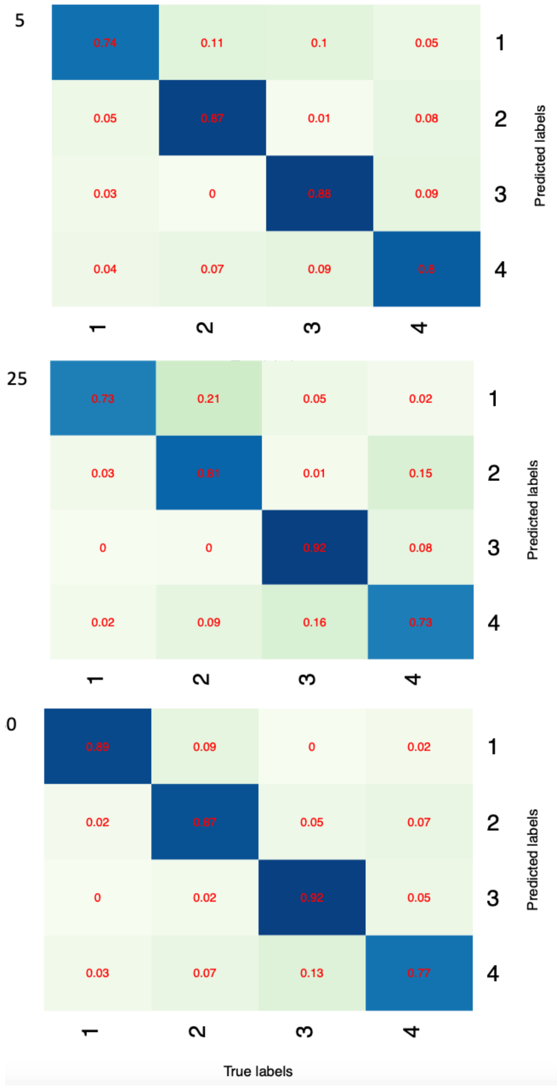


**Figure S20.** Confusion matrices of models with different numbers of diploids sampled from a path1 population.

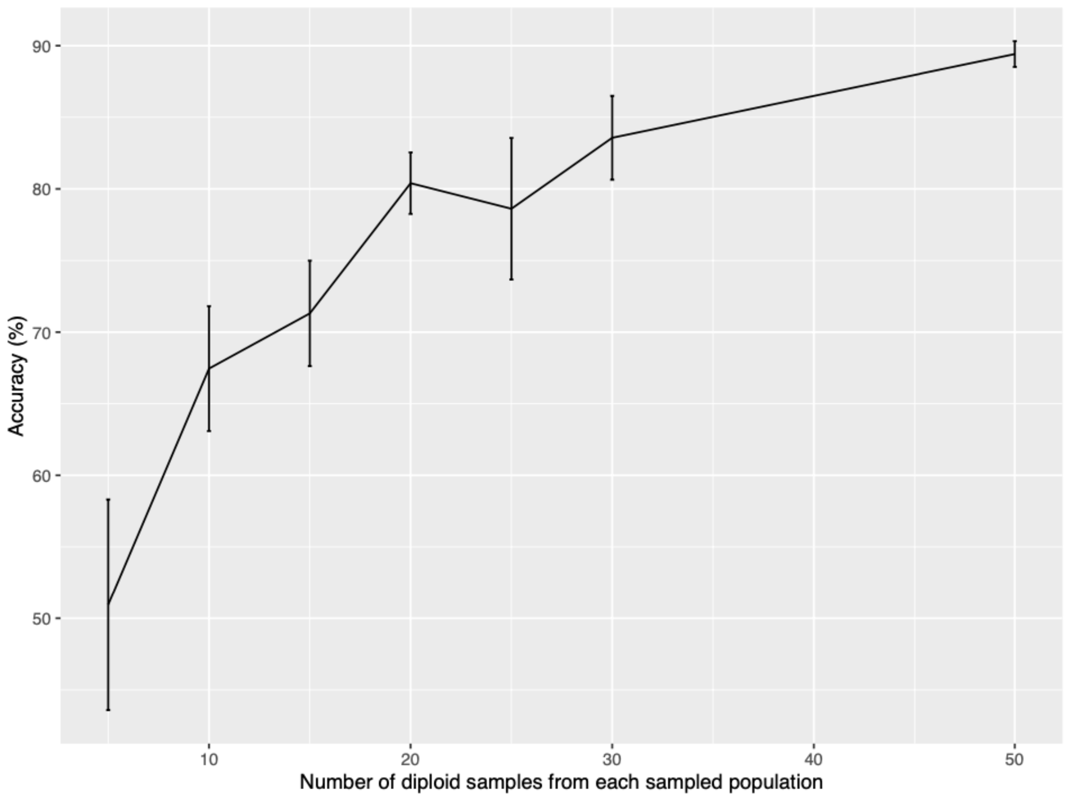


**Figure S21.** Change in accuracy as the number of diploids sampled from all admixed populations increases. Error bars show the standard deviation from the mean, over five testing tree sequences.

##### 3.3 Testing model misspecification

The neural network for determining European local ancestry is trained on simulated data that we believe to match the real data well enough. However, the true history of any population is never known exactly, and the following section explores how a classifier copes when it is trained on data that does not match the testing data in various ways. For all investigations we used demographies with four paths and 1500 generations of path separation time. Except where specified, simulations were carried out in the same way as described in Supplementary Note 3.2.

For each parameter under scrutiny, we simulated two datasets with only the parameter under scrutiny differing between the two, and trained a classifier on each. Testing within the datasets was performed as in Supplementary Note 3.2. When testing between datasets, the classifier of one was applied to each of the five testing tree sequences from the other dataset. This results in a mean accuracy and standard deviation for all four combinations of classifiers and testing data, to which we applied two sample t-tests to determine whether there is a significant difference in classifier accuracy when applied to testing data that does not match the training data. The two classifiers were then applied to pooled testing data in all four combinations to produce confusion matrices.

##### 3.3.1 Inference with different population size

Using the same simulations and classifiers as those used for testing an imbalance in population size along paths, we tested the two classifiers trained on a 10k model and an imbalanced 50k model on GNNs extracted from the alternative demography. A difference in population size in a ‘path’ population will change the GNN structure, where a smaller population size will produce more coalescence events in the path population and a larger population size will conversely produce fewer coalescences. A difference in either direction in the GNN structure between training and testing data will result in the paths being less easy to recognise by the classifier. Predictably there is a significant decrease in accuracy when test data is classed by the classifier trained on different training data to when classed by the classifier trained on the corresponding training data (Table 17). Two sample t-tests produce p-values of 0.007 and 0.0067.

There is no significant decrease in accuracy when the 50k classifier is applied to 10k data compared to 50k data (p-value = 0.418). However, there is a significant decrease in accuracy (p-value = 9.661e-05) when the 10k classifier is applied to 50k data compared to 10k data. An unexpected deficit of coalescences along a path is more confusing to a classifier than a surplus, i.e., the absence of a defining GNN is more detrimental than the presence of an extra GNN.

**Table 17.** Table displaying the mean and standard deviation of accuracy of classifiers trained on models with the same or different population size than the testing data, along path 3.

| Testing data<br>Classifier | 10k all | 50k imbalanced |
|---|---|---|
| 10k all | 80.66. +/- 2.77 | 68.95 +/- 2.12 |
| 50k imbalanced | 74.52 +/- 2.61 | 73.45 +/- 0.71 |

##### 3.3.2 Inference with different admixture times

To explore how the classifier generalises to data with different admixture times, we simulated from one demographic model with admixture events occurring every 15 generations back in time from the present day and another every 30 generations. So, the latter did not result in a smaller time while the 'path' populations were diverged, which would confound the results, we increased the time of the most recent population split in that demographic model.

Table 18 shows that the accuracy for the model with 30 generations separating admixture events is not significantly different from the accuracy of the model with 15 generations, when tested on their corresponding testing data and trained classifier (p-value = 0.976). When we swap the classifiers, to test the 30 generation separated testing data with a 15 generation separated classifier and vice versa, there is no significant change in accuracy in either case compared to testing corresponding data and classifiers (p-values = 0.977, 0.716). The classifiers are able to generalise and compensate for the difference in admixture times. This indicates that, despite providing the node ages in the GNNs, the classifiers are largely using the topology of trees and not the coalescence times.

**Table 18.** Table displaying the mean and standard deviation of accuracy of classifiers trained on models with the same or different admixture times than the testing data.

| Testing data / Classifier | 30 gen. admixture | 15 gen. admixture |
|---|---|---|
| 30 gen. admixture | 80.58 +/- 4.81 | 81.27 +/- 2.33 |
| 15 gen. admixture | 80.49 +/- 4.65 | 80.66 +/- 2.77 |

##### S1cS3.3 Inference with different admixture fractions

We next investigated how different admixture fractions would change the classification accuracy. We simulated a dataset with all admixture fractions 50/50 and another with all 25/75. There is no significant change in accuracy when the 50/50 classifier is applied to 25/75 testing data compared to testing the corresponding 25/75 data and classifier (p-value = 0.393). Neither is there a significant change in accuracy when the 25/75 classifier is applied to 50/50 data compared to testing the corresponding 50/50 data and classifier (p-value = 0.589). This suggests that classifiers are able to generalise when the admixture fractions are misspecified.

##### 3.3.4 Inference when samples are drifted from ancestral populations

For all models we have been simulating samples that are taken directly from the simulated populations. The ancient samples we have in the MesoNeo dataset are unlikely to be individuals from the ancestral populations involved in the admixture events themselves, but instead more or less closely related to them. To investigate the effect of drift between the true ancestral admixing populations and sampled populations, we simulated population splits so that the ancient samples were taken from ‘hanging branches’, slightly diverged from the admixing lineages. A separation time of ten generations from the true ancestral populations simulates approximately 300 years of drift. We trained classifiers using a demographic model where the ancient samples are taken directly from the ancestral admixing populations and tested its performance on GNNs from the drifted model.

**Table 19.** Table displaying the mean and standard deviation of accuracy of classifiers trained on models with the same or different admixture fractions than the testing data.

| Classifier \ Testing data | 50/50 | 25/75 |
|---|---|---|
| 50/50 | 80.66 +/- 2.77 | 85.53 +/- 1.95 |
| 25/75 | 79.58 +/- 3.27 | 86.82 +/- 2.51 |

Table 20 shows the accuracy results when classifiers are testing on different testing data. There is no significant change in accuracy between any two pairs of testing data and classifier combinations demonstrating that classifiers are able to generalise between drifted and directly sampled models (p-values = 0.106, 0.314, 0.634, 0.320, 0.733, 0.122).

**Table 20.** Table displaying the mean and standard deviation of accuracy of classifiers trained on models with ancient samples that were drifted or not from the true ancestral populations.

| Classifier \ Testing data | Drifted | Not drifted |
|---|---|---|
| Drifted | 83.28 +/- 1.44 | 84.60 +/- 0.78 |
| Not drifted | 81.51 +/- 3.32 | 82.63 +/- 3.79 |

##### 3.3.5 Inference for additional samples

The number of ancient samples that are available continues to increase and so more samples relevant to population histories can be incorporated into datasets. Likewise, samples may be removed from datasets after more stringent filtering. To test whether classifiers can generalise to more or fewer samples, we applied a classifier trained on a simulation with 20 diploids taken from all sampled populations to testing data from a simulation with a different number of diploids per sampled population (5, 10, 15, 25, 30, 50). GNNs were extracted using all samples in the demography, as if testing a corresponding classifier.

Figure S22 shows that there is no significant difference when using the classifier trained on 20 diploids compared to the corresponding classifier for testing data containing 5, 10, 15, 25 and 30 diploids (p-value>0.05). For data containing 50 diploids there is a marginally significant decrease in accuracy using the 20 diploid classifier.

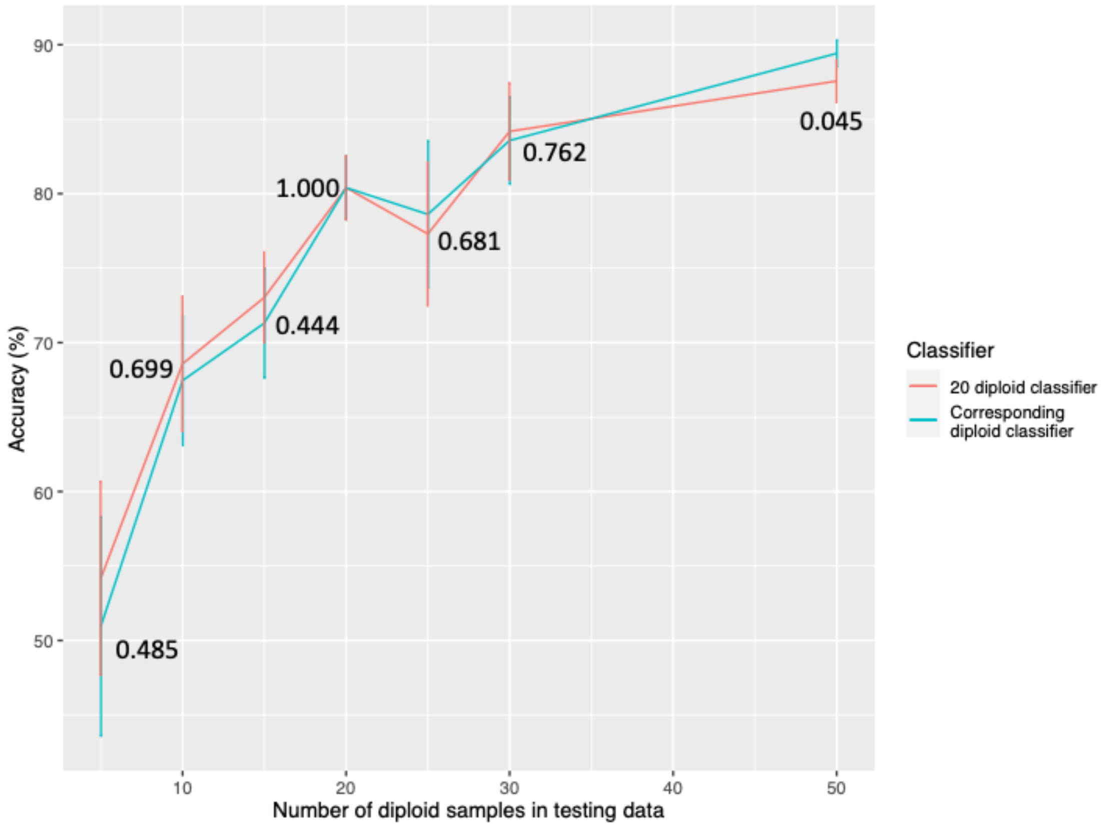


**Figure S22.** Change in accuracy as the number of diploids sampled from all sampled populations increases using a classifier trained on 20 diploids vs the corresponding classifier trained on the same number of diploids as the testing data. Error bars show the standard deviation from the mean over five testing tree sequences. Numbers show the two sample T-test (two-sided) p-values comparing the two classifiers.

Classifiers were able to generalise to fewer samples than in the training data. When the number of samples is much larger than simulated in the training data, the accuracy begins to decrease. This decrease is only to a small extent even for more than twice the number of samples demonstrating that classifiers are flexible to differences in sample size.

##### 3.3.6 Inference of ghost lineages

Genetic evidence has revealed ghost populations in many species, including humans. This is when a population is inferred to have existed but is not sampled with DNA or in the fossil record. To test whether our method could be used in a case involving a ghost population, we applied a classifier trained on a simulation where one ’path population’ of the four paths present was not sampled.

We tested two scenarios from Figure S17: one where the samples from population 0 were removed meaning path 1 contained the ghost lineage; and one where samples from population 9 were removed meaning path 4 contained the ghost lineage.

Figure S23 shows that the precision in all paths is not decreased when path 4 contains the ghost lineage and the precision of the classifier to predict path 4 actually increases. It appears that the classifier receives enough information from other populations present along path 4 to continue to identify path 4 with high precision.

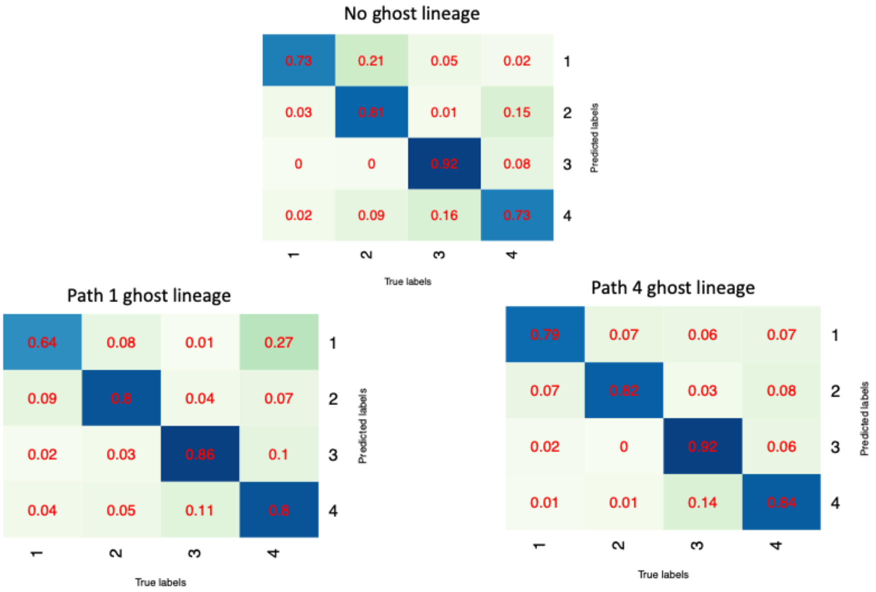


**Figure S23.** Confusion matrices showing the precision of classifiers in each class when various populations were removed from training and testing data to mimic ghost lineages.

When path 1 contains the ghost lineages, the precision of the classifier is decreased in path 1. This is likely because there are no other populations along path 1 that are included in the GNNs, meaning that when population 0 is removed it becomes harder to identify the path.

Overall, the accuracy remains high for identifying ghost lineages. The precision maintained depends on whether there are other sampled populations involved that can inform the classifier of path identity. The ability to identify ghost lineages, even with little to no other populations present along the lineages highlights the advantage of a path structure compared to a framework of single population identity that is used by other local ancestry inference tools.

##### 3.4 A method to estimate time since admixture

We devised an LD-based method to infer the time since admixture of samples from our painted chromosomes. LD-based methods are more robust to noisy ancestry inference and therefore are appropriate to our data. Segments are painted by path, so admixture corresponds to when two paths join in a population history. This means that tracts made up of two or more paths combined can be used to date multiple admixture events, where one event that joins multiple paths follows other events that join two or more paths. For example, the joining of paths 1, 2, 3 and 4 in the Bronze Age admixture event in Figure S7. is preceded by paths 1 and 3 joining in the Neolithic farmers and paths 2 and 4 joining in the Yamnaya.

With empirical sampling we plot the probability of being in the same path as a function of genetic distance and fit an exponential decay curve to the distribution. The parameters of the exponential decay correspond to the values of interest, time since admixture and admixture fraction. This can be done for each sample chromosome individually and the results across the whole genome for each sample combined to give an admixture time estimate and admixture fraction for each individual.

The sampling process is as follows:

1. Sample a starting position uniformly from between 0 and 0.5cM from one end of the chromosome.
2. Record the starting path.
3. Move 1cM towards the end of the chromosome.
4. Record the path 1cM away.
5. Now move to a new starting position 1cM away from the previous starting position.
6. Repeat steps 2-5 until the end of the chromosome.

Repeat the above steps for testing distances 1-50cM in step 3 and 4. Over all autosomes, we can then calculate the probability of being in path y given starting in path x, d cM away, for all path combinations of x and y, including x=y. The shortest distance is 1cM, as in ROLLOFF, to avoid the effect of background LD.

The probability when x=y decays exponentially with genetic distance and the rate of decay depends on the time since admixture. Parameters are determined by nonlinear least squares regression of the data to the formula

$$P(\text{same path}) = \alpha e^{(-\beta d)} + \theta \qquad (1)$$

The probability will asymptote to the probability of being in path x across the whole chromosome, which intuitively is the admixture fraction of path x, therefore the admixture fraction is given by θ. The β parameter represents the time since admixture in generations (Figure S24).

The process is performed over all autosomes for each individual genome and homologous chromosomes are treated as independent. For admixture events between two populations, each individual has two estimates of the admixture age, one calculated per ancestral path. We combined the estimates of the two in a way that minimises the standard error to give a weighted average value of time since admixture for each individual. The time of admixture in generations ago from present-day can therefore be calculated as the sum of the sample age in generations ago and the estimated time since admixture for that sample.

##### 3.4.1 Performance on simulated data

To test the performance of this method, we simulated data from the model of European population structure. We executed the analysis on the admixed populations to infer time since admixture and admixture fractions of paths.

We dated the admixture times of all individuals by extracting the β parameter from fitting exponential decay curves as described above. Counts for all distances were taken across all nine tree sequences. The probability of remaining in the same path was therefore calculated using the total counts from across 18 independent sequences per individual. The method was applied on both the simulated painted tree sequences and on tree sequences that were inferred by RELATE from the simulated data and painted using a classifier. The results are shown in Figure S24 and Tables S7 and S8.

**Table 21.** Table of results of admixture time analysis performed on simulated painted tree sequences. For three admixed populations the mean time of admixture across all simulated samples and the standard deviation of estimates around the mean is shown. All values are in units of generations ago.

| Population | Simulated time of admixture | Mean time of admixture | Standard deviation (+/-) |
|---|---|---|---|
| Bronze Age | 166 | 166.03 | 3.86 |
| Neolithic farmers | 259 | 258.54 | 12.96 |
| Yamnaya | 177 | 177.74 | 1.66 |

In Figure S24, as the sample ages become more recent and further from the admixture time, the variance of estimates around the true value increases for both simulated and RELATE inferred results. Likewise, Figure S25 shows how the standard error of estimates increases as the sample ages get further from the time of admixture. More generations since admixture mean that more recombination events have occurred resulting in shorter ancestry tracts.

Shorter tracts make it harder to differentiate between background LD inherited from the parent populations from the relevant admixture LD. Therefore, the standard error for the β parameters when fitting exponential decay curves is greater and the variance in β parameters estimates is greater. This is the case for both simulated data and RELATE inferred data showing there is a limit to the length of admixture tracts, even in data with no noise, for inferring time since admixture accurately.

Noise appears as short tracts of misclassified correlated trees in RELATE inferred data, which starts to become difficult to differentiate from admixture LD when the admixture tracts become comparable in length. The standard error values for the analysis of tree sequences inferred by RELATE up to 100 generations since admixture are very similar to those from the analysis of the simulated data (Figure S25). This suggests that up to approximately 100 generations since admixture the method is robust to noise introduced by classification error in the painted chromosomes. Similar results can be seen in Figure S24, as the sample ages become more recent and further from the admixture time, the deviation of estimates from the true value increases in both sets of data but to a greater extent in RELATE inferred data. Likewise, the overall standard deviation of the mean time since admixture estimate across all samples is slightly larger in the RELATE inferred data compared to the simulated data for all populations (Tables S7 and S8).

**Table 22.** Table of results of admixture time analysis performed on tree sequences that were inferred by RELATE from the simulated data and painted using a classifier. For three admixed populations, the mean time of admixture estimate across all simulated samples and the standard deviation of estimates around the mean is shown. All values are in units of generations ago.

| Population | Simulated time of admixture | Mean time of admixture | Standard deviation (+/-) |
|---|---|---|---|
| Bronze Age | 166 | 166.45 | 4.56 |
| Neolithic farmers | 259 | 257.28 | 16.67 |
| Yamnaya | 177 | 177.24 | 2.00 |

In results of analysis of the simulated data, the Bronze Age mean admixture time predicted across all samples matches the true value of 166 (Table 21) and a small standard deviation of estimates of +/- 3.86 generations. Neolithic Farmers display a mean admixture time slightly under the true value of 259 and a larger standard deviation of estimates of 12.96 +/- generations. The slightly poorer ability to estimate Neolithic farmer admixture time is because some samples have ages that are further from admixture compared to the Bronze Age population, resulting in the mean being slightly under the true admixture time and a greater standard deviation. The error may also be increased by the ancestry proportions of 75/25 Anatolian and WHG respectively, as the exponential decays are harder to fit when the difference between the starting value, near 1, and the asymptotic value of 0.75 is not as large. Yamnaya samples are a maximum of 17 generations from the true admixture time of 177. The mean predicted admixture date matches that simulated and there is a small standard deviation of estimates of +/- 1.66 generations.

Results of analysis of RELATE inferred data were very similar to that of the simulated data (Table 22). The Neolithic farmer's mean β estimate is slightly further from the truth in the RELATE inferred data analysis than in the simulated data analysis due the same reasons explained above; more samples with a greater time since admixture, above 100 generations, meaning noise introduced by classification has more of an effect of decreasing the accuracy of the β estimates.

For the present-day samples that are all 166 generations since admixture, in both simulated and RELATE inferred tree sequences, the admixture tracts have become too short to produce a reliable estimate of time.

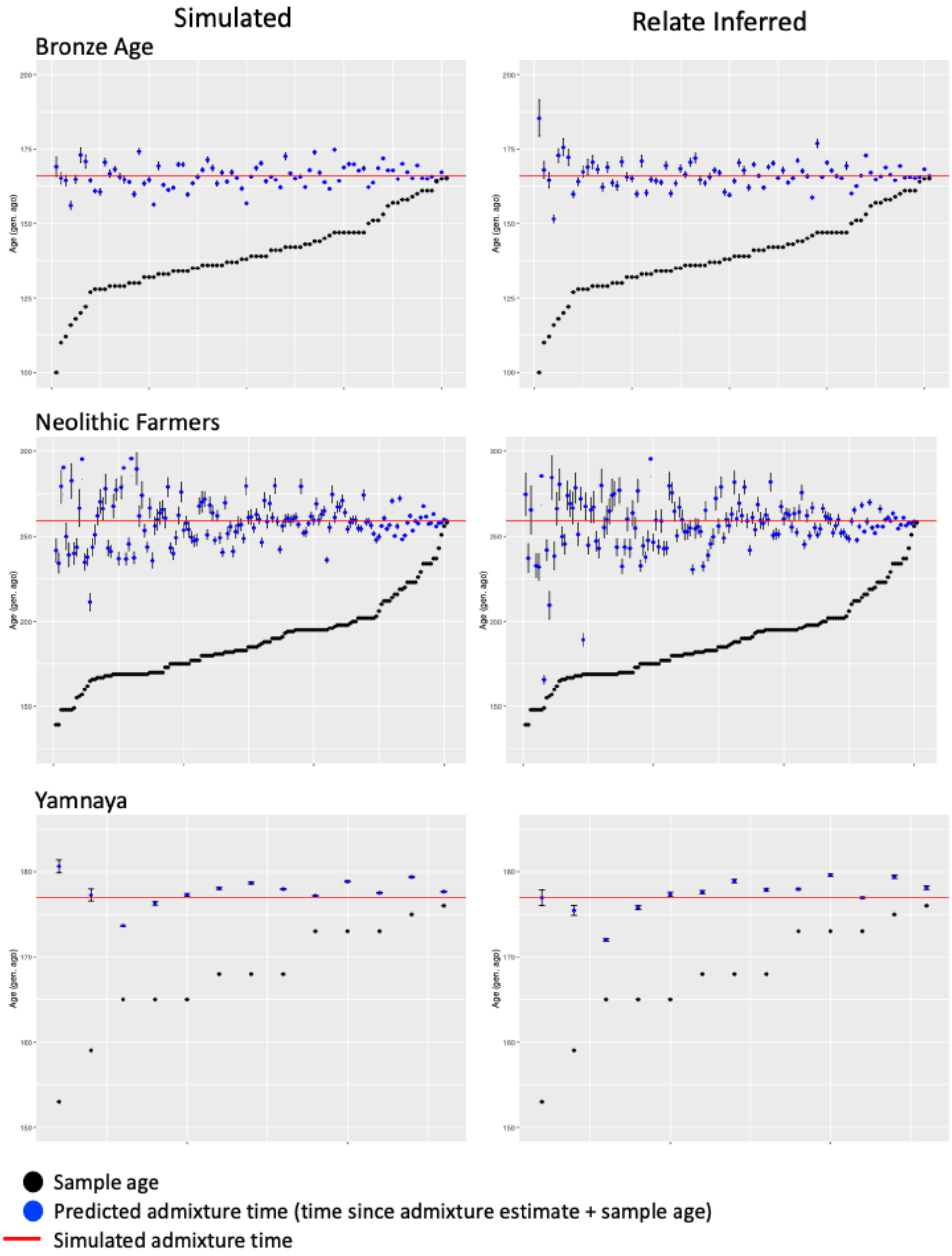


**Figure S24.** Plots showing the predicted mean admixture time with standard error bars and the sample age for three simulated admixed populations. The time since admixture from each painted individual was calculated from the β parameter described in Supplementary Note 3.4 and the time of admixture plotted from the sum of the time since admixture and sample age in generations ago.

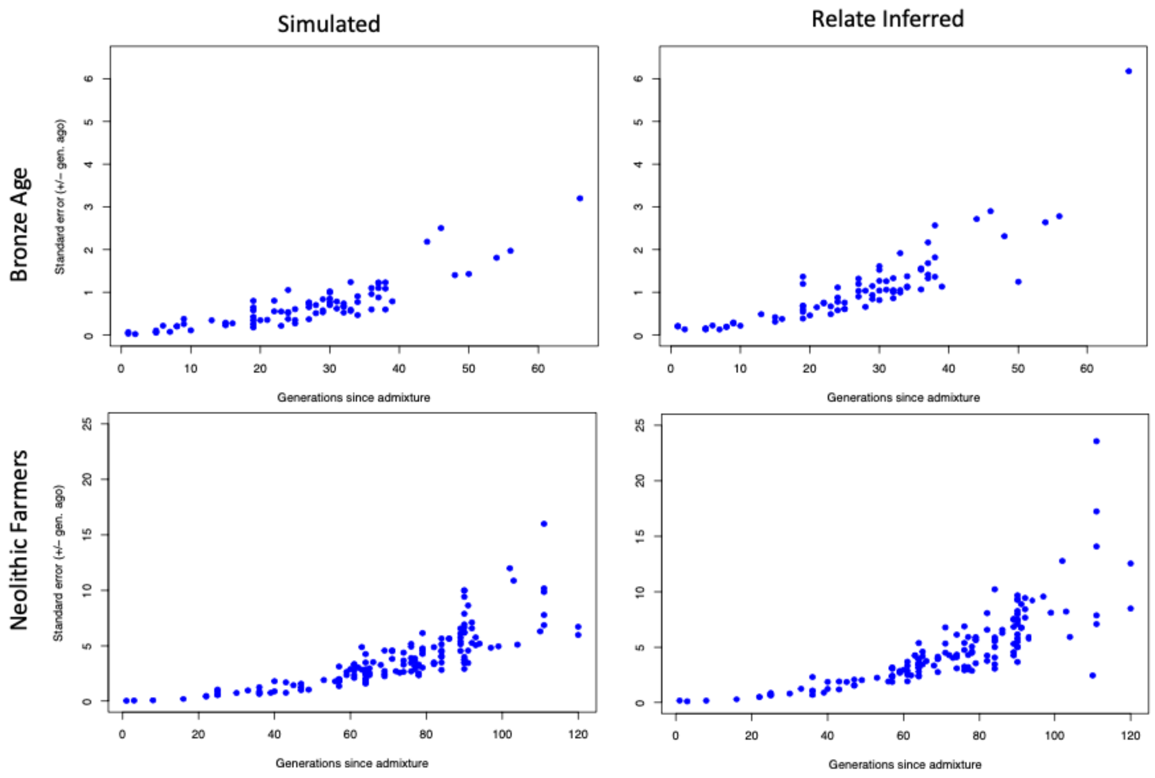


**Figure S25.** Plots showing the decrease in standard error of admixture time estimates as sample age increases and gets closer to the simulated time of admixture for Bronze Age and Neolithic farmer populations. The left column shows the results for admixture time analysis on the simulated tree sequences and the right shows the results for tree sequences inferred by RELATE from the same simulated data and painted using a classifier.

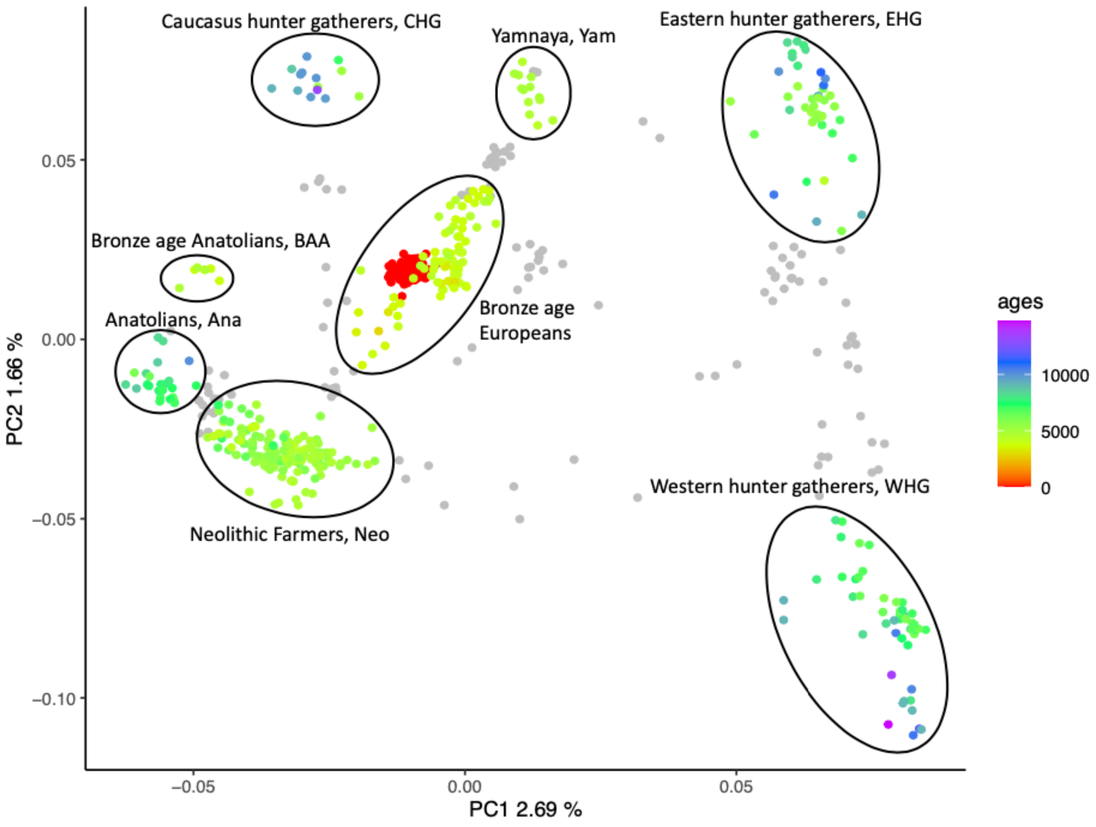


**Figure S26.** Subset of MesoNeo genomes, plotted by their first two principal components. Samples that are diagnostic of each ancient European group are coloured by their radiocarbon age in years BP. The samples that fall in between circled samples and were removed from further analysis, coloured in grey.

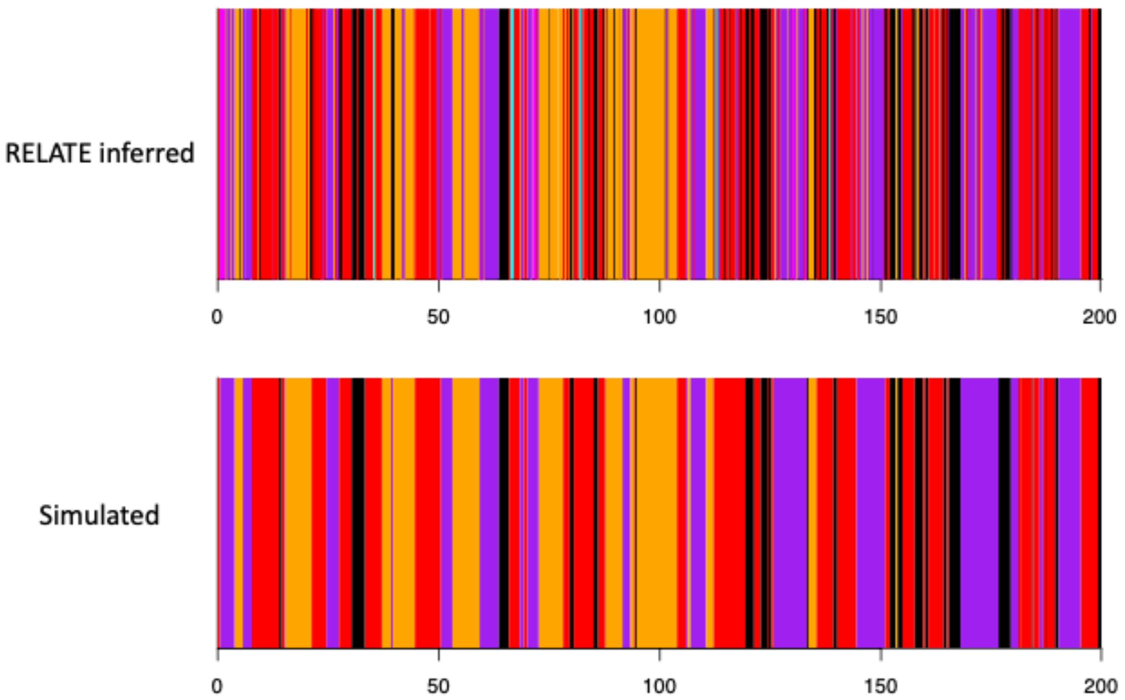


**Figure S27.** Example simulated painted haploid chromosomes. Painted haploid chromosomes from a Bronze Age individual. The top chromosome shows the true simulated painting and the bottom chromosome shows the corresponding RELATE inferred chromosome, painted by classification.

**Table 23.** Summary of longitude and latitude linear regression models for 173 Neolithic farmers samples.

| Coefficients | Estimate | Std. Error | P-value |
|---|---|---|---|
| Inferred admixture time ~ longitude + latitude | | | |
| Intercept | 9092.66 | 450.59 | < 2e-16 |
| Longitude | -25.66 | 4.87 | 4.08e-07 |
| Latitude | -42.74 | 8.98 | 4.18e-06 |
| R2-adjusted = 0.234 | | Model p-value = 5.947e-11 | |
| Sample age ~ longitude + latitude | | | |
| Intercept | 7394.92 | 488.34 | < 2e-16 |
| Longitude | -11.7 | 5.65 | 0.0396 |
| Latitude | -35.25 | 9.89 | 0.000471 |
| R2-adjusted = 0.084 | | Model p-value = 0.0013 | |

**Table S10.** Summary of longitude and latitude linear regression models for 97 Bronze Age samples.

| Coefficients | Estimate | Std. Error | P-value |
|---|---|---|---|
| Inferred admixture time ~ longitude + latitude | | | |
| Intercept | 3894.05 | 443.23 | 7.57e-14 |
| Longitude | 2.98 | 5.77 | 0.607 |
| Latitude | 16.15 | 9.08 | 0.0785 |

#### References

1. Slatkin M, Racimo F. Ancient DNA and human history. *Proceedings of the National Academy of Sciences*. 2016 2022/08/29;113(23):6380–6387. Available from: https://doi.org/10.1073/pnas.1524306113

2. Lazaridis I, Patterson N, Mittnik A, Renaud G, Mallick S, Kirsanow K, et al. Ancient human genomes suggest three ancestral populations for present-day Europeans. *Nature*. 2014 Sep;513(7518):409–413

3. Marnetto D, Pärna K, Läll K, Molinaro L, Montinaro F, Haller T, et al. Ancestry deconvolution and partial polygenic score can improve susceptibility predictions in recently admixed individuals. *Nature Communications*. 2020;11(1):1628. Available from: https://doi.org/10.1038/s41467-020-15464-w

4. Atkinson EG, Maihofer AX, Kanai M, Martin AR, Karczewski KJ, Santoro ML, et al. Tractor uses local ancestry to enable the inclusion of admixed individuals in GWAS and to boost power. *Nature Genetics*. 2021;53(2):195–204. Available from: https://doi.org/10.1038/s41588-020-00766-y

5. Falush D, Stephens M, Pritchard JK. Inference of population structure using multilocus genotype data: linked loci and correlated allele frequencies. *Genetics*. 2003 Aug;164(4):1567–1587

6. Patterson N, Hattangadi N, Lane B, Lohmueller KE, Hafler DA, Oksenberg JR, et al. Methods for high-density admixture mapping of disease genes. *Am J Hum Genet.* 2004 May;74(5):979–1000

7. Sankararaman S, Sridhar S, Kimmel G, Halperin E. Estimating local ancestry in admixed populations. *Am J Hum Genet*. 2008 Feb;82(2):290–303

8. Pasaniuc B, Sankararaman S, Kimmel G, Halperin E. Inference of locus-specific ancestry in closely related populations. *Bioinformatics*. 2009 Jun;25(12):i213–21

9. Tang H, Coram M, Wang P, Zhu X, Risch N. Reconstructing genetic ancestry blocks in admixed individuals. *Am J Hum Genet.* 2006 Jul;79(1):1–12

10. Price AL, Tandon A, Patterson N, Barnes KC, Rafaels N, Ruczinski I, et al. Sensitive Detection of Chromosomal Segments of Distinct Ancestry in Admixed Populations. *PLOS Genetics.* 2009 06;5(6). Available from: https://doi.org/10.1371/journal.pgen.1000519

11. Baran Y, Pasaniuc B, Sankararaman S, Torgerson DG, Gignoux C, Eng C, et al. Fast and accurate inference of local ancestry in Latino populations. *Bioinformatics*. 2012 May;28(10):1359–1367

12. Maples BK, Gravel S, Kenny EE, Bustamante CD. RFMix: a discriminative modeling approach for rapid and robust local-ancestry inference. *Am J Hum Genet.* 2013 Aug;93(2):278–288

13. Hilmarsson H, Kumar AS, Rastogi R, Bustamante CD, Montserrat DM, Ioannidis AG. High Resolution Ancestry Deconvolution for Next Generation Genomic Data. *bioRxiv*. 2021 01; Available from: http://biorxiv.org/content/early/2021/09/21/2021.09.19.460980.abstract

14. Speidel L, Forest M, Shi S, Myers SR. A method for genome-wide genealogy estimation for thousands of samples. *Nature Genetics.* 2019;51(9):1321–1329. Available from: https://doi.org/10.1038/s41588-019-0484-x

15. Speidel L, Cassidy L, Davies RW, Hellenthal G, Skoglund P, Myers SR. Inferring Population Histories for Ancient Genomes Using Genome-Wide Genealogies. *Molecular Biology and Evolution*. 2021 06;38(9):3497–3511. Available from: https://doi.org/10.1093/molbev/msab174

16. Allentoft ME, Sikora M, Sjögren KG, Rasmussen S, Rasmussen M, Stenderup J, et al. Population genomics of Bronze Age Eurasia. *Nature*. 2015;522(7555):167–172. Available from: https://doi.org/10.1038/nature14507

17. Jones ER, Gonzalez-Fortes G, Connell S, Siska V, Eriksson A, Martiniano R, et al. Upper Palaeolithic genomes reveal deep roots of modern Eurasians. *Nature Communications*. 2015;6(1):8912. Available from: https://doi.org/10.1038/ncomms9912

18. Kelleher J, Etheridge AM, McVean G. Efficient Coalescent Simulation and Genealogical Analysis for Large Sample Sizes. *PLOS Computational Biology*. 2016 05;12(5):e1004842–. Available from: https://doi.org/10.1371/journal.pcbi.1004842

19. Lazaridis I, Nadel D, Rollefson G, Merrett DC, Rohland N, Mallick S, et al. Genomic insights into the origin of farming in the ancient Near East. *Nature*. 2016 Aug;536(7617):419–424

20. Mathieson I, Alpaslan-Roodenberg S, Posth C, Szécsényi-Nagy A, Rohland N, Mallick S, et al. The genomic history of southeastern Europe. *Nature*. 2018;555(7695):197–203. Available from: https://doi.org/10.1038/nature25778

21. Hofmanová Z, Kreutzer S, Hellenthal G, Sell C, Diekmann Y, Díez-Del-Molino D, et al. Early farmers from across Europe directly descended from Neolithic Aegeans. *Proceedings of the National Academy of Sciences.* 2016 Jun;113(25):6886–6891

22. Brace S, Diekmann Y, Booth TJ, van Dorp L, Faltyskova Z, Rohland N, et al. Ancient genomes indicate population replacement in Early Neolithic Britain. *Nature Ecology & Evolution*. 2019;3(5):765–771. Available from: https://doi.org/10.1038/s41559-019-0871-9

23. Haak W, Lazaridis I, Patterson N, Rohland N, Mallick S, Llamas B, et al. Massive migration from the steppe was a source for Indo-European languages in Europe. *Nature*. 2015;522(7555):207–211. Available from: https://doi.org/10.1038/nature14317

24.Olalde I, Brace S, Allentoft ME, Armit I, Kristiansen K, Booth T, et al. The Beaker phenomenon and the genomic transformation of northwest Europe. *Nature*. 2018;555(7695):190–196. Available from: https://doi.org/10.1038/nature25738

25. Cassidy LM, Martiniano R, Murphy EM, Teasdale MD, Mallory J, Hartwell B, et al. Neolithic and Bronze Age migration to Ireland and establishment of the insular Atlantic genome [doi: 10.1073/pnas.1518445113]. *Proceedings of the National Academy of Sciences*. 2016 2022/09/01;113(2):368–373. Available from: https://doi.org/10.1073/pnas.1518445113

26. Lazaridis I, Alpaslan-Roodenberg S, Acar A, Açikkol A, Agelarakis A, Aghikyan L, et al. The genetic history of the Southern Arc: A bridge between West Asia and Europe. *Science*. 2022;377(6609):eabm4247. Available from: https://www.science.org/doi/abs/10.1126/science.abm4247

27. Allentoft ME, Sikora M, Refoyo-Martínez A, Irving-Pease EK, Fischer A, Barrie W, et al. Population Genomics of Stone Age Eurasia. *bioRxiv*. 2022 01;Available from: https://doi.org/10.1101/2022.05.04.490594

28. Price AL, Patterson NJ, Plenge RM, Weinblatt ME, Shadick NA, Reich D. Principal components analysis corrects for stratification in genome-wide association studies. *Nature* Genetics. 2006;38(8):904–909. Available from: https://doi.org/10.1038/ng1847

29. Kelleher J, Wong Y, Wohns AW, Fadil C, Albers PK, McVean G. Inferring whole-genome histories in large population datasets. *Nat Genet.* 2019 Sep;51(9):1330–1338

30. Schroeder H, Margaryan A, Szmyt M, Theulot B, Włodarczak P, Rasmussen S, et al. Unraveling ancestry, kinship, and violence in a Late Neolithic mass grave [doi: 10.1073/pnas.1820210116]. *Proceedings of the National Academy of Sciences*. 2019 2022/09/16;116(22):10705–10710. Available from: https://doi.org/10.1073/pnas.1820210116

31. Racimo F, Woodbridge J, Fyfe RM, Sikora M, Sjögren KG, Kristiansen K, et al. The spatiotemporal spread of human migrations during the European Holocene [doi: 10.1073/pnas.1920051117]. *Proceedings of the National Academy of Sciences*. 2020 2022/09/09;117(16):8989–9000. Available from: https://doi.org/10.1073/pnas.1920051117

32. Morgunova NL, Khokhlova OS. Chronology and Periodization of the Pit-Grave Culture in the Region Between the Volga and Ural Rivers Based on Radiocarbon Dating and Paleopedological Research. *Radiocarbon*. 2013;55(3):1286–1296. Available from: https://doi.org/10.1017/S0033822200048190

33. Ali AT, Liebert A, Lau W, Maniatis N, Swallow DM. The hazards of genotype imputation in chromosomal regions under selection: A case study using the Lactase gene region [https://doi.org/10.1111/ahg.12444]. *Annals of Human Genetics*. 2022 2022/10/16;86(1):24–33. Available from: https://doi.org/10.1111/ahg.12444

34. Chintalapati M, Patterson N, Moorjani P. The spatiotemporal patterns of major human admixture events during the European Holocene. *eLife*. 2022 May;11

### 4) Estimating allele frequency trajectories of trait-associated variants

Evan K. Irving-Pease[1], Aaron J. Stern[2], Rasmus Nielsen[1,2], Fernando Racimo[1]

[1] Lundbeck Foundation GeoGenetics Centre, GLOBE Institute, University of Copenhagen, Copenhagen, Denmark

[2] Department of Integrative Biology, University of California, Berkeley

#### Introduction

Genome-wide association studies (GWAS) of present-day human populations have identified large numbers of genetic variants associated with complex traits. However, the extent to which these variants have been under positive selection during recent human evolution is unclear. We aimed to model the allele frequency trajectories and selection coefficients of GWAS variants through time, using genomic data from both present-day populations and ancient individuals sampled across West Eurasia during the Holocene. We used the software *CLUES* [1], which supports inference of allele frequency trajectories from marginal trees sampled from a reconstruction of the ancestral recombination graph (ARG) [2] for a set of genomic sequences, in combination with genotype likelihoods from serially sampled ancient DNA (aDNA).

To account for population structure in our samples, we applied a novel chromosome painting technique (Supplementary Note 3). This technique is based on inference of a sample's nearest neighbours in the marginal trees of an ARG that contains labelled individuals. In our case, the labelling corresponds to ancestral populations that predate the main episodes of admixture in West Eurasia (Supplementary Note 3). This method allows us to accurately assign ancestral population labels to haplotypes found in both ancient and present-day individuals. By conditioning our selection analyses on these haplotype backgrounds, we can infer the selection trajectories of GWAS risk alleles in a manner that is approximately invariant to change in the admixture proportions through time. These ancestry specific allele trajectories reveal many novel aspects about the dynamic interplay between selection and admixture in West Eurasia throughout the Holocene.

#### Methods

The computational pipeline to perform all analyses was written in the `snakemake` workflow management system [3]. For a full list of all the software and versions used, see Table 25. The directed acyclic graph (DAG) of the computational pipeline is shown in Figure S28. All pipeline code, custom scripts and a `conda` environment to replicate the analyses are available in the GitHub repository (https://github.com/ekirving/mesoneo_paper).

##### SNP Ascertainment

###### GWAS SNPs

We ascertained -a list of GWAS targets by downloading version v1.0.2 (r2020-06-04) of the NHGRI-EBI GWAS Catalog [4]; containing 187,403 GWAS associations for 3,735 traits. To account for the varying significance thresholds used in the 4,007 published studies included in the catalogue, we restricted our analysis to SNPs with a genome-wide significance threshold of $p<5e-8$. We further filtered the catalogue to retain only single-nucleotide polymorphisms (SNPs) with a valid dbSNP Reference SNP identifier (rsID); resulting in 121,795 GWAS associations for 70,224 rsIDs. For each of the retained associations, we retrieved the trait ontology hierarchy by querying the EMBL-EBI Ontology Lookup Service [5]. We then queried the Ensembl REST API [6], to retrieve metadata about each rsID; including chromosome and position in the GRCh37 assembly, ancestral allele, and nearest genes.

###### Control SNPs

To determine the extent to which GWAS variants are enriched for selection, we paired each GWAS SNP with a unique “Control SNP”. Control SNPs were ascertained by selecting all biallelic SNPs within the imputed dataset [7] and excluding any that fell within +/- 50 kb of a GWAS SNP or a gene region. Gene annotations for GRCh37 were downloaded from Ensembl (release 87) [8]. Control SNPs were grouped into bins based on their derived allele frequency (DAF), rounded to the nearest 1%, and paired randomly (without replacement) with GWAS SNPs in the same chromosome and DAF bin.

###### Simulated Neutral SNPs

To measure the effects of demography on the modelled allele frequency trajectories, we frequency paired GWAS SNPs with neutral SNPs, simulated under the demography used to train the chromosome painting model (Supplementary Note 3). Neutral simulations were performed with *msprime* [9], for genomes of length 198 Mbp, with sample sizes and ages based on those in the empirical aDNA dataset. We ascertained SNPs by frequency pairing the DAF in the last generation of our neutral simulation with the GWAS SNPs from chr1 of the imputed dataset.

#### 1000G ARG

We built genome-wide genealogies for all samples in the 1000 Genomes Project (1000G) Phase 3 release [10] using the software *Relate* (1.1.3) [2,11].

##### Data pre-processing

Prior to inference of the ARG, we converted VCF files into HAPS format, removed all non-biallelic SNPs, polarised SNPs against the ancestral allele calls from the Ensembl Compara 71 database (ens-staging2:3306) [12], filtered sites using the 1000 Genomes StrictMask (20140520) and generated SNP annotations using `Relate`.

##### ARG Inference

We jointly inferred genome-wide genealogies for all 1000G Phase 3 samples, using `Relate`, assuming a mutation rate of 1.25e-8 and an Ne of 30,000. From this ARG, we extracted subtrees containing samples belonging to three European (EUR) populations: (i) Finnish in Finland (FIN); (ii) British in England and Scotland (GBR); and (iii) Toscani in Italia (TSI). We used these subtrees to jointly reinfer branch lengths and to infer a population size history for the EUR metapopulation. Lastly, we remapped all SNPs which had been pruned during inference of the ARG onto the branch-length calibrated EUR subtrees.

#### Modifications to CLUES

Here we describe several modifications made to *CLUES* (see the GitHub wiki page for more information https://github.com/standard-aaron/clues/wiki). For validation and benchmarking of the modifications see Supplementary Note 5.

##### ARG sampling using Relate

Instead of sampling ARGs using *ARGweaver* [13], we sample ARGs using *Relate* [2] for scalability reasons. `Relate` differs from *ARGweaver* in that it assumes a continuous-time coalescent process (vs discrete-time for *ARGweaver*); hence we modified the hidden Markov model (HMM) used by *CLUES* to (1) take time steps every generation, *vs* over a smaller number (~10-50) of timesteps; and (2) within time steps, the probability density of coalescence is calculated using the approach of references [14] and [15], *vs* the discrete-time lines-of-descent approach used in references [13] and [1].

##### Ancient DNA samples

We also introduced a new feature that allows the user to specify a time series of ancient genotype likelihoods which are incorporated into the HMM. We incorporate these samples by, for a given timestep *t,* including (i.e., multiplying by) a Binomial(n=2, p=$X_t$) emission probability for each ancient sampled during timestep *t* in the HMM, where $X_t$ is the latent allele frequency during timestep *t.* In this particular application, we supplanted genotype likelihoods with genotype posterior probabilities (which should be identical under a uniform prior); we confirmed through tests that this did not yield any systematic biases.

#### Selection Analysis

##### CLUES with Modern 1000G data

For each modelled SNP, we used `Relate` to draw 100 samples from the MCMC posterior distribution of trees at that locus. Trees were sampled assuming a mutation rate of 1.25e-8 [2,16] and using the population size history from the EUR calibrated subtrees.

We ran *CLUES* to infer allele frequency trajectories and selection coefficients from modern 1000G data, using: (i) the 100 sampled trees from *Relate* (`--times`); (ii) the inferred EUR population size history (`--coal`); (iii) with trajectories polarised the by the derived allele (`--A1`); (iv) a terminal frequency equal to the DAF of each SNP in EUR (`--popFreq`); and (v) constrained selection to a single epoch spanning the last 15,000 years (`--timeBins`).

We ran these models for all GWAS and Control SNPs present in the imputed dataset for which `Relate` was able to confidently map a mutation to the inferred trees.

##### CLUES with aDNA Time Series

We also ran *CLUES* in an alternative mode, excluding the modern ARG data, and replacing them with aDNA time series data, using: (i) the time series of aDNA genotype probabilities (`--ancientSamps`) (ii) the inferred EUR population size history (`--coal`); (iii) a terminal frequency equal to the DAF of each SNP in EUR (`--popFreq`); and (iv) constraining selection to a single epoch spanning the last 15,000 years (`--timeBins`). We converted the calendrical ages of the samples into generations by assuming a generation time of 28 years [17].

We ran these models for all GWAS and Control SNPs present in the imputed dataset, irrespective of their mappability in the `Relate` analyses (n=73,988), as well as for a set of Simulated SNPs that were frequency paired with GWAS SNPs from chr1 (n=2,948).

##### CLUES with aDNA Ancestral Paintings

We also ran four additional models for each GWAS and Control SNP, in which we conditioned the time series of aDNA genotype probabilities on one of the four ancestral pathways leading to present-day Europeans (Supplementary Note 3):

1. ANA (Anatolian Farmers -> Neolithic)
2. CHG (Caucasus Hunter-gatherers -> Yamnaya)
3. WHG (Western Hunter-gatherers -> Neolithic)
4. EHG (Eastern Hunter-gatherers -> Yamnaya)

All other particulars of these models were identical to the earlier aDNA analyses, except that genotypes were passed to *CLUES* in haploid mode (`--ancientHaps`), even when both haplotypes in an individual shared the same painting.

For the simulated SNPs, we ran these analyses on a dataset in which the pathways were inferred by the chromosome painting model, using inferred tree sequences from running `Relate` on the simulated VCF (i.e., following the same analysis pipeline as the empirical data).

##### Reference and mapping bias filters

To address issues of mapping biases that may cause artifactual changes in allele frequency at individual sites, we also constructed a causal model for distinguishing direct effects of age on allele frequency from indirect effects mediated by read depth, read length, and/or error rates (Supplementary Note 6). We then filtered out SNPs in which more than half of the signal for temporal allele frequency change was driven by non-biological artefacts ($0.5 \geq F_j < 1.0$). Furthermore, we implemented an additional mapping bias test, in which we compared the inferred present-day frequency of all SNPs based on (i) a *CLUES* model of the aDNA time-series data which was conditioned on the present-day frequency of each variant in the three EUR populations (see above); and (ii) a simpler *CLUES* model containing the aDNA time-series data alone. We then filtered out all SNPs in which the addition of the modern data resulted in an absolute present-day frequency difference of > 0.1 between the two models and increased the significance of the test statistic. We also filtered out SNPs in which the observed pattern of genotypes in modern individuals was inconsistent with the marginal trees inferred from the surrounding haplotypes, as determined by `Relate` [2].

##### Genome-wide selection

We aggregated the results of all *CLUES* models and converted the likelihood-ratio scores into p-values using `2*logLR` as the test statistic, assuming a chi-squared distribution with one degree of freedom; which we validate in Supplementary Note 5. To identify independent genome-wide selection peaks we performed hierarchical clustering, using the 'single linkage' method, to group adjacent genome-wide significant SNPs. We cut the resulting tree at a maximum height of 1 Mb, and filtered for a minimum of six genome-wide significant SNPs in each resulting cluster.

We also obtained a list of putatively selected SNPs and their p-values, inferred in an earlier aDNA study [18]. To ascertain which sweep loci were novel to our study, we used the same hierarchical clustering approach to infer sweeps from the published p-values, rather than rely on the published sweep regions (although our inferred loci were almost identical in range). Additionally, we used *CLUES* to infer new p-values for all genome-wide significant SNPs ($p<5e-8$; $n=381$) from the earlier study [18], as not all SNPs found to be significant in that study were present in our GWAS/Control ascertainment.

**Table 25.** Software and versions used in the allele trajectory pipeline

| Software | Version | URL | Reference |
|---|---|---|---|
| bcftools | 1.10.2 | https://github.com/samtools/bcftools | [19] |
| bedtools | 2.29.2 | https://github.com/arq5x/bedtools2 | [20] |
| biopython | 1.76 | https://github.com/biopython/biopython | [21] |
| clues | 36cb7de | https://github.com/35ajstern/clues | [1] |
| conda | 4.9.0 | https://github.com/conda/conda | [22] |
| msprime | | https://github.com/tskit-dev/msprime | [9] |
| numpy | 1.17.0 | https://github.com/numpy/numpy | [23] |
| pandas | 1.0.4 | https://github.com/pandas-dev/pandas | [24] |
| pysam | 0.15.3 | https://github.com/pysam-developers/pysam | [25] |
| python | 3.6.7 | https://www.python.org | [26] |
| r-base | 3.6.1 | https://www.r-project.org/ | [27] |
| r-bedr | 1.0.7 | https://github.com/cran/bedr | [28] |
| r-dplyr | 0.8.0.1 | https://github.com/tidyverse/dplyr | [29] |
| r-ggplot2 | 3.1.1 | https://github.com/tidyverse/ggplot2 | [30] |
| r-ggrastr | 0.2.1 | https://github.com/VPetukhov/ggrastr | [31] |
| r-ggrepel | 0.8.2 | https://github.com/slowkow/ggrepel | [32] |
| r-ggridges | 0.5.1 | https://github.com/wilkelab/ggridges | [33] |
| r-stringr | 1.4.0 | https://github.com/tidyverse/stringr | [34] |
| relate | 1.1.3 | https://myersgroup.github.io/relate | [2] |
| scipy | 1.4.1 | https://github.com/scipy/scipy | [35] |
| snakemake | 5.12.3 | https://github.com/snakemake/snakemake | [3] |

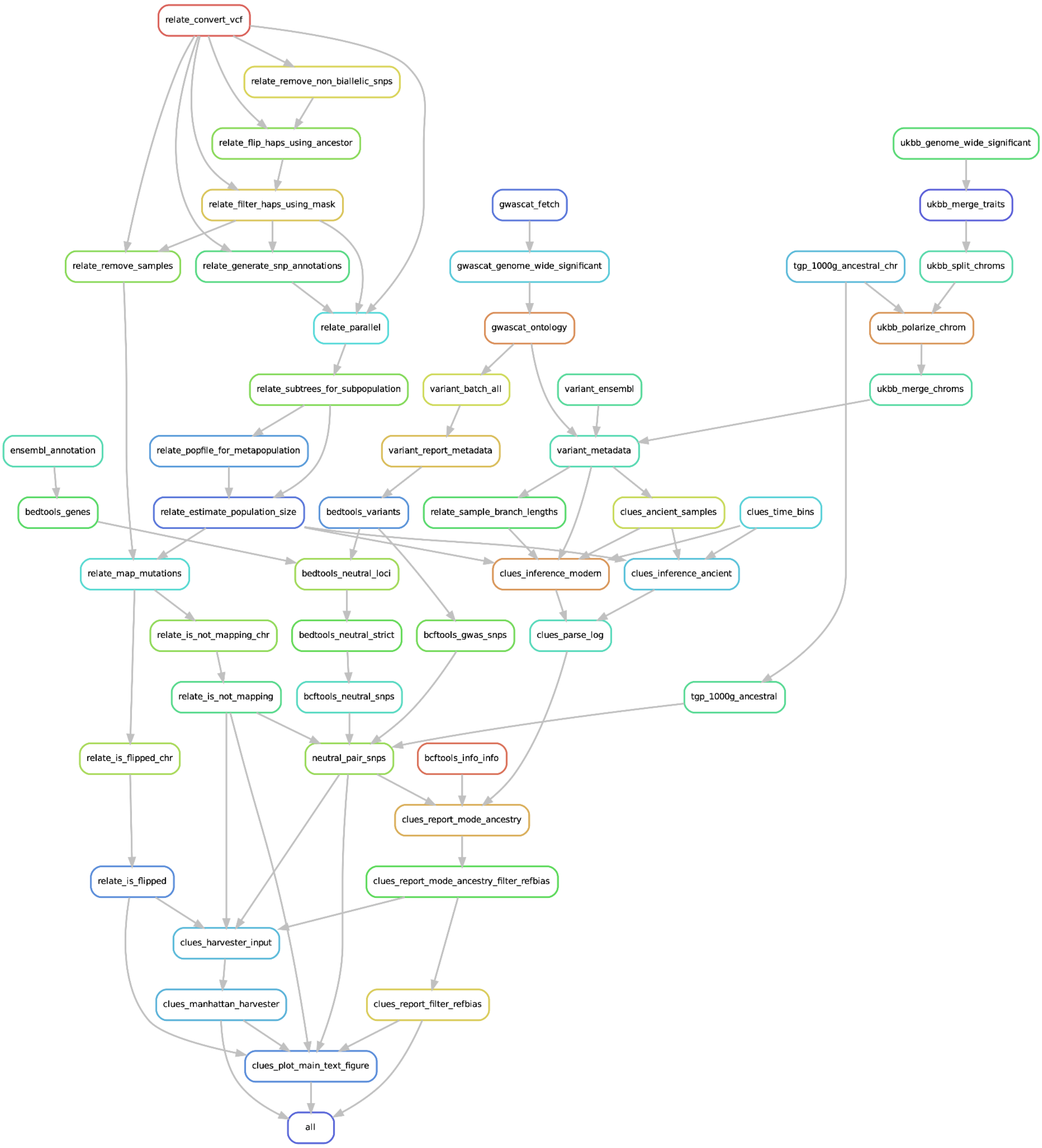


**Figure S28.** Directed acyclic graph (DAG) of the allele frequency trajectory analysis pipeline

#### Results

##### Selection in 1000G EUR

CLUES analysis of all GWAS (n=32,079) and Control group SNPs (n=32,079), which passed all quality control filters, in the 1000G Project populations FIN, GBR, and TSI, identified 9 genome-wide significant SNPs (p<5e-8); 9 in the GWAS group (100%) and none in the Control group (0%) (Supplementary Table 4). Within the GWAS group, we identified no genome-wide significant sweep regions, and none in the Control group (Figure S29). Despite the general lack of genome-wide significant SNPs, the GWAS group was significantly enriched for evidence of selection when compared to the Control group (Wilcoxon signed-rank test, p-value < 7.29e-35).

The most significant SNP was rs12465802 (*R3HDM1*; p=1.09e-10; s=0.011), an intron variant associated with gut microbiota abundance (genus Bifidobacterium id.436) [36], lactase-phlorizin hydrolase levels (LCT.9017.58.3) [37], mosquito bite size [38], and urinary metabolite levels in chronic kidney disease [39] in the GWAS Catalog (r2023-04-07). Other genome-wide significant SNPs within the surrounding region include the lactase persistence SNP rs4988235 (p=3.62e-10; s=0.010), which has been widely reported as a target of strong selection in West Eurasians [40,41].

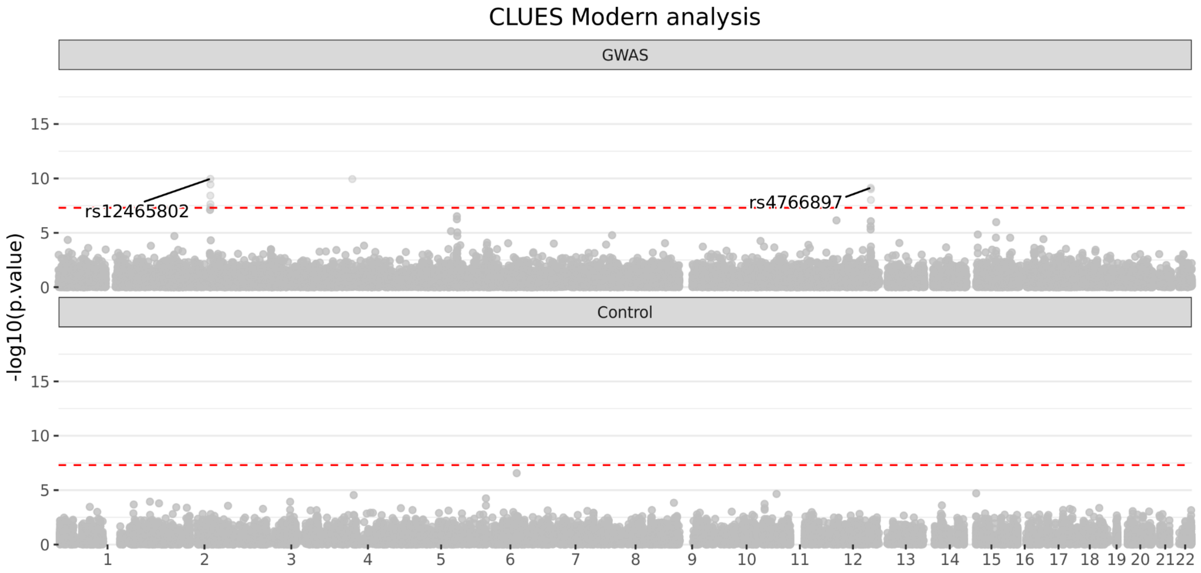

**Figure S29.** Manhattan plot of the p-values from running CLUES on an ARG containing all samples in FIN, GBR, and TSI from 1000G Phase 3, for (a) GWAS SNPs from the GWAS Catalog; and (b) Control SNPs, frequency paired with the GWAS SNPs.

#### Selection in aDNA Time Series

CLUES analysis of all GWAS (n=33,341) and Control group SNPs (n=33,341), which passed all quality control filters in the aDNA time-series dataset, identified 527 genome-wide significant SNPs (p<5e-8); 476 in the GWAS group (90.32%) and 51 in the Control group (9.68%) (Supplementary Table 6). Within the GWAS group, we identified 11 genome-wide significant sweep regions, and none in the Control group (Figure S30).

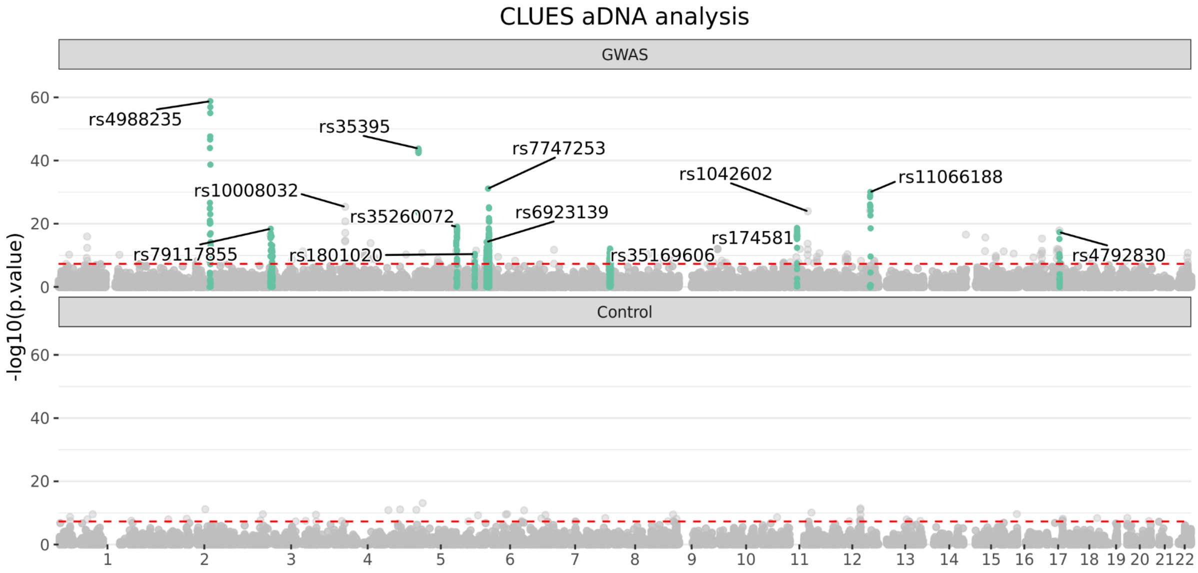


**Figure S30.** Manhattan plot of the p-values from running CLUES on an aDNA time series from all West Eurasian samples in the imputed dataset, for (a) GWAS SNPs from the GWAS Catalog; and (b) Control SNPs, frequency paired with the GWAS SNPs.

#### Peak 1: MCM6

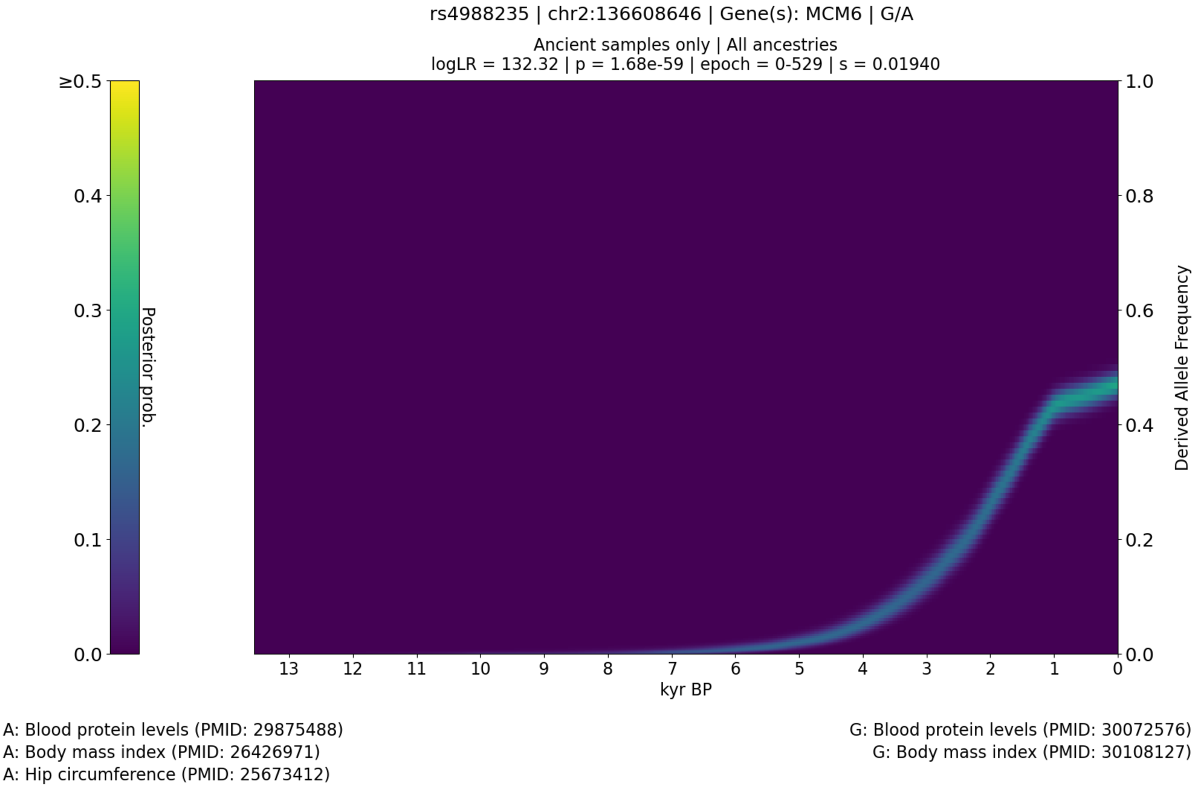


**Figure S31.** CLUES plot of the aDNA time series analysis for all West Eurasian samples in the imputed dataset, showing the posterior probability of the derived allele frequency trajectory for rs4988235 the most significant SNP in the selection peak spanning chr2:134963892-137613935.

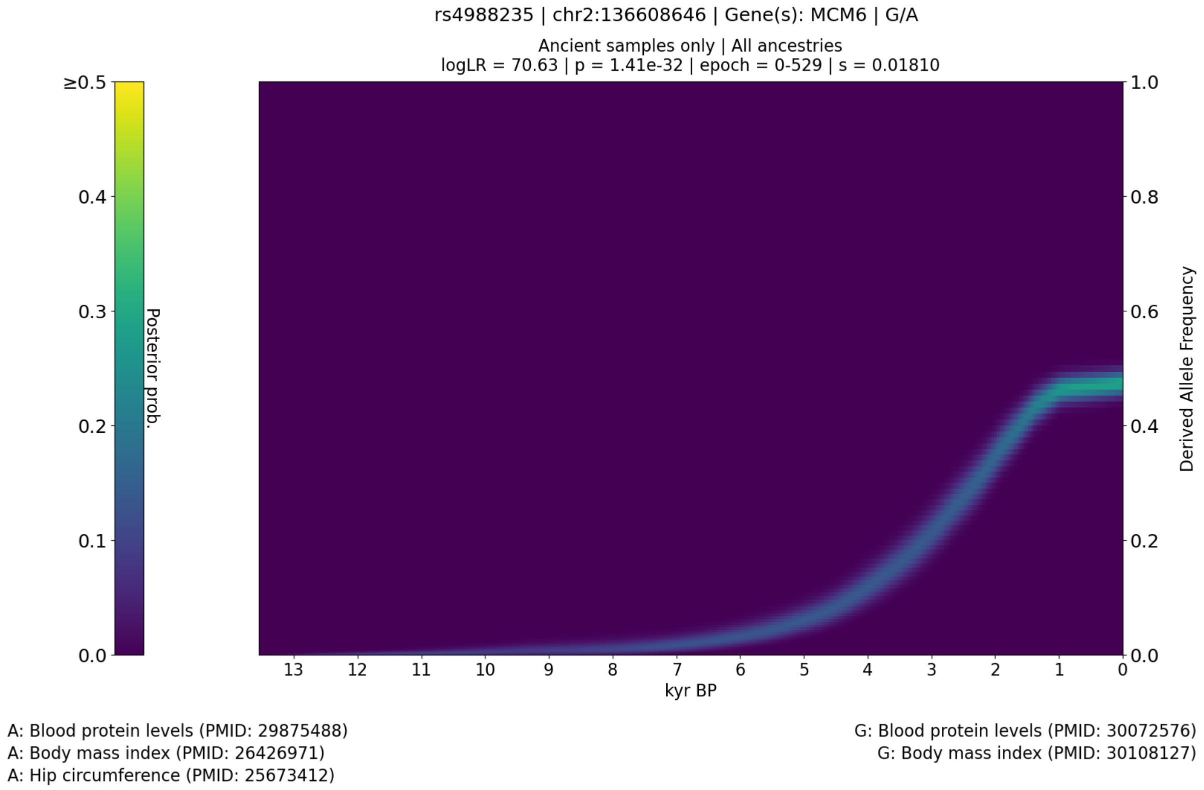


**Figure S32.** CLUES plot of the aDNA time series analysis for all West Eurasian samples in the genotype likelihood dataset (i.e., non-imputed), showing the posterior probability of the derived allele frequency trajectory for rs4988235.

The first peak spanned the region chr2:134963892-137613935, with the most significant SNP being rs4988235 (*MCM6*; p=1.68e-59; s=0.019), an intron variant associated with lactase persistence, 1,5-anhydroglucitol levels [42,43], body mass index [44,45], acetate levels [46], bacilli A abundance in stool [47], bifidobacterium bifidum abundance in stool [47], blood protein levels [48], body fat percentage and LDL-C (pairwise) [49], body mass index (MTAG) [50], body mass index and LDL-C (pairwise) [49], brevibacillaceae abundance in stool [47], brevibacillales abundance in stool [47], broad bean liking [51], degree of unsaturation [46], free cholesterol to total lipids ratio in small HDL [46], gut microbiota abundance (phylum Actinobacteria id.400) [36], hip circumference [52], indolepropionate levels [42], lactase-phlorizin hydrolase levels [53], lactase-phlorizin hydrolase levels (LCT.9017.58.3) [37], lactobacillus B abundance in stool [47], lactobacillus B ruminis abundance in stool [47], medication use for hyperlipidemia (number of purchases) [54], phospholipids to total lipids ratio in medium HDL [46], turicibacter sp001543345 abundance in stool [47], and x-11795 levels [42] in the GWAS Catalog (r2023-04-07).

Peak 2: CCDC12

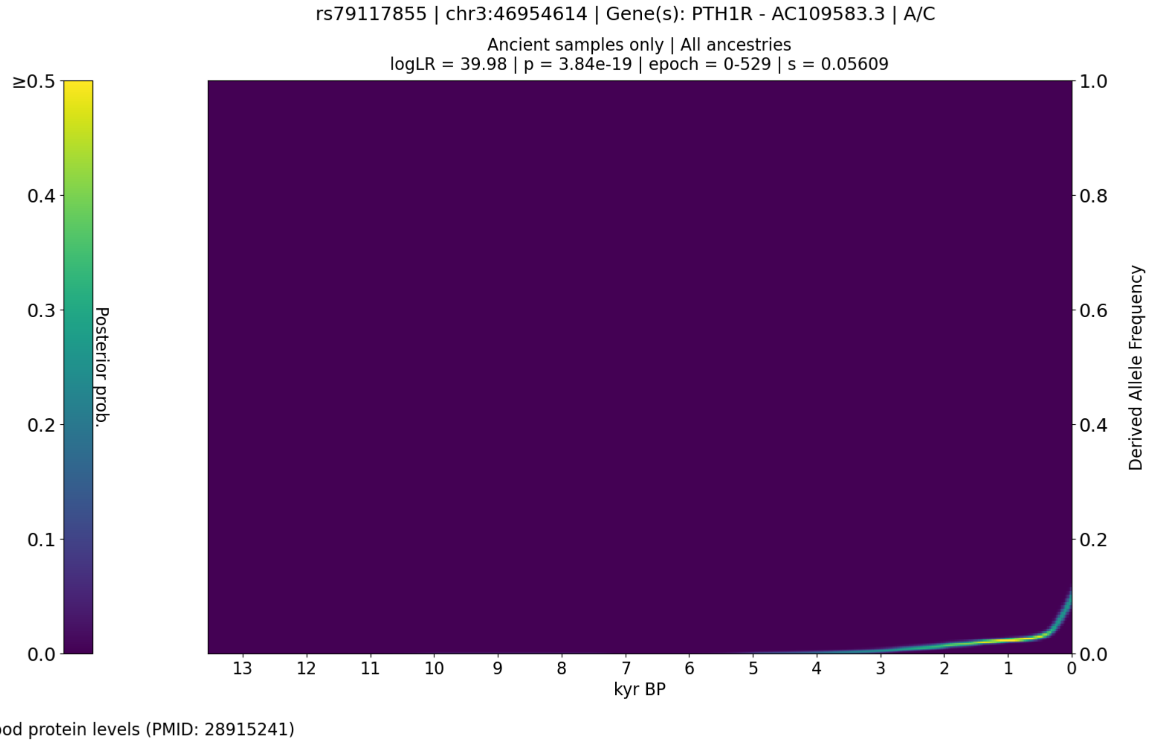


**Figure S33.** CLUES plot of the aDNA time series analysis for all West Eurasian samples in the imputed dataset, showing the posterior probability of the derived allele frequency trajectory for rs79117855 the most significant SNP in the selection peak spanning chr3:45943595-52029991.

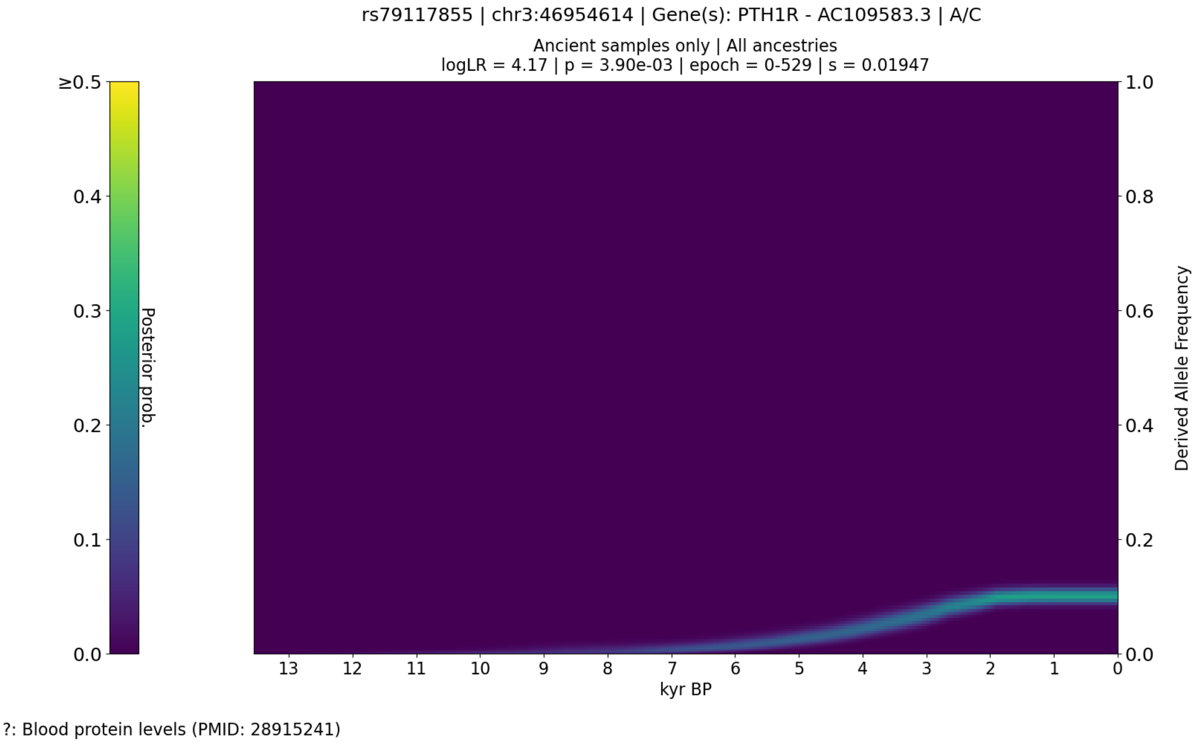


**Figure S34.** CLUES plot of the aDNA time series analysis for all West Eurasian samples in the genotype likelihood dataset (i.e., non-imputed), showing the posterior probability of the derived allele frequency trajectory for rs79117855.

The second peak spanned the region chr3:45943595-52029991, with the most significant SNP being rs79117855 (*CCDC12*; p=3.84e-19; s=0.056), an intergenic variant associated with blood protein levels [55] in the GWAS Catalog (r2023-04-07).

Peak 3: SLC45A2

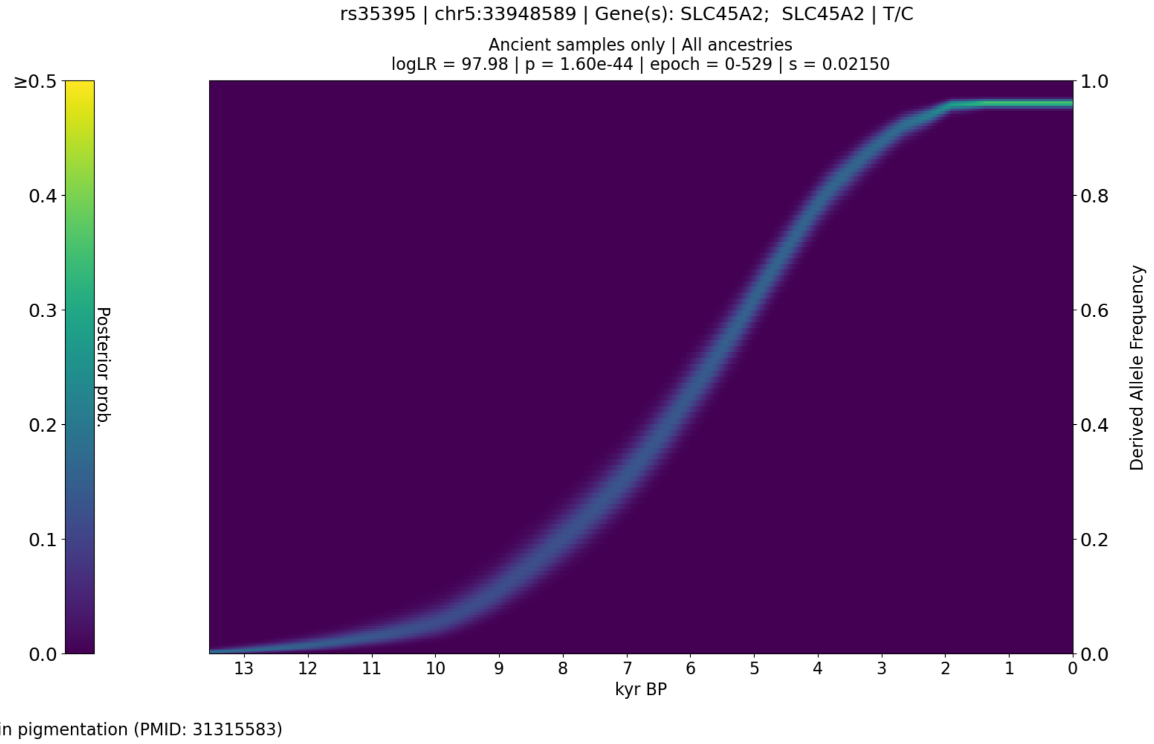


**Figure S35.** CLUES plot of the aDNA time series analysis for all West Eurasian samples in the imputed dataset, showing the posterior probability of the derived allele frequency trajectory for rs35395 the most significant SNP in the selection peak spanning chr5:33946571-33964210.

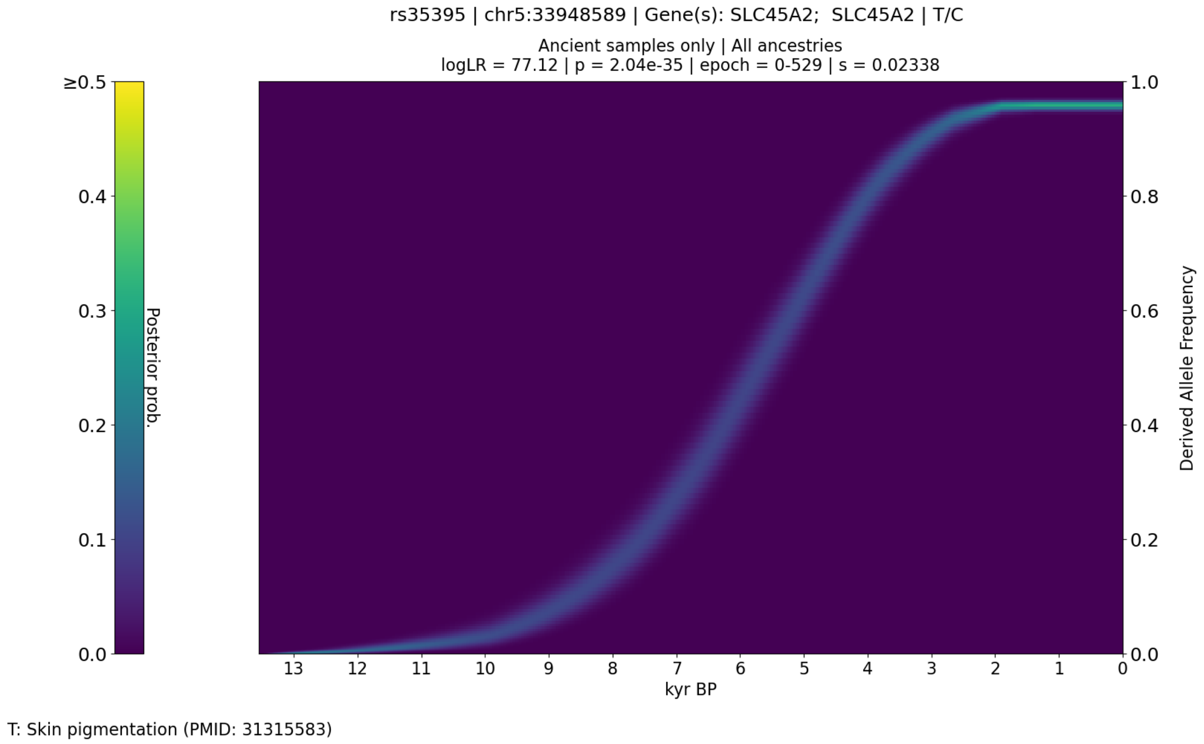


**Figure S36.** CLUES plot of the aDNA time series analysis for all West Eurasian samples in the genotype likelihood dataset (i.e., non-imputed), showing the posterior probability of the derived allele frequency trajectory for rs35395.

The third peak spanned the region chr5:33946571-33964210, with the most significant SNP being rs35395 (*SLC45A2*; p=1.60e-44; s=0.021), an intron variant associated with skin pigmentation [56] in the GWAS Catalog (r2023-04-07).

#### Peak 4: P4HA2 / SLC22A4

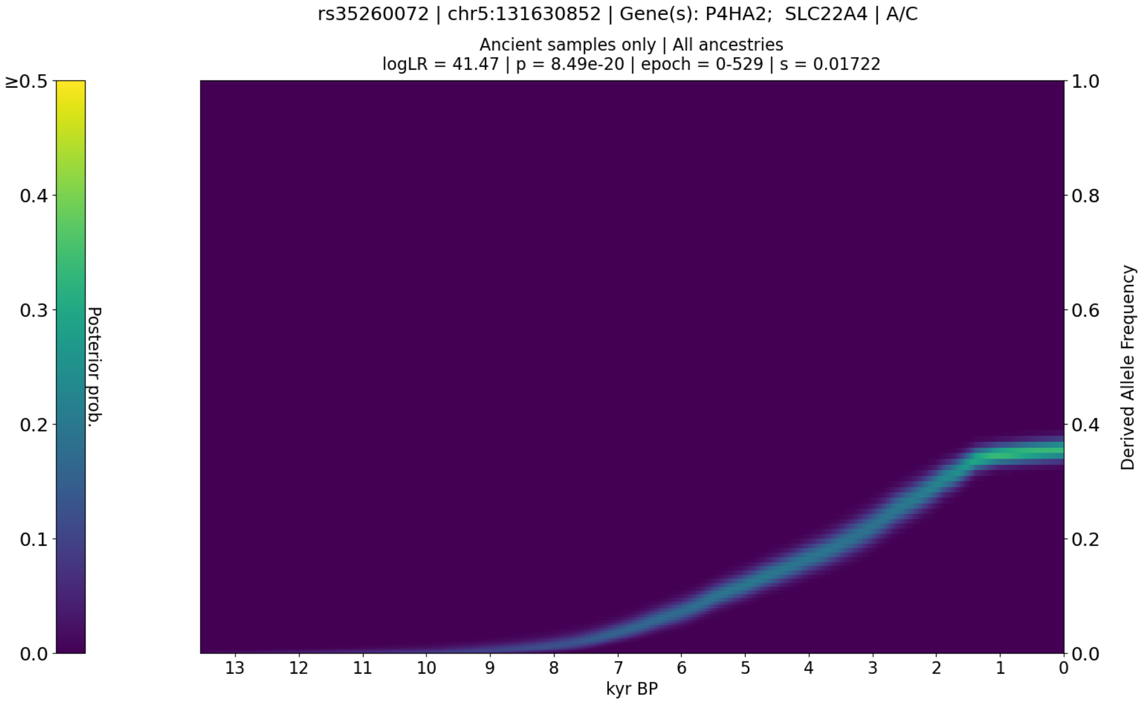


C: Itch intensity from mosquito bite (PMID: 28199695)

**Figure S37.** CLUES plot of the aDNA time series analysis for all West Eurasian samples in the imputed dataset, showing the posterior probability of the derived allele frequency trajectory for rs35260072 the most significant SNP in the selection peak spanning chr5:129573666-131833599.

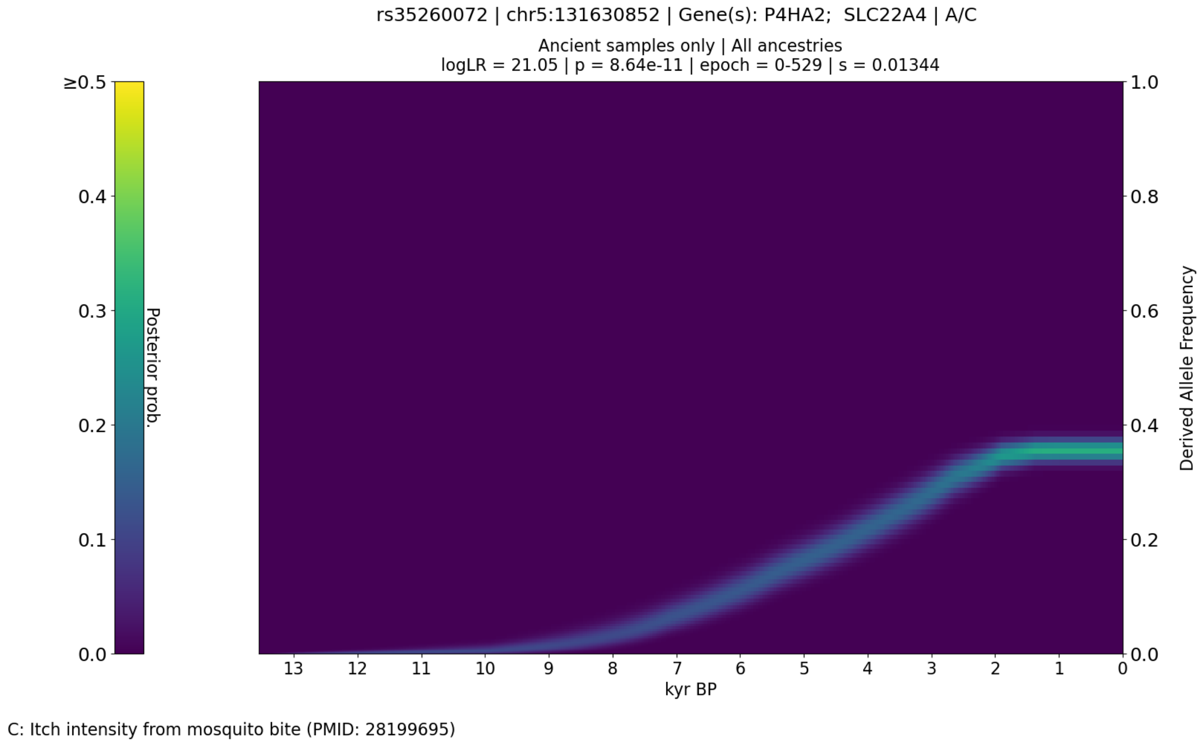


**Figure S38.** CLUES plot of the aDNA time series analysis for all West Eurasian samples in the genotype likelihood dataset (i.e., non-imputed), showing the posterior probability of the derived allele frequency trajectory for rs35260072.

The fourth peak spanned the region chr5:129573666-131833599, with the most significant SNP being rs35260072 (*P4HA2* / *SLC22A4*; p=8.49e-20; s=0.017), an intron variant associated with histidine betaine (hercynine) levels [43] and itch intensity from mosquito bite [38] in the GWAS Catalog (r2023-04-07).

Peak 5: F12 / GRK6

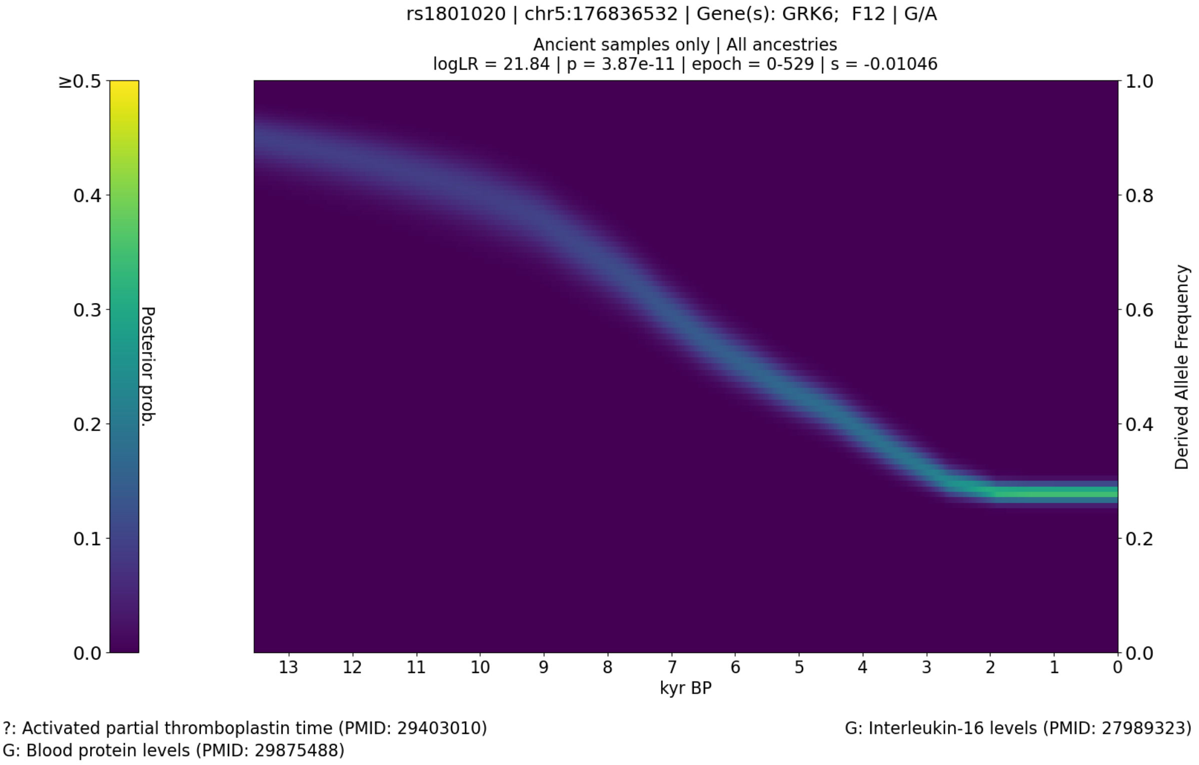


**Figure S39.** CLUES plot of the aDNA time series analysis for all West Eurasian samples in the imputed dataset, showing the posterior probability of the derived allele frequency trajectory for rs1801020 the most significant SNP in the selection peak spanning chr5:176509193-176842474.

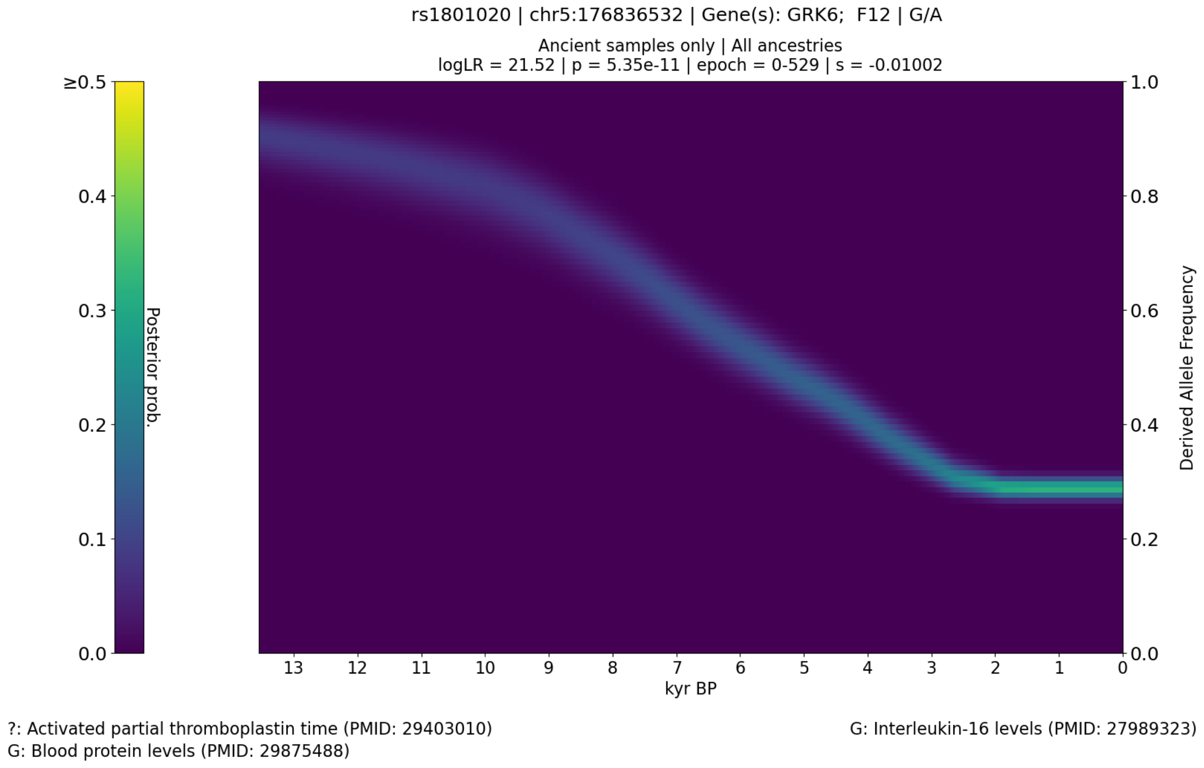


**Figure S40.** CLUES plot of the aDNA time series analysis for all West Eurasian samples in the genotype likelihood dataset (i.e., non-imputed), showing the posterior probability of the derived allele frequency trajectory for rs1801020.

The fifth peak spanned the region chr5:176509193-176842474, with the most significant SNP being rs1801020 (*F12 / GRK6*; p=3.87e-11; s=-0.010), a 5-prime UTR variant associated with activated partial thromboplastin time [57], alpha-2-macroglobulin levels [58], taurine levels [59], aspartate levels [59], blood pressure [60], bMP and activin membrane-bound inhibitor homolog level [61], bone morphogenetic protein 6 levels [62], bone morphogenetic protein 7 levels [58], cadherin-15 levels [37], glucosidase 2 subunit beta levels [37], glycoprotein acetyls levels [46], heme oxygenase 1 levels [62], interleukin-16 levels [63], kininostatin levels [53], lipoprotein lipase levels [62], metabolite peak levels [64], parathyroid hormone-related protein levels [58], plasma serine protease inhibitor levels [58], serine/threonine-protein kinase ULK3 levels [53], serum levels of protein ADM [65], serum levels of protein ATP13A1 [65], serum levels of protein CAPN1;CAPNS1 [65], serum levels of protein CDH15 [65], serum levels of protein DCK [65], serum levels of protein DPEP2 [65], serum levels of protein ETHE1 [65], serum levels of protein GPNMB [65], serum levels of protein ITIH4 [65], serum levels of protein LAMTOR3 [65], serum levels of protein LGMN [65], serum levels of protein PDCL2 [65], serum levels of protein PLXDC1 [65], serum levels of protein SIRT5 [65], serum levels of protein TMEM87B [65], serum lipopolysaccharide activity [66], serum metabolite levels [67], superoxide dismutase mitochondrial levels [37], superoxide dismutase mitochondrial levels [53,68], and thrombin levels [53,58] in the GWAS Catalog (r2023-04-07).

#### Peak 6: HIST1H3PS1

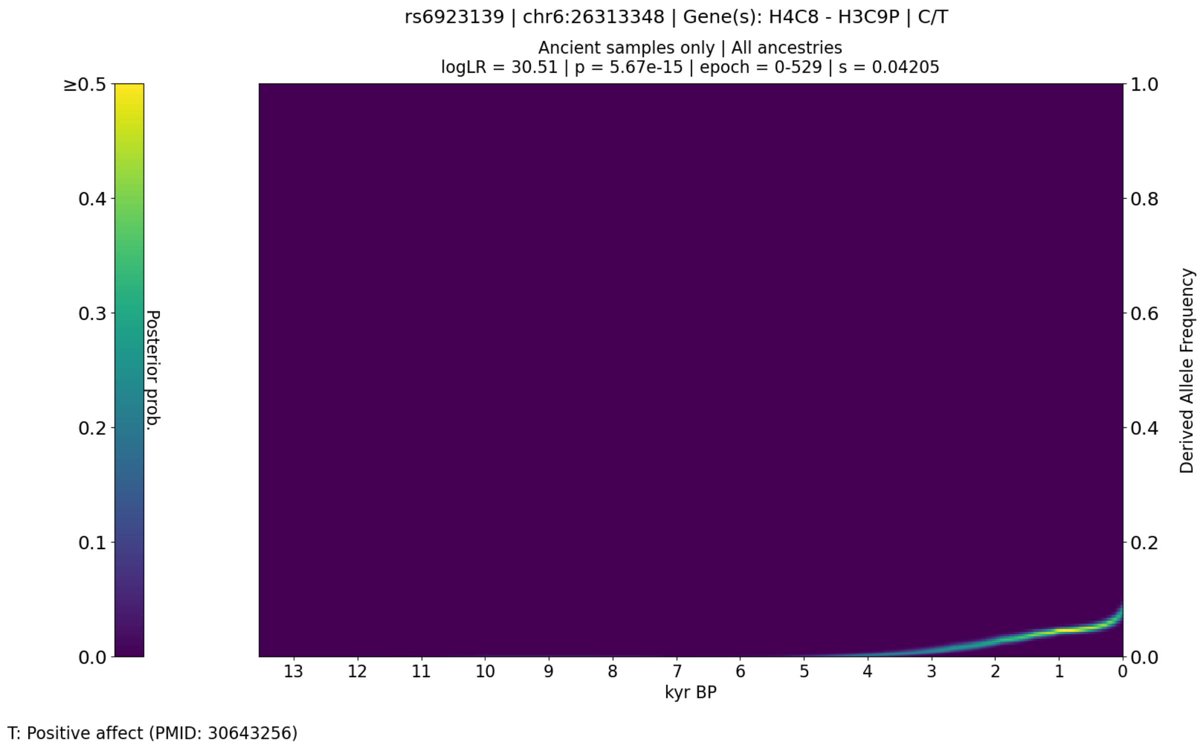


**Figure S41.** CLUES plot of the aDNA time series analysis for all West Eurasian samples in the imputed dataset, showing the posterior probability of the derived allele frequency trajectory for rs6923139 the most significant SNP in the selection peak spanning chr6:25417423-28546952.

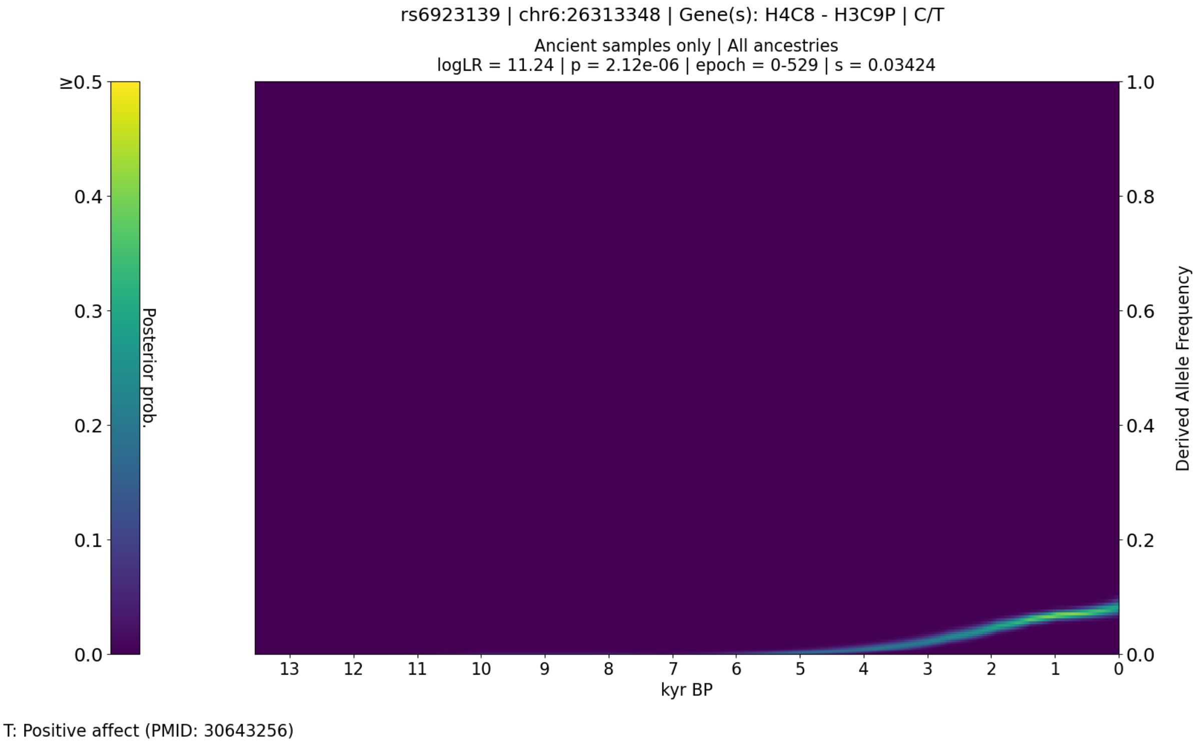


**Figure S42.** CLUES plot of the aDNA time series analysis for all West Eurasian samples in the genotype likelihood dataset (i.e., non-imputed), showing the posterior probability of the derived allele frequency trajectory for rs6923139.

The sixth peak spanned the region chr6:25417423-28546952, with the most significant SNP being rs6923139 (*HIST1H3PS1*; p=5.67e-15; s=0.042), an intergenic variant associated with positive affect [69] in the GWAS Catalog (r2023-04-07).

#### Peak 7: HLA-W

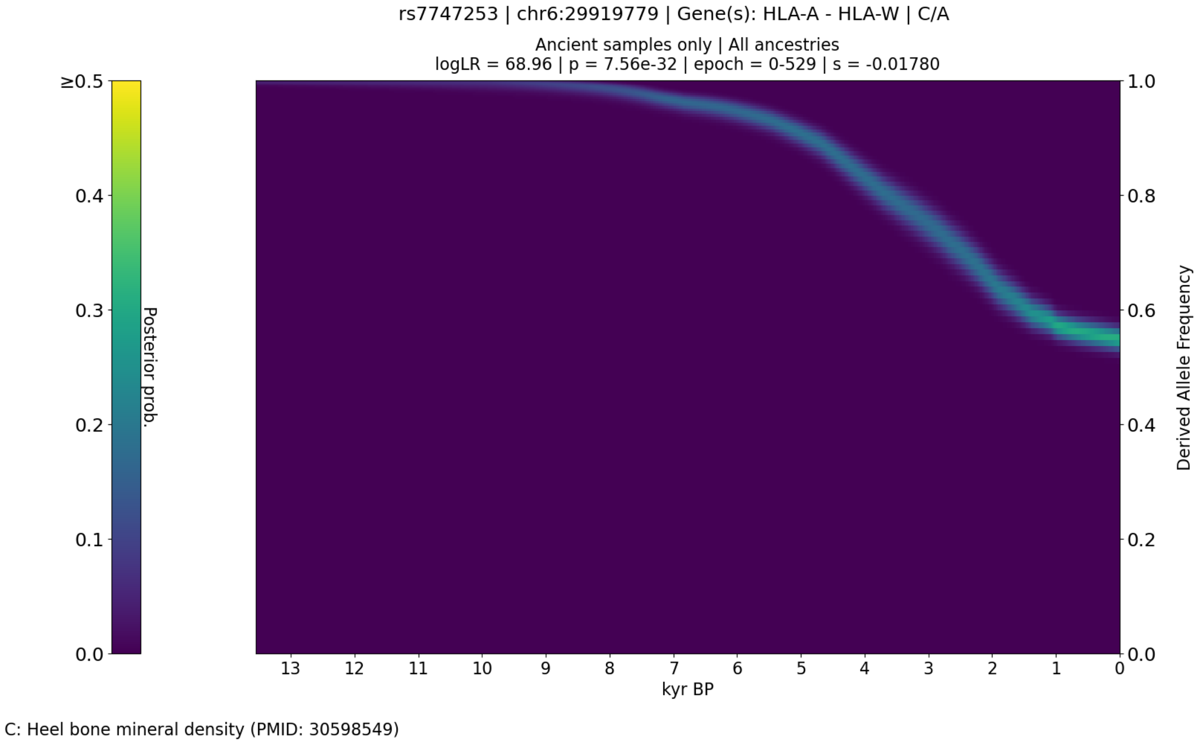


**Figure S43.** CLUES plot of the aDNA time series analysis for all West Eurasian samples in the imputed dataset, showing the posterior probability of the derived allele frequency trajectory for rs7747253 the most significant SNP in the selection peak spanning chr6:29705659-32832786.

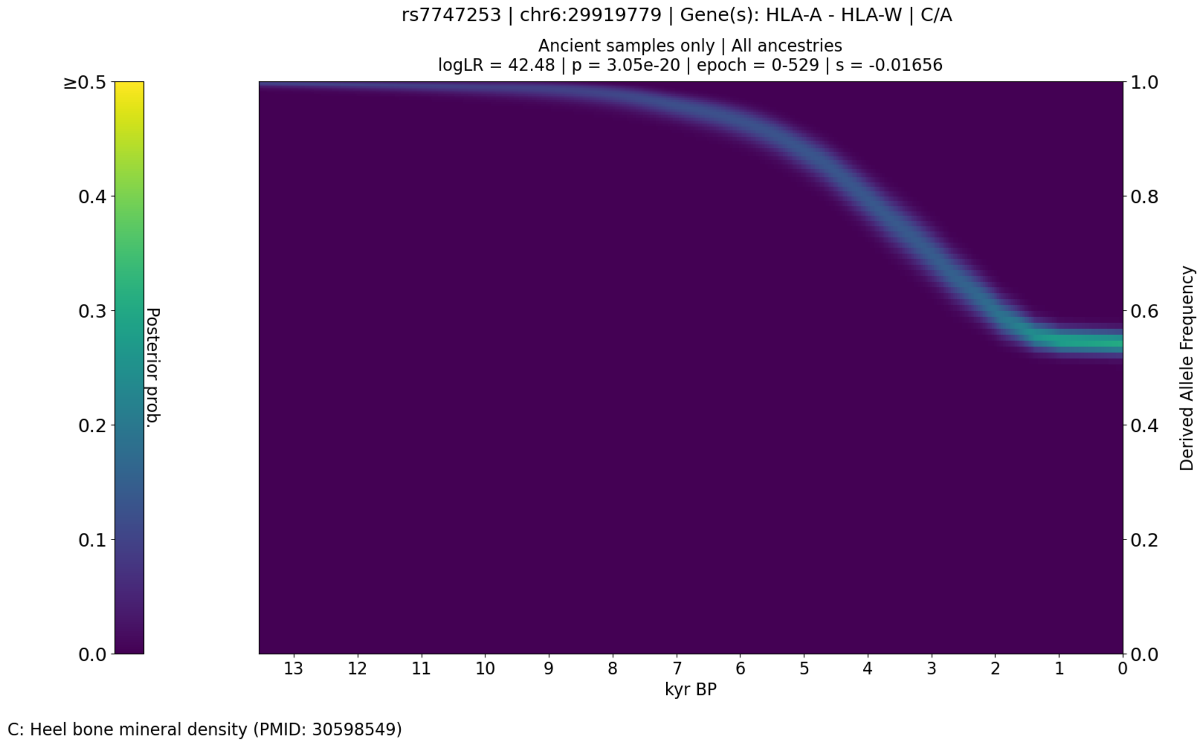


**Figure S44.** CLUES plot of the aDNA time series analysis for all West Eurasian samples in the genotype likelihood dataset (i.e., non-imputed), showing the posterior probability of the derived allele frequency trajectory for rs7747253.

The seventh peak spanned the region chr6:29705659-32832786, with the most significant SNP being rs7747253 (*HLA-W*; p=7.56e-32; s=-0.018), an upstream gene variant/intergenic variant associated with heel bone mineral density [70] in the GWAS Catalog (r2023-04-07).

Peak 8: TNKS

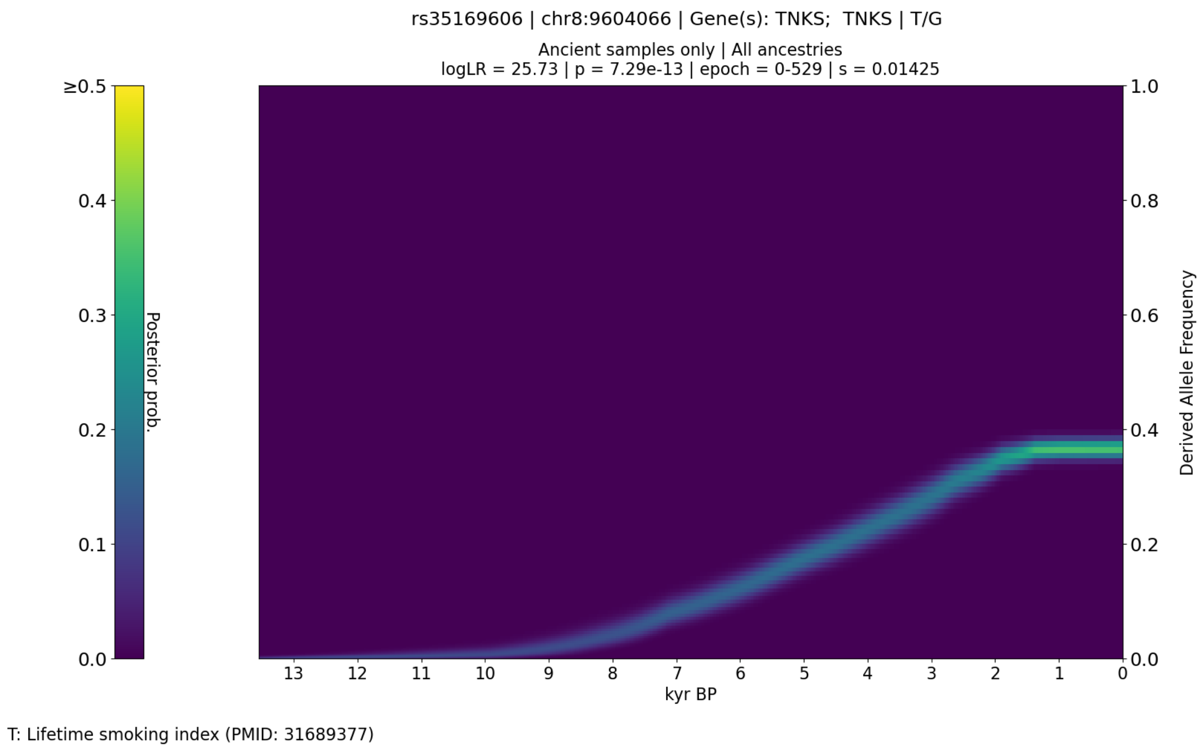


**Figure S45.** CLUES plot of the aDNA time series analysis for all West Eurasian samples in the imputed dataset, showing the posterior probability of the derived allele frequency trajectory for rs35169606 the most significant SNP in the selection peak spanning chr8:8168474-11856864.

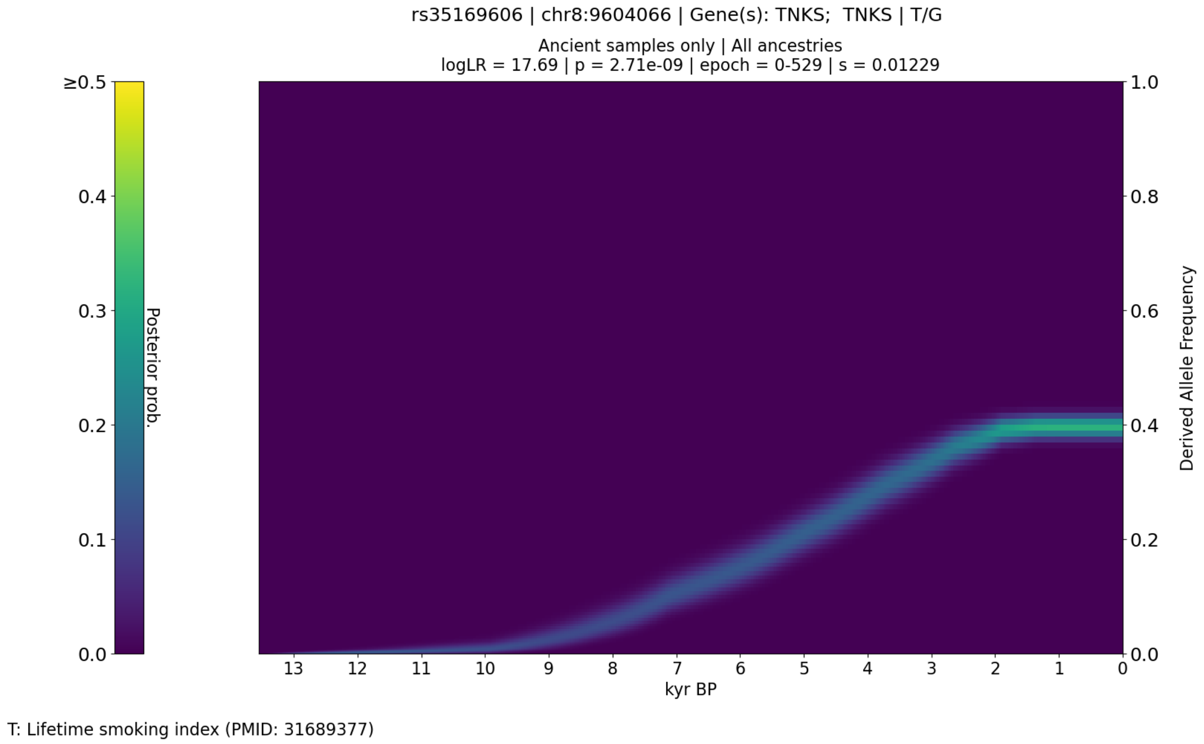


**Figure S46.** CLUES plot of the aDNA time series analysis for all West Eurasian samples in the genotype likelihood dataset (i.e., non-imputed), showing the posterior probability of the derived allele frequency trajectory for rs35169606.

The eighth peak spanned the region chr8:8168474-11856864, with the most significant SNP being rs35169606 (*TNKS*; p=7.29e-13; s=0.014), an intron variant associated with a body shape index [71], lifetime smoking index [72], waist circumference adjusted for body mass index [71], waist-hip index [71], and waist-to-hip ratio adjusted for BMI [71] in the GWAS Catalog (r2023-04-07).

#### Peak 9: FADS2

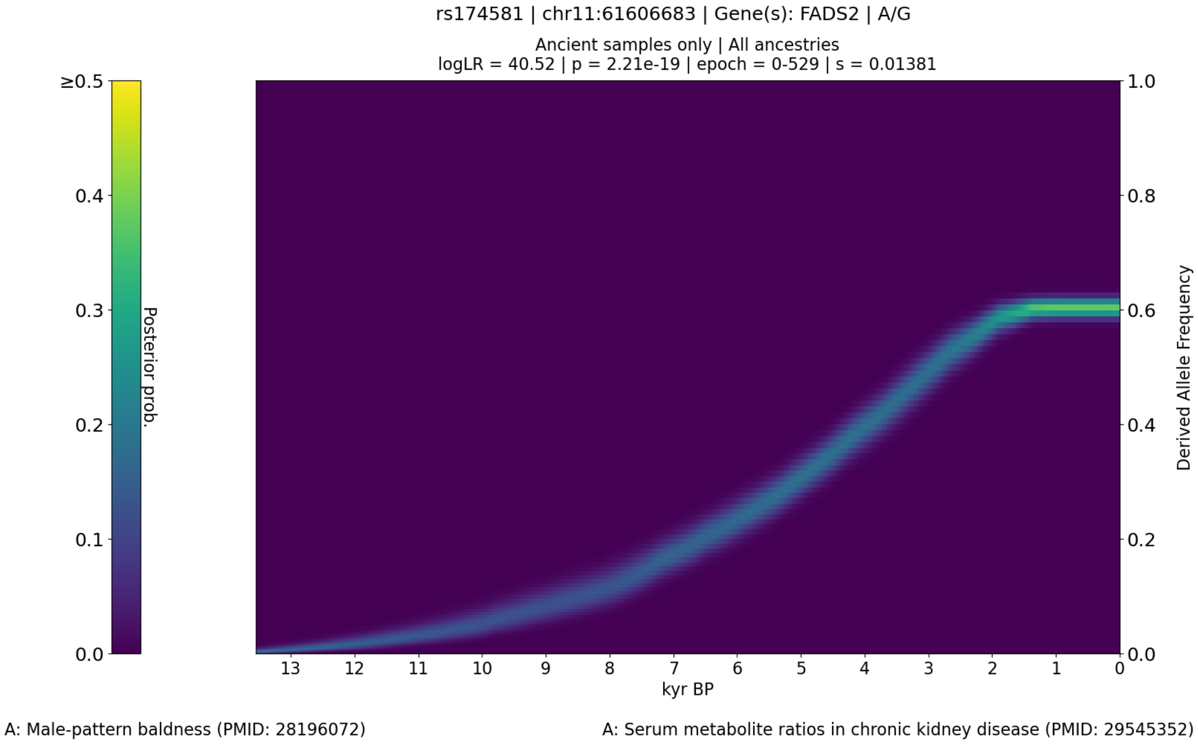


**Figure S47.** CLUES plot of the aDNA time series analysis for all West Eurasian samples in the imputed dataset, showing the posterior probability of the derived allele frequency trajectory for rs174581 the most significant SNP in the selection peak spanning chr11:61543499-61663691.

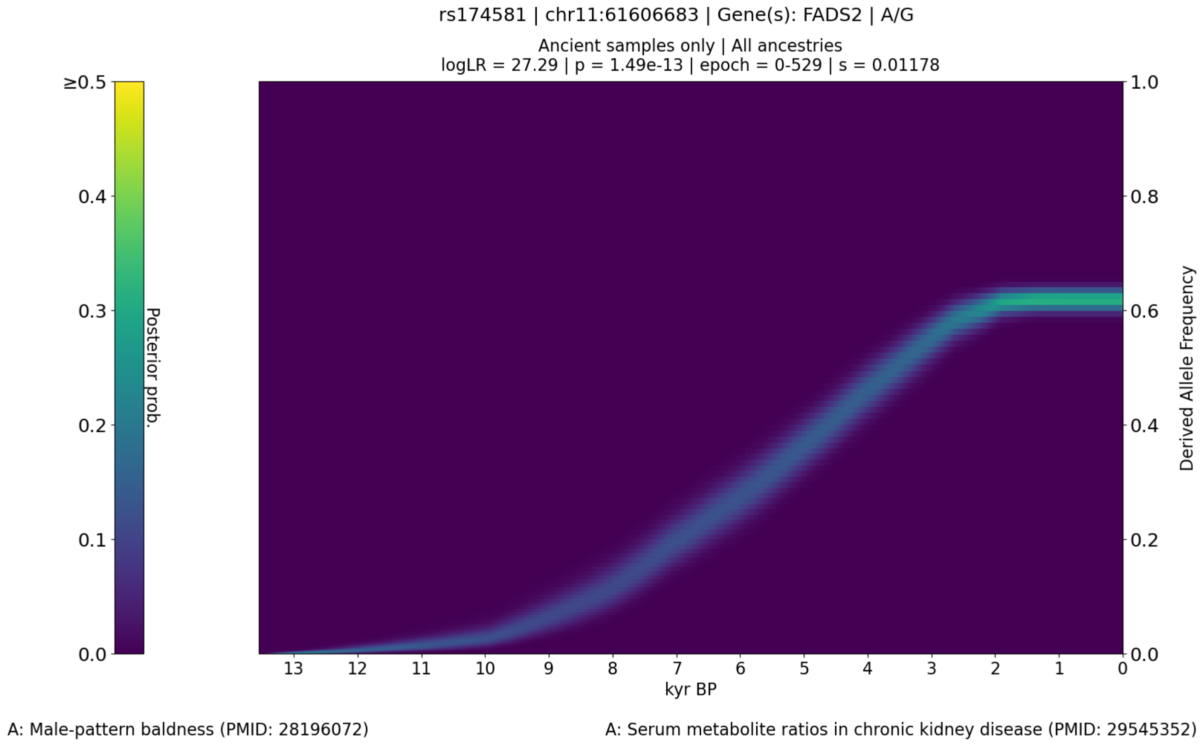


**Figure S48.** CLUES plot of the aDNA time series analysis for all West Eurasian samples in the genotype likelihood dataset (i.e., non-imputed), showing the posterior probability of the derived allele frequency trajectory for rs174581.

The ninth peak spanned the region chr11:61543499-61663691, with the most significant SNP being rs174581 (*FADS2*; p=2.21e-19; s=0.014), an intron variant associated with 1-margaroyl-2-linoleoyl-GPC levels [42], 1-oleoyl-2-eicosapentaenoyl-GPC levels [42], 1-palmitoyl-2-linoleoyl-gpc levels [42], 1-pentadecanoyl-2-linoleoyl-GPC levels [42], 1-stearoyl-2-linoleoyl-gpc levels [42], arachidonoylcholine levels [42], cholesteryl ester levels [73,74], diacylglycerol levels [74], fatty acid levels [74], lysophosphatidylcholine levels [74], male-pattern baldness [75], phosphatidate levels [74], phosphatidylcholine levels [73,74], phosphatidylethanolamine levels [74], phosphatidylglycerol levels [74], phosphatidylinositol levels [74], phosphatidylserine levels [74], phospholipids to total lipids ratio in small VLDL [46], serum metabolite ratios in chronic kidney disease [76], sphingomyelin levels [74], and triacylglycerol levels [74] in the GWAS Catalog (r2023-04-07).

#### Peak 10: HECTD4

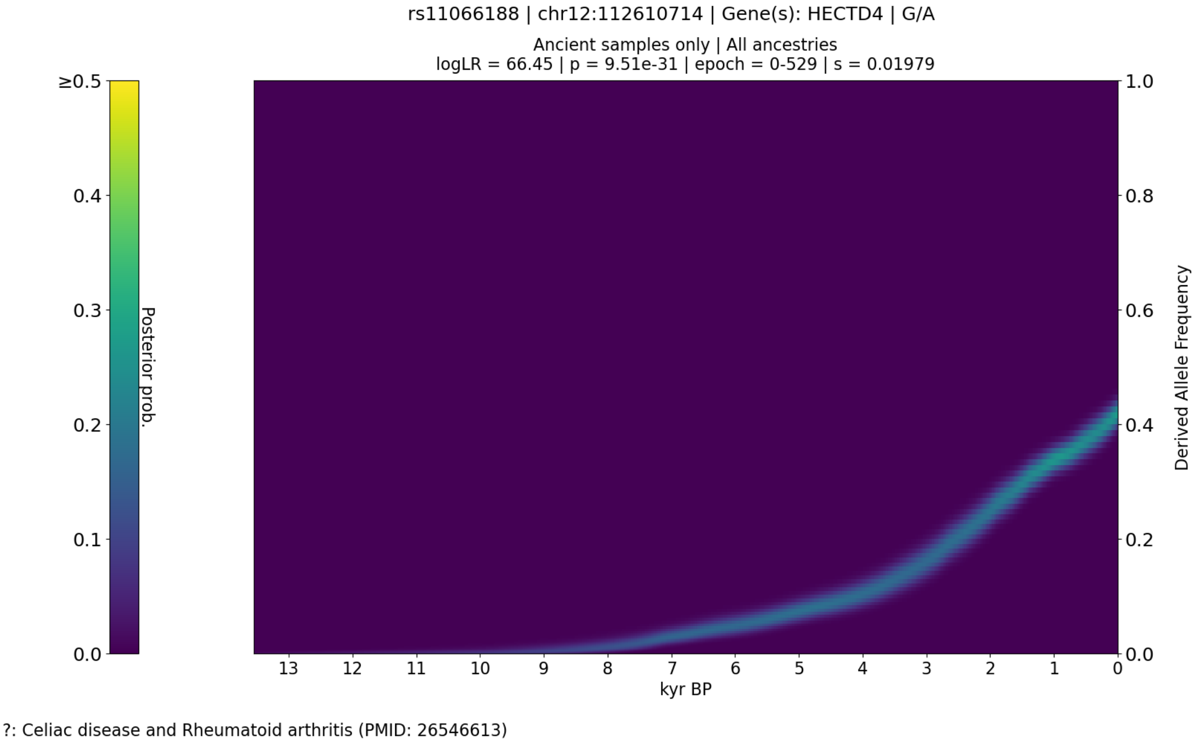


**Figure S49.** CLUES plot of the aDNA time series analysis for all West Eurasian samples in the imputed dataset, showing the posterior probability of the derived allele frequency trajectory for rs11066188 the most significant SNP in the selection peak spanning chr12:111833788-113359157.

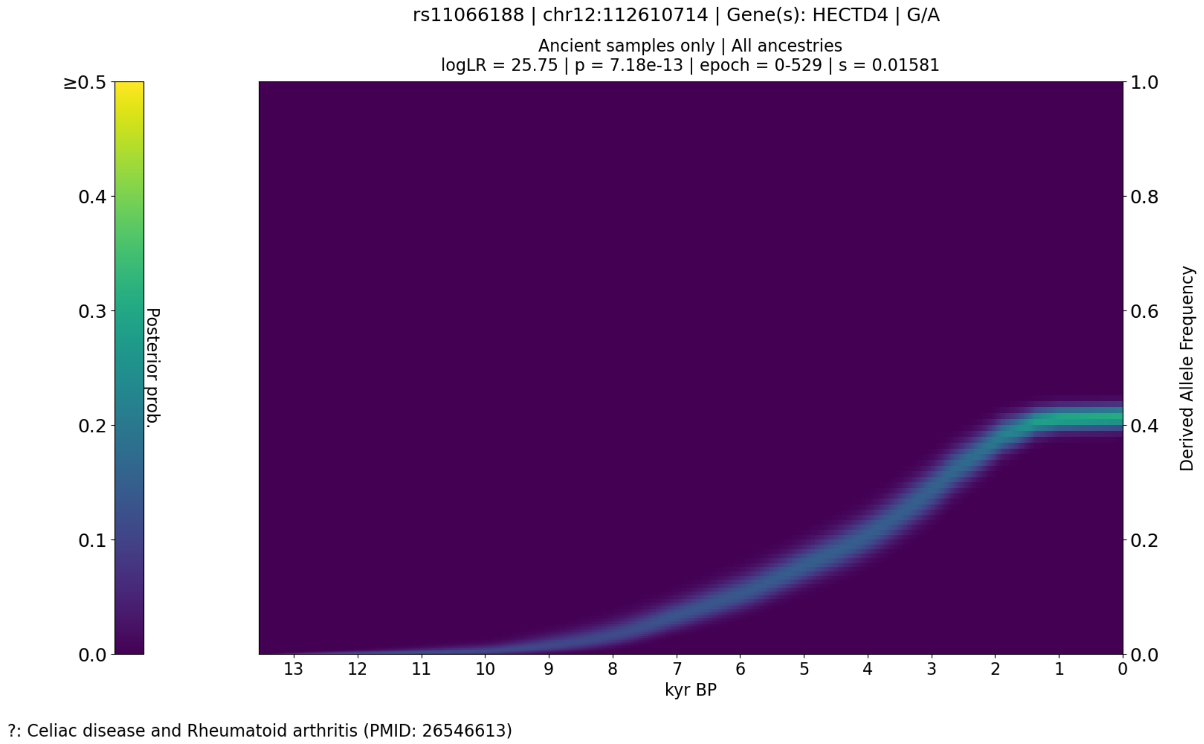


**Figure S50.** CLUES plot of the aDNA time series analysis for all West Eurasian samples in the genotype likelihood dataset (i.e., non-imputed), showing the posterior probability of the derived allele frequency trajectory for rs11066188.

The tenth peak spanned the region chr12:111833788-113359157, with the most significant SNP being rs11066188 (*HECTD4*; p=9.51e-31; s=0.020), an intron variant associated with body mass index (MTAG) [50], celiac disease and Rheumatoid arthritis [77], diastolic blood pressure x depressive symptoms interaction (2df test) [78], heart failure [79], ischemic stroke or factor VII levels [80], ischemic stroke or factor VIII levels [80], ischemic stroke or factor XI levels [80], ischemic stroke or fibrinogen levels [80], ischemic stroke or plasminogen activator inhibitor 1 levels [80], ischemic stroke or tissue plasminogen activator levels [80], ischemic stroke or von Willebrand factor levels [80], and spleen volume [81] in the GWAS Catalog (r2023-04-07).

Peak 11: KANSL1

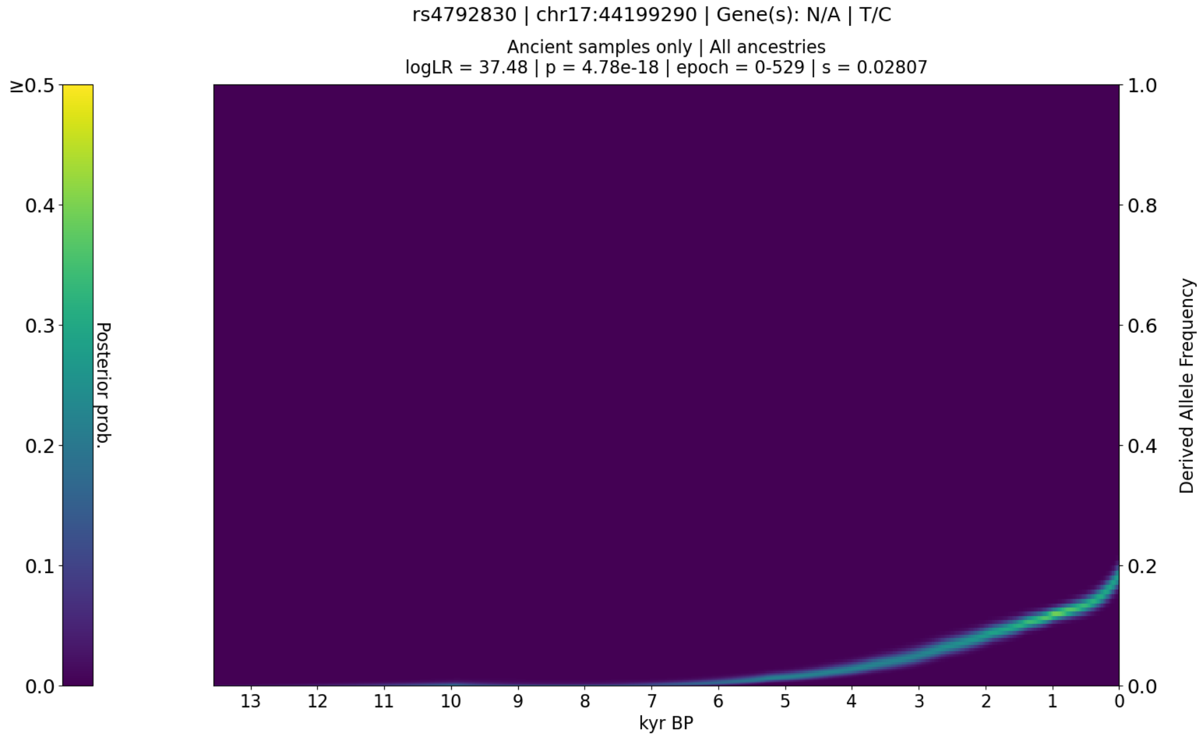


**Figure S51.** CLUES plot of the aDNA time series analysis for all West Eurasian samples in the imputed dataset, showing the posterior probability of the derived allele frequency trajectory for rs4792830 the most significant SNP in the selection peak spanning chr17:44086267-45055107.

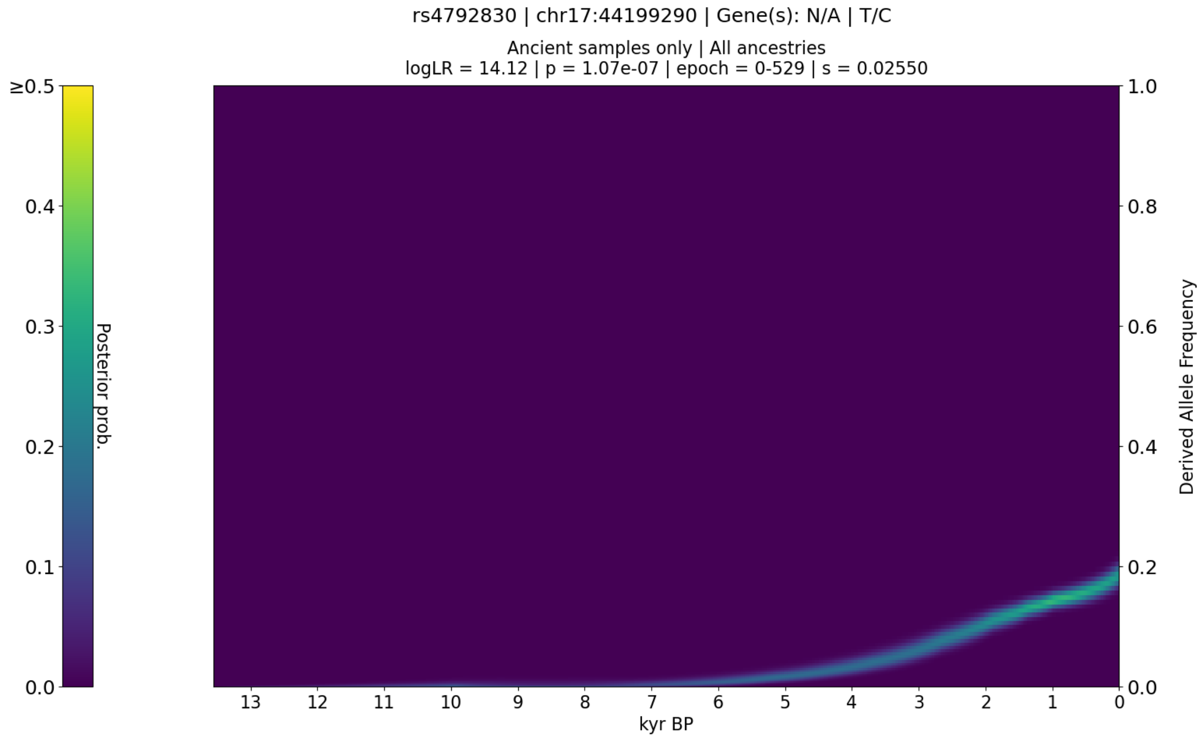


**Figure S52.** CLUES plot of the aDNA time series analysis for all West Eurasian samples in the genotype likelihood dataset (i.e., non-imputed), showing the posterior probability of the derived allele frequency trajectory for rs4792830.

The eleventh peak spanned the region chr17:44086267-45055107, with the most significant SNP being rs4792830 (*KANSL1*; p=4.78e-18; s=0.028), an intron variant associated with pulse pressure [82] in the GWAS Catalog (r2023-04-07).

#### Selection in simulations with Ancestral Paintings

CLUES analysis of all simulated SNPs (n=2,948), in the frequency-paired aDNA simulation, identified 22 (0.75%) genome-wide significant SNPs (p<5e-8) (Supplementary Table 5) We identified no genome-wide significant sweep regions across all ancestries (Figure S53), indicating that the false positive rate of our ancestry stratified selection analysis is low.

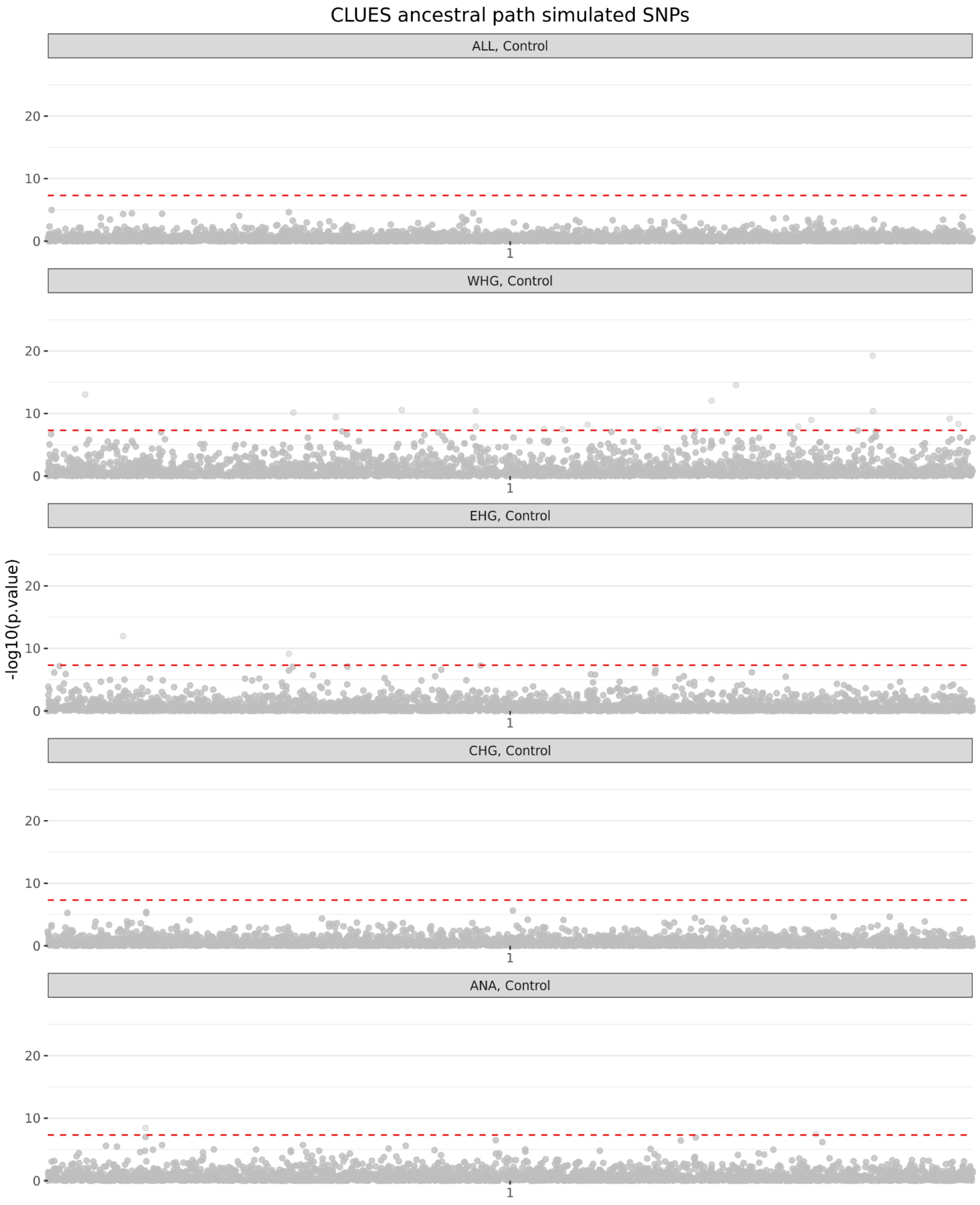


**Figure S53.** Manhattan plot of the p-values from running CLUES on a neutral simulation of chr3, using the inferred ancestral paths of each ancestry. The first row shows results for all ancient samples considered in aggregate, and each subsequent row shows the results conditional on one of the four specific ancestral paintings:  WHG (Western Hunter-gatherers), EHG (Eastern Hunter-gatherers), CHG (Caucasus Hunter-gatherers), and ANA (Anatolian Farmers).

##### Selection in aDNA with Ancestral Paintings

CLUES analysis of all GWAS (n=33,341) and Control group SNPs (n=33,341) in the aDNA with Ancestral Paintings dataset identified 1,914 genome-wide significant SNPs ($p<5e-8$); 1,357 in the GWAS group (70.9%) and 557 in the Control group (29.1%) (Supplementary Table 7). Within the GWAS group, we identified 19 non-overlapping genome-wide significant sweep regions across all ancestries, and 2 in the Control group (Figure S54 and S55).

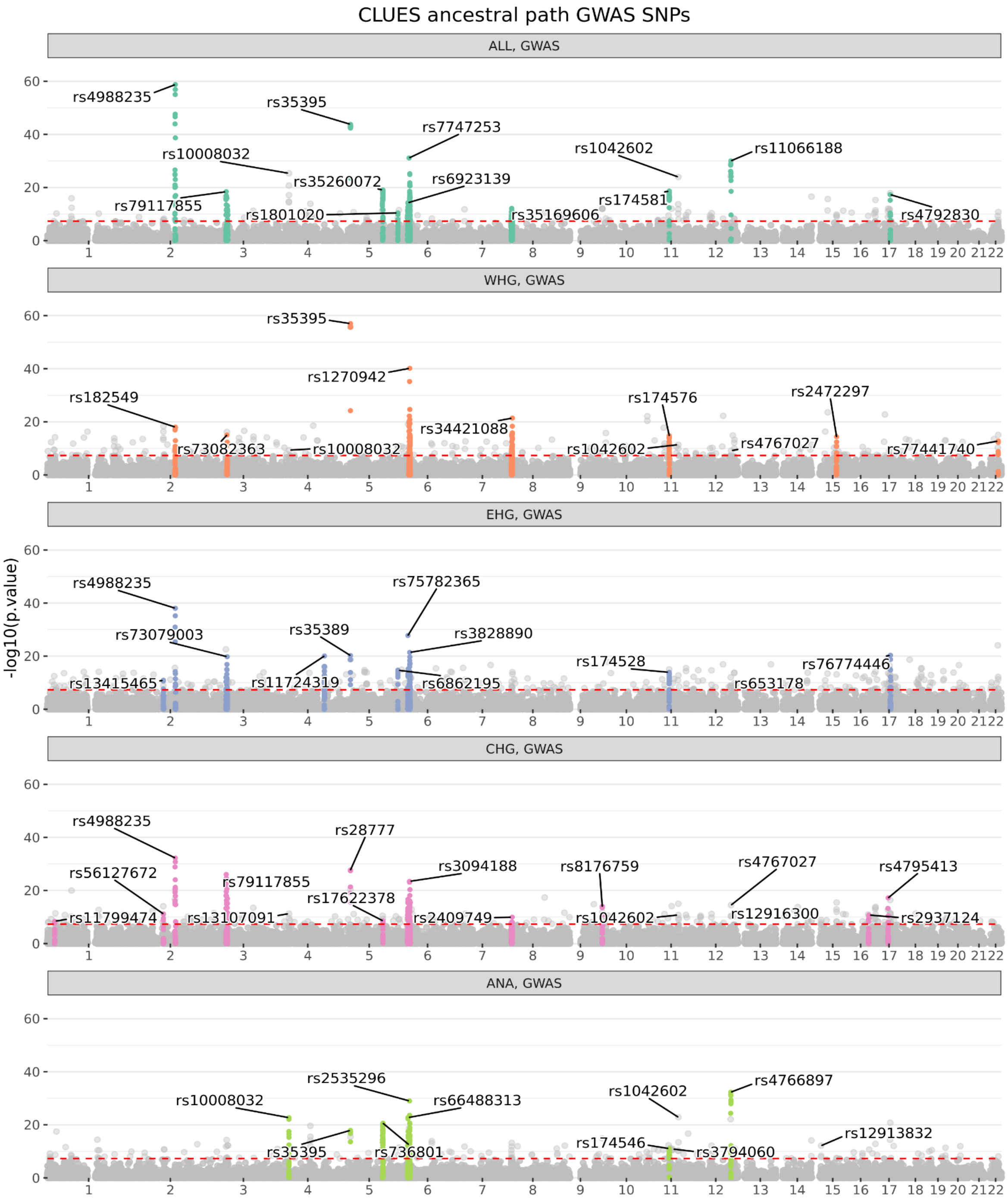


**Figure S54.** Manhattan plot of the p-values from running CLUES on an aDNA time series conditioned on ancestry paintings from all West Eurasian samples in the imputed dataset for GWAS SNPs from the GWAS Catalog. The first row shows results for all ancient samples considered in aggregate, and each subsequent row shows the results conditional on one of the four specific ancestral paintings: WHG (Western Hunter-gatherers), EHG (Eastern Hunter-gatherers), CHG (Caucasus Hunter-gatherers), and ANA (Anatolian Farmers).

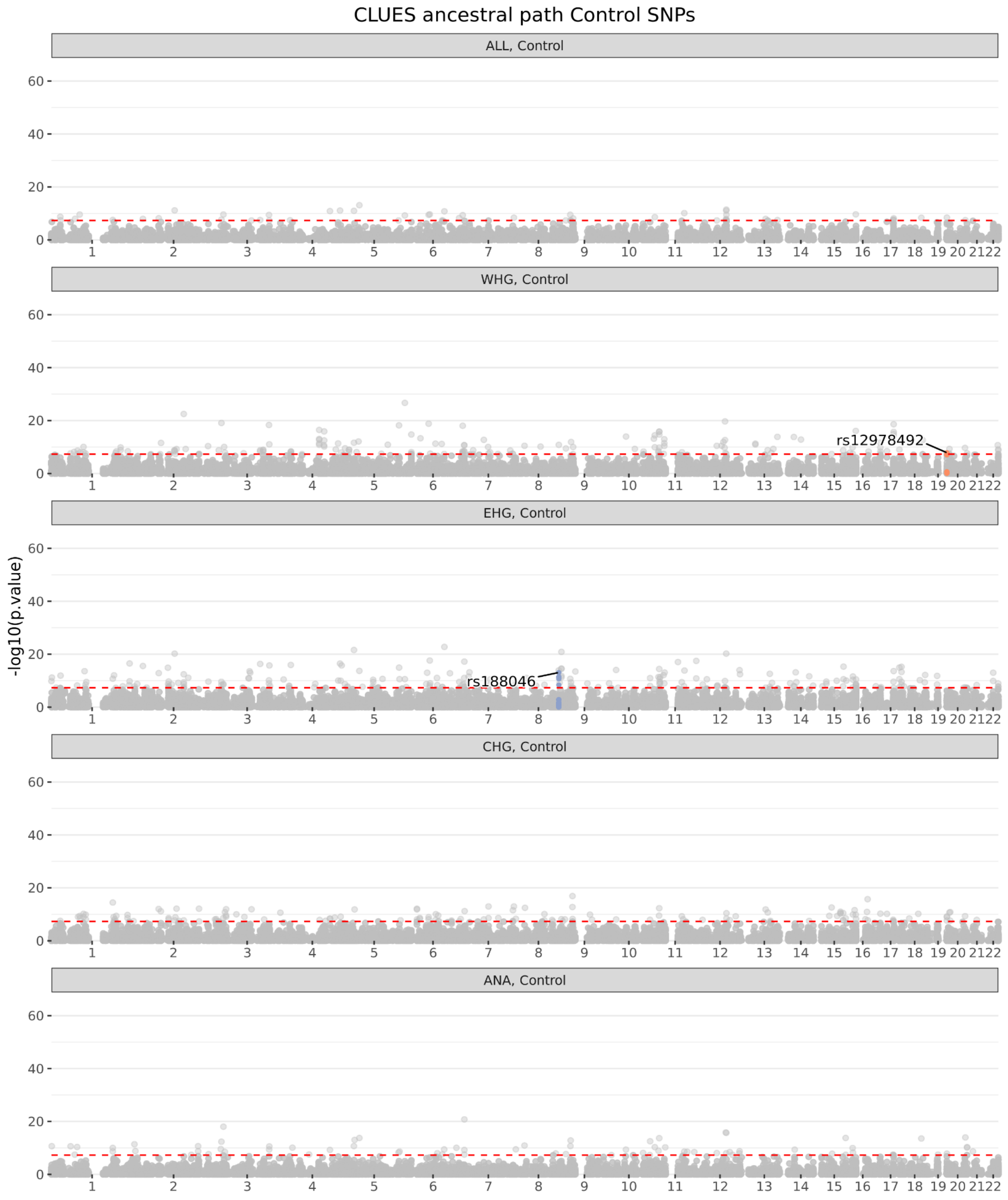


**Figure S55.** Manhattan plot of the p-values from running CLUES on an aDNA time series conditioned on ancestry paintings from all West Eurasian samples in the imputed dataset for Control SNPs. The first row shows results for all ancient samples considered in aggregate, and each subsequent row shows the results conditional on one of the four specific ancestral paintings: WHG (Western Hunter-gatherers), EHG (Eastern Hunter-gatherers), CHG (Caucasus Hunter-gatherers), and ANA (Anatolian Farmers).

#### Peak 1: RP11-415K20.1

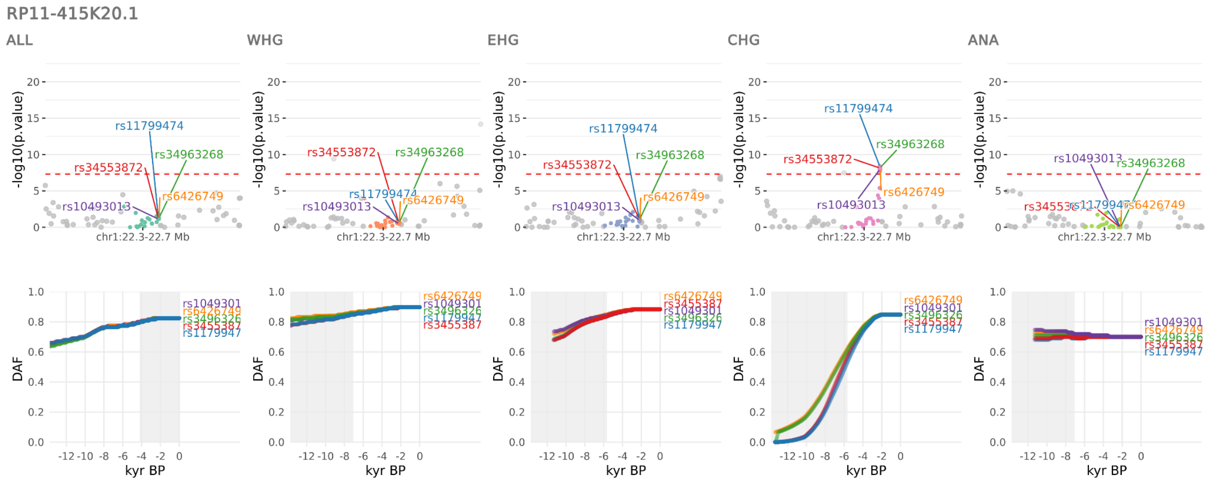


**Figure S56.** Selection at the *RP11-415K20.1* locus, spanning chr1:22266330-22711473. Results for the pan-ancestry analysis (ALL) plus each of the four marginal ancestries: Western hunter-gatherers (WHG), Eastern hunter-gatherers (EHG), Caucasus hunter-gatherers (CHG) and Anatolian farmers (ANA). Row one shows zoomed Manhattan plots of the p-values for each ancestry, and row two shows allele trajectories for the top SNPs across all ancestries.

The first peak spanned the region chr1:22266330-22711473, with the five most significant SNPs (across ancestries) and their associations in the GWAS Catalog (r2023-04-07) being:

1. rs11799474 (*RP11-415K20.1*; CHG: p=3.89e-09; s=0.015), an intergenic variant associated with red cell distribution width [83,84].

2. rs34963268 (*RP11-415K20.1*; CHG: p=6.27e-09; s=0.014), an intergenic variant associated with heel bone mineral density [85,86], colorectal cancer [87], mean reticulocyte volume [88], and red blood cell count [89].

3. rs34553872 (*RP11-415K20.1*; CHG: p=6.92e-09; s=0.015), a regulatory region variant associated with heel bone mineral density [70].

4. rs6426749 (*RP11-415K20.1*; CHG: p=1.07e-08; s=0.013), an intergenic variant associated with femoral neck bone mineral density [90] and red cell distribution width [91].

5. rs10493013 (*RP11-415K20.1*; CHG: p=1.21e-08; s=0.015), an intergenic variant associated with total body bone mineral density [92].

#### Peak 2: LINC01104

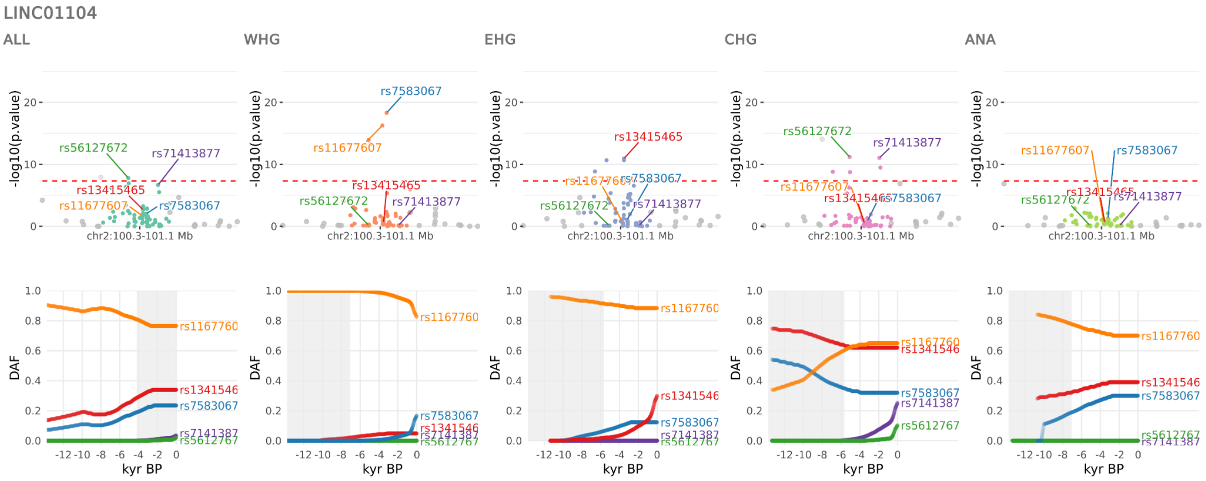


**Figure S57.** Selection at the *LINC01104* locus, spanning chr2:100309124-101050353. Results for the pan-ancestry analysis (ALL) plus each of the four marginal ancestries: Western hunter-gatherers (WHG), Eastern hunter-gatherers (EHG), Caucasus hunter-gatherers (CHG) and Anatolian farmers (ANA). Row one shows zoomed Manhattan plots of the p-values for each ancestry, and row two shows allele trajectories for the top SNPs across all ancestries.

The second peak spanned the region chr2:100309124-101050353, with the five most significant SNPs (across ancestries) and their associations in the GWAS Catalog (r2023-04-07) being:

1. rs7583067 (*LINC01104*; WHG: p=4.98e-19; s=0.044), an intergenic variant associated with general cognitive ability [93] and HDL cholesterol levels [94].
2. rs56127672 (*AFF3*; CHG: p=6.45e-12; s=0.072), an intron variant associated with intelligence (MTAG) [95].
3. rs13415465 (*AFF3*; EHG: p=1.18e-11; s=0.027), an upstream gene variant associated with autoimmune traits (pleiotropy) [96], red blood cell count [89], rheumatoid arthritis [96], and serum total protein level [97].
4. rs11677607 (*AFF3*; WHG: p=5.47e-17; s=-0.036), an intron variant associated with body mass index [98].

5. rs71413877 (*LONRF2*; CHG: p=9.26e-12; s=0.030), an intron variant associated with educational attainment (years of education) [99,100], attention deficit hyperactivity disorder or autism spectrum disorder or intelligence [101], educational attainment (MTAG) [99], general cognitive ability [93], highest math class taken (MTAG) [99], household income (MTAG) [102], intelligence [103], and intelligence (MTAG) [95].

#### Peak 3: MCM6

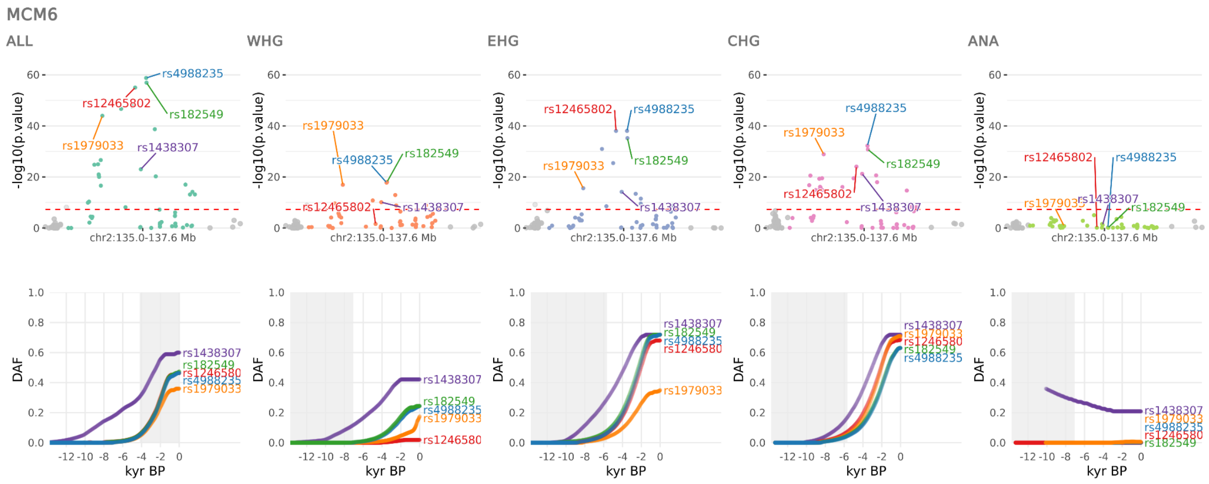


**Figure S58.** Selection at the *MCM6* locus, spanning chr2:134963892-137613935. Results for the pan-ancestry analysis (ALL) plus each of the four marginal ancestries: Western hunter-gatherers (WHG), Eastern hunter-gatherers (EHG), Caucasus hunter-gatherers (CHG) and Anatolian farmers (ANA). Row one shows zoomed Manhattan plots of the p-values for each ancestry, and row two shows allele trajectories for the top SNPs across all ancestries.

The third peak spanned the region chr2:134963892-137613935, with the five most significant SNPs (across ancestries) and their associations in the GWAS Catalog (r2023-04-07) being:

1. rs4988235 (*MCM6*; ALL: p=1.68e-59; s=0.019), an intron variant associated with lactase persistence, 1,5-anhydroglucitol levels [42,43], body mass index [44,45], acetate levels [46], bacilli A abundance in stool [47], bifidobacterium bifidum abundance in stool [47], blood protein levels [48], body fat percentage and LDL-C (pairwise) [49], body mass index (MTAG) [50], body mass index and LDL-C (pairwise) [49], brevibacillaceae abundance in stool [47], brevibacillales abundance in stool [47], broad bean liking [51], degree of unsaturation [46], free cholesterol to total lipids ratio in small HDL [46], gut microbiota abundance (phylum Actinobacteria) [36], hip circumference [52],

indolepropionate levels [42], lactase-phlorizin hydrolase levels [53], lactase-phlorizin hydrolase levels [37], lactobacillus B abundance in stool [47], lactobacillus B ruminis abundance in stool [47], medication use for hyperlipidemia (number of purchases) [54], phospholipids to total lipids ratio in medium HDL [46], turicibacter sp001543345 abundance in stool [47], and x-11795 levels [42].

2. rs182549 (*MCM6*; ALL: p=1.19e-57; s=0.019), an intron variant associated with 1,5-anhydroglucitol levels [104], bifidobacterium adolescentis abundance in stool [47], bread consumption (slices per week) [105], gut microbiota abundance (class Actinobacteria) [36], gut microbiota abundance (family Bifidobacteriaceae) [36], gut microbiota abundance (genus Bifidobacterium) [36], gut microbiota abundance (k_Bacteria.p_Actinobacteria.c_Actinobacteria. o_Bifidobacteriales.f_Bifidobacteriaceae.g_Bifidobacterium) [106], gut microbiota abundance (k_Bacteria.p_Actinobacteria.c_Actinobacteria. o_Bifidobacteriales.f_Bifidobacteriaceae) [106], gut microbiota abundance (k_Bacteria.p_Actinobacteria.c_Actinobacteria. o_Bifidobacteriales)[106], gut microbiota abundance (k_Bacteria.p_Actinobacteria.c_Actinobacteria) [106], gut microbiota abundance (k_Bacteria.p_Actinobacteria) [106], gut microbiota abundance (order Bifidobacteriales) [36], milky sweets liking [51], negativibacillus abundance in stool [47], ratio of polyunsaturated fatty acids to monounsaturated fatty acids [46], and ratio of polyunsaturated fatty acids to total fatty acids [46].

3. rs12465802 (*R3HDM1*; ALL: p=9.56e-56; s=0.019), an intron variant associated with gut microbiota abundance (genus Bifidobacterium) [36], lactase-phlorizin hydrolase levels [37], mosquito bite size [38], and urinary metabolite levels in chronic kidney disease [39].

4. rs1979033 (*CCNT2*; ALL: p=1.12e-44; s=0.020), an intron variant associated with age at menopause [89], gut microbiota abundance (family Bifidobacteriaceae) [36], gut microbiota abundance (genus Bifidobacterium) [36], gut microbiota abundance (order Bifidobacteriales) [36], and lactase-phlorizin hydrolase level in Chronic kidney disease with hypertension and no diabetes [61].

5. rs6759321 (R3HDM1; ALL: p=2.32e-48; s=0.018), an intron variant associated with hand grip strength [107].

#### Peak 4: CCDC12

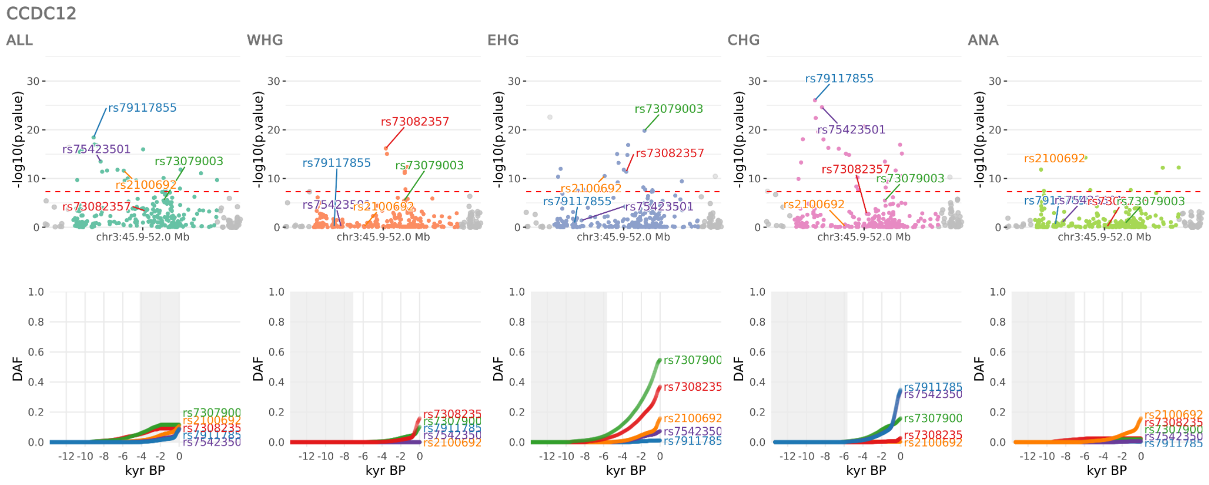

**Figure S59.** Selection at the *CCDC12* locus, spanning chr3:45943595-52029991. Results for the pan-ancestry analysis (ALL) plus each of the four marginal ancestries: Western hunter-gatherers (WHG), Eastern hunter-gatherers (EHG), Caucasus hunter-gatherers (CHG) and Anatolian farmers (ANA). Row one shows zoomed Manhattan plots of the p-values for each ancestry, and row two shows allele trajectories for the top SNPs across all ancestries.

The fourth peak spanned the region chr3:45943595-52029991, with the five most significant SNPs (across ancestries) and their associations in the GWAS Catalog (r2023-04-07) being:

1. rs79117855 (*CCDC12*; CHG: p=8.95e-27; s=0.034), an intergenic variant associated with blood protein levels [55].
2. rs73079003 (*CDHR4*; EHG: p=1.61e-20; s=0.021), a missense variant associated with glucose levels [97] and thioredoxin domain-containing protein 12 levels [37].
3. rs73082357 (*QRICH1*; WHG: p=6.71e-17; s=0.054), an intron variant associated with mood instability [108].
4. rs2100692 (*CDC25A*; ANA: p=5.18e-15; s=0.035), a downstream gene variant associated with body mass index [109] and HDL cholesterol levels [94].
5. rs75423501 (*KIF9-AS1*; CHG: p=2.37e-25; s=0.034), an intron variant associated with blood protein levels [55].

#### Peak 5: RNA5SP158

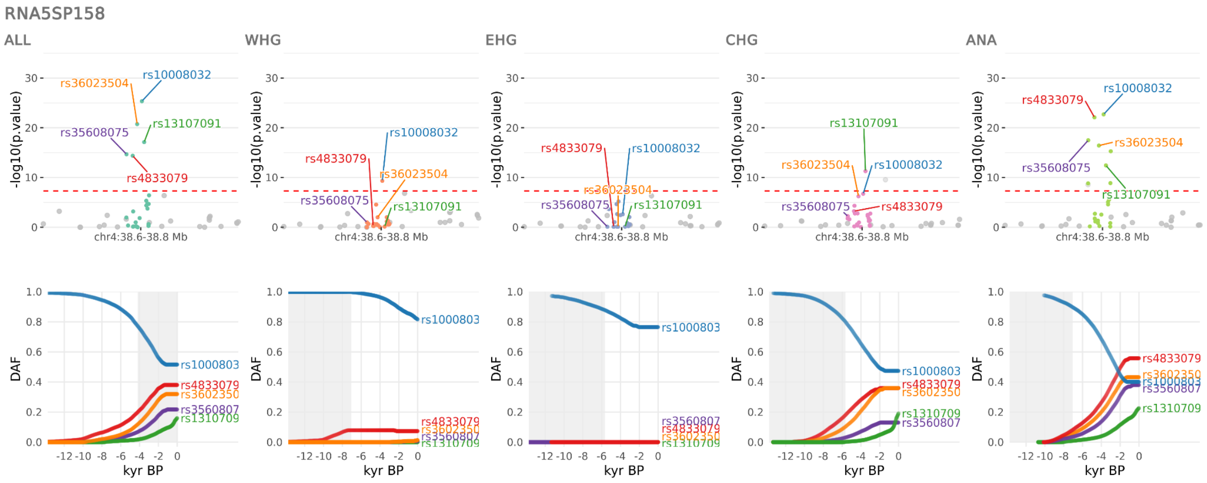

**Figure S60.** Selection at the *RNA5SP158* locus, spanning chr4:38591188-38816338. Results for the pan-ancestry analysis (ALL) plus each of the four marginal ancestries: Western hunter-gatherers (WHG), Eastern hunter-gatherers (EHG), Caucasus hunter-gatherers (CHG) and Anatolian farmers (ANA). Row one shows zoomed Manhattan plots of the p-values for each ancestry, and row two shows allele trajectories for the top SNPs across all ancestries.

The fifth peak spanned the region chr4:38591188-38816338, with the five most significant SNPs (across ancestries) and their associations in the GWAS Catalog (r2023-04-07) being:

1. rs10008032 (RNA5SP158; ALL: p=4.57e-26; s=-0.016), a regulatory region variant associated with allergic disease (asthma, hay fever or eczema) [110].

2. rs13107091 (*TLR10*; ALL: p=7.67e-18; s=0.030), an intergenic variant associated with atopic asthma [109].

3. rs4833079 (*AC021860.1 / RP11-617D20.1*; ANA: p=8.03e-23; s=0.017), an intron variant associated with body mass index [45] and neutrophil-to-lymphocyte ratio [111].

4. rs36023504 (*KLF3*; ALL: p=1.82e-21; s=0.017), an 3-prime UTR variant associated with adult body size [112], body mass index [89], red cell distribution width [89], reticulocyte count [88], reticulocyte fraction of red cells [88], and serum alkaline phosphatase levels [97].

5. rs35608075 (*RP11-213G21.2*; ANA: p=3.22e-18; s=0.018), an intergenic variant associated with male-pattern baldness [113].

Peak 6: KRT18P51

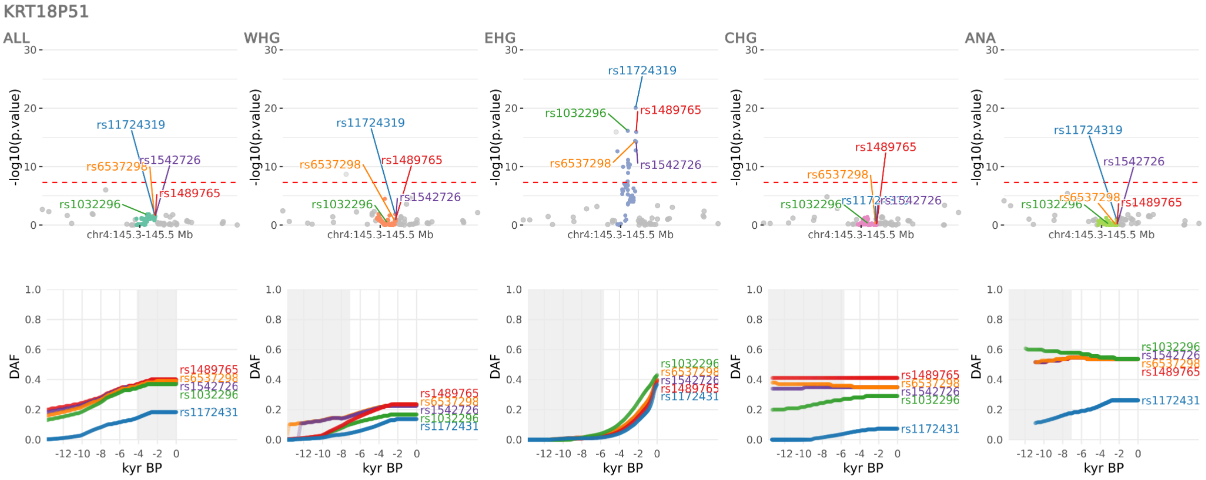


**Figure S61.** Selection at the *KRT18P51* locus, spanning chr4:145321006-145516378. Results for the pan-ancestry analysis (ALL) plus each of the four marginal ancestries: Western hunter-gatherers (WHG), Eastern hunter-gatherers (EHG), Caucasus hunter-gatherers (CHG) and Anatolian farmers (ANA). Row one shows zoomed Manhattan plots of the p-values for each ancestry, and row two shows allele trajectories for the top SNPs across all ancestries.

The sixth peak spanned the region chr4:145321006-145516378, with the five most significant SNPs (across ancestries) and their associations in the GWAS Catalog (r2023-04-07) being:

1. rs11724319 (*KRT18P51*; EHG: p=8.45e-21; s=0.025), an intergenic variant associated with post bronchodilator FEV1/FVC ratio [114].

2. rs1032296 (*RP11-361D14.2*; EHG: p=7.31e-17; s=0.020), a downstream gene variant associated with BMI at 6 months old [115], kidney volume [81], post bronchodilator FEV1/FVC ratio [114], waist-hip index [71], and waist-to-hip ratio adjusted for BMI [71].

3. rs1489765 (*KRT18P51*; EHG: p=1.15e-16; s=0.023), an intergenic variant associated with aparc-DKTatlas lh volume cuneus [116] and post bronchodilator FEV1/FVC ratio [114].

4. rs6537298 (*KRT18P51*; EHG: p=3.99e-15; s=0.022), an intergenic variant associated with aparc-a2009s lh volume G-cuneus [116], post bronchodilator FEV1 [114], and post bronchodilator FEV1/FVC ratio [114].

5. rs1542726 (*KRT18P51*; EHG: p=5.68e-15; s=0.023), an intergenic variant associated with haemorrhoidal disease [117], hip circumference adjusted for BMI [71], and post bronchodilator FEV1/FVC ratio [114].

#### Peak 7: SLC45A2

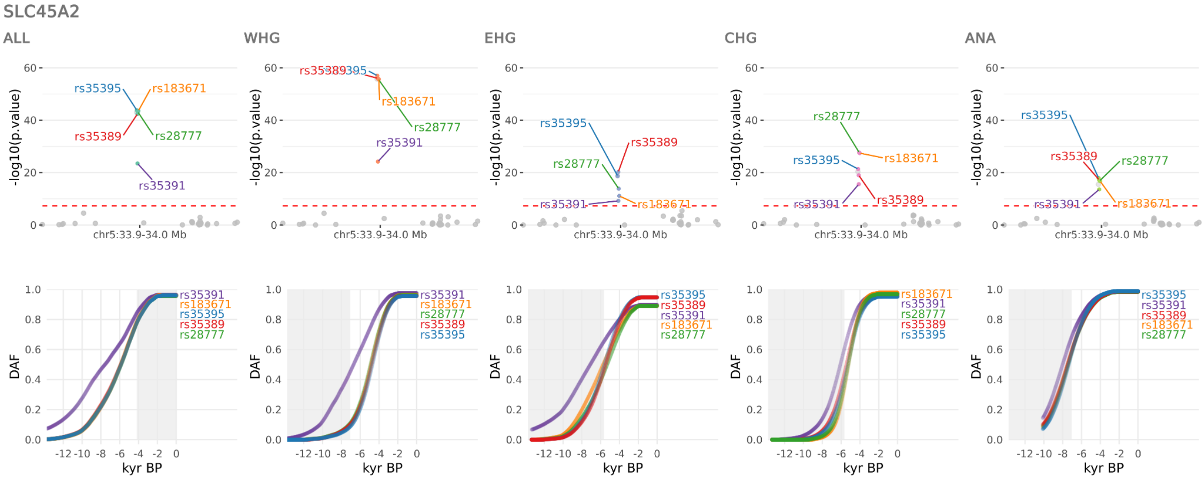


**Figure S62.** Selection at the *SLC45A2* locus, spanning chr5:33946571-33964210. Results for the pan-ancestry analysis (ALL) plus each of the four marginal ancestries: Western hunter-gatherers (WHG), Eastern hunter-gatherers (EHG), Caucasus hunter-gatherers (CHG) and Anatolian farmers (ANA). Row one shows zoomed Manhattan plots of the p-values for each ancestry, and row two shows allele trajectories for the top SNPs across all ancestries.

The seventh peak spanned the region chr5:33946571-33964210, with the five most significant SNPs (across ancestries) and their associations in the GWAS Catalog (r2023-04-07) being:

1. rs35395 (SLC45A2; WHG: p=9.79e-58; s=0.030), an intron variant associated with skin pigmentation [56].

2. rs28777 (SLC45A2; WHG: p=2.07e-56; s=0.029), an intron variant associated with black vs. blond hair color [118], black vs. red hair color [118], obstructive sleep apnea trait [119], and skin, hair and eye pigmentation [120].

3. rs35389 (*SLC45A2*; WHG: p=1.06e-56; s=0.029), an intron variant with no associations in the GWAS Catalog.

4. rs183671 (*SLC45A2*; WHG: p=2.20e-56; s=0.030), an intron variant associated with hair color [121], skin colour saturation [122], and skin pigmentation traits [123].

5. rs35391 (*SLC45A2*; WHG: p=6.25e-25; s=0.020), an intron variant associated with blond vs. brown/black hair color [124] and tanning [125].

#### Peak 8: IRF1

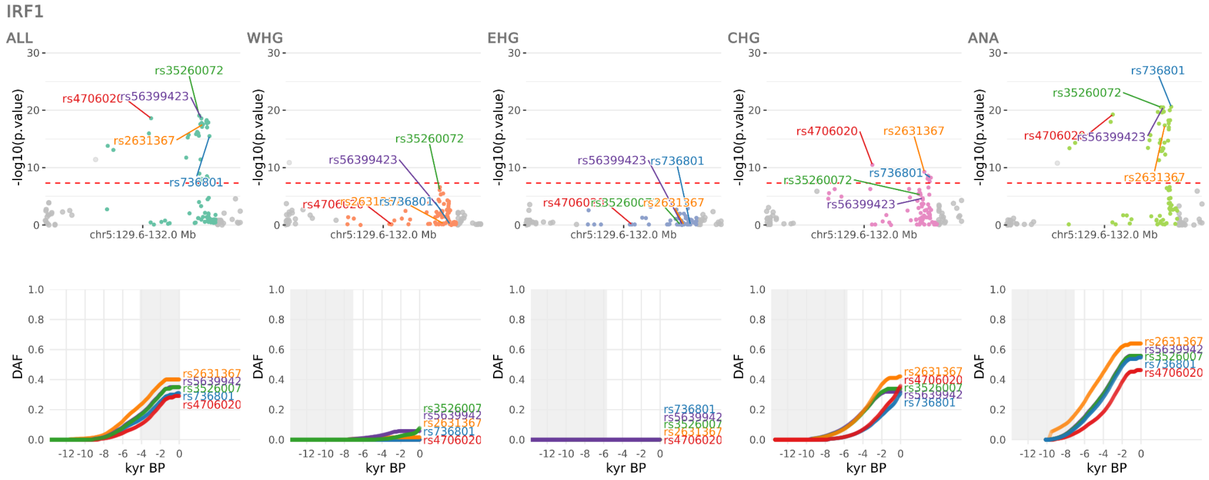


**Figure S63.** Selection at the *IRF1* locus, spanning chr5:129573666-131953510. Results for the pan-ancestry analysis (ALL) plus each of the four marginal ancestries: Western hunter-gatherers (WHG), Eastern hunter-gatherers (EHG), Caucasus hunter-gatherers (CHG) and Anatolian farmers (ANA). Row one shows zoomed Manhattan plots of the p-values for each ancestry, and row two shows allele trajectories for the top SNPs across all ancestries.

The eighth peak spanned the region chr5:129573666-131953510, with the five most significant SNPs (across ancestries) and their associations in the GWAS Catalog (r2023-04-07) being:

1. rs736801 (*IRF1*; ANA: p=2.48e-21; s=0.016), an intergenic variant associated with mosquito bite size [38].

2. rs35260072 (*P4HA2 / SLC22A4*; ANA: p=2.95e-21; s=0.016), an intron variant associated with histidine betaine (hercynine) levels [43] and itch intensity from mosquito bite [38].

3. rs4706020 (*CDC42SE2*; ANA: p=5.46e-20; s=0.017), an intron variant associated with itch intensity from mosquito bite [38] and itch intensity from mosquito bite adjusted by bite size [38].

4. rs2631367 (*AC034220.3 / SLC22A5*; ALL: p=2.88e-18; s=0.016), a 5-prime UTR variant associated with white blood cell count [88,89,97], monocyte count [88,91], acetylcarnitine levels [43], hexanoylcarnitine levels [43], linolenoylcarnitine levels [43], mean platelet volume [91], neutrophil count [97], palmitoylcarnitine levels (Metabolon platform) [43], pyruvate levels [46], and serum metabolite levels [67].

5. rs56399423 (*AC034220.3 / SLC22A4*; ANA: p=3.06e-21; s=0.016), an intron variant associated with inflammatory bowel disease [126].

#### Peak 9: SLC34A1

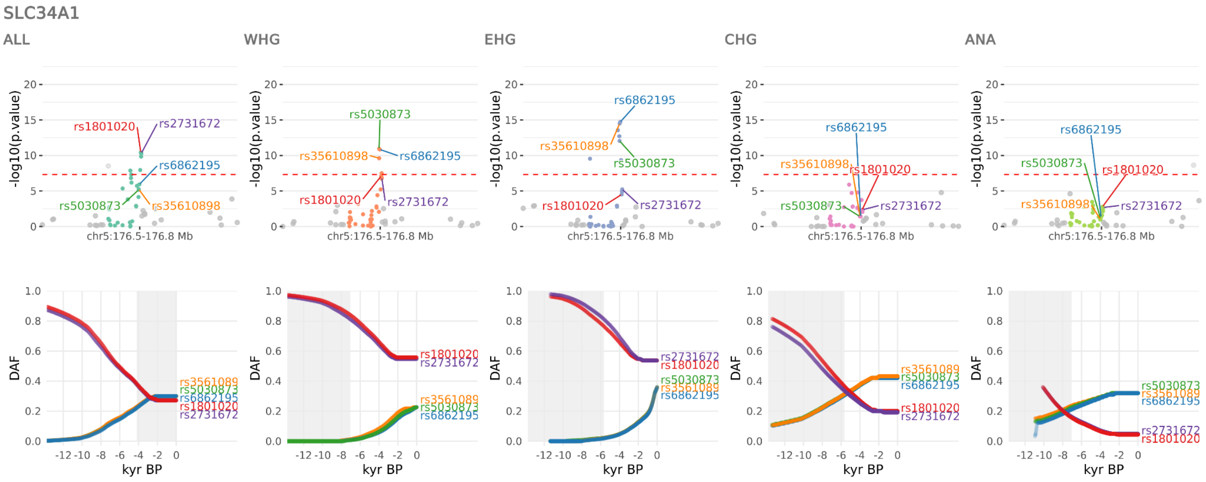


**Figure S64.** Selection at the *SLC34A1* locus, spanning chr5:176509193-176842474. Results for the pan-ancestry analysis (ALL) plus each of the four marginal ancestries: Western hunter-gatherers (WHG), Eastern hunter-gatherers (EHG), Caucasus hunter-gatherers (CHG) and Anatolian farmers (ANA). Row one shows zoomed Manhattan plots of the p-values for each ancestry, and row two shows allele trajectories for the top SNPs across all ancestries.

The ninth peak spanned the region chr5:176509193-176842474, with the five most significant SNPs (across ancestries) and their associations in the GWAS Catalog (r2023-04-07) being:

1. rs6862195 (*SLC34A1*; EHG: p=1.92e-15; s=0.025), an intron variant associated with estimated glomerular filtration rate [127].

2. rs5030873 (*SLC34A1*; EHG: p=8.83e-13; s=0.025), an synonymous variant associated with creatinine levels [46,128].

3. rs1801020 (*F12 / GRK6*; ALL: p=3.87e-11; s=-0.010), a 5-prime UTR variant associated with superoxide dismutase mitochondrial levels [53,68], thrombin levels [53,58], activated partial thromboplastin time [57], alpha-2-macroglobulin levels [58], aspartate levels [59], blood pressure [60], bMP and activin membrane-bound inhibitor homolog level in Chronic kidney disease with hypertension and no diabetes [61], bone morphogenetic protein 6 levels [62], bone morphogenetic protein 7 levels [58], cadherin-15 levels [37], glucosidase 2 subunit beta levels [37], glycoprotein acetyls levels [46], heme oxygenase 1 levels [62], interleukin-16 levels [63], kininostatin levels [53], lipoprotein lipase levels [62], metabolite peak levels (QI7471) [64], metabolite peak levels (QI7638) [64], metabolite peak levels (QI8556) [64], metabolite peak levels (QI8596) [64], parathyroid hormone-related protein levels [58], plasma serine protease inhibitor levels [58], serine/threonine-protein kinase ULK3 levels [53], serum levels of protein ADM [65], serum levels of protein ATP13A1 [65], serum levels of protein CAPN1;CAPNS1 [65], serum levels of protein CDH15 [65], serum levels of protein DCK [65], serum levels of protein DPEP2 [65], serum levels of protein ETHE1 [65], serum levels of protein GPNMB [65], serum levels of protein ITIH4 [65], serum levels of protein LAMTOR3 [65], serum levels of protein LGMN [65], serum levels of protein PDCL2 [65], serum levels of protein PLXDC1 [65], serum levels of protein SIRT5 [65], serum levels of protein TMEM87B [65], serum lipopolysaccharide activity [66], serum metabolite levels [67], superoxide dismutase mitochondrial levels [37], and taurine levels [59].

4. rs35610898 (*SLC34A1*; EHG: p=3.21e-15; s=0.025), an intron variant associated with blood urea nitrogen levels [97], estimated glomerular filtration rate [127], hematocrit [91], and serum creatinine levels [97].

5. rs2731672 (*GRK6*; ALL: p=4.82e-11; s=-0.009), an intron variant associated with phenylalanine levels [129,130], activated partial thromboplastin time [131], circulating chromogranin peptide levels [132], circulating vasoactive peptide levels [133], high-sensitivity cardiac troponin I concentration [134], interleukin-2 levels [58], matrix extracellular phosphoglycoprotein levels [62], metabolite levels [135], plexin domain-

containing protein 1 levels [53], serum levels of protein DBNL [65], serum levels of protein HBEGF [65], serum levels of protein HTN3 [65], superoxide dismutase mitochondrial level in Chronic kidney disease with hypertension and no diabetes [61], x-11792 levels [136], and x-12038-to-bradykinin, des-arg 9 ratio [137].

#### Peak 10: HLA

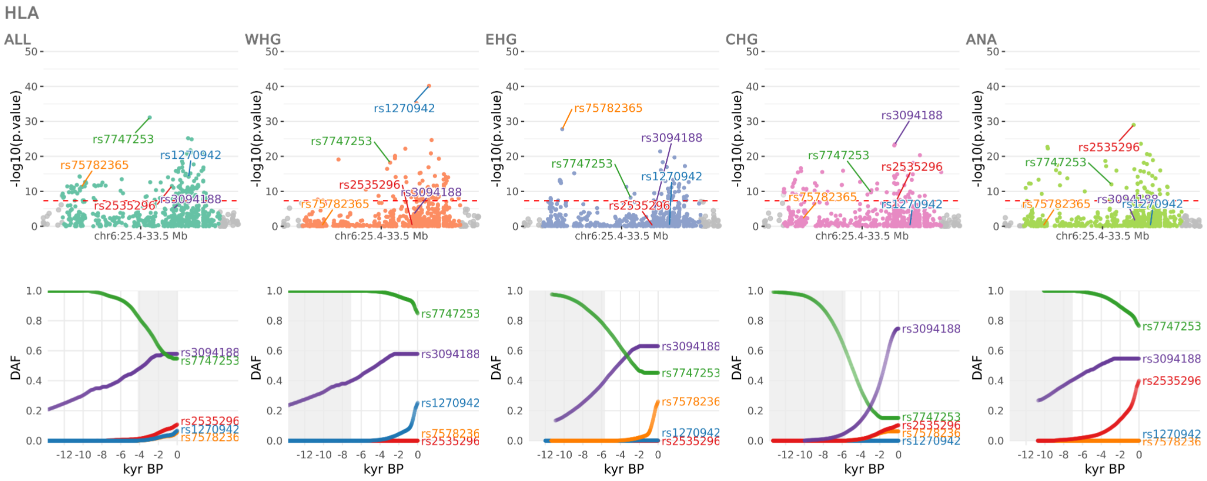

**Figure S65.** Selection at the *HLA* locus, spanning chr6:25417423-33524820. Results for the pan-ancestry analysis (ALL) plus each of the four marginal ancestries: Western hunter-gatherers (WHG), Eastern hunter-gatherers (EHG), Caucasus hunter-gatherers (CHG) and Anatolian farmers (ANA). Row one shows zoomed Manhattan plots of the p-values for each ancestry, and row two shows allele trajectories for the top SNPs across all ancestries.

The tenth peak spanned the region chr6:25417423-33524820, with the five most significant SNPs (across ancestries) and their associations in the GWAS Catalog (r2023-04-07) being:

1. rs1270942 (*CFB*; WHG: p=6.82e-41; s=0.036), a non-coding transcript exon variant associated with hemoglobin concentration [138], iDP dMRI TBSS ICVF Retrolenticular part of internal capsule R [116], inguinal hernia [139], systemic lupus erythematosus [140], and type 1 diabetes and autoimmune thyroid diseases [141].

2. rs7747253 (*HLA-W*; ALL: p=7.56e-32; s=-0.018), an upstream gene variant/intergenic variant associated with heel bone mineral density [70].

3. rs2535296 (*RNU6-1133P*; ANA: p=1.03e-29; s=0.024), a regulatory region variant/downstream gene variant/intergenic variant associated with pneumonia [142].

4. rs75782365 (*BTN3A1*; EHG: p=1.67e-28; s=0.049), an intron variant associated with schizophrenia (MTAG) [143,144], a body shape index [71], autism spectrum disorder or schizophrenia [145], bipolar disorder (MTAG) [143], depression (broad) [146], and help-seeking from a GP [147].

5. rs3094188 (*POU5F1 / PSORS1C3*; CHG: p=3.64e-24; s=0.023), an intron variant associated with drug-induced Stevens-Johnson syndrome or toxic epidermal necrolysis [148].

#### Peak 11: GATA4

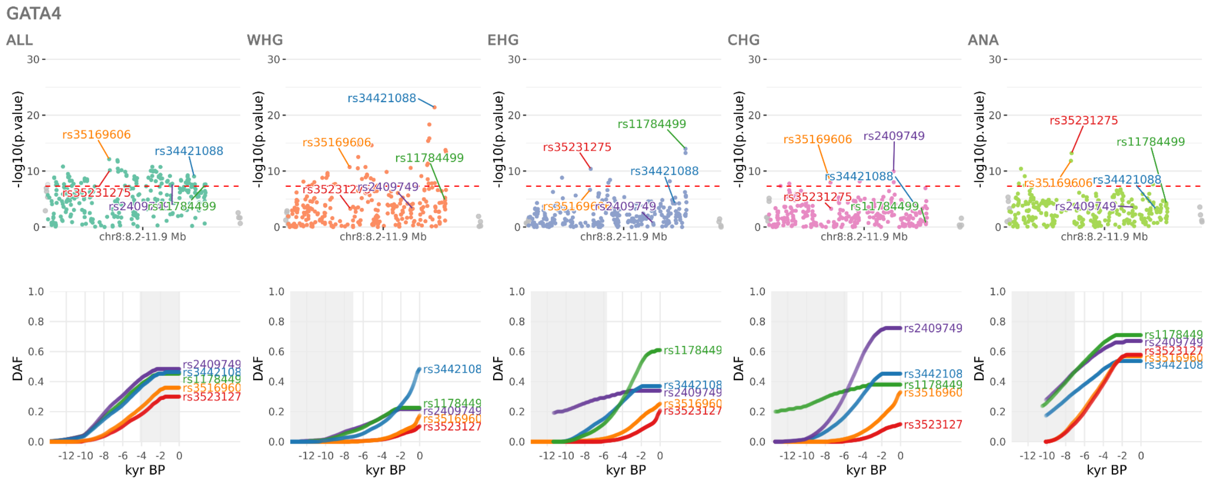


**Figure S66.** Selection at the *GATA4* locus, spanning chr8:8168474-11856864. Results for the pan-ancestry analysis (ALL) plus each of the four marginal ancestries: Western hunter-gatherers (WHG), Eastern hunter-gatherers (EHG), Caucasus hunter-gatherers (CHG) and Anatolian farmers (ANA). Row one shows zoomed Manhattan plots of the p-values for each ancestry, and row two shows allele trajectories for the top SNPs across all ancestries.

The eleventh peak spanned the region chr8:8168474-11856864, with the five most significant SNPs (across ancestries) and their associations in the GWAS Catalog (r2023-04-07) being:

1. rs34421088 (*GATA4*; WHG: p=3.92e-22; s=0.021), an intron variant associated with average diameter for VLDL particles [46], body mass index [97], general factor of neuroticism [149], height [97], neuroticism [150], and triglyceride levels in large VLDL [46].

2. rs11784499 (*DEFB136*; EHG: p=1.00e-14; s=0.018), an upstream gene variant associated with medication use (thyroid preparations) [151].

3. rs35231275 (*TNKS*; ANA: p=6.29e-14; s=0.014), an intron variant associated with pulse pressure x alcohol consumption interaction (2df test) [152].

4. rs35169606 (*TNKS*; ALL: p=7.29e-13; s=0.014), an intron variant associated with a body shape index [71], lifetime smoking index [72], waist circumference adjusted for body mass index [71], waist-hip index [71], and waist-to-hip ratio adjusted for BMI [71].

5. rs2409749 (*AF131215.8*; CHG: p=1.11e-10; s=0.016), an intergenic variant associated with neuroticism [150].

#### Peak 12: CTD-2008O4.1

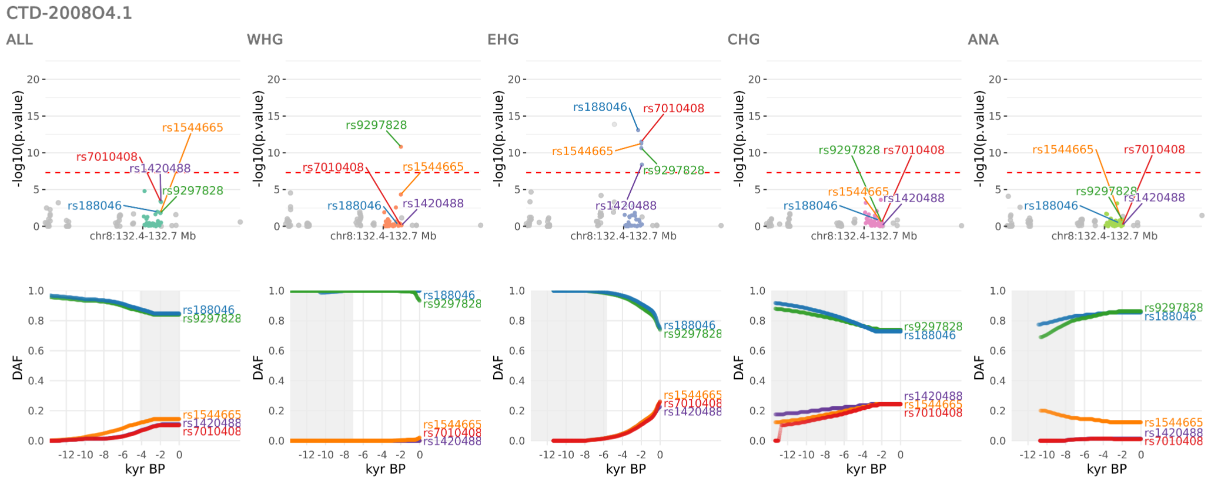


**Figure S67.** Selection at the *CTD-2008O4.1* locus, spanning chr8:132393015-132669706. Results for the pan-ancestry analysis (ALL) plus each of the four marginal ancestries: Western hunter-gatherers (WHG), Eastern hunter-gatherers (EHG), Caucasus hunter-gatherers (CHG) and Anatolian farmers (ANA). Row one shows zoomed Manhattan plots of the p-values for each ancestry, and row two shows allele trajectories for the top SNPs across all ancestries.

The twelfth peak spanned the region chr8:132393015-132669706, with the five most significant SNPs (across ancestries) and their associations in the GWAS Catalog (r2023-04-07) being:

1. rs188046 (*CTD-2008O4.1*; EHG: p=8.26e-14; s=-0.028), an intergenic variant with no associations in the GWAS Catalog.

2. rs9297828 (*CTD-2008O4.1*; WHG: p=1.52e-11; s=-0.100), an intergenic variant with no associations in the GWAS Catalog.
3. rs7010408 (*CTD-2008O4.1*; EHG: p=3.01e-12; s=0.026), an intergenic variant with no associations in the GWAS Catalog.
4. rs1544665 (*CTD-2008O4.1*; EHG: p=5.49e-12; s=0.025), an intergenic variant with no associations in the GWAS Catalog.
5. rs1420488 (*CTD-2008O4.1*; EHG: p=4.04e-09; s=0.025), an intergenic variant with no associations in the GWAS Catalog.

#### Peak 13: ABO

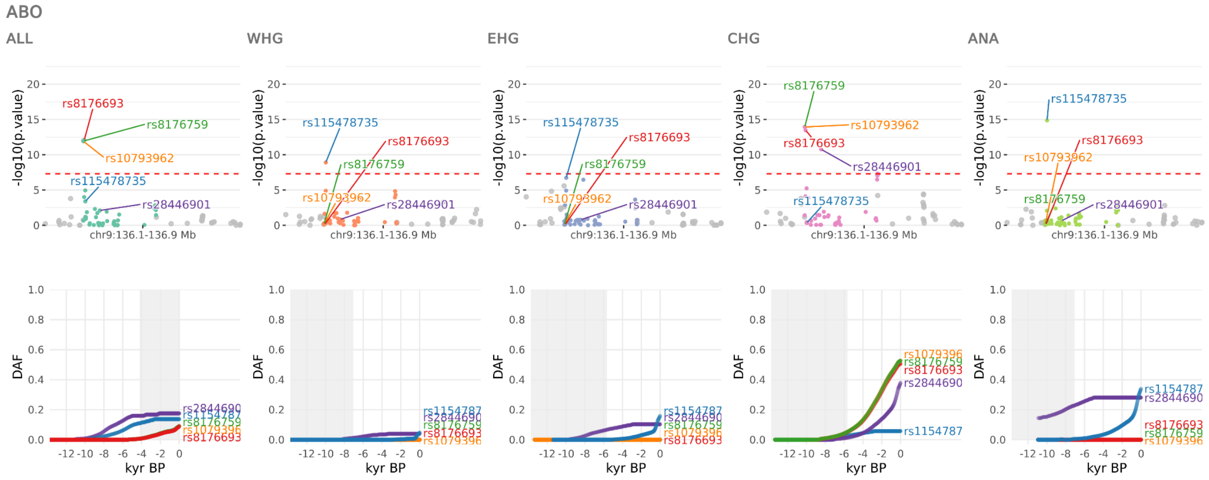


**Figure S68.** Selection at the ABO locus, spanning chr9:136128000-136930134. Results for the pan-ancestry analysis (ALL) plus each of the four marginal ancestries: Western hunter-gatherers (WHG), Eastern hunter-gatherers (EHG), Caucasus hunter-gatherers (CHG) and Anatolian farmers (ANA). Row one shows zoomed Manhattan plots of the p-values for each ancestry, and row two shows allele trajectories for the top SNPs across all ancestries.

The thirteenth peak spanned the region chr9:136128000-136930134, with the five most significant SNPs (across ancestries) and their associations in the GWAS Catalog (r2023-04-07) being:

1. rs115478735 (*ABO*; ANA: p=1.34e-15; s=0.027), an intron variant associated with apolipoprotein B levels [46,94], IDL cholesterol levels [46,94], linoleic acid levels [46,153], angiotensin-converting enzyme levels [53], cholesterol levels in IDL [46], cholesterol

levels in large LDL [46], cholesterol levels in medium LDL [46], cholesterol levels in medium VLDL [46], cholesterol levels in small LDL [46], cholesterol levels in small VLDL [46], cholesterol levels in very small VLDL [46], cholesterol to total lipids ratio in chylomicrons and extremely large VLDL [46], cholesteryl ester levels in IDL [46], cholesteryl ester levels in large LDL [46], cholesteryl ester levels in LDL [46], cholesteryl ester levels in medium LDL [46], cholesteryl ester levels in medium VLDL [46], cholesteryl ester levels in small LDL [46], cholesteryl ester levels in small VLDL [46], cholesteryl ester levels in very small VLDL [46], cholesteryl ester levels in VLDL [46], cholesteryl esters to total lipids ratio in chylomicrons and extremely large VLDL [46], cholesteryl esters to total lipids ratio in small VLDL [46], clinical LDL cholesterol levels [46], concentration of IDL particles [46], concentration of large LDL particles [46], concentration of LDL particles [46], concentration of medium LDL particles [46], concentration of small LDL particles [46], concentration of very small VLDL particles [46], free cholesterol levels in large LDL [46], free cholesterol levels in LDL [46], free cholesterol levels in medium LDL [46], free cholesterol levels in medium VLDL [46], free cholesterol levels in small LDL [46], free cholesterol levels in small VLDL [46], free cholesterol levels in very small VLDL [46], free cholesterol to total lipids ratio in chylomicrons and extremely large VLDL [46], galectin-8 levels [53], HDL cholesterol [97], immunoglobulin superfamily containing leucine-rich repeat protein 2 levels (ISLR2.13124.20.3) [37], low-density lipoprotein receptor-related protein 8 levels [53], medication use for hyperlipidemia [54], omega-6 fatty acid levels [46], phospholipid levels in IDL [46], phospholipid levels in large LDL [46], phospholipid levels in LDL [46], phospholipid levels in medium LDL [46], phospholipid levels in medium VLDL [46], phospholipid levels in small LDL [46], phospholipid levels in very small VLDL [46], phospholipids to total lipids ratio in small HDL [46], platelet endothelial aggregation receptor 1 levels [53], polyunsaturated fatty acid levels [46], protein-tyrosine sulfotransferase 2 levels [37], ratio of apolipoprotein B to apolipoprotein A1 levels [46], ratio of linoleic acid to total fatty acids [46], remnant cholesterol (non-HDL, non-LDL - cholesterol) [46], total cholesterol levels [46], total cholesterol minus HDL-C levels [46], total esterified cholesterol levels [46], total free cholesterol levels [46], total lipid levels in IDL [46], total lipid levels in large LDL [46], total lipid levels in LDL [46], total lipid levels in medium LDL [46], total lipid levels in small LDL [46], total lipid levels in very small VLDL [46], total omega-6 fatty acid levels [153], triglycerides to total lipids ratio in chylomicrons and extremely large VLDL [46], triglycerides to total lipids ratio in large VLDL [46], and vLDL cholesterol levels [46].

2. rs8176759 (*ABO / RP11-430N14.4*; CHG: p=1.05e-14; s=0.020), a non-coding transcript exon variant/3-prime UTR variant associated with granulocyte percentage of myeloid white cells [83], mean corpuscular hemoglobin [91], platelet endothelial cell adhesion molecule levels [62], plateletcrit [83], red cell distribution width [91], and von Willebrand factor levels [58].

3. rs8176693 (*ABO*; CHG: p=3.29e-14; s=0.020), an intron variant associated with cadherin-5 levels [53,58], tyrosine-protein kinase receptor Tie-1, soluble levels [58,68], angiopoietin-1 receptor, soluble levels [37], basal Cell Adhesion Molecule levels [53], blood protein levels in cardiovascular risk [154], cadherin-17 level in Chronic kidney disease with hypertension and no diabetes [61], endothelial growth factor levels [155], high serum lipase activity [156], klotho levels [53], mean corpuscular hemoglobin concentration [88], neurogenic locus notch homolog protein 1 levels [68], platelet glycoprotein 4 levels (CD36.2973.15.2) [37], semaphorin-6B levels [58], serum albumin levels [97], serum lipase activity [156], sex hormone-binding globulin levels [157], and sex hormone-binding globulin levels adjusted for BMI [157].

4. rs10793962 (*ABO / RP11-430N14.4*; CHG: p=1.16e-14; s=0.020), a non-coding transcript exon variant/3-prime UTR variant associated with intraocular pressure [158,159], adhesion G protein-coupled receptor F5 levels [53], angiopoietin-1 receptor, soluble levels [68], cadherin-5 levels [62], cD109 antigen levels [68], immunoglobulin superfamily containing leucine-rich repeat protein 2 levels (ISLR2.13124.20.3) [37], interleukin-27 receptor subunit alpha levels [58], ischemic stroke or von Willebrand factor levels (pleiotropy) [80], mean corpuscular hemoglobin concentration [97], mean corpuscular volume [97], platelet endothelial aggregation receptor 1 levels [53], and red cell distribution width [91].

5. rs28446901 (*ADAMTS13*; CHG: p=1.68e-11; s=0.024), an intron variant associated with tyrosine-protein kinase receptor Tie-1, soluble levels [68] and venous thromboembolism [160].

#### Peak 14: FADS2

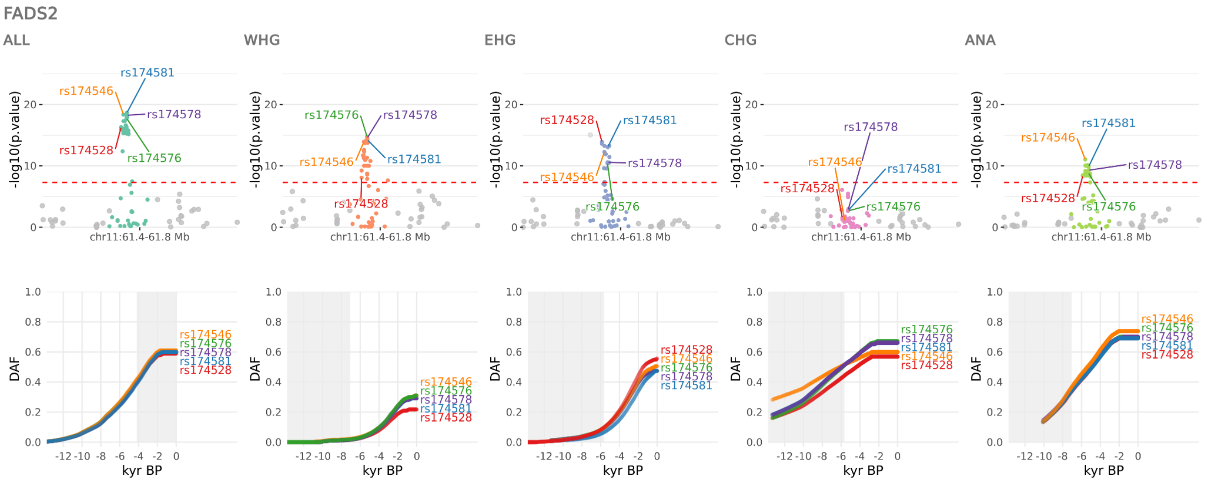


**Figure S69.** Selection at the *FADS2* locus, spanning chr11:61416970-61830169. Results for the pan-ancestry analysis (ALL) plus each of the four marginal ancestries: Western hunter-gatherers (WHG), Eastern hunter-gatherers (EHG), Caucasus hunter-gatherers (CHG) and Anatolian farmers (ANA). Row one shows zoomed Manhattan plots of the p-values for each ancestry, and row two shows allele trajectories for the top SNPs across all ancestries.

The fourteenth peak spanned the region chr11:61416970-61830169, with the five most significant SNPs (across ancestries) and their associations in the GWAS Catalog (r2023-04-07) being:

1. rs174581 (*FADS2*; ALL: p=2.21e-19; s=0.014) an intron variant associated with 1-margaroyl-2-linoleoyl-GPC levels [42], 1-oleoyl-2-eicosapentaenoyl-GPC levels [42], 1-palmitoyl-2-linoleoyl-gpc levels [42], 1-pentadecanoyl-2-linoleoyl-GPC levels [42], 1-stearoyl-2-linoleoyl-gpc levels [42], arachidonoylcholine levels [42], cholesteryl ester levels [73], cholesteryl ester levels [74], diacylglycerol levels [74], fatty acid levels [74], lysophosphatidylcholine levels [74], male-pattern baldness [75], phosphatidate levels [74], phosphatidylcholine levels [73], phosphatidylethanolamine levels [74], phosphatidylglycerol levels [74], phosphatidylinositol levels [74], phosphatidylserine levels [74], phospholipids to total lipids ratio in small VLDL [46], serum metabolite ratios in chronic kidney disease [76], sphingomyelin levels [74], and triacylglycerol levels [74].

2. rs174576 (*FADS2*; ALL: p=1.70e-18; s=0.012) an intron variant associated with 1-lignoceroyl-GPC levels [161], 1-stearoyl-2-docosapentaenoyl-GPC levels [161], 2-linoleoyl-GPE levels [161], alanine aminotransferase levels [162], alkenylphosphatidylcholine levels

[73], alkenylphosphatidylethanolamine levels [73], bipolar disorder [163], cholesterol levels in large HDL [46], cholesterol to total lipids ratio in large LDL [46], cholesteryl ester levels in large HDL [46], cholesteryl ester levels [74], diacylglycerol levels [74], fatty acid levels [74], free cholesterol to total lipids ratio in large VLDL [46], lysoPhosphatidylcholine acyl C18:0 levels [129], lysophosphatidylcholine levels [74], metabolite levels [164], phosphatidate levels [74], phosphatidylcholine levels [74,129,136,165], phosphatidylethanolamine levels [74], phosphatidylglycerol levels [74], phosphatidylinositol levels [74], phosphatidylserine levels [74], phospholipid levels in large HDL [46], phospholipids to total lipids ratio in chylomicrons and extremely large VLDL [46], platelet count [91], serum metabolite levels (CMS) [166], serum metabolite ratios in chronic kidney disease [76], sphingomyelin levels [74], total lipid levels in large HDL [46], trans fatty acid levels [167], and triacylglycerol levels [74].

3. rs174528 (*MYRF / TMEM258*; ALL: p=4.90e-17; s=0.013) an intron variant associated with 1-arachidonylglycerol levels [42], circulating docosahexaenoic acid levels [153], coronary artery disease or von Willebrand factor levels (pleiotropy) [80], degree of unsaturation [46], docosahexaenoic acid levels [46], venous thromboembolism or von Willebrand factor levels [80], fasting apolipoprotein A-I [168], fasting total cholesterol [168], gondoic acid levels [169], hemoglobin concentration [91], lysophosphatidylcholine levels [73], omega-3 fatty acid levels [46], phosphatidylcholine diacyl levels [129], phosphatidylcholine-ether levels [170], phosphatidylinositol levels [73], plasma omega-6 polyunsaturated fatty acid levels (arachidonic acid) [171], postprandial cholesterol esters in very large HDL [168], postprandial total cholesterol [168], ratio of docosahexaenoic acid to total fatty acid levels [46], ratio of linoleic acid to total fatty acids [46], ratio of monounsaturated fatty acids to total fatty acids [46], ratio of omega-6 fatty acids to omega-3 fatty acids [46], ratio of omega-6 fatty acids to total fatty acids [46], serum metabolite ratios in chronic kidney disease [76], sex hormone-binding globulin levels [157], stem cell factor levels [172], trans fatty acid levels [167], and vaccenic acid levels [169].

4. rs174546 (*FADS1 / FADS2*; ALL: p=4.41e-19; s=0.013) an 3-prime UTR variant associated with 1-linoleoyl-2-linolenoyl-GPC levels [43], 1-linoleoyl-gpc levels [43], 1-linoleoyl-GPE levels [43], 1-linoleoyl-GPI levels [161], 1-linoleoylglycerophosphoethanolamine levels [136], 1-oleoyl-2-linoleoyl-GPE levels [43], 1-palmitoyl-2-dihomo-linolenoyl-GPC levels [43], 1-palmitoyl-2-docosahexaenoyl-gpc levels [43], 1-stearoyl-2-docosahexaenoyl-gpc levels [43], 1-stearoyl-2-linoleoyl-gpc levels

[43], 1,2-dilinoleoyl-GPC levels [43], adrenate levels [43], c-reactive protein levels [173], change in serum metabolite levels [166], cholesterol total [174,175], cholesteryl ester levels [74], diacylglycerol levels [74], eicosapentaenoate levels [43], fatty acid levels [74], HDL cholesterol levels [174–176], high density lipoprotein cholesterol levels [177], iron status biomarkers (total iron binding capacity) [178], IDL cholesterol [174,175], IDL cholesterol levels [176], low density lipoprotein cholesterol levels [50,177,179], lysophosphatidylcholine levels [74], lysophosphatidylethanolamine levels [74], phosphatidate levels [74], phosphatidylcholine levels [74,129,136,165], phosphatidylethanolamine levels [73,74], phosphatidylglycerol levels [74], phosphatidylinositol levels [74], phosphatidylserine levels [74], plasma omega-6 polyunsaturated fatty acid levels (gamma-linolenic acid) [171], platelet count [97], qT interval [180], serum metabolite levels [166], sphingomyelin levels [74], total cholesterol levels [177], trans fatty acid levels [167], triacylglycerol levels [74], triglyceride levels [176,179], and triglycerides [174,175,177].

5. rs174578 (*FADS2*; ALL: p=5.42e-19; s=0.014) an intron variant associated with 1-arachidonoylglycerophosphoethanolamine levels [137], 1-arachidonoylglycerophosphoinositol levels [137], apolipoprotein A1 levels [46], cholesterol levels in medium HDL [46], cholesteryl ester levels [46,74], concentration of HDL particles [46], diacylglycerol levels [74], fatty acid levels [74], HDL cholesterol levels [46], hematocrit [91], hemoglobin concentration [91], lysophosphatidylcholine levels [74], phosphatidate levels [74], phosphatidylcholine levels [73,74,129,165], phosphatidylethanolamine levels [74], phosphatidylglycerol levels [74], phosphatidylinositol levels [74], phosphatidylserine levels [74], phospholipids to total lipids ratio in very small VLDL [46], plasma omega-6 polyunsaturated fatty acid levels (linoleic acid) [171], serum metabolite ratios in chronic kidney disease [76], sphingomyelin levels [74], total concentration of lipoprotein particles [46], trans fatty acid levels [167], and triacylglycerol levels [74].

#### Peak 15: ACAD10

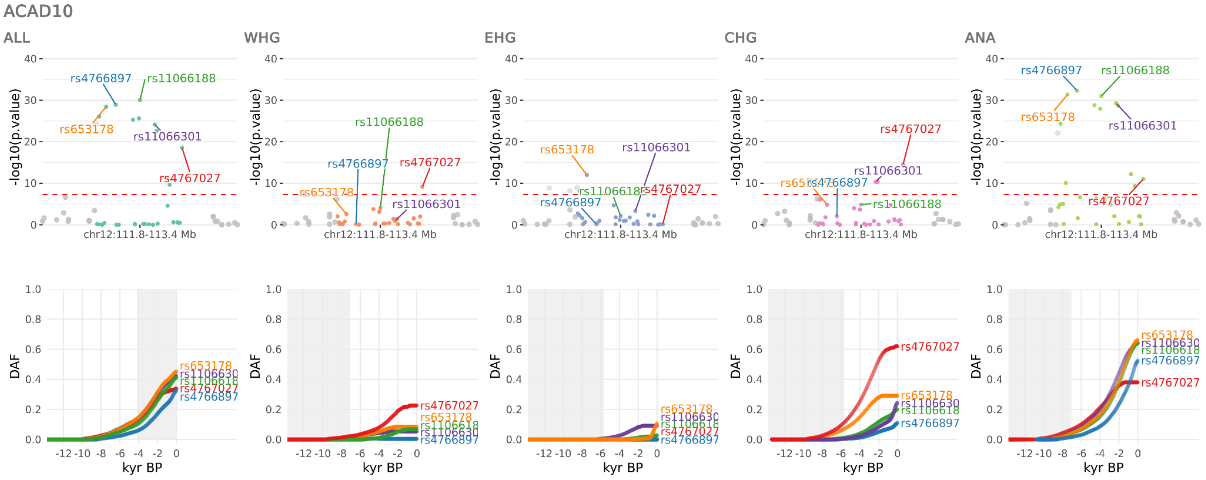


**Figure S70.** Selection at the *ACAD10* locus, spanning chr12:111833788-113359157. Results for the pan-ancestry analysis (ALL) plus each of the four marginal ancestries: Western hunter-gatherers (WHG), Eastern hunter-gatherers (EHG), Caucasus hunter-gatherers (CHG) and Anatolian farmers (ANA). Row one shows zoomed Manhattan plots of the p-values for each ancestry, and row two shows allele trajectories for the top SNPs across all ancestries.

The fifteenth peak spanned the region chr12:111833788-113359157, with the five most significant SNPs (across ancestries) and their associations in the GWAS Catalog (r2023-04-07) being:

1. rs4766897 (*ACAD10*; ANA: p=4.95e-33; s=0.022), an intron variant associated with cystatin C plasma levels [181], fibrinogen levels [182], and hemoglobin concentration [138].

2. rs11066188 (*HECTD4*; ANA: p=9.89e-32; s=0.021), an intron variant associated with body mass index (MTAG) [50], celiac disease and Rheumatoid arthritis [77], diastolic blood pressure x depressive symptoms interaction (2df test) [78], heart failure [79], ischemic stroke or factor VII levels (pleiotropy) [80], ischemic stroke or factor VIII levels (pleiotropy) [80], ischemic stroke or factor XI levels (pleiotropy) [80], ischemic stroke or fibrinogen levels (pleiotropy) [80], ischemic stroke or plasminogen activator inhibitor 1 levels (pleiotropy) [80], ischemic stroke or tissue plasminogen activator levels (pleiotropy) [80], ischemic stroke or von Willebrand factor levels (pleiotropy) [80], and spleen volume [81].

3. rs4767027 (*OAS1 / RP1-71H24.1*; ALL: p=2.75e-19; s=0.018), an intron variant associated with 2'-5'-oligoadenylate synthase 1 levels (OAS1.10361.25.3) [37].

4. rs653178 (*ATXN2*; ANA: p=4.23e-32; s=0.021), an intron variant associated with alanine aminotransferase levels [183], allergic disease (asthma, apolipoprotein A1 levels [183], apolipoprotein B levels [183], aspartate aminotransferase levels [183], aspartate aminotransferase to alanine aminotransferase ratio [183], asthma [184], asthma or COVID-19 (pleiotropy) [185], beta-2-microglobulin levels [53], blood pressure [186], brain morphology (MOSTest) [187], brain shape (segment 1) [188], c-X-C motif chemokine 10 levels [53], c-X-C motif chemokine 16 levels [53], cD45RA- CD4+ T cell Absolute Count [189], celiac disease [190], celiac disease or Rheumatoid arthritis [191], cholesterol levels [46], chronic kidney disease [192], colorectal cancer [87], concentration of IDL particles [46], concentration of large HDL particles [46], concentration of very large HDL particles [46], cortical surface area [193], cortical thickness [193], cystatin C levels [181,183], diastolic blood pressure [194–196], diastolic blood pressure x depressive symptoms interaction (2df test) [78], direct bilirubin levels [183], eczema [89], eosinophil counts [83], free cholesterol levels [46], glycated hemoglobin levels [183], hay fever and/or eczema [197], hay fever or eczema) [197], hemoglobin concentration [138], high density lipoprotein cholesterol levels [183], inflammatory bowel disease [126,198], kynurenine levels [129], IDL cholesterol [199], lifetime smoking [200], low density lipoprotein cholesterol levels [183,201], lymphocyte activation gene 3 protein levels [53], lymphocyte count [91], lymphocyte percentage of white cells [88], mean arterial pressure [202], monocyte count [83], myocardial infarction [203], neutrophil percentage of granulocytes [83], neutrophil percentage of white cells [88], neutrophil-to-lymphocyte ratio [111], non-albumin protein levels [183], non-HDL cholesterol levels [201], phospholipid levels [46], procollagen C-endopeptidase enhancer 2 levels [53], psoriasis or type 2 diabetes [204], remnant cholesterol [46], right ventricular stroke volume [205], sarcoidosis [206], serum albumin levels [183], serum alkaline phosphatase levels [183], serum phosphate levels [183], serum total protein level [183], sphingomyelin levels [46], sum eosinophil basophil counts [83], systemic lupus erythematosus [207], thyroid peroxidase antibody positivity [208], tonsillectomy [142], total bilirubin levels [183], total cholesterol levels [46,183,199,201], total cholines levels [46], total esterified cholesterol levels [46], total free cholesterol levels [46], total lipid levels in IDL [46], total phospholipid levels in lipoprotein particles [46], triglyceride levels in non-type 2 diabetes [209], triglycerides to total lipids ratio [46], type 1 diabetes [210], urate levels [183,211,212], vertex-wise cortical surface area [213],

vertex-wise cortical thickness [213], vertex-wise sulcal depth [213], and whole brain free water diffusion [214].

5. rs11066301 (*PTPN11*; ANA: p=4.84e-30; s=0.020), an intron variant associated with alanine aminotransferase levels [183], apolipoprotein A1 levels [183], apolipoprotein B levels [183], aspartate aminotransferase levels [183], cystatin C levels [183], cystatin C plasma levels [181], direct bilirubin levels [183], eosinophil counts [91], glycated hemoglobin levels [183], hematological parameters [215], hemoglobin concentration [138], high density lipoprotein cholesterol levels [183], low density lipoprotein cholesterol levels [183], mean arterial pressure [202], medication use (beta blocking agents) [151], non-albumin protein levels [183], peripheral artery disease [216,217], serum albumin levels [183], serum alkaline phosphatase levels [183], serum phosphate levels [183], serum total protein level [183], systemic seropositive rheumatic diseases [218], total bilirubin levels [183], total cholesterol levels [183], and urate levels [183].

#### Peak 16: CYP1A1

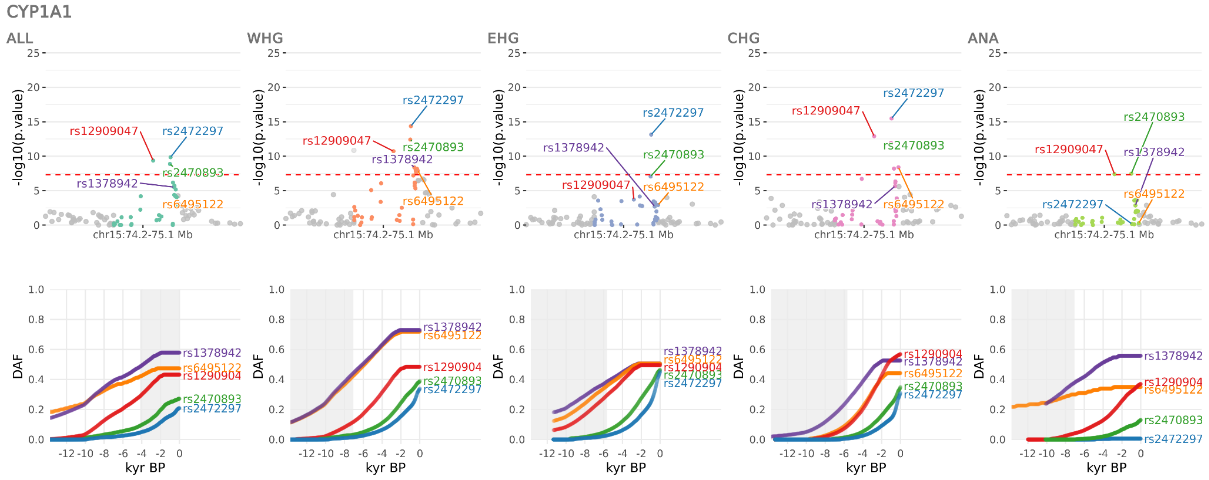


**Figure S71.** Selection at the *CYP1A1* locus, spanning chr15:74223118-75125645. Results for the pan-ancestry analysis (ALL) plus each of the four marginal ancestries: Western hunter-gatherers (WHG), Eastern hunter-gatherers (EHG), Caucasus hunter-gatherers (CHG) and Anatolian farmers (ANA). Row one shows zoomed Manhattan plots of the p-values for each ancestry, and row two shows allele trajectories for the top SNPs across all ancestries.

The sixteenth peak spanned the region chr15:74223118-75125645, with the five most significant SNPs (across ancestries) and their associations in the GWAS Catalog (r2023-04-07) being:

1. rs2472297 (*CYP1A1*; CHG: p=3.33e-16; s=0.028), an intergenic variant associated with coffee consumption (cups per day) [105,219,220], estimated glomerular filtration rate [127,183,221], alcohol consumption (drinks per week) (MTAG) [222,223], coffee consumption [224,225], estimated glomerular filtration rate (creatinine) [226,227], plasma clozapine levels in treatment-resistant schizophrenia [228,229], tea consumption [225,230], urate levels [183,231], urinary albumin-to-creatinine ratio [232,233], urinary sodium excretion [183,234], 1-methylxanthine levels [43], 1,3,7-trimethylurate levels [42], 1,7-dimethylurate levels [42], 5-acetylamino-6-amino-3-methyluracil levels [43], 5-hydroxy-2-methylpyridine sulfate levels [43], alcohol consumption (drinks per week) [222], bitter beverage consumption [225], bitter non-alcoholic beverage consumption [225], body mass index [235], caffeine consumption from coffee [236], caffeine consumption from coffee or tea [236], caffeine consumption from tea [236], caffeine levels [42], caffeine metabolism (plasma 1,3,7-trimethylxanthine (caffeine) level) [237], caffeine metabolism (plasma 1,7-dimethylxanthine (paraxanthine) to 1,3,7-trimethylxanthine (caffeine) ratio) [237], clozapine concentration in schizophrenia [238], coffee max liking [51], creatinine levels [183], cystatin C levels [183], estimated glomerular filtration rate in non-diabetics [127], f-chocolate/coffee liking (derived food-liking factor) [51], f-coffee/alcohol liking (derived food-liking factor) [51], milk chocolate liking [51], milky sweets liking [51], paraxanthine levels [42], plasma norclozapine levels in treatment-resistant schizophrenia [229], predicted visceral adipose tissue [239], relative carbohydrate intake [240], serum phosphate levels [183], theophylline levels [42], urea levels [183], urinary albumin excretion [241], urinary creatinine levels [183], urinary potassium excretion [234], and x-21286 levels [42].

2. rs2470893 (*CYP1A1*; WHG: p=3.86e-13; s=0.020), an upstream gene variant associated with coffee consumption [219,242], blood urea nitrogen levels [221], body mass index (MTAG) [50], caffeine consumption [243], caffeine metabolism (plasma 1,3,7-trimethylxanthine (caffeine) level) [237], caffeine metabolism (plasma 1,7-dimethylxanthine (paraxanthine) to 1,3,7-trimethylxanthine (caffeine) ratio) [237], microalbuminuria [244], platelet distribution width [83], skin autofluorescence [245], urinary albumin excretion (no hypertensive medication) [241], and urinary albumin-to-creatinine ratio [244].

3. rs12909047 (*RP11-10O17.1 / RP11-10O17.3*; CHG: p=1.25e-13; s=0.019), an intron variant associated with caffeine metabolism (plasma 1,3-dimethylxanthine (theophylline) level) [237] and caffeine metabolism (plasma 1,3,7-trimethylxanthine (caffeine) level) [237].

4. rs6495122 (*CPLX3*; CHG: p=4.13e-09; s=0.016), a downstream gene variant associated with diastolic blood pressure [82,246], total cholesterol levels [97,201], coffee consumption [242], diastolic blood pressure (cigarette smoking interaction) [247], mean arterial pressure [202], non-HDL cholesterol levels [201], systolic blood pressure [82], and systolic blood pressure (cigarette smoking interaction) [247].

5. rs1378942 (*CSK*; WHG: p=4.35e-09; s=0.010), an intron variant associated with diastolic blood pressure [194,248–250], systolic blood pressure [248,250,251], blood pressure [186], coronary artery disease and total cholesterol levels (multivariate analysis) [252], diastolic blood pressure (cigarette smoking interaction) [247], systemic sclerosis [253], and systolic blood pressure (cigarette smoking interaction) [247].

#### Peak 17: WWP2

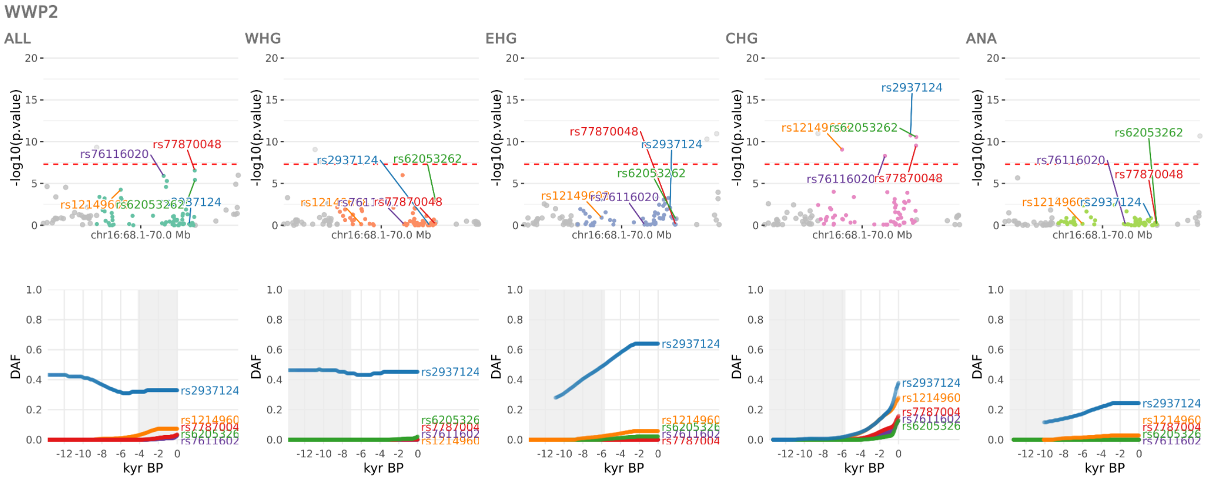


**Figure S72.** Selection at the *WWP2* locus, spanning chr16:68138091-69969299. Results for the pan-ancestry analysis (ALL) plus each of the four marginal ancestries: Western hunter-gatherers (WHG), Eastern hunter-gatherers (EHG), Caucasus hunter-gatherers (CHG) and Anatolian farmers (ANA). Row one shows zoomed Manhattan plots of the p-values for each ancestry, and row two shows allele trajectories for the top SNPs across all ancestries.

The seventeenth peak spanned the region chr16:68138091-69969299, with the five most significant SNPs (across ancestries) and their associations in the GWAS Catalog (r2023-04-07) being:

1. rs2937124 (*WWP2*; CHG: p=1.76e-11; s=0.025), an intron variant associated with triglycerides [199].

2. rs62053262 (*WWP2*; CHG: p=2.78e-11; s=0.057), an intron variant associated with ascending aorta maximum area [254,255], ascending aorta minimum area [254,255], ascending aorta diameter [256], ascending aorta maximum area (MTAG) [255], ascending thoracic aortic diameter [257], descending thoracic aortic diameter [257], and pulse pressure [251].

3. rs77870048 (*WWP2*; CHG: p=2.99e-10; s=0.034), an intron variant associated with myocardial infarction [97,258], systolic blood pressure [89,97], alanine aminotransferase levels [183], ascending aorta distensibility (MTAG) [255], ascending aorta minimum area [255], descending aorta maximum area [255], descending aorta maximum area (MTAG) [255], descending aorta minimum area [255], descending thoracic aortic diameter [257], diastolic blood pressure [251], and pulse pressure [97].

4. rs12149608 (*ZFP90*; CHG: p=8.74e-10; s=0.024), an intron variant associated with ulcerative colitis [126].

5. rs76116020 (*RP11-343C2.9 / TMED6*; CHG: p=5.04e-09; s=0.040), a missense variant associated with high density lipoprotein cholesterol levels [183,259], total cholesterol levels [183,201], apolipoprotein A1 levels [183], cholesteryl esters to total lipids ratio in large LDL [46], omega-6 fatty acid levels [46], polyunsaturated fatty acid levels [46], total lipid levels in lipoprotein particles [46], and total omega-6 fatty acid levels [153].

#### Peak 18: RAPGEFL1

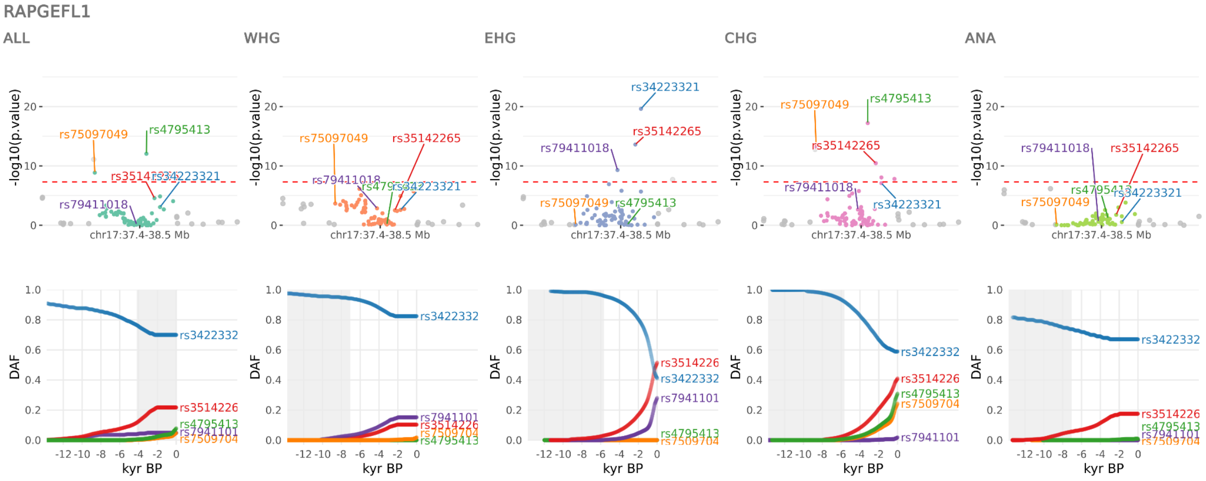


**Figure S73.** Selection at the *RAPGEFL1* locus, spanning chr17:37372773-38545193. Results for the pan-ancestry analysis (ALL) plus each of the four marginal ancestries: Western hunter-gatherers (WHG), Eastern hunter-gatherers (EHG), Caucasus hunter-gatherers (CHG) and Anatolian farmers (ANA). Row one shows zoomed Manhattan plots of the p-values for each ancestry, and row two shows allele trajectories for the top SNPs across all ancestries.

The eighteenth peak spanned the region chr17:37372773-38545193, with the five most significant SNPs (across ancestries) and their associations in the GWAS Catalog (r2023-04-07) being:

1. rs34223321 (*RAPGEFL1*; EHG: p=2.45e-20; s=-0.023), a downstream gene variant associated with white blood cell count [88,89].

2. rs4795413 (*PSMD3*; CHG: p=6.17e-18; s=0.028), a non-coding transcript exon variant associated with atopic asthma [109].

3. rs35142265 (*MSL1*; EHG: p=2.50e-14; s=0.023), an TF binding site variant associated with asthma (childhood onset) [109].

4. rs75097049 (*RP11-690G19.4*; CHG: p=5.37e-14; s=0.030), a regulatory region variant associated with asthma (childhood onset) [109] and neutrophil count [111].

5. rs79411018 (*IKZF3*; EHG: p=5.01e-10; s=0.036), an intron variant associated with atopic asthma [109] and nonatopic asthma [109].

#### Peak 19: ARL17B

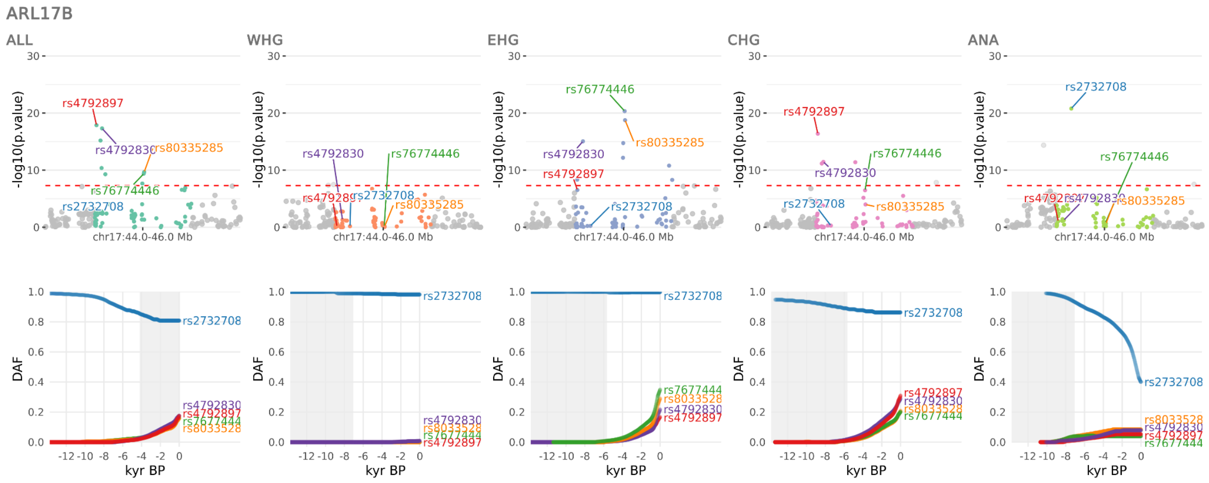


**Figure S74.** Selection at the *ARL17B* locus, spanning chr17:44038785-46012502. Results for the pan-ancestry analysis (ALL) plus each of the four marginal ancestries: Western hunter-gatherers (WHG), Eastern hunter-gatherers (EHG), Caucasus hunter-gatherers (CHG) and Anatolian farmers (ANA). Row one shows zoomed Manhattan plots of the p-values for each ancestry, and row two shows allele trajectories for the top SNPs across all ancestries.

The nineteenth peak spanned the region chr17:44038785-46012502, with the five most significant SNPs (across ancestries) and their associations in the GWAS Catalog (r2023-04-07) being:

1. rs2732708 (*ARL17B*; ANA: p=1.61e-21; s=-0.021), a downstream gene variant/intergenic variant associated with feeling miserable [260] and neuroticism [150].

2. rs76774446 (*GOSR2 / RP11-156P1.2*; EHG: p=4.52e-21; s=0.027), an intron variant associated with cardioembolic stroke (MTAG) [261], atrial fibrillation [262], heart failure (multivariate analysis) [79], and right ventricular end diastolic volume [205].

3. rs4792897 (*MAPT*; ALL: p=1.33e-18; s=0.030), an intron variant associated with snoring [263].

4. rs80335285 (*GOSR2 / RP11-156P1.2*; EHG: p=1.73e-19; s=0.031), an intron variant associated with pulse pressure [251].

5. rs4792830 (*KANSL1*; ALL: p=4.78e-18; s=0.028), an intron variant associated with pulse pressure [82].

#### Peak 20: CTC-258N23.3

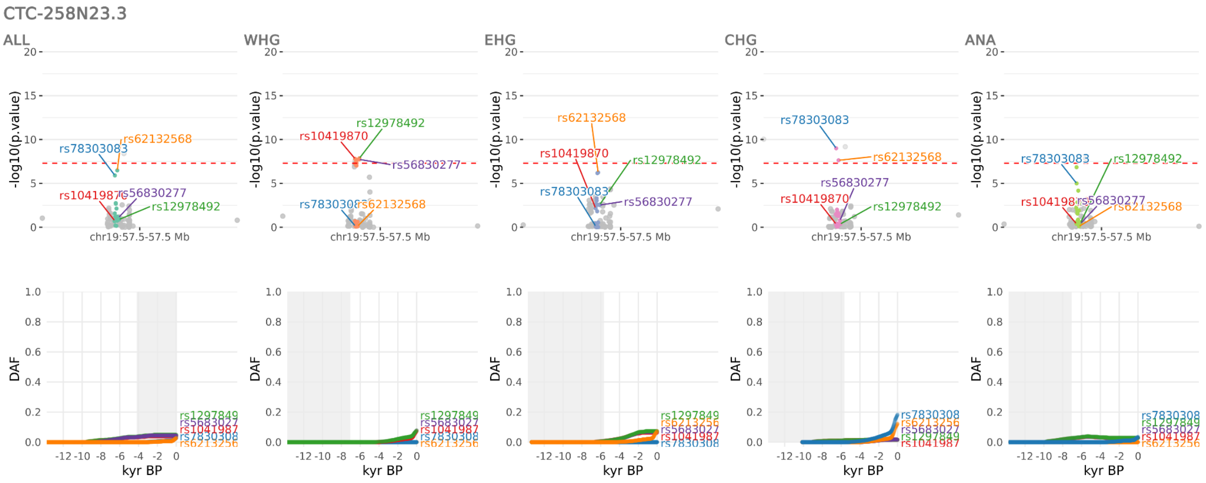


**Figure S75.** Selection at the *CTC-258N23.3* locus, spanning chr19:57495154-57502383. Results for the pan-ancestry analysis (ALL) plus each of the four marginal ancestries: Western hunter-gatherers (WHG), Eastern hunter-gatherers (EHG), Caucasus hunter-gatherers (CHG) and Anatolian farmers (ANA). Row one shows zoomed Manhattan plots of the p-values for each ancestry, and row two shows allele trajectories for the top SNPs across all ancestries.

The twentieth peak spanned the region chr19:57495154-57502383, with the five most significant SNPs (across ancestries) and their associations in the GWAS Catalog (r2023-04-07) being:

1. rs78303083 (*CTC-258N23.3*; CHG: p=9.98e-10; s=0.039), an intergenic variant with no associations in the GWAS Catalog.

2. rs12978492 (*CTC-258N23.3*; WHG: p=1.37e-08; s=0.048), an intergenic variant with no associations in the GWAS Catalog.

3. rs10419870 (*CTC-258N23.3*; WHG: p=1.59e-08; s=0.056), an intergenic variant with no associations in the GWAS Catalog.

4. rs62132568 (*CTC-258N23.3*; CHG: p=2.34e-08; s=0.057), an intergenic variant with no associations in the GWAS Catalog.

5. rs56830277 (*CTC-258N23.3*; WHG: p=1.83e-08; s=0.050), an intergenic variant with no associations in the GWAS Catalog.

Peak 21: CENPM

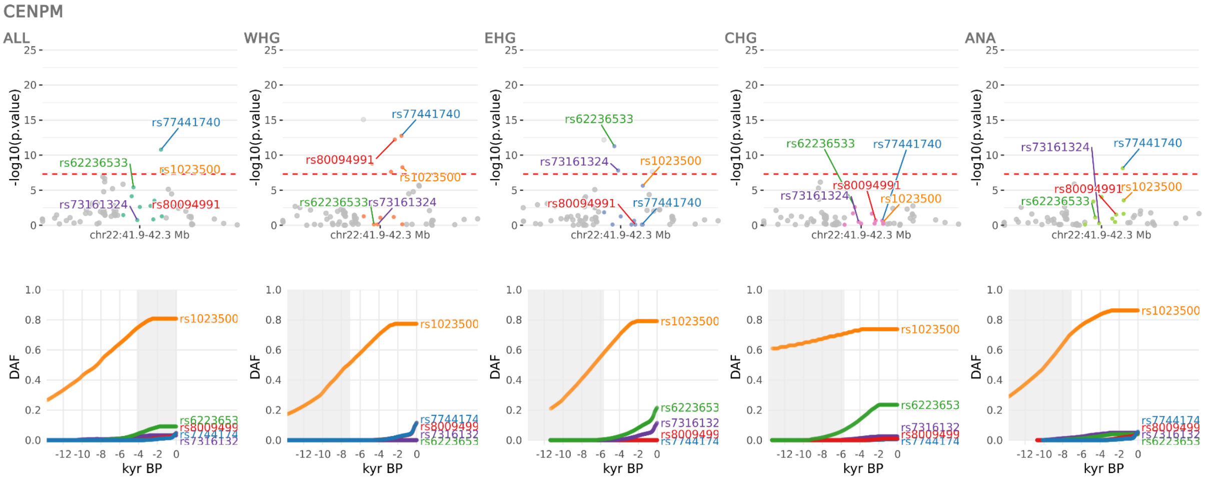


**Figure S76.** Selection at the *CENPM* locus, spanning chr22:41864190-42340844. Results for the pan-ancestry analysis (ALL) plus each of the four marginal ancestries: Western hunter-gatherers (WHG), Eastern hunter-gatherers (EHG), Caucasus hunter-gatherers (CHG) and Anatolian farmers (ANA). Row one shows zoomed Manhattan plots of the p-values for each ancestry, and row two shows allele trajectories for the top SNPs across all ancestries.

The twenty-first peak spanned the region chr22:41864190-42340844, with the five most significant SNPs (across ancestries) and their associations in the GWAS Catalog (r2023-04-07) being:

1. rs77441740 (*CENPM*; WHG: p=1.72e-13; s=0.047), an intergenic variant associated with schizophrenia [264].
2. rs62236533 (*DESI1*; EHG: p=5.29e-12; s=0.029), a downstream gene variant associated with intelligence [103,265], attention deficit hyperactivity disorder or autism spectrum disorder or intelligence [101], cognitive aspects of educational attainment [266], cognitive performance [99], and general cognitive ability [93].
3. rs80094991 (*SREBF2*; WHG: p=5.90e-13; s=0.048), an intron variant associated with highest math class taken (MTAG) [99] and schizophrenia [267].
4. rs1023500 (*CENPM*; WHG: p=5.48e-09; s=0.009), an intron variant associated with schizophrenia [143,268].

5. rs73161324 (*XRCC6*; EHG: p=1.52e-08; s=0.042), an intron variant associated with breast cancer [269] and pulse pressure [270].

##### Selection at the LCT/MCM6 locus

To investigate the timing of the commencement of selection at the LCT/MCM6 sweep locus (chr2:134963892-137613935), we conducted an additional scan in which we ran CLUES without ancestral paintings on all SNPs located within the sweep locus that passed quality control filters (n=6,943). We observed 552 SNPs with a genome-wide significant p-value (p < 5e-8) and confirmed that rs4988235 had the lowest p-value of all SNPs tested (Supplementary Table 8).

We then extracted the maximum likelihood estimate of the starting frequency for each of the 553 genome-wide significant SNPs and calculated the earliest time point by which the selected allele had changed in frequency by >10% in the direction of selection. We then ranked all SNPs by this date metric to determine the relative temporal ordering of major allele frequency changes due to selection. We observed that the vast majority of genome-wide significant selected SNPs began rising in frequency earlier than the lactase persistence allele. Of the 552 SNPs tested, rs4988235 was ranked 475th under this metric (i.e., 86% of selected SNPs showed an earlier rise in frequency). We then inspected the trajectories of higher ranked SNPs and observed that rs1438307 (p=9.77e-24; s=0.0146) began rising in frequency c. 12,000 years ago; approximately 6,000 years earlier than rs4988235 (Figure S78 and Figure S79).

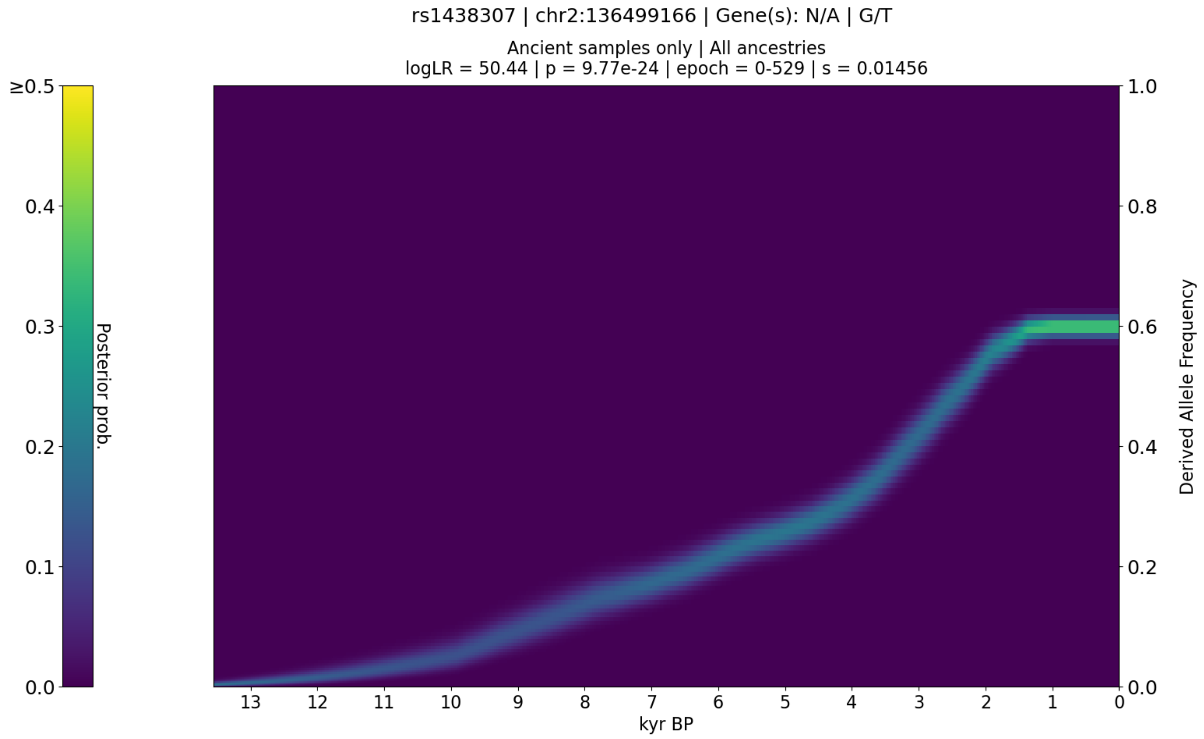


**Figure S77.** CLUES plot of the aDNA time series analysis for all West Eurasian samples in the imputed dataset, showing the posterior probability of the derived allele frequency trajectory for rs1438307.

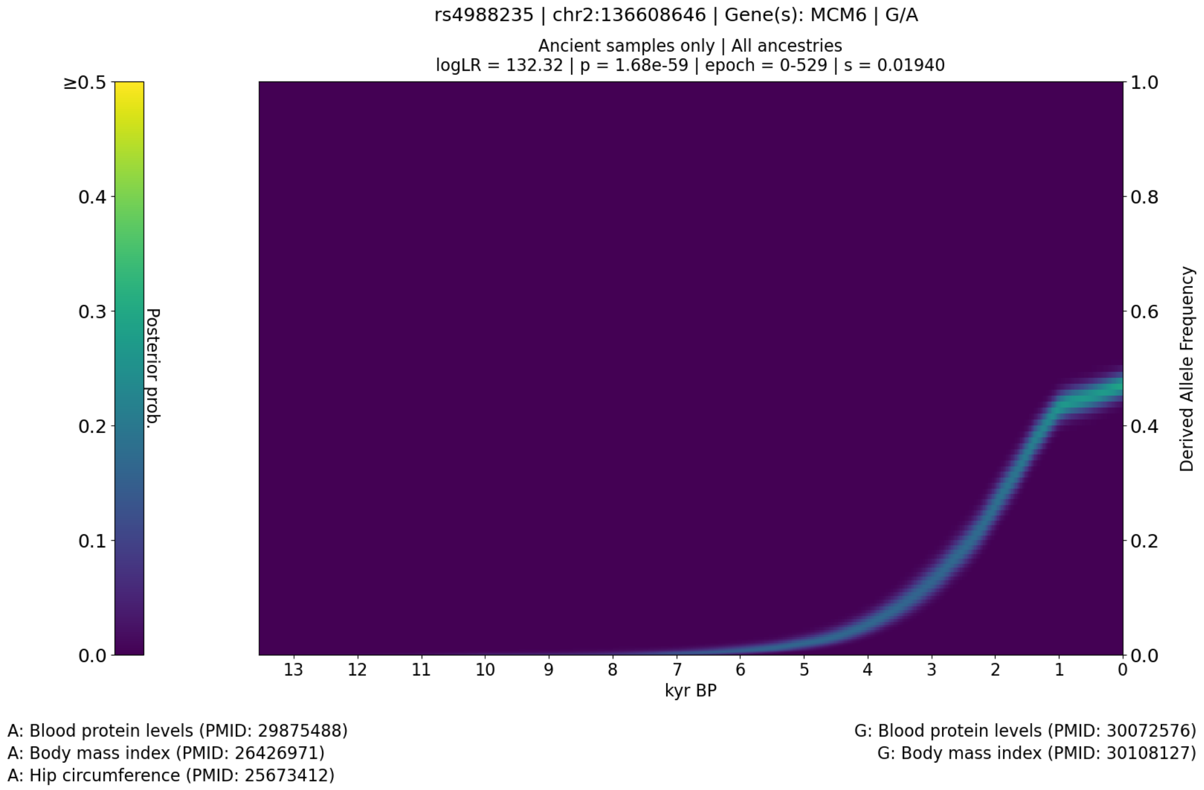


**Figure S78.** CLUES plot of the aDNA time series analysis for all West Eurasian samples in the imputed dataset, showing the posterior probability of the derived allele frequency trajectory for rs4988235.

To control for potential bias introduced by imputation, we replicated our CLUES analyses for both rs4988235 and rs1438307 using genotype likelihoods, instead of imputed genotype probabilities. We observed that trajectories for the imputed and likelihood models were almost identical, indicating that imputation does not bias the selection results for these SNPs (Figure S80 and Figure S81).

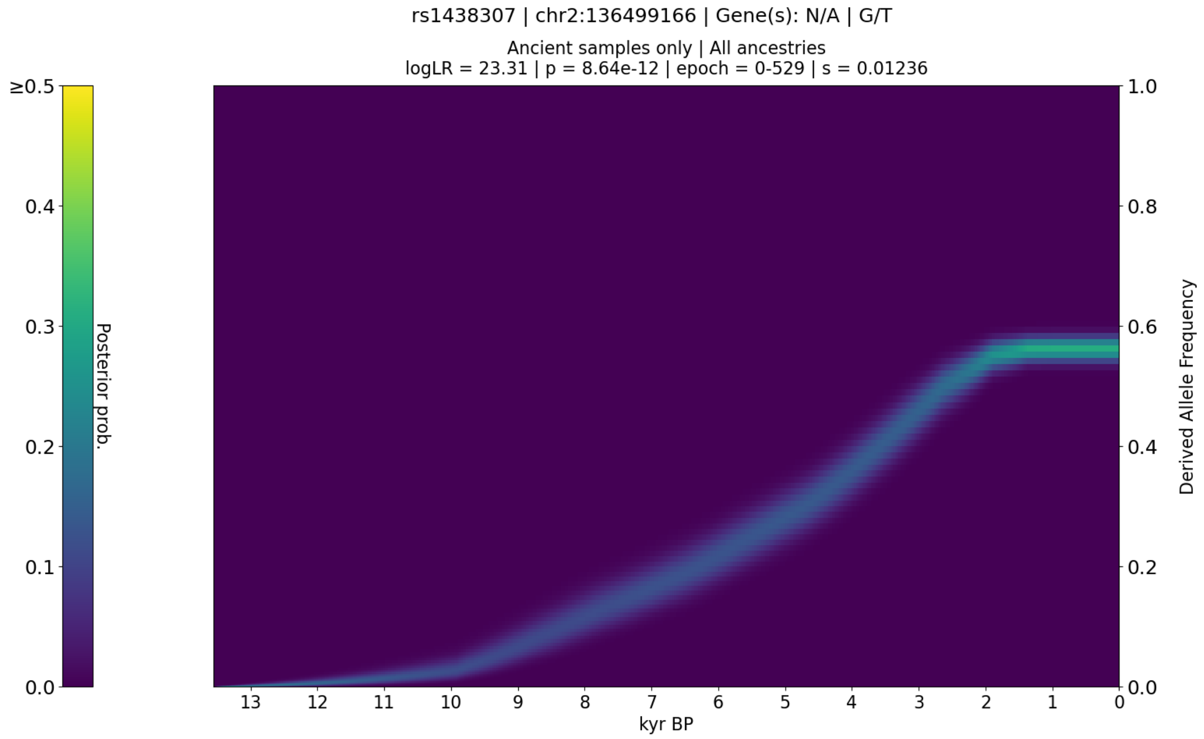


**Figure S79.** *CLUES* plot of the aDNA time series analysis for all West Eurasian samples in the genotype likelihood dataset, showing the posterior probability of the derived allele frequency trajectory for rs1438307.

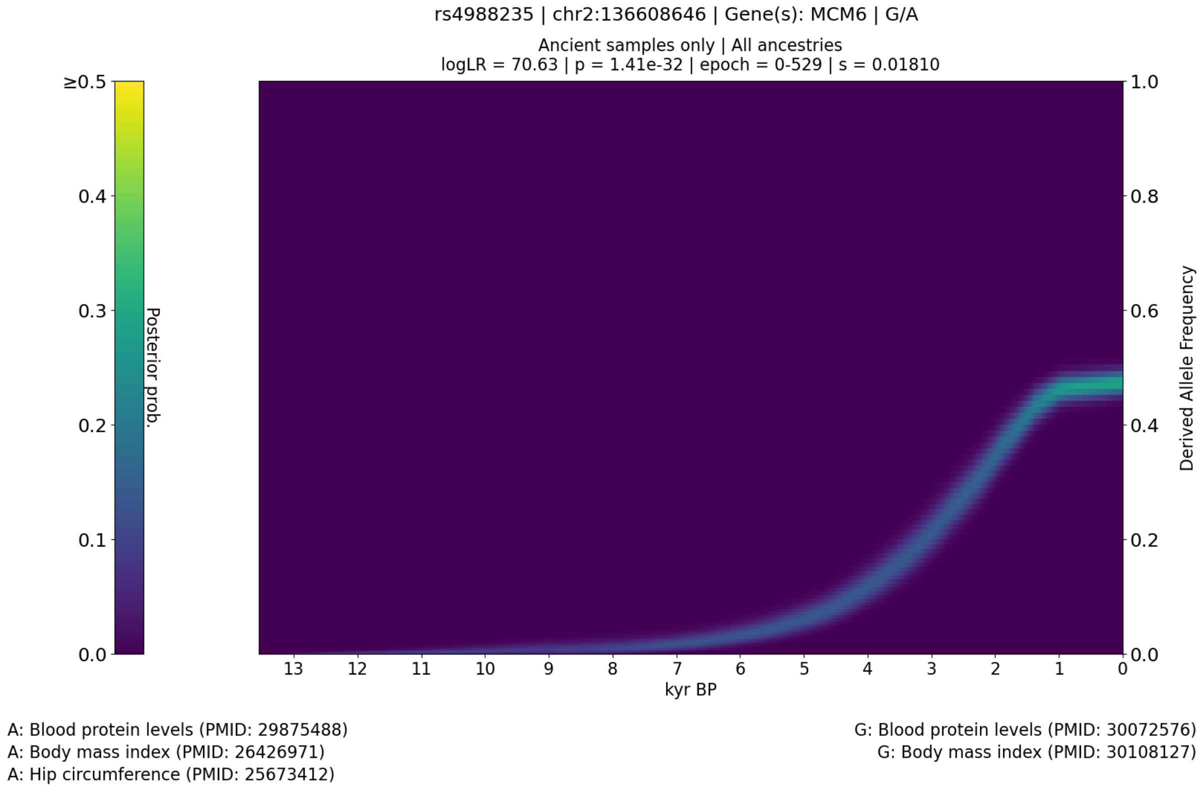

**Figure S80.** *CLUES* plot of the aDNA time series analysis for all West Eurasian samples in the genotype likelihood dataset, showing the posterior probability of the derived allele frequency trajectory for rs4988235.

To control for potential model bias from our modified CLUES software, we extracted both imputed genotype calls, and maximum likelihood calls from the original shotgun dataset, and calculated derived allele frequencies (DAF) in one-thousand-year bins. We plotted these DAF estimates for both SNPs, and both datasets, with a weighted loess regression (Figure **S81**). We observed in both the imputed and non-imputed callsets, that rs1438307 begins rising in frequency thousands of years prior to rs4988235, and that around 6,000 years ago, both SNPs begin to rise rapidly in parallel, due to selection for a haplotype containing both derived alleles; suggestive of a combined fitness benefit to rs1438307 and rs4988235.

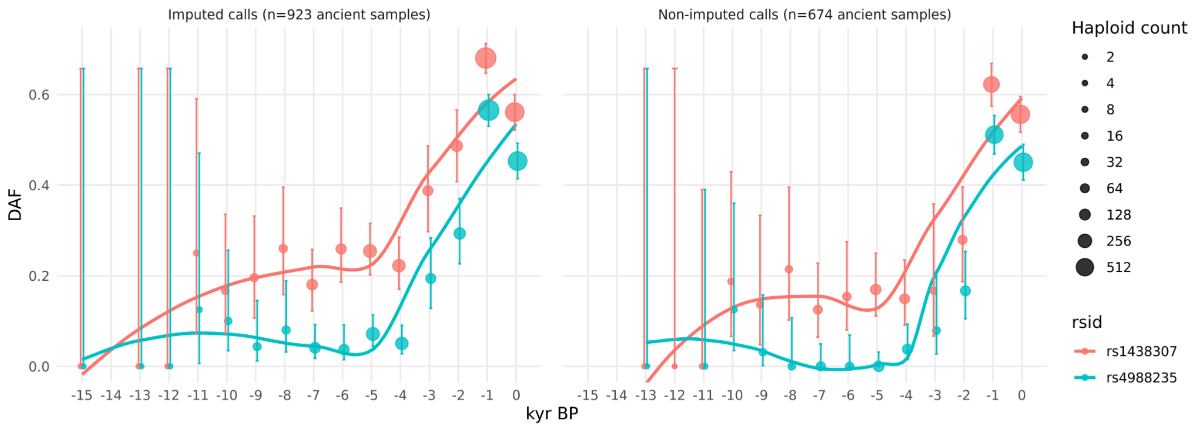


**Figure S81.** Scatter plot of the derived allele frequencies (DAF) for rs1438307 and rs4988235, calculated in one-thousand-year bins. The left-hand panel shows DAF calculated from hard calls in the imputed dataset, and the right-hand panel shows DAF calculated from hard calls in the raw shotgun dataset. Solid lines depict weighted loess regression fits to the observed DAF estimates in each time bin, and error bars show the 95% confidence interval of the DAF estimate in each bin.

We also independently replicated these results using publicly available data from the 1240k capture array. We downloaded genotypes from the Allen Ancient DNA Resource (version 52.2) [271] and filtered for the set of 1,291 West Eurasian samples used in [272], then calculated DAF in one thousand year bins, and plotted a weighted loess regression (Figure S82).

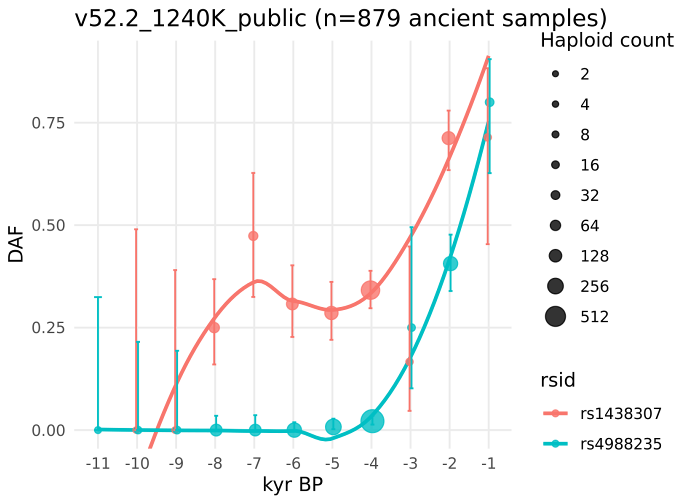


**Figure S82.** Scatter plot of the derived allele frequencies (DAF) for rs1438307 and rs4988235, calculated in one thousand year bins, using publicly available data from the 1240k capture array (Allen Ancient DNA Resource, version 52.2). Solid lines depict weighted loess regression fits to the observed DAF estimates in each time bin, and error bars show the 95% confidence interval of the DAF estimate in each bin.

#### Selection at the APOE locus

To investigate selection at the APOE locus, which is known to significantly mediate risk of developing Alzheimer's disease [273,274], we constructed additional CLUES models in which we called phased haplotypes for three gene alleles: ApoE2 (rs429358:T and rs7412:T), ApoE3 (rs429358:T and rs7412:C), and ApoE4 (rs429358:C and rs7412:C).

Our pan-ancestry analysis identified positive selection favouring ApoE2 (p=6.99e-3; s=0.0130), beginning c. 7,000 years ago and plateauing c. 2,500 years ago (Figure S83). However, we did not identify evidence of selection for either ApoE3 or ApoE4 (Figure S84 and S85).

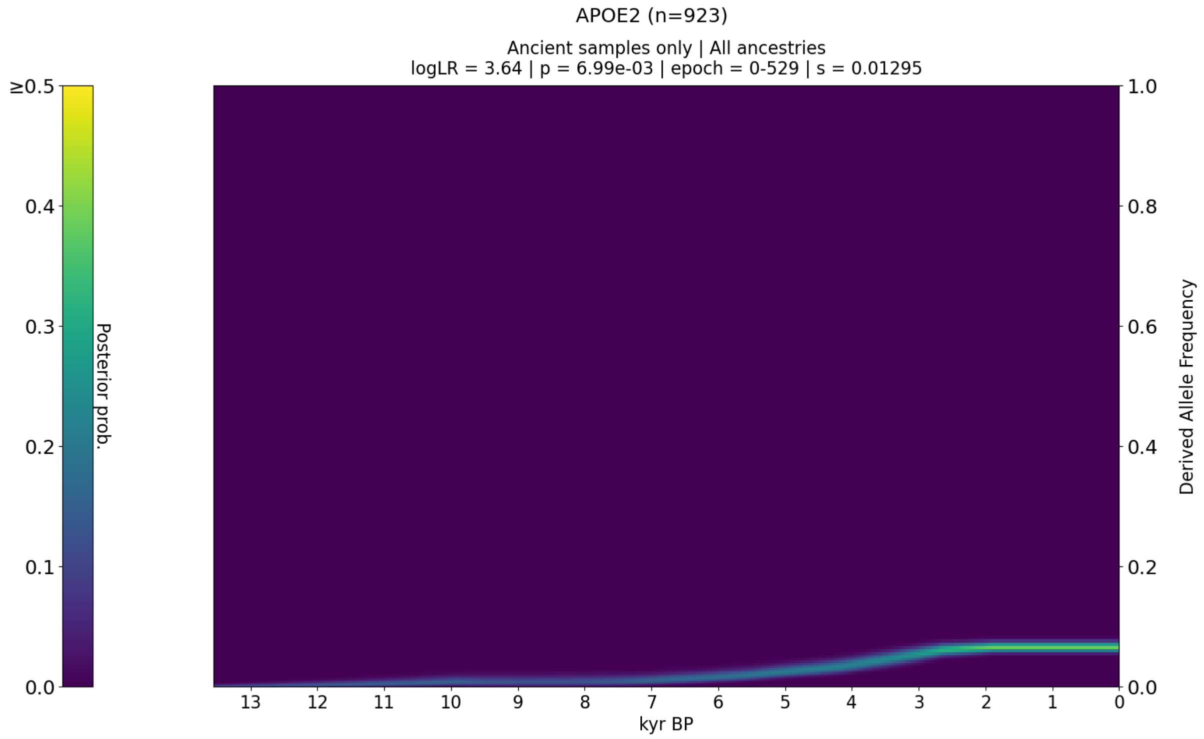


**Figure S83.** CLUES plot of the aDNA time series analysis for all West Eurasian samples in the imputed dataset, showing the posterior probability of the allele frequency trajectory for APOE2.

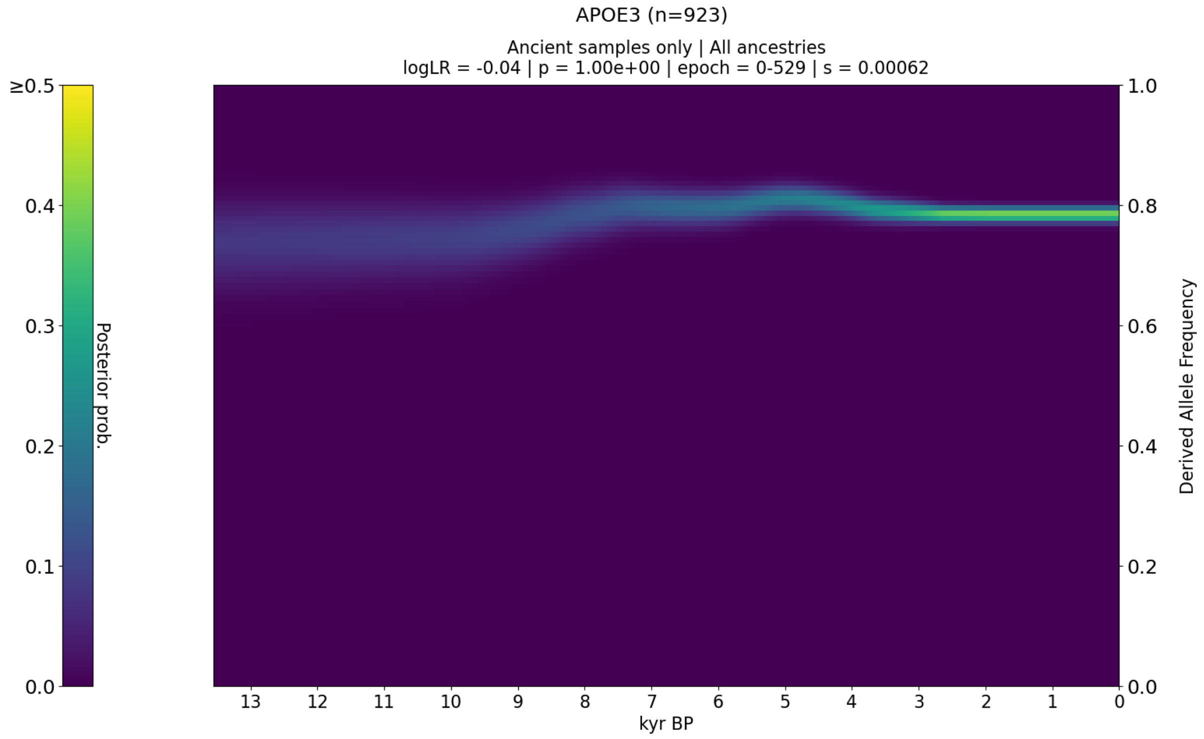

**Figure S84.** CLUES plot of the aDNA time series analysis for all West Eurasian samples in the imputed dataset, showing the posterior probability of the allele frequency trajectory for APOE3.

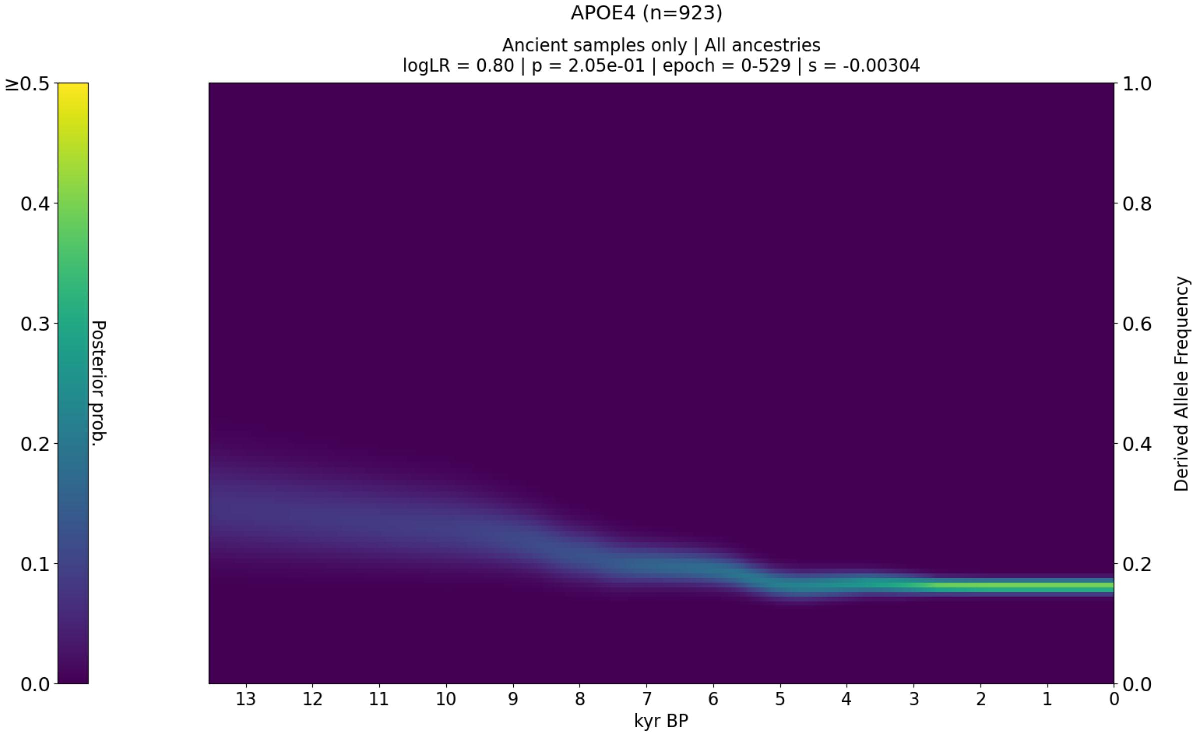


**Figure S85.** CLUES plot of the aDNA time series analysis for all West Eurasian samples in the imputed dataset, showing the posterior probability of the allele frequency trajectory for APOE4.

#### Discussion

Using our ancient genomic panel, we sought to identify phenotype-associated variants that have evidence for directional selection over the last 13,000 years. To estimate allele frequency trajectories and selection coefficients of trait-associated variants through time, we used the software *CLUES* [1] which can perform inference of allele frequency trajectories using marginal trees sampled from a reconstruction of an ancestral recombination graph (ARG) [2] for a set of genomic sequences, in combination with genotype likelihoods from serially sampled ancient DNA (aDNA).

Our results show that the incorporation of ancient DNA considerably improves our power to detect variants under selection, compared to a method that only uses the ARG inferred from present-day data alone. Using genomes from the 1,000 Genomes Project project (populations GBR, FIN and TSI), we inferred allele trajectories and selection coefficients for 32,079 phenotype-associated variants, ascertained from the GWAS Catalog [4], along with an equal number of putatively neutral “control” variants. Our analysis identified no genome-wide significant selective sweeps (p < 5e-8) using present-day data alone. However, the trait-associated variants were significantly enriched for evidence of selection when compared to the control group (Wilcoxon signed-rank test, p < 7.29e-35).

##### Pan-ancestry selection

In contrast to patterns observed in present-day genomes, our selection analysis based on a time-series of ancient DNA genotype probabilities identified 11 genome-wide significant (p < 5e-8) selective sweeps in the GWAS variants, and none in the control group; consistent with widespread selection acting on trait-associated variants. This analysis confirms many of the previously reported selection loci in West Eurasians, identified from present-day and ancient DNA [18,275–277], and reveals novel selective sweeps, while refining the temporal dynamics of the selected alleles.

The strongest overall signal of selection in the pan-ancestry analysis is at the LCT / MCM6 locus (rs4988235:A; p=1.68e-59; s=0.0194), the derived allele of which is casual for lactase persistence [40,41]. The inferred trajectory indicates that this allele began rising in frequency c. 6,000 years ago, and has continued to rise in frequency up to the present (Figure S31). However, we also observe that there is evidence of at least two selective sweeps at this locus, targeting variants with strikingly different metabolic effects. We identified evidence of selection at this locus beginning c. 6,000 years prior to the emergence of the lactase persistence allele, favouring a variant rs1438307 (p=9.77e-24; s=0.0146) which has been hypothesised as an ancient adaptation to famine [278]. We replicated these findings with a CLUES model using genotype-likelihoods (called directly from the aDNA sequencing reads) (Figure S79-S80), and with binned allele frequencies called from both the imputed and shotgun datasets (Figure S81), and using an independent dataset genotyped with difference sequencing technology (1240k capture array) (Figure S82).

We find a strong signal of selection at the *FADS1* (rs174546:C; p=4.41e-19; s=0.0126) and *FADS2* (rs174581:G; p=2.21e-19; s=0.0138) locus, associated with fatty acid metabolism [166,173,175,279–281]. The trajectories for these variants indicate a rise in frequency, beginning around 13,000 years ago, and continuing up until c. 2,000 years ago, after which their frequencies plateaued (Figure S47).

We detect an 8 megabase (Mb) wide selection sweep signal in chromosome 6 (chr6:25.4-33.5 Mb), spanning the human leukocyte antigen (HLA) region. The selection trajectories of the variants within this locus support multiple independent sweeps, occurring at different times and with differing intensities. The strongest signal of selection at this locus in the pan-ancestry analysis is at an intergenic variant, located between *HLA-A* and *HLA-W* (rs7747253:A; p=7.56e-32; s=-0.0178), associated with heel bone mineral density [70], the derived allele of which rapidly reduced in frequency, beginning c. 8,000 years ago. In contrast, the signal of selection at C2 (rs9267677:C; p= 6.60e-26; s= 0.0441), also found within this sweep shows a gradual increase in frequency beginning c. 4,000 years ago, before rising more rapidly c. 1,000 years ago (Figure S43). In this case, the favoured allele is associated with protection against some sexually transmitted diseases [282], and with increased psoriasis risk [283,284].

We also identified selection signals at the *SLC22A4* (rs35260072:C; p=8.49e-20; s=0.0172) locus, associated with increased itch intensity from mosquito bites [38] protection against childhood and adult asthma [282] and asthma-related infections [282] and we find that the derived variant has been steadily rising in frequency since c. 9,000 years ago (Figure S37). However, in the same *SLC22A4* candidate region as rs35260072, we find that the frequency of the previously reported allele rs1050152:T (which also protects against asthma and related infections [282]) plateaued c. 1,500 years ago, contrary to previous reports suggesting a recent rise in frequency [18]. Similarly, we detect selection at the *HECTD4* (rs11066188:A; p=9.51e-31 s=0.0198) and *ATXN2* (rs653178:C; p=3.73e-29; s=0.0189) loci, both of which have been rising in frequency for c. 9,000 years (Figure S70), also contrary to previous reports of a more recent rise in frequency [18].

We detect strong selection at the *SLC45A2* locus (rs35395:C; p=1.60e-44; s=0.0215) associated with skin pigmentation [56,285], and find that the selected allele began increasing in frequency from c. 13,000 years ago, until plateauing c. 2,000 years ago (Figure S35). The predominant hypothesis is that high melanin levels in the skin are important in equatorial regions owing to its protection against UV radiation, whereas lighter skin has been selected for at higher latitudes (where UV radiation is less intense) because some UV penetration is required for cutaneous synthesis of vitamin D [286,287]. Our findings confirm pigmentation alleles as major targets of selection during the Holocene [18,275,276] particularly on a small proportion of loci with large effect sizes [285].

We further detect signs of strong selection in a 2 Mb sweep on chromosome 17 (chr17:44.0-46.0), spanning a locus on 17q21.3, implicated in neurodegenerative and developmental disorders. The locus includes an inversion and other structural polymorphisms with indications of a recent positive selection sweep in some human populations [288,289]. The strongest signal of selection observed in the pan-ancestry analysis at this locus is at MAPT (rs4792897:G; p=1.33e-18; s=0.0299 (Figure S74), which codes for the tau protein [290], and is associated with protection against mumps [282] and increased risk of snoring [263]. More generally, polymorphisms in MAPT have been associated with increased risk of a number of neurodegenerative disorders, including Alzheimer's disease and Parkinson's disease [291–295]. However, we caution that this region is also enriched for evidence of reference bias in our dataset—especially around the KANSL1 gene—due to complex structural polymorphisms (Supplementary Note 10).

#### Ancestry stratified selection trajectories

To account for population structure in our samples, we also applied a novel chromosome painting technique, based on inference of a sample's nearest neighbours in the marginal trees of an ARG that contains individuals classified into different ancient populations (Supplementary Note 3). This method allows us to accurately assign ancestral population labels to haplotypes found in both ancient and present-day individuals. By conditioning our selection analyses on these haplotype backgrounds, we can infer the selection trajectories of GWAS risk alleles in a manner that is approximately invariant to change in the admixture proportions through time. These ancestry specific allele trajectories reveal many novel aspects about the dynamic interplay between selection and admixture in West Eurasia throughout the Holocene. We find that the allele trajectories of directionally selected sites become much more apparent once we perform this ancestry partitioning. We often find variants with strong allele frequency changes in one ancestral population but not another, and analysing all ancient individuals without accounting for their ancestry composition leads to a decrease in our ability to identify selected variants, and a blurring of the temporal signal of allele frequency changes.

When conditioned on one of our four marginal ancestries—Western hunter-gatherers (WHG), Eastern hunter-gatherers (EHG), Caucasus hunter-gatherers (CHG) and Anatolian farmers (ANA)—we find 21 genome-wide significant selection peaks (almost twice as many as in our pan-ancestry analysis), indicating that admixture between ancestral populations has masked evidence of selection at many loci. Furthermore, we find that some strong signals of selection identified in the pan-ancestry analysis are driven by sweeps in the marginal ancestries which substantially differ in their most significant SNPs, suggesting that multiple selected alleles may be common within genome-wide significant sweep loci.

Our analysis suggests that there are several reasons why we detect more selection when conditioning on ancestry; one of which is increased statistical power. In cases where the selected allele is not segregating in all ancestral backgrounds (e.g., if it is private to one ancestral path), stratification by ancestry increases our statistical power to detect selection, as it allows us to separate haplotypes in which the selected allele is absent (i.e., where selection can have no observable effect). Similarly, in cases where selection is influenced by epistasis, stratification by ancestry may allow us to separate haplotypes that contain only a subset of the adaptive markers.

However, the main reason we detect more selection is due to the effects of multiple waves of admixture. Our pan-ancestry analysis spans three major waves of admixture (Figure **S7**), which coincide with dramatic changes in subsistence strategy, as well as large movements of people into new environmental niches. In cases where selection is acting in only one population, admixture can confound analyses based on time-series data, especially when the admixing populations have substantially different allele frequencies. Stratifying by ancestry controls for this effect, as it allows us to model changes in allele frequency independently of changes in admixture fraction.

##### Robustness of analysis

To test for potential effects of imputation bias on our selection analyses, we ran additional models for each of the top SNPs in the pan-ancestry analysis, using genotype likelihoods (GL) called directly from the aDNA sequencing reads. Because our chromosome painting model requires phased haplotypes, this replication test was limited to pan-ancestry models only. When comparing the imputed and GL selection models, we observed that the posterior likelihood densities of the allele frequency trajectories are highly correlated, as are the selection coefficients. Due to the smaller sample sizes in the GL callset we had less power to reject neutrality, and the inferred log-likelihood ratio test statistics were consistently lower than in the imputed models, but all remained high. Overall, we did not detect any notable bias when comparing these two sets of models.

To test for potential effects of painting bias on our selection analyses, we performed neutral simulations using the same demographic model used to train the classifier. We applied the chromosome painting model to the simulated VCF, and used CLUES to infer allele frequency trajectories and selection coefficients for frequency paired simulated SNPs, in both a pan-ancestry analysis and stratified by each of the four ancestral paths. We then applied the same thresholds used in the empirical analysis to detect selective sweep loci. We observed zero genome-wide significant selective sweep loci across all five analyses. At a SNP level we observed 22 false-positive SNPs (0.75%) across all analyses (Figure S53). Due to the stochastic occurrence of false-positive SNPs along the chromosome, no false-positive sweep loci were detected. Detection of a false-positive sweep locus would require a cluster of at least 6 genome-wide significant SNPs within a +/- 1 Mb locus. From this analysis we conclude that the low rate of error in our chromosome painting model is unlikely to bias the inference of sweep loci, but may produce a small number of randomly occurring false-positive SNPs.

To control for errors specific to ancient DNA damage, we developed two new quality-control metrics for filtering sites with evidence of potential bias. Firstly, we developed a novel statistic ($F_j$) for detecting correlations between characteristics of aDNA damage (i.e., depth of coverage, read length and error rate) and changes in allele frequencies over time (Supplementary Note 6). The purpose of the $F_j$ statistic is to identify SNPs where the observed time-series of aDNA genotypes may be biased by age dependent preservation characteristics. We also developed a second metric, intended to detect reference and mapping bias when analysing ancient and modern DNA together. Due to the characteristics of aDNA damage, some sites may be enriched for mapping bias, which favours observations of the reference allele. This can result in systematic differences between allele frequencies calculated from aDNA and modern data. To control for this, we developed a test to filter out sites exhibiting substantial differences between present-day allele frequencies inferred from ancient and modern data respectively.

Finally, to test for potential effects from unmodeled phenomena, we ascertained a set of putatively neutral “control” SNPs. These SNPs were drawn at random from regions of the genome at least 50 kb from a GWAS SNP or gene region, and frequency paired with each GWAS SNP based on their derived allele frequency (DAF). We then ran the Control SNPs through the same selection pipeline as the GWAS SNPs. Our ancestry stratified results detected 19 genome-wide significant selective sweeps in the GWAS group, and 2 in the Control group. Upon further investigation, one of the two sweeps identified in the Control group (chr19:57.5-57.5 Mb) contains genome-wide significant SNPs that were not reported in the GWAS Catalog, but are reported in the UK Biobank (i.e., rs959939, rs2102540, rs56830277 and rs12978492 for phenotype code '20024_1121'). Interpretation of the remaining sweep is less clear, as it is entirely possible that a non-GWAS locus may be a target of selection. Overall, these results indicate that error propagation between analysis steps is not a major source of bias, as the Control SNPs are themselves subject to the same phasing, imputation, painting and selection analyses as the GWAS SNPs and yet we find a 20-to-1 ratio of sweep loci that contain significant GWAS trait associations.

Other recent papers have also modelled selection in West Eurasia during the Holocene [272,296]; however, it is difficult to directly compare results due to substantial differences in methodology and sampling. Lee et al. [272] use an updated version of the mixture model developed in Mathieson et al. [18], which relies on differences in allele frequencies postdating admixture. As such, they are best powered to detect rapid episodes of selection following admixture between populations. The selection test used by Kerner et al. [296] is based on choosing variants with an estimated selection coefficient above the 99th quantile from their simulations, and is therefore best-powered to detect cases of strong selection. In our analyses, we used a selection test that is well-powered to detect both weak and strong selection, and we used local ancestry inference to deconvolute the effects of changes in admixture proportions through time, allowing us to detect selection in a broader range of demographic scenarios.

Another key difference is in sampling. Both Lee et al. [272] and Kerner et al. [296] used pseudohaploid data from the 1240k capture array, which is affected by allelic bias, due to the capture chemistry [297,298]. It remains unclear how sensitive selection results from the 1240k array are to systematic bias in the recovery of some alleles; however, Kerner et al. [296] found that nine of the top 10 variants in their capture dataset had a frequency trajectory inconsistent with their shotgun dataset. This suggests that allelic bias from the 1240k capture chemistry may be a major confounder for tests of selection. In comparison to shotgun data, Rohland et al. [297] found that 61.7% of the SNPs on the 1240k capture array exhibit evidence of allelic bias (n=757,587 with `PassFilterForMetaAnalysisBias==0`).

A compounding factor may also be systematic differences in capture efficiency between sites, which results in much smaller sample sizes than the reported number of ancient individuals. For example, in our analysis of selection at the LCT locus using the 1240k dataset (Figure S82)—which used the same 1,291 samples as Lee et al. (2022)—we observed that there were 838 pseudohaploid calls for rs4988235, but only 476 for rs1438307, indicating capture efficiency varies greatly between sites, as well as between alleles at the same site. In comparison, our imputed callset contains 1,015 diploid genotypes for all modelled SNPs, and we show via replication (using genotype-likelihoods) that imputation does not substantively bias our inference of allele frequency trajectories or selection coefficients.

#### References


1. Stern, A. J., Wilton, P. R. & Nielsen, R. An approximate full-likelihood method for inferring selection and allele frequency trajectories from DNA sequence data. *PLoS Genet.* **15**, e1008384 (2019).

2. Speidel, L., Forest, M., Shi, S. & Myers, S. R. A method for genome-wide genealogy estimation for thousands of samples. *Nat. Genet.* **51**, 1321–1329 (2019).

3. Köster, J. & Rahmann, S. Snakemake--a scalable bioinformatics workflow engine. *Bioinformatics* **28**, 2520–2522 (2012).

4. Buniello, A. *et al.* The NHGRI-EBI GWAS Catalog of published genome-wide association studies, targeted arrays and summary statistics 2019. *Nucleic Acids Res.* **47**, D1005–D1012 (2019).

5. Jupp, S., Burdett, T., Leroy, C. & Parkinson, H. E. A new Ontology Lookup Service at EMBL-EBI. in *SWAT4LS* 118–119 (ceur-ws.org, 2015).

6. Yates, A. *et al.* The Ensembl REST API: Ensembl Data for Any Language. *Bioinformatics* **31**, 143–145 (2015).

7. Allentoft, M. E., Sikora, M. & Refoyo-Martínez, A. Population Genomics of Stone Age Eurasia. *bioRxiv* (2022).

8. Aken, B. L. *et al.* The Ensembl gene annotation system. *Database* **2016**, (2016).

9. Kelleher, J., Etheridge, A. M. & McVean, G. Efficient Coalescent Simulation and Genealogical Analysis for Large Sample Sizes. *PLoS Comput. Biol.* **12**, e1004842 (2016).

10. 1000 Genomes Project Consortium *et al.* A global reference for human genetic variation. *Nature* **526**, 68–74 (2015).

11. Speidel, L. *et al.* Inferring Population Histories for Ancient Genomes Using Genome-Wide Genealogies. *Mol. Biol. Evol.* **38**, 3497–3511 (2021).

12. Vilella, A. J. *et al.* EnsemblCompara GeneTrees: Complete, duplication-aware phylogenetic trees in vertebrates. *Genome Res.* **19**, 327–335 (2009).

13. Rasmussen, M. D., Hubisz, M. J., Gronau, I. & Siepel, A. Genome-wide inference of ancestral recombination graphs. *PLoS Genet.* **10**, e1004342 (2014).

14. Hein, J., Schierup, M. & Wiuf, C. *Gene Genealogies, Variation and Evolution: A primer in coalescent theory*. (Oxford University Press, USA, 2004).

15. Stern, A. J., Speidel, L., Zaitlen, N. A. & Nielsen, R. Disentangling selection on genetically correlated polygenic traits via whole-genome genealogies. *Am. J. Hum. Genet.* (2021) doi:10.1016/j.ajhg.2020.12.005.

16. 1000 Genomes Project Consortium *et al.* A map of human genome variation from population-scale sequencing. *Nature* **467**, 1061–1073 (2010).

17. Moorjani, P. *et al.* A genetic method for dating ancient genomes provides a direct estimate of human generation interval in the last 45,000 years. *Proc. Natl. Acad. Sci. U. S. A.* **113**, 5652–5657 (2016).

18. Mathieson, I. *et al.* Genome-wide patterns of selection in 230 ancient Eurasians. *Nature* **528**, 499–503 (2015).

19. Li, H. A statistical framework for SNP calling, mutation discovery, association mapping and population genetical parameter estimation from sequencing data. *Bioinformatics* **27**, 2987–2993 (2011).

20. Quinlan, A. R. & Hall, I. M. BEDTools: a flexible suite of utilities for comparing genomic features. *Bioinformatics* **26**, 841–842 (2010).

21. Cock, P. J. A. *et al.* Biopython: freely available Python tools for computational

molecular biology and bioinformatics. *Bioinformatics* **25**, 1422–1423 (2009).

22. Anaconda Software Distribution. *Anaconda Documentation* Preprint at https://docs.anaconda.com/ (2020).

23. Harris, C. R. *et al.* Array programming with NumPy. *Nature* **585**, 357–362 (2020).

24. McKinney, W. & Others. Data structures for statistical computing in python. in *Proceedings of the 9th Python in Science Conference* vol. 445 51–56 (Austin, TX, 2010).

25. Li, H. *et al.* The Sequence Alignment/Map format and SAMtools. *Bioinformatics* **25**, 2078–2079 (2009).

26. Van Rossum, G. & Drake, F. L. *Python 3 Reference Manual: (Python Documentation Manual Part 2).* (CreateSpace Independent Publishing Platform, 2009).

27. R Core Team. R: A Language and Environment for Statistical Computing. Preprint at https://www.R-project.org/ (2019).

28. Haider, S. *et al.* A bedr way of genomic interval processing. *Source Code Biol. Med.* **11**, 14 (2016).

29. Wickham, H., François, R., Henry, L. & Müller, K. dplyr: A Grammar of Data Manipulation. Preprint at https://CRAN.R-project.org/package=dplyr (2019).

30. Wickham, H. *ggplot2: Elegant Graphics for Data Analysis*. (Springer, 2016).

31. Petukhov, V., van den Brand, T. & Biederstedt, E. ggrastr: Raster Layers for ‘ggplot2’. Preprint at https://CRAN.R-project.org/package=ggrastr (2020).

32. Slowikowski, K. ggrepel: Automatically Position Non-Overlapping Text Labels with ‘ggplot2’. Preprint at https://CRAN.R-project.org/package=ggrepel (2020).

33. Wilke, C. O. ggridges: Ridgeline Plots in ‘ggplot2’. Preprint at https://CRAN.R-project.org/package=ggridges (2018).

34. Wickham, H. stringr: Simple, Consistent Wrappers for Common String

Operations. Preprint at https://CRAN.R-project.org/package=stringr (2019).

35. Virtanen, P. *et al.* SciPy 1.0: fundamental algorithms for scientific computing in Python. *Nat. Methods* **17**, 261–272 (2020).

36. Kurilshikov, A. *et al.* Large-scale association analyses identify host factors influencing human gut microbiome composition. *Nat. Genet.* **53**, 156–165 (2021).

37. Sun, B. B. *et al.* Genomic atlas of the human plasma proteome. *Nature* **558**, 73–79 (2018).

38. Jones, A. V. *et al.* GWAS of self-reported mosquito bite size, itch intensity and attractiveness to mosquitoes implicates immune-related predisposition loci. *Hum. Mol. Genet.* **26**, 1391–1406 (2017).

39. Schlosser, P. *et al.* Genetic studies of urinary metabolites illuminate mechanisms of detoxification and excretion in humans. *Nat. Genet.* **52**, 167–176 (2020).

40. Enattah, N. S. *et al.* Identification of a variant associated with adult-type hypolactasia. *Nat. Genet.* **30**, 233–237 (2002).

41. Bersaglieri, T. *et al.* Genetic signatures of strong recent positive selection at the lactase gene. *Am. J. Hum. Genet.* **74**, 1111–1120 (2004).

42. Yin, X. *et al.* Genome-wide association studies of metabolites in Finnish men identify disease-relevant loci. *Nat. Commun.* **13**, 1644 (2022).

43. Chen, Y. *et al.* Genomic atlas of the plasma metabolome prioritizes metabolites implicated in human diseases. *Nat. Genet.* **55**, 44–53 (2023).

44. Winkler, T. W. *et al.* The Influence of Age and Sex on Genetic Associations with Adult Body Size and Shape: A Large-Scale Genome-Wide Interaction Study. *PLoS Genet.* **11**, e1005378 (2015).

45. Hoffmann, T. J. *et al.* A Large Multiethnic Genome-Wide Association Study of Adult Body Mass Index Identifies Novel Loci. *Genetics* **210**, 499–515 (2018).

46. Richardson, T. G. *et al.* Characterising metabolomic signatures of lipid-modifying therapies through drug target mendelian randomisation. *PLoS Biol.* **20**, e3001547 (2022).

47. Qin, Y. *et al.* Combined effects of host genetics and diet on human gut microbiota and incident disease in a single population cohort. *Nat. Genet.* **54**, 134–142 (2022).

48. Emilsson, V. *et al.* Co-regulatory networks of human serum proteins link genetics to disease. *Science* **361**, 769–773 (2018).

49. Huang, L. O. *et al.* Genome-wide discovery of genetic loci that uncouple excess adiposity from its comorbidities. *Nat Metab* **3**, 228–243 (2021).

50. Koskeridis, F. *et al.* Pleiotropic genetic architecture and novel loci for C-reactive protein levels. *Nat. Commun.* **13**, 6939 (2022).

51. May-Wilson, S. *et al.* Large-scale GWAS of food liking reveals genetic determinants and genetic correlations with distinct neurophysiological traits. *Nat. Commun.* **13**, 2743 (2022).

52. Shungin, D. *et al.* New genetic loci link adipose and insulin biology to body fat distribution. *Nature* **518**, 187–196 (2015).

53. Pietzner, M. *et al.* Mapping the proteo-genomic convergence of human diseases. *Science* **374**, eabj1541 (2021).

54. Kiiskinen, T. *et al.* Genetic predictors of lifelong medication-use patterns in cardiometabolic diseases. *Nat. Med.* **29**, 209–218 (2023).

55. Ahsan, M. *et al.* The relative contribution of DNA methylation and genetic variants on protein biomarkers for human diseases. *PLoS Genet.* **13**, e1007005 (2017).

56. Lona-Durazo, F. *et al.* Meta-analysis of GWA studies provides new insights on the genetic architecture of skin pigmentation in recently admixed populations. *BMC Genet.* **20**, 59 (2019).

57. Kanai, M. *et al.* Genetic analysis of quantitative traits in the Japanese population links cell types to complex human diseases. *Nat. Genet.* **50**, 390–400 (2018).

58. Katz, D. H. *et al.* Whole Genome Sequence Analysis of the Plasma Proteome in Black Adults Provides Novel Insights Into Cardiovascular Disease. *Circulation* **145**, 357–370 (2022).

59. König, E. *et al.* Whole Exome Sequencing Enhanced Imputation Identifies 85 Metabolite Associations in the Alpine CHRIS Cohort. *Metabolites* **12**, (2022).

60. Zhu, X., Zhu, L., Wang, H., Cooper, R. S. & Chakravarti, A. Genome-wide pleiotropy analysis identifies novel blood pressure variants and improves its polygenic risk scores. *Genet. Epidemiol.* **46**, 105–121 (2022).

61. Surapaneni, A. *et al.* Identification of 969 protein quantitative trait loci in an African American population with kidney disease attributed to hypertension. *Kidney Int.* **102**, 1167–1177 (2022).

62. Png, G. *et al.* Identifying causal serum protein-cardiometabolic trait relationships using whole genome sequencing. *Hum. Mol. Genet.* **32**, 1266–1275 (2023).

63. Ahola-Olli, A. V. *et al.* Genome-wide Association Study Identifies 27 Loci Influencing Concentrations of Circulating Cytokines and Growth Factors. *Am. J. Hum. Genet.* **100**, 40–50 (2017).

64. Tahir, U. A. *et al.* Whole Genome Association Study of the Plasma Metabolome Identifies Metabolites Linked to Cardiometabolic Disease in Black Individuals. *Nat. Commun.* **13**, 4923 (2022).

65. Gudjonsson, A. *et al.* A genome-wide association study of serum proteins reveals shared loci with common diseases. *Nat. Commun.* **13**, 480 (2022).

66. Leskelä, J. *et al.* Genetic Profile of Endotoxemia Reveals an Association With

Thromboembolism and Stroke. *J. Am. Heart Assoc.* **10**, e022482 (2021).

67. Feofanova, E. V. *et al.* A Genome-wide Association Study Discovers 46 Loci of the Human Metabolome in the Hispanic Community Health Study/Study of Latinos. *Am. J. Hum. Genet.* **107**, 849–863 (2020).

68. Thareja, G. *et al.* Differences and commonalities in the genetic architecture of protein quantitative trait loci in European and Arab populations. *Hum. Mol. Genet.* **32**, 907–916 (2023).

69. Baselmans, B. M. L. *et al.* Multivariate genome-wide analyses of the well-being spectrum. *Nat. Genet.* **51**, 445–451 (2019).

70. Morris, J. A. *et al.* An atlas of genetic influences on osteoporosis in humans and mice. *Nat. Genet.* **51**, 258–266 (2019).

71. Christakoudi, S., Evangelou, E., Riboli, E. & Tsilidis, K. K. GWAS of allometric body-shape indices in UK Biobank identifies loci suggesting associations with morphogenesis, organogenesis, adrenal cell renewal and cancer. *Sci. Rep.* **11**, 10688 (2021).

72. Wootton, R. E. *et al.* Evidence for causal effects of lifetime smoking on risk for depression and schizophrenia: a Mendelian randomisation study. *Psychol. Med.* **50**, 2435–2443 (2020).

73. Cadby, G. *et al.* Comprehensive genetic analysis of the human lipidome identifies loci associated with lipid homeostasis with links to coronary artery disease. *Nat. Commun.* **13**, 3124 (2022).

74. Harshfield, E. L. *et al.* Genome-wide analysis of blood lipid metabolites in over 5000 South Asians reveals biological insights at cardiometabolic disease loci. *BMC Med.* **19**, 232 (2021).

75. Hagenaars, S. P. *et al.* Genetic prediction of male pattern baldness. *PLoS Genet.* **13**, e1006594 (2017).

76. Li, Y. *et al.* Genome-Wide Association Studies of Metabolites in Patients with CKD Identify Multiple Loci and Illuminate Tubular Transport Mechanisms. *J. Am. Soc. Nephrol.* **29**, 1513–1524 (2018).

77. Gutierrez-Achury, J. *et al.* Functional implications of disease-specific variants in loci jointly associated with coeliac disease and rheumatoid arthritis. *Hum. Mol. Genet.* **25**, 180–190 (2016).

78. Sun, D. *et al.* Multi-Ancestry Genome-wide Association Study Accounting for Gene-Psychosocial Factor Interactions Identifies Novel Loci for Blood Pressure Traits. *HGG Adv* **2**, (2021).

79. Levin, M. G. *et al.* Genome-wide association and multi-trait analyses characterize the common genetic architecture of heart failure. *Nat. Commun.* **13**, 6914 (2022).

80. Temprano-Sagrera, G. *et al.* Multi-phenotype analyses of hemostatic traits with cardiovascular events reveal novel genetic associations. *J. Thromb. Haemost.* **20**, 1331–1349 (2022).

81. Liu, Y. *et al.* Genetic architecture of 11 organ traits derived from abdominal MRI using deep learning. *Elife* **10**, (2021).

82. Hoffmann, T. J. *et al.* Genome-wide association analyses using electronic health records identify new loci influencing blood pressure variation. *Nat. Genet.* **49**, 54–64 (2017).

83. Astle, W. J. *et al.* The Allelic Landscape of Human Blood Cell Trait Variation and Links to Common Complex Disease. *Cell* **167**, 1415–1429.e19 (2016).

84. Pilling, L. C. *et al.* Red blood cell distribution width: Genetic evidence for aging pathways in 116,666 volunteers. *PLoS One* **12**, e0185083 (2017).

85. Kemp, J. P. *et al.* Identification of 153 new loci associated with heel bone mineral density and functional involvement of GPC6 in osteoporosis. *Nat. Genet.* **49**,

1468–1475 (2017).

86. Kim, S. K. Identification of 613 new loci associated with heel bone mineral density and a polygenic risk score for bone mineral density, osteoporosis and fracture. *PLoS One* **13**, e0200785 (2018).

87. Fernandez-Rozadilla, C. *et al.* Deciphering colorectal cancer genetics through multi-omic analysis of 100,204 cases and 154,587 controls of European and east Asian ancestries. *Nat. Genet.* **55**, 89–99 (2023).

88. Vuckovic, D. *et al.* The Polygenic and Monogenic Basis of Blood Traits and Diseases. *Cell* **182**, 1214–1231.e11 (2020).

89. Kichaev, G. *et al.* Leveraging Polygenic Functional Enrichment to Improve GWAS Power. *Am. J. Hum. Genet.* **104**, 65–75 (2019).

90. Estrada, K. *et al.* Genome-wide meta-analysis identifies 56 bone mineral density loci and reveals 14 loci associated with risk of fracture. *Nat. Genet.* **44**, 491–501 (2012).

91. Chen, M.-H. *et al.* Trans-ethnic and Ancestry-Specific Blood-Cell Genetics in 746,667 Individuals from 5 Global Populations. *Cell* **182**, 1198–1213.e14 (2020).

92. Medina-Gomez, C. *et al.* Life-Course Genome-wide Association Study Meta-analysis of Total Body BMD and Assessment of Age-Specific Effects. *Am. J. Hum. Genet.* **102**, 88–102 (2018).

93. Davies, G. *et al.* Study of 300,486 individuals identifies 148 independent genetic loci influencing general cognitive function. *Nat. Commun.* **9**, 2098 (2018).

94. Richardson, T. G. *et al.* Evaluating the relationship between circulating lipoprotein lipids and apolipoproteins with risk of coronary heart disease: A multivariable Mendelian randomisation analysis. *PLoS Med.* **17**, e1003062 (2020).

95. Hill, W. D. *et al.* A combined analysis of genetically correlated traits identifies 187 loci and a role for neurogenesis and myelination in intelligence. *Mol. Psychiatry* **24**,

169–181 (2019).

96. Márquez, A. *et al.* Meta-analysis of Immunochip data of four autoimmune diseases reveals novel single-disease and cross-phenotype associations. *Genome Med.* **10**, 97 (2018).

97. Sakaue, S. *et al.* A cross-population atlas of genetic associations for 220 human phenotypes. *Nat. Genet.* **53**, 1415–1424 (2021).

98. Pulit, S. L. *et al.* Meta-analysis of genome-wide association studies for body fat distribution in 694 649 individuals of European ancestry. *Hum. Mol. Genet.* **28**, 166–174 (2019).

99. Lee, J. J. *et al.* Gene discovery and polygenic prediction from a genome-wide association study of educational attainment in 1.1 million individuals. *Nat. Genet.* **50**, 1112–1121 (2018).

100. Okbay, A. *et al.* Genome-wide association study identifies 74 loci associated with educational attainment. *Nature* **533**, 539–542 (2016).

101. Rao, S., Baranova, A., Yao, Y., Wang, J. & Zhang, F. Genetic Relationships between Attention-Deficit/Hyperactivity Disorder, Autism Spectrum Disorder, and Intelligence. *Neuropsychobiology* **81**, 484–496 (2022).

102. Hill, W. D. *et al.* Genome-wide analysis identifies molecular systems and 149 genetic loci associated with income. *Nat. Commun.* **10**, 5741 (2019).

103. Savage, J. E. *et al.* Genome-wide association meta-analysis in 269,867 individuals identifies new genetic and functional links to intelligence. *Nat. Genet.* **50**, 912–919 (2018).

104. Li, M. *et al.* Genome-wide association study of 1,5-anhydroglucitol identifies novel genetic loci linked to glucose metabolism. *Sci. Rep.* **7**, 2812 (2017).

105. Cole, J. B., Florez, J. C. & Hirschhorn, J. N. Comprehensive genomic analysis of dietary habits in UK Biobank identifies hundreds of genetic associations. *Nat.*

*Commun.* **11**, 1467 (2020).

106. Lopera-Maya, E. A. *et al.* Effect of host genetics on the gut microbiome in 7,738 participants of the Dutch Microbiome Project. *Nat. Genet.* **54**, 143–151 (2022).

107. Tikkanen, E. *et al.* Biological Insights Into Muscular Strength: Genetic Findings in the UK Biobank. *Sci. Rep.* **8**, 6451 (2018).

108. Ward, J. *et al.* The genomic basis of mood instability: identification of 46 loci in 363,705 UK Biobank participants, genetic correlation with psychiatric disorders, and association with gene expression and function. *Mol. Psychiatry* **25**, 3091–3099 (2020).

109. Zhu, Z. *et al.* Shared genetic and experimental links between obesity-related traits and asthma subtypes in UK Biobank. *J. Allergy Clin. Immunol.* **145**, 537–549 (2020).

110. Zhu, Z. *et al.* A genome-wide cross-trait analysis from UK Biobank highlights the shared genetic architecture of asthma and allergic diseases. *Nat. Genet.* **50**, 857–864 (2018).

111. Kachuri, L. *et al.* Genetic determinants of blood-cell traits influence susceptibility to childhood acute lymphoblastic leukemia. *Am. J. Hum. Genet.* **108**, 1823–1835 (2021).

112. Richardson, T. G., Sanderson, E., Elsworth, B., Tilling, K. & Davey Smith, G. Use of genetic variation to separate the effects of early and later life adiposity on disease risk: mendelian randomisation study. *BMJ* **369**, m1203 (2020).

113. Yap, C. X. *et al.* Dissection of genetic variation and evidence for pleiotropy in male pattern baldness. *Nat. Commun.* **9**, 5407 (2018).

114. Lutz, S. M. *et al.* A genome-wide association study identifies risk loci for spirometric measures among smokers of European and African ancestry. *BMC Genet.* **16**, 138 (2015).

115. Helgeland, Ø. *et al.* Characterization of the genetic architecture of infant and

early childhood body mass index. *Nat Metab* **4**, 344–358 (2022).

116. Smith, S. M. *et al.* An expanded set of genome-wide association studies of brain imaging phenotypes in UK Biobank. *Nat. Neurosci.* **24**, 737–745 (2021).

117. Zheng, T. *et al.* Genome-wide analysis of 944 133 individuals provides insights into the etiology of haemorrhoidal disease. *Gut* **70**, 1538–1549 (2021).

118. Han, J. *et al.* A genome-wide association study identifies novel alleles associated with hair color and skin pigmentation. *PLoS Genet.* **4**, e1000074 (2008).

119. Cade, B. E. *et al.* Whole-genome association analyses of sleep-disordered breathing phenotypes in the NHLBI TOPMed program. *Genome Med.* **13**, 136 (2021).

120. Adhikari, K. *et al.* A GWAS in Latin Americans highlights the convergent evolution of lighter skin pigmentation in Eurasia. *Nat. Commun.* **10**, 358 (2019).

121. Adhikari, K. *et al.* A genome-wide association scan in admixed Latin Americans identifies loci influencing facial and scalp hair features. *Nat. Commun.* **7**, 10815 (2016).

122. Liu, F. *et al.* Genetics of skin color variation in Europeans: genome-wide association studies with functional follow-up. *Hum. Genet.* **134**, 823–835 (2015).

123. Galván-Femenía, I. *et al.* Multitrait genome association analysis identifies new susceptibility genes for human anthropometric variation in the GCAT cohort. *J. Med. Genet.* **55**, 765–778 (2018).

124. Lona-Durazo, F. *et al.* A large Canadian cohort provides insights into the genetic architecture of human hair colour. *Commun Biol* **4**, 1253 (2021).

125. Nan, H. *et al.* Genome-wide association study of tanning phenotype in a population of European ancestry. *J. Invest. Dermatol.* **129**, 2250–2257 (2009).

126. Liu, J. Z. *et al.* Association analyses identify 38 susceptibility loci for inflammatory bowel disease and highlight shared genetic risk across populations. *Nat. Genet.* **47**, 979–986 (2015).

127. Hellwege, J. N. *et al.* Mapping eGFR loci to the renal transcriptome and phenome in the VA Million Veteran Program. *Nat. Commun.* **10**, 3842 (2019).

128. Graham, S. E. *et al.* Sex-specific and pleiotropic effects underlying kidney function identified from GWAS meta-analysis. *Nat. Commun.* **10**, 1847 (2019).

129. Lotta, L. A. *et al.* A cross-platform approach identifies genetic regulators of human metabolism and health. *Nat. Genet.* **53**, 54–64 (2021).

130. Kettunen, J. *et al.* Genome-wide study for circulating metabolites identifies 62 loci and reveals novel systemic effects of LPA. *Nat. Commun.* **7**, 11122 (2016).

131. Houlihan, L. M. *et al.* Common variants of large effect in F12, KNG1, and HRG are associated with activated partial thromboplastin time. *Am. J. Hum. Genet.* **86**, 626–631 (2010).

132. Benyamin, B. *et al.* Identification of novel loci affecting circulating chromogranins and related peptides. *Hum. Mol. Genet.* **26**, 233–242 (2017).

133. Verweij, N. *et al.* Genome-wide association study on plasma levels of midregional-proadrenomedullin and C-terminal-pro-endothelin-1. *Hypertension* **61**, 602–608 (2013).

134. Moksnes, M. R. *et al.* Genome-wide association study of cardiac troponin I in the general population. *Hum. Mol. Genet.* **30**, 2027–2039 (2021).

135. Inouye, M. *et al.* Novel Loci for metabolic networks and multi-tissue expression studies reveal genes for atherosclerosis. *PLoS Genet.* **8**, e1002907 (2012).

136. Yet, I. *et al.* Genetic Influences on Metabolite Levels: A Comparison across Metabolomic Platforms. *PLoS One* **11**, e0153672 (2016).

137. Shin, S.-Y. *et al.* An atlas of genetic influences on human blood metabolites. *Nat. Genet.* **46**, 543–550 (2014).

138. Timoteo, V. J., Chiang, K.-M., Yang, H.-C. & Pan, W.-H. Common and ethnic-specific genetic determinants of hemoglobin concentration between Taiwanese Han

Chinese and European Whites: findings from comparative two-stage genome-wide association studies. *J. Nutr. Biochem.* **111**, 109126 (2023).

139. Wei, J. *et al.* Identification of fifty-seven novel loci for abdominal wall hernia development and their biological and clinical implications: results from the UK Biobank. *Hernia* **26**, 335–348 (2022).

140. Bentham, J. *et al.* Genetic association analyses implicate aberrant regulation of innate and adaptive immunity genes in the pathogenesis of systemic lupus erythematosus. *Nat. Genet.* **47**, 1457–1464 (2015).

141. Tomer, Y. *et al.* Genome wide identification of new genes and pathways in patients with both autoimmune thyroiditis and type 1 diabetes. *J. Autoimmun.* **60**, 32–39 (2015).

142. Tian, C. *et al.* Genome-wide association and HLA region fine-mapping studies identify susceptibility loci for multiple common infections. *Nat. Commun.* **8**, 599 (2017).

143. Wu, Y. *et al.* Multi-trait analysis for genome-wide association study of five psychiatric disorders. *Transl. Psychiatry* **10**, 209 (2020).

144. Wang, H., Yi, Z. & Shi, T. Novel loci and potential mechanisms of major depressive disorder, bipolar disorder, and schizophrenia. *Sci. China Life Sci.* **65**, 167–183 (2022).

145. Autism Spectrum Disorders Working Group of The Psychiatric Genomics Consortium. Meta-analysis of GWAS of over 16,000 individuals with autism spectrum disorder highlights a novel locus at 10q24.32 and a significant overlap with schizophrenia. *Mol. Autism* **8**, 21 (2017).

146. Howard, D. M. *et al.* Genome-wide association study of depression phenotypes in UK Biobank identifies variants in excitatory synaptic pathways. *Nat. Commun.* **9**, 1470 (2018).

147. Cai, N. *et al.* Minimal phenotyping yields genome-wide association signals of

low specificity for major depression. *Nat. Genet.* **52**, 437–447 (2020).

148. Génin, E. *et al.* Genome-wide association study of Stevens-Johnson Syndrome and Toxic Epidermal Necrolysis in Europe. *Orphanet J. Rare Dis.* **6**, 52 (2011).

149. Hill, W. D. *et al.* Genetic contributions to two special factors of neuroticism are associated with affluence, higher intelligence, better health, and longer life. *Mol. Psychiatry* **25**, 3034–3052 (2020).

150. Luciano, M. *et al.* Association analysis in over 329,000 individuals identifies 116 independent variants influencing neuroticism. *Nat. Genet.* **50**, 6–11 (2018).

151. Wu, Y. *et al.* Genome-wide association study of medication-use and associated disease in the UK Biobank. *Nat. Commun.* **10**, 1891 (2019).

152. Feitosa, M. F. *et al.* Novel genetic associations for blood pressure identified via gene-alcohol interaction in up to 570K individuals across multiple ancestries. *PLoS One* **13**, e0198166 (2018).

153. Borges, M. C. *et al.* Role of circulating polyunsaturated fatty acids on cardiovascular diseases risk: analysis using Mendelian randomization and fatty acid genetic association data from over 114,000 UK Biobank participants. *BMC Med.* **20**, 210 (2022).

154. Folkersen, L. *et al.* Mapping of 79 loci for 83 plasma protein biomarkers in cardiovascular disease. *PLoS Genet.* **13**, e1006706 (2017).

155. Lieb, W. *et al.* Genome-wide association study for endothelial growth factors. *Circ. Cardiovasc. Genet.* **8**, 389–397 (2015).

156. Weiss, F. U. *et al.* Fucosyltransferase 2 (FUT2) non-secretor status and blood group B are associated with elevated serum lipase activity in asymptomatic subjects, and an increased risk for chronic pancreatitis: a genetic association study. *Gut* **64**, 646–656 (2015).

157. Ruth, K. S. *et al.* Using human genetics to understand the disease impacts of testosterone in men and women. *Nat. Med.* **26**, 252–258 (2020).

158. Gao, X. R., Huang, H., Nannini, D. R., Fan, F. & Kim, H. Genome-wide association analyses identify new loci influencing intraocular pressure. *Hum. Mol. Genet.* **27**, 2205–2213 (2018).

159. Khawaja, A. P. *et al.* Genome-wide analyses identify 68 new loci associated with intraocular pressure and improve risk prediction for primary open-angle glaucoma. *Nat. Genet.* **50**, 778–782 (2018).

160. Lindström, S. *et al.* Genomic and transcriptomic association studies identify 16 novel susceptibility loci for venous thromboembolism. *Blood* **134**, 1645–1657 (2019).

161. Hysi, P. G. *et al.* Metabolome Genome-Wide Association Study Identifies 74 Novel Genomic Regions Influencing Plasma Metabolites Levels. *Metabolites* **12**, (2022).

162. Nielsen, J. B. *et al.* Loss-of-function genomic variants highlight potential therapeutic targets for cardiovascular disease. *Nat. Commun.* **11**, 6417 (2020).

163. Ikeda, M. *et al.* A genome-wide association study identifies two novel susceptibility loci and trans population polygenicity associated with bipolar disorder. *Mol. Psychiatry* **23**, 639–647 (2018).

164. Rhee, E. P. *et al.* A genome-wide association study of the human metabolome in a community-based cohort. *Cell Metab.* **18**, 130–143 (2013).

165. Draisma, H. H. M. *et al.* Genome-wide association study identifies novel genetic variants contributing to variation in blood metabolite levels. *Nat. Commun.* **6**, 7208 (2015).

166. Gallois, A. *et al.* A comprehensive study of metabolite genetics reveals strong pleiotropy and heterogeneity across time and context. *Nat. Commun.* **10**, 4788 (2019).

167. Mozaffarian, D. *et al.* Genetic loci associated with circulating phospholipid trans fatty acids: a meta-analysis of genome-wide association studies from the

CHARGE Consortium. *Am. J. Clin. Nutr.* **101**, 398–406 (2015).

168. Li-Gao, R. *et al.* Genetic Studies of Metabolomics Change After a Liquid Meal Illuminate Novel Pathways for Glucose and Lipid Metabolism. *Diabetes* **70**, 2932–2946 (2021).

169. Hu, Y. *et al.* Discovery and fine-mapping of loci associated with MUFAs through trans-ethnic meta-analysis in Chinese and European populations. *J. Lipid Res.* **58**, 974–981 (2017).

170. Tabassum, R. *et al.* Genetic architecture of human plasma lipidome and its link to cardiovascular disease. *Nat. Commun.* **10**, 4329 (2019).

171. Dorajoo, R. *et al.* A genome-wide association study of n-3 and n-6 plasma fatty acids in a Singaporean Chinese population. *Genes Nutr.* **10**, 53 (2015).

172. Folkersen, L. *et al.* Genomic and drug target evaluation of 90 cardiovascular proteins in 30,931 individuals. *Nat Metab* **2**, 1135–1148 (2020).

173. Ligthart, S. *et al.* Bivariate genome-wide association study identifies novel pleiotropic loci for lipids and inflammation. *BMC Genomics* **17**, 443 (2016).

174. Teslovich, T. M. *et al.* Biological, clinical and population relevance of 95 loci for blood lipids. *Nature* **466**, 707–713 (2010).

175. Willer, C. J. *et al.* Discovery and refinement of loci associated with lipid levels. *Nat. Genet.* **45**, 1274–1283 (2013).

176. Bentley, A. R. *et al.* Multi-ancestry genome-wide gene-smoking interaction study of 387,272 individuals identifies new loci associated with serum lipids. *Nat. Genet.* **51**, 636–648 (2019).

177. Hoffmann, T. J. *et al.* A large electronic-health-record-based genome-wide study of serum lipids. *Nat. Genet.* **50**, 401–413 (2018).

178. Bell, S. *et al.* A genome-wide meta-analysis yields 46 new loci associating with biomarkers of iron homeostasis. *Commun Biol* **4**, 156 (2021).

179. Wojcik, G. L. *et al.* Genetic analyses of diverse populations improves discovery for complex traits. *Nature* **570**, 514–518 (2019).

180. van Setten, J. *et al.* Genome-wide association meta-analysis of 30,000 samples identifies seven novel loci for quantitative ECG traits. *Eur. J. Hum. Genet.* **27**, 952–962 (2019).

181. Jiao, H., Zhang, M., Zhang, Y., Wang, Y. & Li, W.-D. Pathway Association Studies Reveal Gene Loci and Pathway Networks that Associated With Plasma Cystatin C Levels. *Front. Genet.* **12**, 711155 (2021).

182. de Vries, P. S. *et al.* Comparison of HapMap and 1000 Genomes Reference Panels in a Large-Scale Genome-Wide Association Study. *PLoS One* **12**, e0167742 (2017).

183. Sinnott-Armstrong, N. *et al.* Genetics of 35 blood and urine biomarkers in the UK Biobank. *Nat. Genet.* **53**, 185–194 (2021).

184. Han, Y. *et al.* Genome-wide analysis highlights contribution of immune system pathways to the genetic architecture of asthma. *Nat. Commun.* **11**, 1776 (2020).

185. Baranova, A., Cao, H., Chen, J. & Zhang, F. Causal Association and Shared Genetics Between Asthma and COVID-19. *Front. Immunol.* **13**, 705379 (2022).

186. Wain, L. V. *et al.* Genome-wide association study identifies six new loci influencing pulse pressure and mean arterial pressure. *Nat. Genet.* **43**, 1005–1011 (2011).

187. van der Meer, D. *et al.* Boosting Schizophrenia Genetics by Utilizing Genetic Overlap With Brain Morphology. *Biol. Psychiatry* **92**, 291–298 (2022).

188. Naqvi, S. *et al.* Shared heritability of human face and brain shape. *Nat. Genet.* **53**, 830–839 (2021).

189. Orrù, V. *et al.* Complex genetic signatures in immune cells underlie autoimmunity and inform therapy. *Nat. Genet.* **52**, 1036–1045 (2020).

190. Dubois, P. C. A. *et al.* Multiple common variants for celiac disease influencing immune gene expression. *Nat. Genet.* **42**, 295–302 (2010).

191. Zhernakova, A. *et al.* Meta-analysis of genome-wide association studies in celiac disease and rheumatoid arthritis identifies fourteen non-HLA shared loci. *PLoS Genet.* **7**, e1002004 (2011).

192. Köttgen, A. *et al.* New loci associated with kidney function and chronic kidney disease. *Nat. Genet.* **42**, 376–384 (2010).

193. Shadrin, A. A. *et al.* Vertex-wise multivariate genome-wide association study identifies 780 unique genetic loci associated with cortical morphology. *Neuroimage* **244**, 118603 (2021).

194. Newton-Cheh, C. *et al.* Genome-wide association study identifies eight loci associated with blood pressure. *Nat. Genet.* **41**, 666–676 (2009).

195. Kato, N. *et al.* Trans-ancestry genome-wide association study identifies 12 genetic loci influencing blood pressure and implicates a role for DNA methylation. *Nat. Genet.* **47**, 1282–1293 (2015).

196. Plotnikov, D. *et al.* High Blood Pressure and Intraocular Pressure: A Mendelian Randomization Study. *Invest. Ophthalmol. Vis. Sci.* **63**, 29 (2022).

197. Johansson, Å., Rask-Andersen, M., Karlsson, T. & Ek, W. E. Genome-wide association analysis of 350 000 Caucasians from the UK Biobank identifies novel loci for asthma, hay fever and eczema. *Hum. Mol. Genet.* **28**, 4022–4041 (2019).

198. de Lange, K. M. *et al.* Genome-wide association study implicates immune activation of multiple integrin genes in inflammatory bowel disease. *Nat. Genet.* **49**, 256–261 (2017).

199. Klarin, D. *et al.* Genetics of blood lipids among ~300,000 multi-ethnic participants of the Million Veteran Program. *Nat. Genet.* **50**, 1514–1523 (2018).

200. Pasman, J. A. *et al.* Genetic Risk for Smoking: Disentangling Interplay

Between Genes and Socioeconomic Status. *Behav. Genet.* **52**, 92–107 (2022).

201. Graham, S. E. *et al.* The power of genetic diversity in genome-wide association studies of lipids. *Nature* **600**, 675–679 (2021).

202. Liu, C. *et al.* Meta-analysis identifies common and rare variants influencing blood pressure and overlapping with metabolic trait loci. *Nat. Genet.* **48**, 1162–1170 (2016).

203. Nikpay, M. *et al.* A comprehensive 1,000 Genomes-based genome-wide association meta-analysis of coronary artery disease. *Nat. Genet.* **47**, 1121–1130 (2015).

204. Patrick, M. T. *et al.* Causal Relationship and Shared Genetic Loci between Psoriasis and Type 2 Diabetes through Trans-Disease Meta-Analysis. *J. Invest. Dermatol.* **141**, 1493–1502 (2021).

205. Aung, N. *et al.* Genome-wide association analysis reveals insights into the genetic architecture of right ventricular structure and function. *Nat. Genet.* **54**, 783–791 (2022).

206. Fischer, A. *et al.* Identification of Immune-Relevant Factors Conferring Sarcoidosis Genetic Risk. *Am. J. Respir. Crit. Care Med.* **192**, 727–736 (2015).

207. Langefeld, C. D. *et al.* Transancestral mapping and genetic load in systemic lupus erythematosus. *Nat. Commun.* **8**, 16021 (2017).

208. Medici, M. *et al.* Identification of novel genetic Loci associated with thyroid peroxidase antibodies and clinical thyroid disease. *PLoS Genet.* **10**, e1004123 (2014).

209. Selvaraj, M. S. *et al.* Genome-wide discovery for diabetes-dependent triglycerides-associated loci. *PLoS One* **17**, e0275934 (2022).

210. Onengut-Gumuscu, S. *et al.* Fine mapping of type 1 diabetes susceptibility loci and evidence for colocalization of causal variants with lymphoid gene enhancers. *Nat. Genet.* **47**, 381–386 (2015).

211. Köttgen, A. *et al.* Genome-wide association analyses identify 18 new loci associated with serum urate concentrations. *Nat. Genet.* **45**, 145–154 (2013).

212. Gill, D. *et al.* Urate, Blood Pressure, and Cardiovascular Disease: Evidence From Mendelian Randomization and Meta-Analysis of Clinical Trials. *Hypertension* **77**, 383–392 (2021).

213. van der Meer, D. *et al.* The genetic architecture of human cortical folding. *Sci Adv* **7**, eabj9446 (2021).

214. Fan, C. C. *et al.* Multivariate genome-wide association study on tissue-sensitive diffusion metrics highlights pathways that shape the human brain. *Nat. Commun.* **13**, 2423 (2022).

215. Soranzo, N. *et al.* A genome-wide meta-analysis identifies 22 loci associated with eight hematological parameters in the HaemGen consortium. *Nat. Genet.* **41**, 1182–1190 (2009).

216. Klarin, D. *et al.* Genome-wide association study of peripheral artery disease in the Million Veteran Program. *Nat. Med.* **25**, 1274–1279 (2019).

217. van Zuydam, N. R. *et al.* Genome-Wide Association Study of Peripheral Artery Disease. *Circ Genom Precis Med* **14**, e002862 (2021).

218. Acosta-Herrera, M. *et al.* Genome-wide meta-analysis reveals shared new loci in systemic seropositive rheumatic diseases. *Ann. Rheum. Dis.* **78**, 311–319 (2019).

219. Coffee and Caffeine Genetics Consortium *et al.* Genome-wide meta-analysis identifies six novel loci associated with habitual coffee consumption. *Mol. Psychiatry* **20**, 647–656 (2015).

220. Kennedy, O. J. *et al.* Coffee Consumption and Kidney Function: A Mendelian Randomization Study. *Am. J. Kidney Dis.* **75**, 753–761 (2020).

221. Wuttke, M. *et al.* A catalog of genetic loci associated with kidney function from analyses of a million individuals. *Nat. Genet.* **51**, 957–972 (2019).

222. Liu, M. *et al.* Association studies of up to 1.2 million individuals yield new insights into the genetic etiology of tobacco and alcohol use. *Nat. Genet.* **51**, 237–244 (2019).

223. Zhou, H. *et al.* Genome-wide meta-analysis of problematic alcohol use in 435,563 individuals yields insights into biology and relationships with other traits. *Nat. Neurosci.* **23**, 809–818 (2020).

224. Sulem, P. *et al.* Sequence variants at CYP1A1-CYP1A2 and AHR associate with coffee consumption. *Hum. Mol. Genet.* **20**, 2071–2077 (2011).

225. Zhong, V. W. *et al.* A genome-wide association study of bitter and sweet beverage consumption. *Hum. Mol. Genet.* **28**, 2449–2457 (2019).

226. Stanzick, K. J. *et al.* Discovery and prioritization of variants and genes for kidney function in >1.2 million individuals. *Nat. Commun.* **12**, 4350 (2021).

227. Liu, H. *et al.* Epigenomic and transcriptomic analyses define core cell types, genes and targetable mechanisms for kidney disease. *Nat. Genet.* **54**, 950–962 (2022).

228. Pardiñas, A. F. *et al.* Pharmacogenomic Variants and Drug Interactions Identified Through the Genetic Analysis of Clozapine Metabolism. *Am. J. Psychiatry* **176**, 477–486 (2019).

229. Pardiñas, A. F. *et al.* Pharmacokinetics and pharmacogenomics of clozapine in an ancestrally diverse sample: a longitudinal analysis and genome-wide association study using UK clinical monitoring data. *Lancet Psychiatry* **10**, 209–219 (2023).

230. Pirastu, N. *et al.* Using genetic variation to disentangle the complex relationship between food intake and health outcomes. *PLoS Genet.* **18**, e1010162 (2022).

231. Tin, A. *et al.* Target genes, variants, tissues and transcriptional pathways influencing human serum urate levels. *Nat. Genet.* **51**, 1459–1474 (2019).

232. Zanetti, D. *et al.* Identification of 22 novel loci associated with urinary

biomarkers of albumin, sodium, and potassium excretion. *Kidney Int.* **95**, 1197–1208 (2019).

233. Casanova, F. *et al.* A genome-wide association study implicates multiple mechanisms influencing raised urinary albumin-creatinine ratio. *Hum. Mol. Genet.* **28**, 4197–4207 (2019).

234. Pazoki, R. *et al.* GWAS for urinary sodium and potassium excretion highlights pathways shared with cardiovascular traits. *Nat. Commun.* **10**, 3653 (2019).

235. Huang, J. *et al.* Genomics and phenomics of body mass index reveals a complex disease network. *Nat. Commun.* **13**, 7973 (2022).

236. Said, M. A., van de Vegte, Y. J., Verweij, N. & van der Harst, P. Associations of Observational and Genetically Determined Caffeine Intake With Coronary Artery Disease and Diabetes Mellitus. *J. Am. Heart Assoc.* **9**, e016808 (2020).

237. Cornelis, M. C. *et al.* Genome-wide association study of caffeine metabolites provides new insights to caffeine metabolism and dietary caffeine-consumption behavior. *Hum. Mol. Genet.* **25**, 5472–5482 (2016).

238. Smith, R. L. *et al.* Identification of a novel polymorphism associated with reduced clozapine concentration in schizophrenia patients-a genome-wide association study adjusting for smoking habits. *Transl. Psychiatry* **10**, 198 (2020).

239. Karlsson, T. *et al.* Contribution of genetics to visceral adiposity and its relation to cardiovascular and metabolic disease. *Nat. Med.* **25**, 1390–1395 (2019).

240. Meddens, S. F. W. *et al.* Genomic analysis of diet composition finds novel loci and associations with health and lifestyle. *Mol. Psychiatry* **26**, 2056–2069 (2021).

241. Haas, M. E. *et al.* Genetic Association of Albuminuria with Cardiometabolic Disease and Blood Pressure. *Am. J. Hum. Genet.* **103**, 461–473 (2018).

242. Amin, N. *et al.* Genome-wide association analysis of coffee drinking suggests association with CYP1A1/CYP1A2 and NRCAM. *Mol. Psychiatry* **17**, 1116–1129 (2012).

243. Cornelis, M. C. *et al.* Genome-wide meta-analysis identifies regions on 7p21 (AHR) and 15q24 (CYP1A2) as determinants of habitual caffeine consumption. *PLoS Genet.* **7**, e1002033 (2011).

244. Teumer, A. *et al.* Genome-wide association meta-analyses and fine-mapping elucidate pathways influencing albuminuria. *Nat. Commun.* **10**, 4130 (2019).

245. Vollenbrock, C. E., Roshandel, D., van der Klauw, M. M., Wolffenbuttel, B. H. R. & Paterson, A. D. Genome-wide association study identifies novel loci associated with skin autofluorescence in individuals without diabetes. *BMC Genomics* **23**, 840 (2022).

246. Levy, D. *et al.* Genome-wide association study of blood pressure and hypertension. *Nat. Genet.* **41**, 677–687 (2009).

247. Sung, Y. J. *et al.* A Large-Scale Multi-ancestry Genome-wide Study Accounting for Smoking Behavior Identifies Multiple Significant Loci for Blood Pressure. *Am. J. Hum. Genet.* **102**, 375–400 (2018).

248. International Consortium for Blood Pressure Genome-Wide Association Studies *et al.* Genetic variants in novel pathways influence blood pressure and cardiovascular disease risk. *Nature* **478**, 103–109 (2011).

249. Surendran, P. *et al.* Trans-ancestry meta-analyses identify rare and common variants associated with blood pressure and hypertension. *Nat. Genet.* **48**, 1151–1161 (2016).

250. Wain, L. V. *et al.* Novel Blood Pressure Locus and Gene Discovery Using Genome-Wide Association Study and Expression Data Sets From Blood and the Kidney. *Hypertension* **70**, e4–e19 (2017).

251. Giri, A. *et al.* Trans-ethnic association study of blood pressure determinants in over 750,000 individuals. *Nat. Genet.* **51**, 51–62 (2019).

252. Siewert, K. M. & Voight, B. F. Bivariate Genome-Wide Association Scan

Identifies 6 Novel Loci Associated With Lipid Levels and Coronary Artery Disease. *Circ Genom Precis Med* **11**, e002239 (2018).

253. López-Isac, E. *et al.* GWAS for systemic sclerosis identifies multiple risk loci and highlights fibrotic and vasculopathy pathways. *Nat. Commun.* **10**, 4955 (2019).

254. Benjamins, J. W. *et al.* Genomic insights in ascending aortic size and distensibility. *EBioMedicine* **75**, 103783 (2022).

255. Francis, C. M. *et al.* Genome-wide associations of aortic distensibility suggest causality for aortic aneurysms and brain white matter hyperintensities. *Nat. Commun.* **13**, 4505 (2022).

256. Tcheandjieu, C. *et al.* High heritability of ascending aortic diameter and trans-ancestry prediction of thoracic aortic disease. *Nat. Genet.* **54**, 772–782 (2022).

257. Pirruccello, J. P. *et al.* Deep learning enables genetic analysis of the human thoracic aorta. *Nat. Genet.* **54**, 40–51 (2022).

258. Hartiala, J. A. *et al.* Genome-wide analysis identifies novel susceptibility loci for myocardial infarction. *Eur. Heart J.* **42**, 919–933 (2021).

259. Liu, D. J. *et al.* Exome-wide association study of plasma lipids in >300,000 individuals. *Nat. Genet.* **49**, 1758–1766 (2017).

260. Nagel, M., Watanabe, K., Stringer, S., Posthuma, D. & van der Sluis, S. Item-level analyses reveal genetic heterogeneity in neuroticism. *Nat. Commun.* **9**, 905 (2018).

261. Cárcel-Márquez, J. *et al.* A Polygenic Risk Score Based on a Cardioembolic Stroke Multitrait Analysis Improves a Clinical Prediction Model for This Stroke Subtype. *Front Cardiovasc Med* **9**, 940696 (2022).

262. Roselli, C. *et al.* Multi-ethnic genome-wide association study for atrial fibrillation. *Nat. Genet.* **50**, 1225–1233 (2018).

263. Jansen, P. R. *et al.* Genome-wide analysis of insomnia in 1,331,010

individuals identifies new risk loci and functional pathways. *Nat. Genet.* **51**, 394–403 (2019).

264. Lam, M. *et al.* Comparative genetic architectures of schizophrenia in East Asian and European populations. *Nat. Genet.* **51**, 1670–1678 (2019).

265. Williams, C. M., Labouret, G., Wolfram, T., Peyre, H. & Ramus, F. A General Cognitive Ability Factor for the UK Biobank. *Behav. Genet.* **53**, 85–100 (2023).

266. Demange, P. A. *et al.* Investigating the genetic architecture of noncognitive skills using GWAS-by-subtraction. *Nat. Genet.* **53**, 35–44 (2021).

267. Trubetskoy, V. *et al.* Mapping genomic loci implicates genes and synaptic biology in schizophrenia. *Nature* **604**, 502–508 (2022).

268. Goes, F. S. *et al.* Genome-wide association study of schizophrenia in Ashkenazi Jews. *Am. J. Med. Genet. B Neuropsychiatr. Genet.* **168**, 649–659 (2015).

269. Michailidou, K. *et al.* Association analysis identifies 65 new breast cancer risk loci. *Nature* **551**, 92–94 (2017).

270. Warren, H. R. *et al.* Genome-wide association analysis identifies novel blood pressure loci and offers biological insights into cardiovascular risk. *Nat. Genet.* **49**, 403–415 (2017).

271. Mallick, S. *et al.* The Allen Ancient DNA Resource (AADR): A curated compendium of ancient human genomes. *bioRxiv* 2023.04.06.535797 (2023) doi:10.1101/2023.04.06.535797.

272. Le, M. K. *et al.* 1,000 ancient genomes uncover 10,000 years of natural selection in Europe. *bioRxiv* (2022) doi:10.1101/2022.08.24.505188.

273. Corder, E. H. *et al.* Gene dose of apolipoprotein E type 4 allele and the risk of Alzheimer’s disease in late onset families. *Science* **261**, 921–923 (1993).

274. Strittmatter, W. J. *et al.* Apolipoprotein E: high-avidity binding to beta-amyloid and increased frequency of type 4 allele in late-onset familial Alzheimer disease. *Proc.*

*Natl. Acad. Sci. U. S. A.* **90**, 1977–1981 (1993).

275. Voight, B. F., Kudaravalli, S., Wen, X. & Pritchard, J. K. A map of recent positive selection in the human genome. *PLoS Biol.* **4**, e72 (2006).

276. Sabeti, P. C. *et al.* Genome-wide detection and characterization of positive selection in human populations. *Nature* **449**, 913–918 (2007).

277. Pickrell, J. K. *et al.* Signals of recent positive selection in a worldwide sample of human populations. *Genome Res.* **19**, 826–837 (2009).

278. Wang, L. *et al.* A MicroRNA Linking Human Positive Selection and Metabolic Disorders. *Cell* **183**, 684–701.e14 (2020).

279. Buckley, M. T. *et al.* Selection in Europeans on Fatty Acid Desaturases Associated with Dietary Changes. *Mol. Biol. Evol.* **34**, 1307–1318 (2017).

280. Ye, K., Gao, F., Wang, D., Bar-Yosef, O. & Keinan, A. Dietary adaptation of FADS genes in Europe varied across time and geography. *Nat Ecol Evol* **1**, 167 (2017).

281. Mathieson, S. & Mathieson, I. FADS1 and the Timing of Human Adaptation to Agriculture. *Mol. Biol. Evol.* **35**, 2957–2970 (2018).

282. Kurki, M. I. *et al.* FinnGen provides genetic insights from a well-phenotyped isolated population. *Nature* **613**, 508–518 (2023).

283. Global Biobank Engine. http://gbe.stanford.edu.

284. McInnes, G. *et al.* Global Biobank Engine: enabling genotype-phenotype browsing for biobank summary statistics. *Bioinformatics* **35**, 2495–2497 (2019).

285. Ju, D. & Mathieson, I. The evolution of skin pigmentation-associated variation in West Eurasia. *Proc. Natl. Acad. Sci. U. S. A.* **118**, (2021).

286. Jablonski, N. G. & Chaplin, G. The evolution of human skin coloration. *J. Hum. Evol.* **39**, 57–106 (2000).

287. Engelsen, O. The relationship between ultraviolet radiation exposure and vitamin D status. *Nutrients* **2**, 482–495 (2010).

288. Stefansson, H. *et al.* A common inversion under selection in Europeans. *Nat. Genet.* **37**, 129–137 (2005).

289. Steinberg, K. M. *et al.* Structural diversity and African origin of the 17q21.31 inversion polymorphism. *Nat. Genet.* **44**, 872–880 (2012).

290. Andreadis, A., Brown, W. M. & Kosik, K. S. Structure and novel exons of the human tau gene. *Biochemistry* **31**, 10626–10633 (1992).

291. Alonso, A. C., Grundke-Iqbal, I. & Iqbal, K. Alzheimer’s disease hyperphosphorylated tau sequesters normal tau into tangles of filaments and disassembles microtubules. *Nat. Med.* **2**, 783–787 (1996).

292. Desikan, R. S. *et al.* Genetic overlap between Alzheimer’s disease and Parkinson's disease at the MAPT locus. *Mol. Psychiatry* **20**, 1588–1595 (2015).

293. Satake, W. *et al.* Genome-wide association study identifies common variants at four loci as genetic risk factors for Parkinson’s disease. *Nat. Genet.* **41**, 1303–1307 (2009).

294. Simón-Sánchez, J. *et al.* Genome-wide association study reveals genetic risk underlying Parkinson’s disease. *Nat. Genet.* **41**, 1308–1312 (2009).

295. Shulman, J. M. & De Jager, P. L. Evidence for a common pathway linking neurodegenerative diseases. *Nature genetics* vol. 41 1261–1262 (2009).

296. Kerner, G. *et al.* Genetic adaptation to pathogens and increased risk of inflammatory disorders in post-Neolithic Europe. *Cell Genom* **3**, 100248 (2023).

297. Rohland, N. *et al.* Three Reagents for in-Solution Enrichment of Ancient Human DNA at More than a Million SNPs. *bioRxiv* 2022.01.13.476259 (2022) doi:10.1101/2022.01.13.476259.

298. Davidson, R. *et al.* Allelic bias when performing in-solution enrichment of ancient human DNA. *bioRxiv* 2023.07.04.547445 (2023) doi:10.1101/2023.07.04.547445.

### 5) Validating CLUES on Ancient Genotypes

Andrew H. Vaughn[1], Rasmus Nielsen[2,3]

[1] Center for Computational Biology, University of California Berkeley, Berkeley, USA
[2] Lundbeck Foundation GeoGenetics Centre, GLOBE Institute, University of Copenhagen, Copenhagen, Denmark
[3] Department of Integrative Biology, University of California, Berkeley, USA

#### Introduction

We here validate the usage of CLUES on ancient genotype samples through several benchmarking simulations. We simulate data according to a basic discrete-time Wright-Fisher model. We draw the initial frequency of the derived allele, $x_0$, from Unif(0.05, 0.2). Given $x_k$, $x_{k+1}$ is $A_{k+1}/N$ where $A_{k+1}$ is drawn from Bin(N, $\frac{x_k + sx_k}{1 + sx_k}$). This process is repeated for 500 generations, and every m generations, one individual is sampled and their genotype recorded. We use the list of sampled individuals as input to CLUES via the `--ancientSamps` option.

Specifically, we run CLUES as:

```
python inference.py --popFreq x500 --ancientSamps Samples.txt --N N
```

where $x_{500}$ is the allele frequency at generation 500 of our simulation, Samples.txt is our set of sampled individuals, and N is the population size we used in our simulations. These simulations have three parameters, s, N, and m.

#### Validation of $\chi^2_1$ Test Statistic

We begin by validating the appropriateness of the $\chi^2_1$ distribution for our tests of selection. We run 250 simulations with s = 0, Ne = 30000, and m = 20 and run CLUES on the output of each of them to infer $\hat{s}^{MLE}$. By Wilks' Theorem, 2 log (L($\hat{s}^{MLE}$) / L(s = 0)) should be distributed as $\chi^2_1$ if s is indeed 0. We plot a P-P plot of our empirical distribution of twice the log likelihood ratio to the $\chi^2_1$ distribution. For comparison, we also plot a P-P plot of the results of 250 simulations with s = 0.005, Ne = 30000, and m = 20.

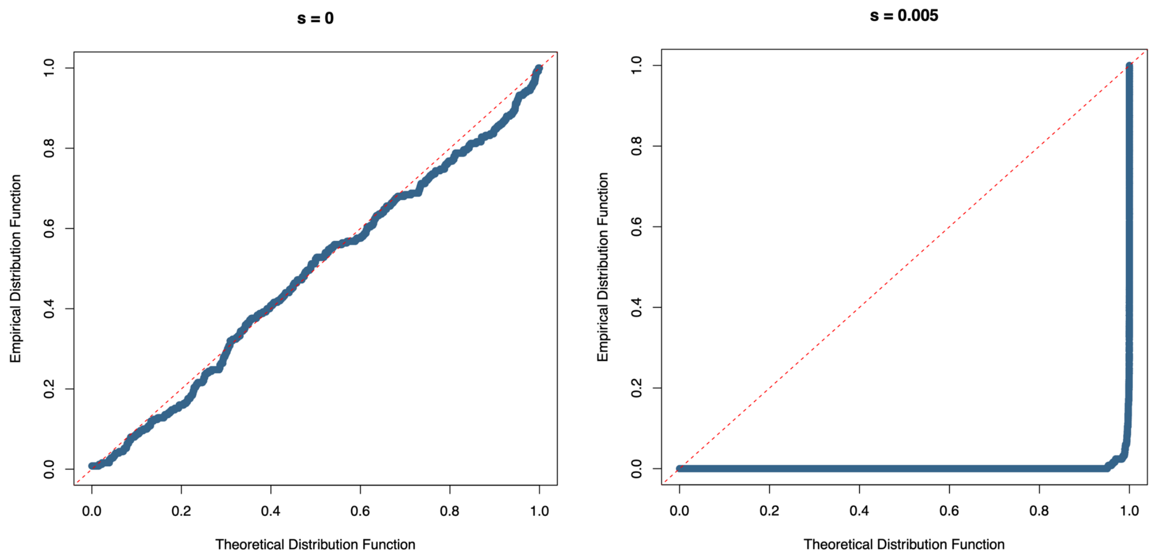


**Figure S86.** Here, we plot P-P plots for our simulations where s = 0 (left) and s = 0.005 (right). A P-P plot compares two distributions by comparing their cumulative distribution functions. We compute the empirical CDF of our data as F(x) = $\frac{1}{n}\sum_{i=1}^{n} 1_{\{X_i \leq x\}}$ and denote the true CDF of the $\chi^2_1$ distribution as G(x). We plot F(x) against G(x) as x ranges from 0 to ∞.

We see that for the simulations run with s = 0, the empirical distribution largely follows a $\chi^2_1$ distribution, while those run with s = 0.005 show an excess of large log likelihood ratios compared to the $\chi^2_1$ distribution. Therefore, we consider the use of the $\chi^2_1$ distribution to obtain p-values for our tests of selection to be justified.

#### Estimation of Selection Coefficients

We also test the accuracy of the estimation of selection coefficients. We plot the distribution of selection coefficients for the simulations described in the previous section with s = 0 and s = 0.005.

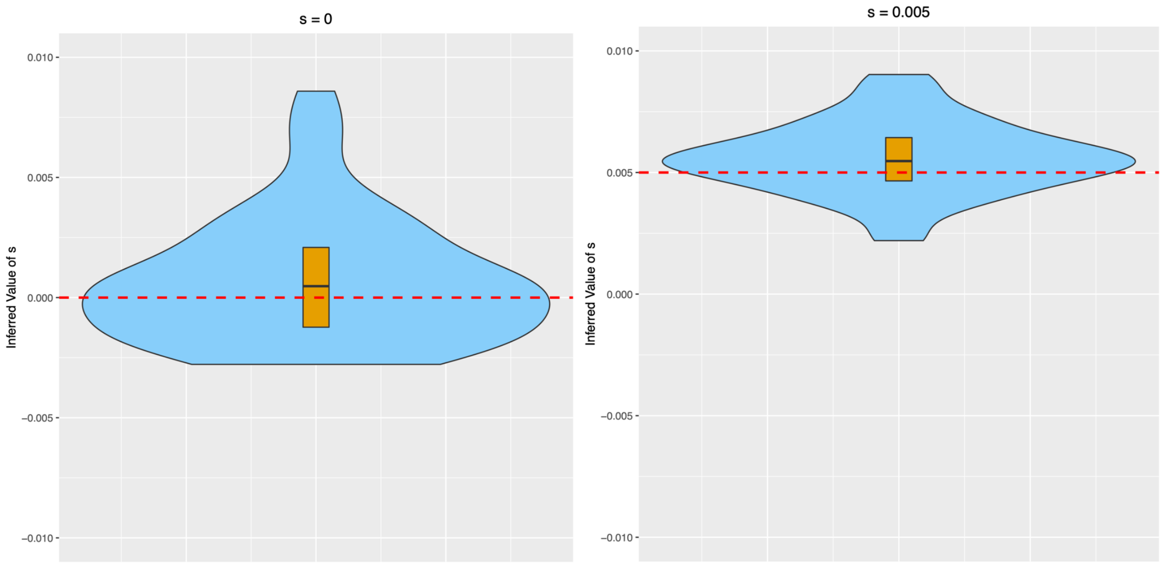


**Figure S87.** Here, we plot violin plots of selection coefficients for our simulations where s = 0 (left) and s = 0.005 (right) for n=250 replicates each. The violin plots are truncated at the extrema of the datasets. Boxplots are overlaid, with the whiskers omitted, showing the median and the first and third quartiles. We plot the true value of s as a dashed horizontal line. Two outliers are omitted for the left plot.

We therefore conclude that the estimation of selection coefficients is quite accurate. If CLUES performs precisely as it should, there will be exactly two sources of error in the estimation of selection coefficients, both contributing to variance rather than bias:

1. Random fluctuations in the true allele frequency caused by genetic drift (which will cause the true allele frequency to be less informative of the true selection coefficient)
2. Noisy estimation of the true allele frequency caused by finite sampling of historic genotypes

We show this explicitly with another round of simulations.

#### Effect of Demography and Sampling Density on Selection Coefficient Estimation

For comparison with our previous results for s = 0, we run two additional sets of 50 simulations with s = 0. One simulation has Ne = 30000 and m = 4, while the other has Ne =

200000 and m = 1. We show the results of both new sets of simulations together with our original simulation, which we call the reference simulation.

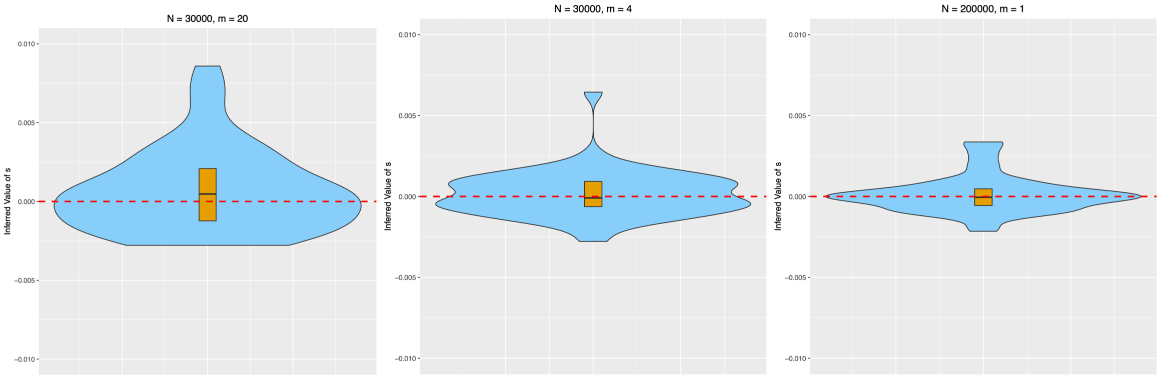


**Figure S88.** Here, we plot violin plots of selection coefficients for our reference simulation (left), our simulation where Ne = 30000 and m = 4 (middle) and our simulation where Ne = 200000 and m = 1 (right), for n=50 replicates each. As before, the violin plots are truncated at the extrema of the datasets. Boxplots are overlaid, with the whiskers omitted, showing the median and the first and third quartiles. We plot the true value of s as a dashed horizontal line. Two outliers are omitted for the left plot.

We do the same thing for the simulations where s = 0.005.

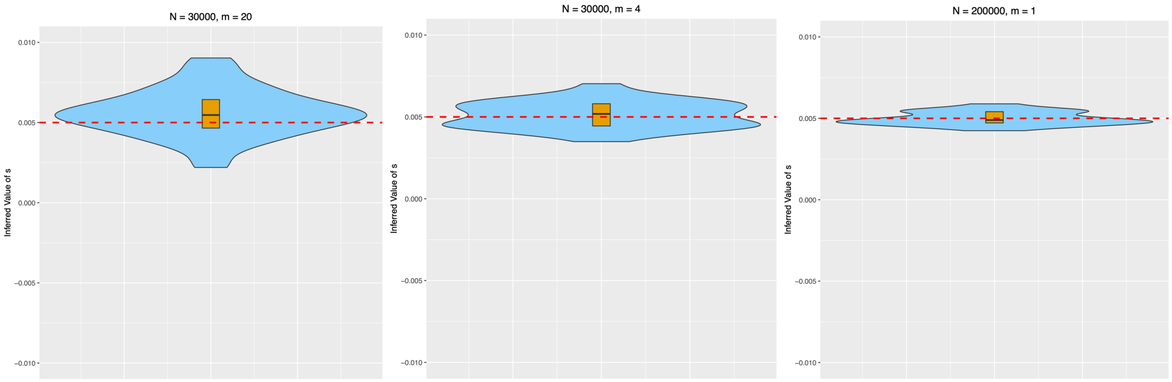


**Figure S89.** Here, we plot violin plots of selection coefficients for our reference simulation (left), our simulation where Ne = 30000 and m = 4 (middle) and our simulation where Ne = 200000 and m = 1 (right), for n=50 replicates each. As before, the violin plots are truncated at the extrema of the datasets. Boxplots are overlaid, with the whiskers omitted, showing the median and the first and third quartiles. We plot the true value of s as a dashed horizontal line.

We therefore conclude that with less genetic drift and denser sampling, CLUES converges to the true value of the selection coefficient.

#### Estimation of Allele Frequencies

Another important functionality of CLUES is to reconstruct historic allele frequencies. We run 3 different simulations to illustrate this functionality. Our first simulation is run with Ne = 30000, s = 0, and m = 2. Our second simulation is run with Ne = 30000, s = 0.005, and m = 2. Our third simulation is run with Ne = 1000, s = 0.005, and m = 2. For each simulation, we record the true historic allele frequency at each generation. We then use the posterior estimates of allele frequencies from CLUES to reconstruct the MAP allele frequency trajectory. We also plot, for each generation, a 95% posterior interval.

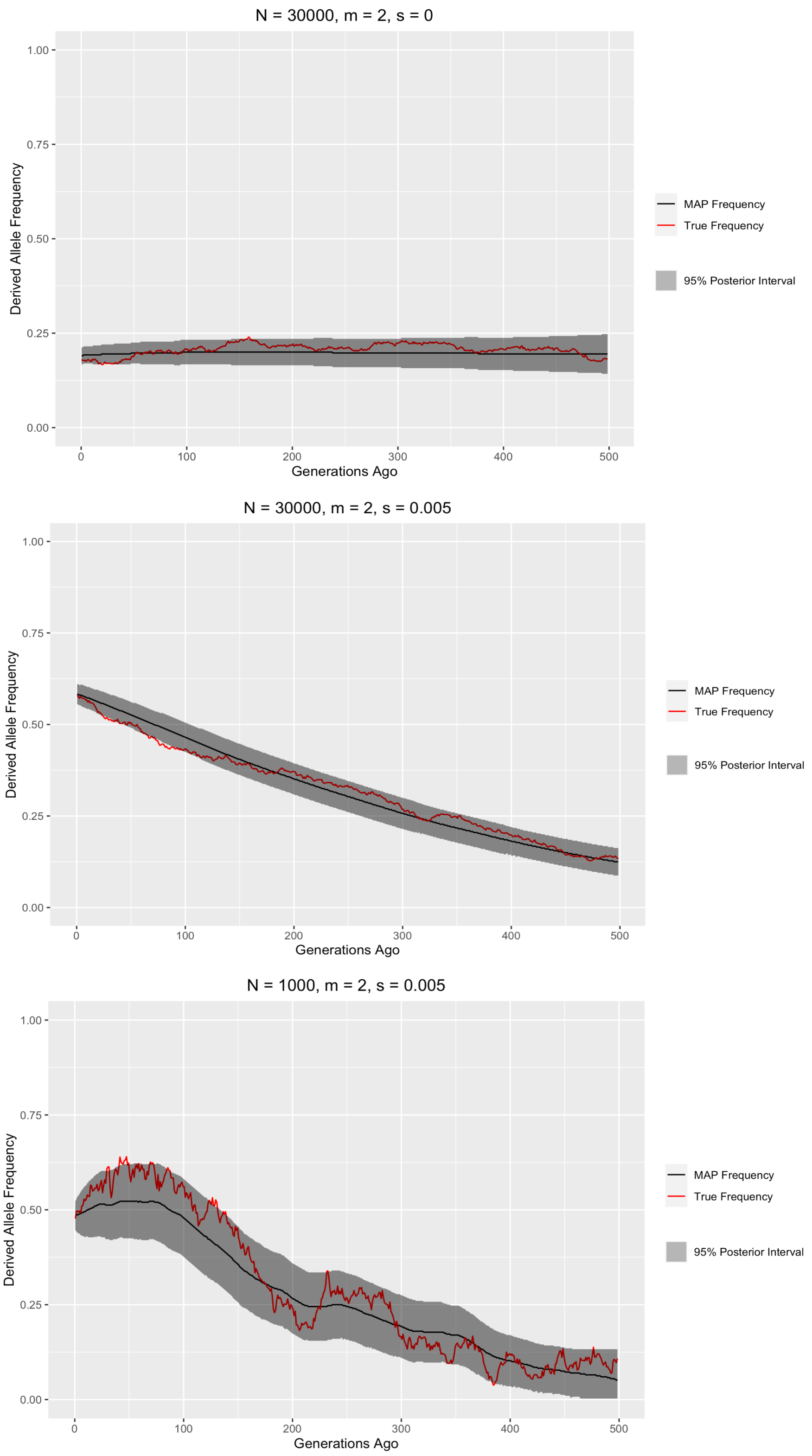


**Figure S90.** Here, we plot allele frequency trajectories and posterior intervals for a set of 3 informative simulations. Error bands show the 95% posterior interval of the model fit.

We see that for the 2 simulations with Ne = 30000 (top and middle), the true allele frequency trajectory has very little deviation from the expected allele trajectory with no drift. We find that CLUES accurately models this certainty, and the 95% posterior interval largely covers the true allele frequency. With our simulation with Ne = 1000 (bottom), there is obviously more drift in the true allele frequency trajectory, and there are significant increases and decreases in the frequency. We see that CLUES recognizes the higher uncertainty inherent to more genetic drift by having larger 95% posterior intervals. Furthermore, we observe that the posterior intervals move up and down roughly in accordance with the major true allele fluctuations. Therefore, we conclude that CLUES accurately estimates historic allele frequencies and correctly models the uncertainty in those estimates.

### 6) Detangling Direct and Indirect impacts of sample age from the Mesolithic-Neolithic data on genotype imputation

Rasmus Amund Henriksen[1], Rasmus Nielsen[1,2], Lei Zhao[1], Thorfinn Sand Korneliussen[1]

[1] Lundbeck Foundation GeoGenetics Centre, GLOBE Institute, University of Copenhagen, Copenhagen, Denmark

[2] Department of Integrative Biology, University of California, Berkeley

#### Introduction

Many factors can influence the genotype imputation/calling at specific sites, especially when present-day reference panels are used to impute missing genotypes for ancient samples. The unique features of aDNA relative to modern DNA include 1) relatively shallower read depths [1], 2) shorter read lengths [2], and 3) different error profiles [3]. Mapping biases that depend on read length and error rates are of particular concern for aDNA and may cause spurious signals of selection. Naturally, read depth may also affect genotype calling. In order to filter SNPs that might be affected by such biases, and to generally correct for and quantify the biases, we develop a causal inference method for distinguishing direct effects of sample age on allele frequency from other (indirect) effects mediated by age-dependent errors, read depth, and read length.

#### Methods

As shown in Figure S91, to distinguish the true selection signal of age on allele frequency, from the signal caused by mapping biases, and other biases, in the ancient samples, we create a model and a workflow which can decompose the influence of sample age ($A_i$, age of individual *i*) on genotype ($G_{ij}$, Genotype of individual *i* at position *j*), into its indirect and direct effects. The imputed genotype for each individual is converted into allele frequencies, representing the homozygous for the reference allele, heterozygous and homozygous for the alternative allele as 0, 0.5 and 1 respectively. The indirect effects are mediated through the three unique features of aDNA as previously described, while the direct impact reflects the true change in allele frequency over the time range from the ancient samples to the modern ones. The three factors are the mean read depth across all sites per individual, the mean read length for each individual at each site, and the third is an overall error estimate for each individual. These three factors are all identified using ANGSD [4] from the aligned BAM files for all individuals used in the study. The depth and length are calculated based solely on the imputed positions (*j)* extracted from the imputed vcf files, whereas the overall error is calculated as described in Orlando et al. 2013 from the entire BAM file containing all the invariant sites.

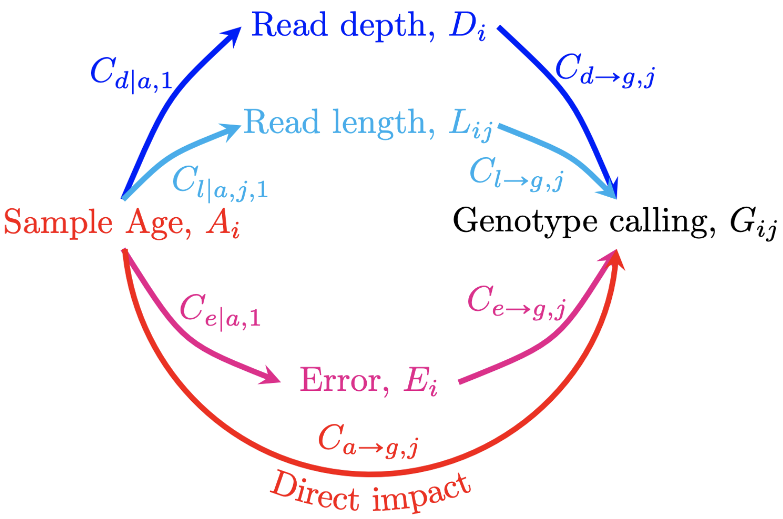


**Figure S91.** Illustrative figure of the factors that influence genotype calling procedure. The denotations with symbol "|" can be obtained by regression, while the denotations with symbol "→" are derived after PCA.

The initial step is to conduct three linear regressions, one for each of the three factors, mean depth (equation 1), mean length (equation 2) and error estimate (equation 3) with respect to one explanatory variable, i.e., sample age (i.e., $A_i$). This step is to investigate the influences of sample age on each of the three factors.

$$D_i \triangleq \overline{D_{ij}} \sim C_{d|a,1}A_i + C_{d|a,0} + \epsilon_{d|a,i} \qquad (1)$$

$$L_{ij} \triangleq \overline{L_{ijk}} \sim C_{l|a,j,1}A_i + C_{l|a,j,0} + \epsilon_{l|a,ij} \qquad (2)$$

$$E_i \sim C_{e|a,1}A_i + C_{e|a,0} + \epsilon_{e|a,i}. \qquad (3)$$

Where $\triangleq$ means definition, $\overline{D_{ij}}$ means the average depth with regard to *j* for a specific individual *i*, and $\overline{L_{ijk}}$ means the average read length, for all reads *k* stretching over site *j* of individual *i*. $E_i$means the overall error rate of individual *i*.

We assume that the sample age influences these factors, as such it is necessary to eliminate this influence as done in equation 4-6.

$$\Delta_i \triangleq D_i - C_{d|a,1}A_i \quad (4)$$

$$\Lambda_{ij} \triangleq L_{ij} - C_{l|a,j,1}A_i \quad (5)$$

$$\Sigma_i \triangleq E_i - C_{e|a,1}A_i \quad (6)$$

A second round of regression is necessary to detect the direct impact of the three factors as well as the direct impact of age on the genotype. The explanatory variables can be the sample age ($A_i$) and the three remainders (or the linear combinations of the three remainders). To avoid potential correlations between the three remainders, we perform a Principal Component Analysis. The relationship between the PCA scores and the remainders can be represented as in equation 7 and 8.

$$\left(\frac{\Lambda_{ij}-\Lambda_j}{\mathrm{sd}(\Lambda_{.j})}, \frac{\Sigma_i-\Sigma}{\mathrm{sd}(\Sigma.)}, \frac{\Delta_i-\Delta}{\mathrm{sd}(\Delta.)}\right)_{n\times 3} = (\alpha_{ij}\,\beta_{ij},\gamma_{ij})_{n\times 3}\begin{pmatrix}\overrightarrow{\omega}'_{j,1}\\ \overrightarrow{\omega}'_{j,2}\\ \overrightarrow{\omega}'_{j,3}\end{pmatrix} \quad (7)$$

$$(\alpha_{ij}\,\beta_{ij},\gamma_{ij})_{n\times 3} = \left(\frac{\Lambda_{ij}-\Lambda_j}{\mathrm{sd}(\Lambda)}, \frac{\Sigma_i-\Sigma}{\mathrm{sd}(\Sigma.)}, \frac{\Delta_i-\Delta}{\mathrm{sd}(\Delta.)}\right)_{n\times 3}(\overrightarrow{\omega}_{j,1}, \overrightarrow{\omega}_{j,2}, \overrightarrow{\omega}_{j,3}) \quad (8)$$

Where $\Lambda_j$and sd($\Lambda_{.j}$) are the mean and the standard deviation of $\Lambda$ across individuals *i* at fixed position *j*. $\Sigma$ and sd($\Sigma_.$) are the mean and the standard deviation of $\Sigma$ across individuals *i*. $\Delta$and sd($\Delta_.$) are the mean and the standard deviation of $\Delta$across individuals *i*. $\overrightarrow{\omega}_{j,2}$, $\overrightarrow{\omega}_{j,1}$, $\overrightarrow{\omega}_{j,3}$ are the three principle directions (column eigenvectors) and the $\overrightarrow{\omega}'_{j,2}$, $\overrightarrow{\omega}'_{j,1}$, $\overrightarrow{\omega}'_{j,3}$ are the corresponding transposed vectors. $\alpha_{ij}, \beta_{ij}, \gamma_{ij}$ are the principal component scores of $\left(\frac{\Lambda_{ij}-\Lambda_j}{sd(\Lambda_{.j})}, \frac{\Sigma_i-\Sigma}{sd(\Sigma_.)}, \frac{\Delta_i-\Delta}{sd(\Delta_.)}\right)$, with *n* being the number of individuals carrying site *j*.

Once all the explanatory variables are independent, the coefficient $C_{g|a,j,1}$represent the total impact of age which can be obtained by conducting a final round of regression (equation 9).

$$G_{ij} \sim C_{g|\alpha,j,1}\alpha_{ij} + C_{g|\beta,j,1}\beta_{ij} + C_{g|\gamma,j,1}\gamma_{ij} + C_{g|a,j,1}A_i + C_{0,j} + \epsilon_{ij} \quad (9)$$

The sum of the first three terms of equation 9 can be obtained by multiplying the coefficients $(C_{g|\alpha,j,1}, C_{g|\beta,j,1}, C_{g|\gamma,j,1})'$ with equation 8 (shown in equation 10). And the effective regression coefficients for $(\Delta_{ij}, \Lambda_i, \Sigma_i)$ will be observed when calculating the linear slopes of the remainders in Equation 10. Such effective coefficients can be viewed as measurements of the direct impact of the corresponding factors, i.e. length ($C_{l\to g,j}$), error ($C_{e\to g,j}$) and depth ($C_{d\to g,j}$), on the genotype calling (equation 11 - 13).

$$(\alpha_{ij}\,\beta_{ij},\gamma_{ij})_{n\times 3}\begin{pmatrix} C_{g|\alpha,j,1} \\ C_{g|\beta,j,1} \\ C_{g|\gamma,j,1} \end{pmatrix} = \begin{pmatrix} \frac{\Lambda_{ij}-\Lambda_j}{\mathrm{sd}(\Lambda_{\cdot j})}, \frac{\Sigma_i-\Sigma}{\mathrm{sd}(\Sigma.)}, \frac{\Delta_i-\Delta}{\mathrm{sd}(\Delta.)} \end{pmatrix}_{n\times 3} (\vec{\omega}_{j,1}, \vec{\omega}_{j,2}, \vec{\omega}_{j,3})_{3\times 3} \begin{pmatrix} C_{g|\alpha,j,1} \\ C_{g|\beta,j,1} \\ C_{g|\gamma,j,1} \end{pmatrix} \quad (10)$$

$$\vec{\omega}_{j,1} = \begin{pmatrix} \omega_{j,1,1} \\ \omega_{j,2,1} \\ \omega_{j,3,1} \end{pmatrix}, \vec{\omega}_{j,2} = \begin{pmatrix} \omega_{j,1,2} \\ \omega_{j,2,2} \\ \omega_{j,3,2} \end{pmatrix}, \vec{\omega}_{j,3} = \begin{pmatrix} \omega_{j,1,3} \\ \omega_{j,2,3} \\ \omega_{j,3,3} \end{pmatrix}$$

$$\mathrm{C}_{l\to g,j} = \frac{1}{\mathrm{sd}(\Lambda_{\cdot j})}\left(\omega_{j,1,1}C_{g|\alpha,j,1} + \omega_{j,1,2}C_{g|\beta,j,1} + \omega_{j,1,3}C_{g|\gamma,j,1}\right) \quad (11)$$

$$\mathrm{C}_{e\to g,j} = \frac{1}{\mathrm{sd}(\Sigma.)}\left(\omega_{j,2,1}C_{g|\alpha,j,1} + \omega_{j,2,2}C_{g|\beta,j,1} + \omega_{j,2,3}C_{g|\gamma,j,1}\right) \quad (12)$$

$$\mathrm{C}_{d\to g,j} = \frac{1}{\mathrm{sd}(\Delta.)}\left(\omega_{j,3,1}C_{g|\alpha,j,1} + \omega_{j,3,2}C_{g|\beta,j,1} + \omega_{j,3,3}C_{g|\gamma,j,1}\right) \quad (13)$$

Any influence of age on the genotype calling imposed through one of the three factors (depth, length, error) is an indirect impact. The measurements of such indirect impacts are calculated by multiplying equations 11-13 with each of the factors corresponding coefficients obtained from our first round of regression (equation 4-6) for each site *j*.

$$\mathrm{C}_{a\to l\to g,j} = C_{l|a,j,1}\cdot \mathrm{C}_{l\to g,j} \quad (14)$$

$$\mathrm{C}_{a\to e\to g,j} = C_{e|a,1}\cdot \mathrm{C}_{e\to g,j} \quad (15)$$

$$\mathrm{C}_{a\to d\to g,j} = C_{d|a,1}\cdot \mathrm{C}_{d\to g,j} \quad (16)$$

Finally, the direct impact of age on the genotype calling at site *j* (equation 17) can be obtained by subtracting all indirect impacts of age (equation 14-16) from the total impact of age $C_{g|a,j,1}$, obtained from the second round of regression (equation 9).

$$C_{a\to g,j} = C_{g|a,j,1} - \mathrm{C}_{a\to l\to g,j} - \mathrm{C}_{a\to e\to g,j} - \mathrm{C}_{a\to d\to g,j} \quad (17)$$

Then to apply all these measurements, we calculate a ratio of the sum of all possible indirect effects and the direct effect for each site (i.e., $R_j$, equation 18). This ratio can be used to quantify the relative contribution of indirect and direct effects on changes in allele frequency.

If the ratio is negative or close to zero, an observed change in allele frequency cannot solely be attributed to biases, and selection at such sites was likely to occur during the course of evolution. While a relatively large positive ratio indicates that the potential selection is indistinguishable with the effects of mapping biases and/or other biases, thus we will filter out such sites while detecting selection signals.

$$R_j = \frac{C_{a\to l\to g,j} + C_{a\to e\to g,j} + C_{a\to d\to g,j}}{C_{a\to g,j}} \qquad (18)$$

$R_j$ can also be converted into a fraction representing the proportion of indirect effects on age relative to the total effect (equation 19), which can equivalently be used for filtering:

$$F_j = \frac{R_j}{1+R_j} = \frac{1}{1+\frac{1}{R_j}} \qquad (19)$$

To validate the proposed regression method, we applied it to simulated datasets. The simulation was conducted based on a simplified model, where the read lengths ($L$) depend on the age of the sample, and the estimated allele frequencies ($G$) depend on both Age and read lengths, so in this simplified model, read length is the only ancient feature considered:

$L = Age_i + \epsilon$

and

$G = C_a Age + C_l L + \eta$

In the simulation, $\epsilon$ and $\eta$ are both normal random variables with mean 0 and standard deviation 100. The coefficient $C_a$ represents the true direct impact of age $C_{a\to g}$ to estimated allele frequencies, which is assumed to be true selection signal, while $C_l$ reflects how the ancient feature read length will affect the estimated allele frequencies, and given $L$, we have the true indirect impact of age given as

$C_{a\to l\to g} = C_l \times 1 = C_l$

Hence the true $F_j$ value of this simulated data will follow

$$F_j = \frac{C_l}{C_a + C_l}$$

We applied our regression method to the datasets simulated following the above description with six different $C_a$ and $C_l$ settings. Case 1: the indirect impact and the direct impact are of the same direction, but the indirect impact contributes more ($C_a = 0.4, C_l = 1$), Case 2: The indirect impact and the direct impact are of the same direction, but the direct impact contributes more ($C_a = -1, C_l = -0.4$), Case 3: The indirect impact and the direct impact are of the same direction, but both impacts contribute equally ($C_a = 0.4, C_l = 0.4$), Case 4: The indirect impact and the direct impact are of different directions ($C_a =$

$0.4, C_l = -1)$, Case 5: The indirect impact and the direct impact are of different directions $(C_a = 1, C_l = -0.4)$ and Case 6 without direct impact $(C_a = 0, C_l = 1)$. In each case, 10000 replicates of simulated datasets containing 5001 samples with the age ranging from 5000 to 10000 are generated, and our method will estimate a $F_j$ for each simulated dataset.

In Figure S92, we present a histogram for the estimated $F_j$ values and its corresponding true value for each case with different parameter settings.

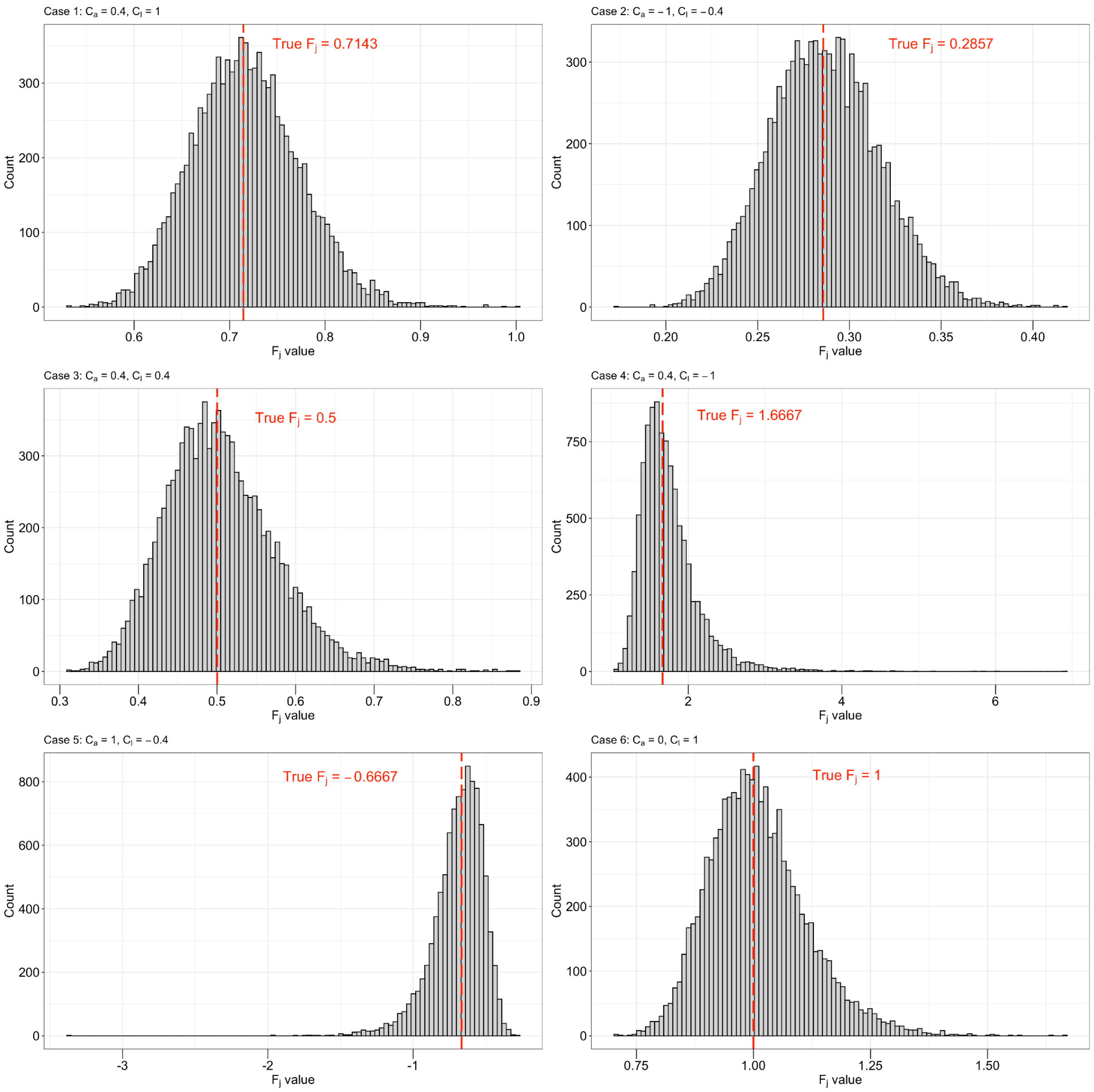


**Figure S92.** The estimated $F_j$ values across six different cases simulating different scenarios of the indirect- and the direct impact.

#### Discussion

The distribution of $F_j$ can be divided into 3 parts, the region $F_j < 0$, maps back to $-1 < R_j < 0$, the region $0 < F_j \leq 1$maps back to $R_j > 0$, and the region $F_j > 1$ maps back to $R_j < -1$. When filtering sites we exclude those sites where the indirect and direct impacts have the same direction, making it impossible to differentiate between the two, i.e., $R_j > 0$. Therefore when the filtering is based on $F_j$ values, we only filter the sites with $F_j$ values within $0 < F_j \leq 1$. When $F_j$ the value is closer to 1, it becomes harder to distinguish the confounding signal from the true temporal signal (e.g., selection), so we use a cutoff c to remove the SNPs where it is unworkable to differentiate them. The value of $F_j$ is used to keep the sites which we believe to be under (relatively large) selection and screen the rest, while the numerator and denominator of $R_j$, as they reflect the absolute strengths of the temporal and confounding signals, are more suitable for detecting unknown selection signals upon sites. To filter out sites in selection analyses that may be affected by biases, we use a fixed threshold of $0.5 < F_j \leq 1$. Choosing a filtering threshold of $F_j$ values above empirical threshold 0.5 removes those sites where indirect effects mediated through the ancient DNA characteristics (biases caused by ancient characteristics) are greater than the true change of allele frequency measured as the direct effect. Their change of frequencies are more likely caused by ancient signals rather than selections.

Figure S92 shows in each of the case, our estimates of $F_j$ are always distributed around their true values, and confirm that $F_j \in (-\infty, 0) \cup (1, +\infty)$ represents the cases that the indirect impact and the direct impact are of different directions; $F_j \in (0{,}0.5)$ represents the case that the indirect impact and the direct impact are of the same direction, but the direct impact contributes more; $F_j \in (0.5{,}1)$ represents the case that the indirect impact and the direct impact are of the same direction, but the indirect impact contributes more. Hence, though empirically set, $F_j \in (0.5{,}1)$ is reasonable evidence that the change of allele frequencies can be mainly due to indirect impacts of age rather than its direct impacts, i.e., true selection signals. Although the simulated model is simplified and does not regard allele frequencies per individual as discrete values within [0,1], which is not biological accurate, we believe the simulations can still validate the basic idea of the proposed regression method and illustrate the logics of setting Fj threshold 0.5.

#### References

1. Shapiro, B. & Hofreiter, M. A paleogenomic perspective on evolution and gene function: new insights from ancient DNA. *Science* **343**, 1236573 (2014).

2. Paabo, S. Ancient DNA: extraction, characterization, molecular cloning, and enzymatic amplification. *Proceedings of the National Academy of Sciences* vol. 86 1939–1943 (1989).

3. Orlando, L. *et al.* Recalibrating Equus evolution using the genome sequence of an early Middle Pleistocene horse. *Nature* **499**, 74–78 (2013).

4. Korneliussen, T. S., Albrechtsen, A. & Nielsen, R. ANGSD: Analysis of Next Generation Sequencing Data. *BMC Bioinformatics* **15**, 356 (2014).

### 7) Identifying candidates for positive selection using patterns of ancient population differentiation

Alba Refoyo Martínez[1], Fernando Racimo[1]

[1] Lundbeck Foundation GeoGenetics Centre, GLOBE Institute, University of Copenhagen, Copenhagen, Denmark

#### Introduction

We aimed to detect whether there is evidence of positive selection in the past 15,000 years by searching for loci with strong differentiation in allele frequencies both between populations and across time.

#### Methods

We worked with the imputed dataset. We applied several variant-level filters:

1. We only used SNPs with MAF > 5%
2. Genotype missingness rate < 50%; and
3. Variants where <10% individuals had post-imputation genotype probability (GP) <= 0.8

We used pcadapt [1] - a method for detection of allele frequency outliers based on principal component analysis - to search for loci that might have evidence for positive selection in the past. After performing a PCA on the ancient genomic data, we used a scree plot to visualise the percentage of variance explained by each PC.

We performed two scans to look for candidates under positive selection. First, we performed a “eurasian” scan, in which we used the first three principal components of a PCA of all ancient genomes (higher components explained less than 1% of the total variance in allele frequencies) (Figures S93, S94 and S95). We also performed a second scan of selection restricting only to ancient West Eurasian individuals (excluding ancient Siberian populations) and we also used the first three components of the PCA (Figures S96, S97 and S98). We call this the “west-eur” scan. We also performed a third scan, which we call “hg-neo” scan, in which we only tested for significant loadings corresponding to the component that separates hunter-gatherers (WHG+EHG) from Neolithic farming peoples in the aforementioned

Western Eurasian PCA (first principal component). The latter scan should serve to find loci with particularly strong allele frequencies between these groups with two distinct modes of subsistence. Figure S94 and S97 shows Manhattan plots for each of the three scans.

We selected the top-scoring 300 SNPs with the lowest p-values and merged them into candidate regions if they were within 100 kb of each other. Each region was labelled with the P-value of its highest scoring SNP, which was then used to rank regions. HGNC protein-coding genes within each region were retrieved using *biomaRt* [2]. Supplementary Table 9 lists the 25 top-scoring candidate regions from the “eurasian” scan, while Supplementary Table 10 and Supplementary Table 11 list the top candidate regions from the “west-eur” and “hg-neo” scan, respectively.

None of our candidate regions seem to be affected by mapping biases. The Fj value for highest scoring SNP in each region is lower than 0.5 (Supplementary Note 6).

#### Results

##### Eurasian scan

In the Eurasian-wide scan (results in Supplementary Table 9), the strongest peak contains the gene SLC24A5, involved in skin and eye pigmentation, and previously reported to be under positive selection in Western Eurasia [3–6] (Figure S94). We also recover the region encompassing the *EDAR* gene which has been implicated in numerous studies of positive selection involving East Asian populations [7–9]. Variants in this gene are associated with numerous ectoderm-related traits, including hair thickness and ectodermal dysplasia [10] and tooth morphology [11].

We also found a candidate peak (chr2:25874547-26568094), containing several genes (ASXL2, RAB10, HADHA, GPR113) involved in glucose homeostasis mainly in response to a high fat diet. The ASXL2 gene regulates skeletal, glucose, and adipocyte homeostasis. It promotes adipogenesis, the formation of osteoclasts and insulin resistance [12–14]. A similar activity is carried out by RAB10, it also regulates glucose homeostasis by improving hyperglycemia, regulating at the skeletal muscle level [15–17]. HADHA participates in long fatty acid oxidation for energy production in different tissues [18–20]. Finally, GPR113 participates in energy expenditure in fat tissue and glucose homeostasis. It improves insulin sensitivity and prevents obesity when it binds to bile acids [21]. High allele frequencies in East Asian populations (Panel B of Figure S99).

We found four peaks containing genes associated with cardiovascular disorders and obesity. One of these (chr20:18991679-19454079) overlaps with the SLC24A3 gene. This is a salt sensitivity gene which is significantly expressed in obese individuals [22–24] and it is associated with hypertension [25]. We also find a peak in an intergenic region (chr3:123433220-123889576) containing ROPN1 and KALRN, two genes involved in vascular disorders [26–28]. The alternative allele at the top SNP in this region is at particularly high frequency in ancient Steppe populations. Another candidate region (chr1:234142067-234549596) contains SLC35F3, which codes for a thiamine transport and has been associated with hypertension in a Han Chinese cohort [29,30]. In the same region, we also find COA6, which has high expression in cardiac pathologies [31]. The alternative allele frequency at the top SNP in the region is high in East Asian populations (Panel B of Figure S99). Finally, the region (chr10:90592757 - 91009553) contains several genes (CH25H, FAS) associated with obesity and lipid metabolism [32–34], and immune responses [35].

One of the top candidate regions (chr11:131077365-131516733) contains a gene - NTM - involved in neuropsychiatric disorders [25,36], while another region (chr11:44634764 - 45073989) contains a gene associated with schizophrenia, TSPAN18. It has previously shown that the schizophrenia-risk SNPs within this region are highly diverged between Europeans and East Asians [37]. In chromosome 7 (chr7:95947959-96347959), SLC25A13, which is highly associated with citrin deficiency in East Asian populations [38,39]. High alternative allele frequencies in East Asians (Panel A of Figure S99) [40,41].

##### West Eurasian scan

In the scan restricting to ancient populations in Western Eurasia (“west-eur”) (top hits in Supplementary Table 10), we recover three regions which are involved in skin, hair and eye pigmentation, and have been previously implicated in differences in these traits across present-day Eurasians: two in chromosome 15 containing the genes *SLC24A5/MYEF2/CTXN2 and OCA2/HERC2* respectively and one in chromosome 5, containing gene *SLC45A2* [3,42–45] (Supplementary Table 10). Alternative allele frequencies shown in Figure S100.

The region containing *LCT/MCM6* - responsible for lactase persistence in Europe - is also a candidate region in the selection scan [46–48]. We also recover the TLR-1-6-10 gene cluster, which is known to be a target of selection in Europe and is associated with the immune response [46–48] (Panel A, Figure S100).

Additionally, we find some new potentially important candidate regions for positive selection. The region showing the strongest evidence of selection is located in chromosome 6 (chr6:134192815-134628278), around the SLC2A12 gene, which codes for a glucose transporter that participates in glucose homeostasis [49,50]. Variants in this gene are associated with mean corpuscular levels, heart diseases and height. The alternative allele of the highest-scoring SNP was at high frequency in hunter-gatherers but at much lower frequencies in Neolithic farmer populations and other, more recent, populations (Panel B, Figure S100).

Another novel candidate region overlaps with the VAMP 5-8 gene cluster in chromosome 2:85369379-85885211 a region associated with cardiovascular diseases [51,52]. In chr9:27009422-27434948, we found a peak overlapping the TEK gene, which codes for a tyrosine kinase receptor expressed in endothelial cells. This gene has an important role in angiogenesis and cardiovascular development and stability, and it is involved in several vascular disorders [53–55]. Recent studies have also investigated its role in asthma and allergic conjunctivitis [53–56]. Intermediate allele frequencies in Eastern hunter-gatherers in both regions (Panel B, Figure S100). Region chr1:227020437 - 227877723 contains CDC42BPA, also known as MRCKα, an important gene involved in iron utilisation [57] is involved in the erythropoiesis regulation [58]. The alternative allele at the top-scoring SNP in this region is at high frequencies found in hunter-gatherer groups, predominantly in eastern hunter-gatherers (Panel B, Figure S100).

In chr15:38464638-38992430, we recovered RASGRP1, associated with immunity and related to systemic lupus erythematosus [59], rheumatoid arthritis [60] or Epstein-Barr virus [61] among other disorders. The alternative allele at this SNP is at high frequencies in eastern hunter gatherers and other ancient Baltic populations (Panel B, Figure S100). We also found a wide candidate region containing several high-scoring SNPs in chromosome 16:66852047-67871804. The gene that falls in the highest peak of the region is ATP6V0D1, and it plays a very important role in the replication of influenza virus [62,63]. In this region, there are also several genes - *TPPP3/ZDHHC1* and *HSD11B2* - associated with obesity, cardiovascular diseases and hypertension [64–70]. The alternative allele at the top-scoring SNP has intermediate allele frequencies in hunter-gatherer populations, and higher frequencies in eastern hunter-gatherers.

#### Neolithic vs. hunter-gatherer scan

The “neo-hg” scan specifically recovers patterns of allele differentiation along the axis separating hunter-gatherer and farmer populations in West Eurasia (Supplementary Table 11). In this scan, we find a large number of high-scoring regions associated with lipid and sugar metabolism, and various metabolic disorders.

For example, we recover the *FADS* gene cluster, involved in lipid metabolism. This region is presumed to be important in the transition to a diet rich in grains, as a consequence of the expansion of agriculture in Europe and/or the demographic transitions subsequent to it [47,71–73]. This region is also found in the “west-eurasia” scan. The alternative allele frequency at the top-scoring SNP is very high in hunter-gatherer populations.

Another region is located in chromosome 22:31353354-31759255 and also contains genes involved in lipid metabolism: PATZ1, LIMK2, MORC2 and PLA2G3. PATZ1 down-regulates FADS1 [74], LIMK2 shows particularly elevated expression in metabolic syndrome [75], MORC2 plays an important role in cellular lipid metabolism [76–78] and PLA2G3 contributes to atherogenesis [79–81]. The top-scoring SNP in this region has high alternative allele frequencies in Neolithic farmer populations (Panel A, Figure S101).

At chromosome 12:6875213-7366672, we find a region with several genes involved in lipid metabolism: PTPN6, EMG1, PHB2, LPCAT3, C1S**.** LPCAT3 is essential in high fat diets and associated with oleic acid levels and linoleic acid [82] and its deficiency alters cholesterol promoting atherosclerosis [83] and plays an important role in hyperuricemia [83,84]. C1S also plays a crucial role in innate immunity [85] and has been recently associated with coronary syndrome [86].

In chromosome 17:8655348-9223981, we found the PIK3 family (PIK3R5, PIK3R6) which is involved in glucose homeostasis and plays an important role in obesity and insulin resistance in type 2 diabetes [87–89].

In chromosome 11:27432440-27832440 we find a region containing BDNF (brain-derived neurotrophic factor), expression of which is associated with obesity [90,91]. It participates in the reduction of free fatty acids, cholesterol and glucose levels and enhances energy expenditure [92]. Its expression has been shown to be suppressed with a high-fat sucrose diet [90,9394]; [90,93]. High alternative allele frequency in HG (Panel A, Figure S101).

One of the strongest peaks in this scan corresponds to SLC2A12, coding for a glucose transporter. The next peak is located in chromosome 17:79055998-79469799. SLC38A10, which falls in the tip of the peak, is involved in absorption of amino acids from the GI tract [95] and has been suggested to play a role in pathways involved in neurotransmission [96,97]. BAIAP2's [98,99] and AATK's expression [100,101], which are also found in the same region, are related to high-fat and omega3 fatty acids. Higher alternative allele frequencies are found in hunter-gatherers, in particular in Eastern-HG (Panel A, Figure S101).

The highest peak in chromosome 4 contains several genes involved in alcohol metabolism, ADH1B, ADH1C and ADH7 [102,103]. In Panel B, Figure S101, we can see the highest alternative allele frequencies for the oldest samples, which correspond to hunter-gatherers.

Two of the top peaks are related to innate immune response in humans. In chromosome 14:73102086-73502086, ZFYVE1 is involved in TLR3-mediated immune response and regulates antiviral response [104,105]. In chromosome 3:98028061-98476960, GPR15 is related to immune tolerance, and it regulates the homeostasis in the intestine mucosa [104,106,107]. In chromosome 18:46132358-46558884, SMAD7 is associated with inflammatory bowel diseases such as Crohn's disease [108–110].

Two regions are related to brain disorders. In chromosome 1:110588519-111127548, several genes regulate neuronal ion channels: KCNC4, SLC6A17, and STRIP1. Mutations in SLC6A17 cause intellectual disabilities associated with speech impairment and behavioural problems [111], while the KCNC gene family has also been associated with intellectual disability [112].

### Figures

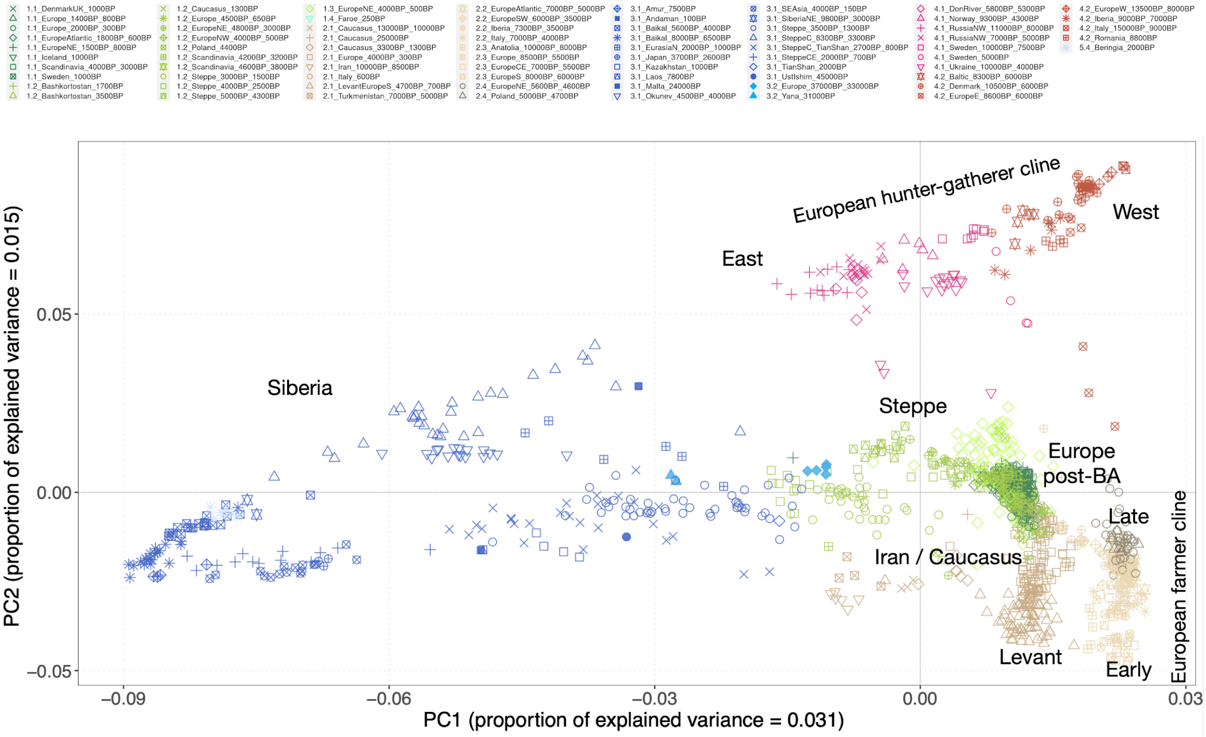


**Figure S93.** Principal component analysis on 1402 Eurasian samples. The first component explains 3.1% of the variance and separates East Asian, Steppe and European samples. The second component separates farmers and Hunter-gatherers (1.5%).

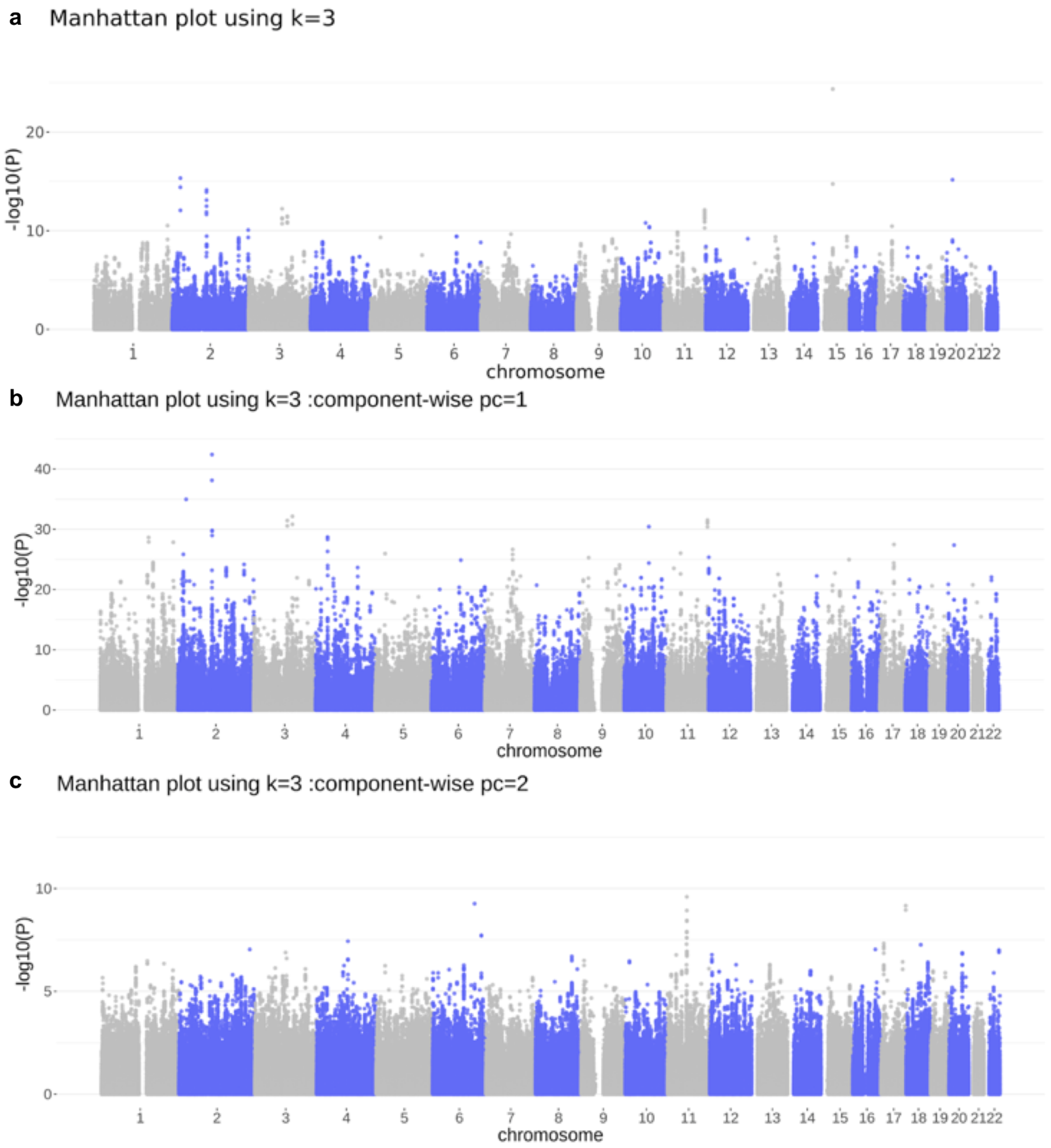


**Figure S94**. Genome scan using pcadapt k=3. **A.** Method: mahalanobis distances. Manhattan plot scanning the whole genome. **B** and **C**. Component-wise genome scans for component PC1 and component PC2, respectively.

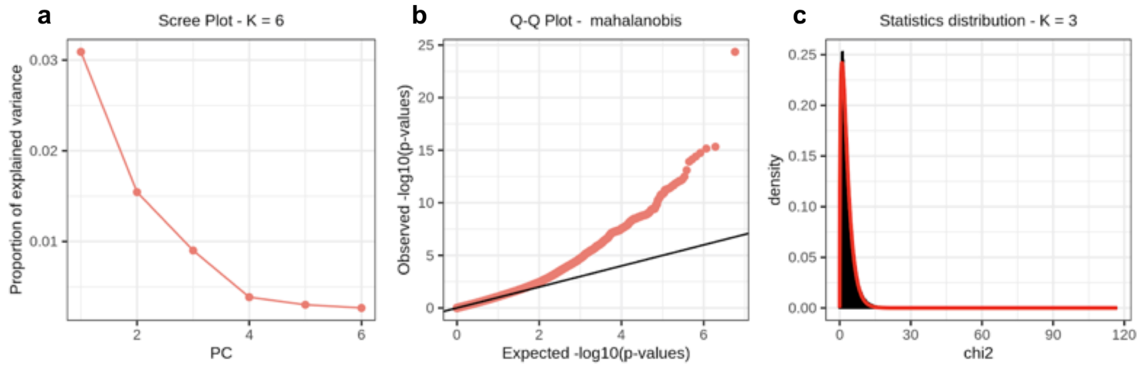


**Figure S95. A.** Scree plot showing proportion of explained variance of the first PCs in the pcadapt analysis. **B.** Q-Q plot using mahalanobis method for K=3. **C.** Distribution of pcadapt scores (k=3) compared to chi-squared distribution with one degree of freedom (red line).

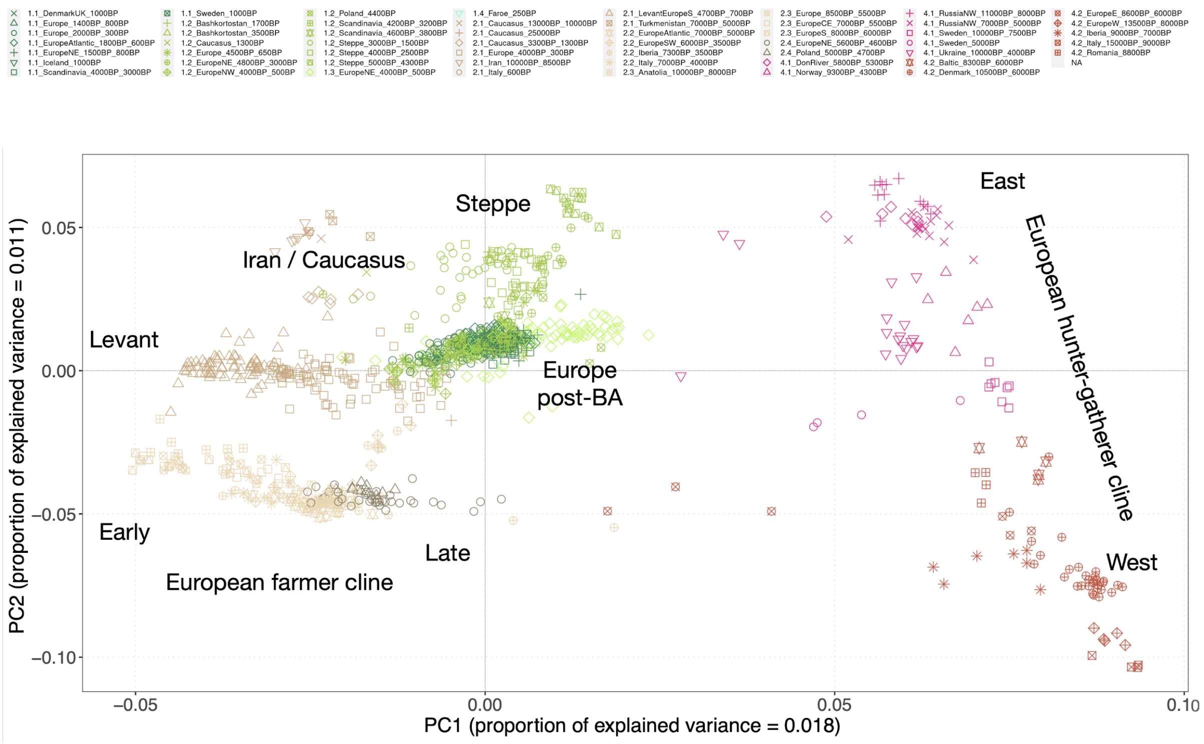


**Figure S96.** Principal component analysis on 1165 Eurasian samples. The first component explains 1.8% of the variance and separates East Asian, Steppe and European samples. The second component separates farmers and Hunter-gatherers (1.1%).

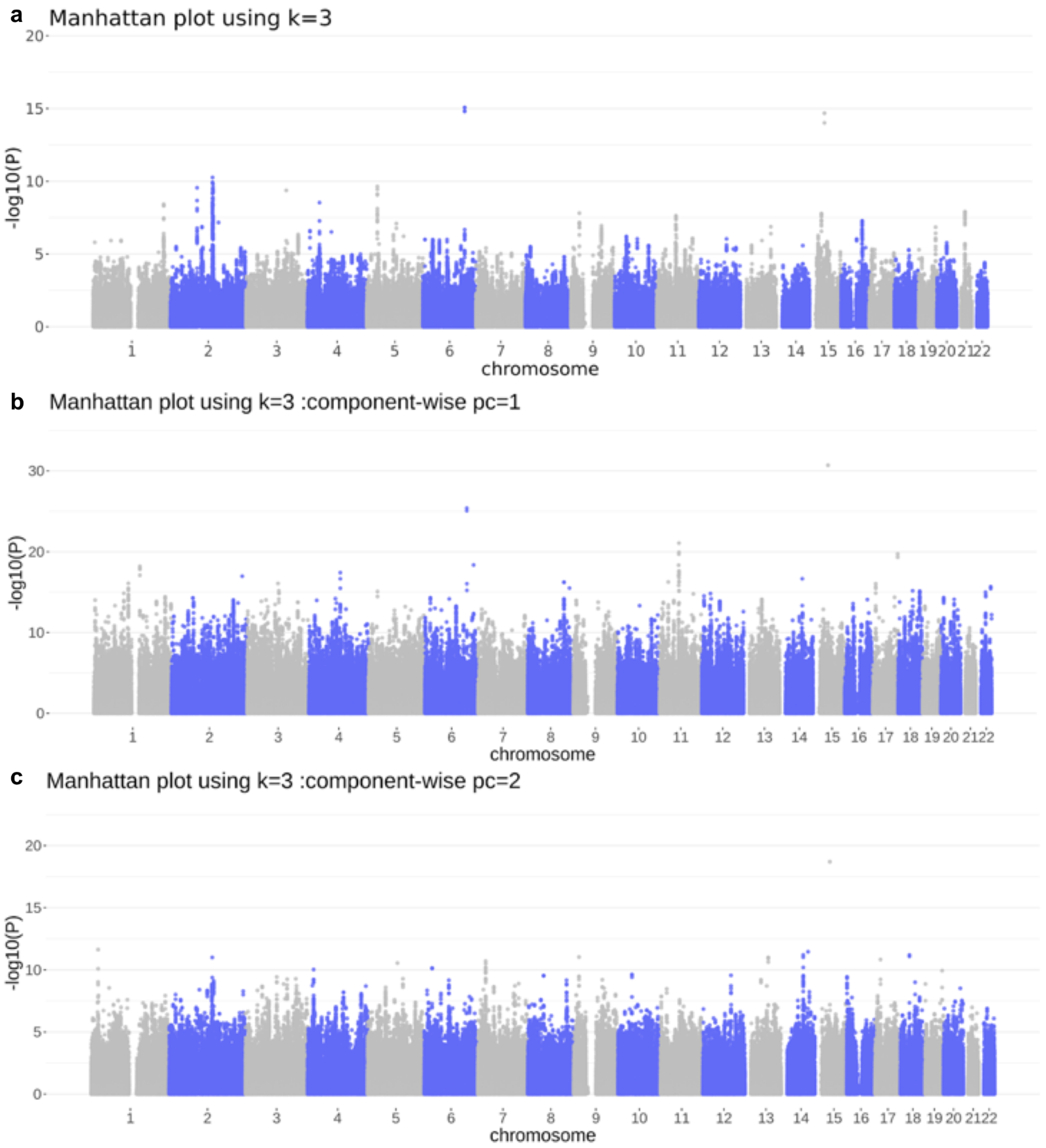


**Figure S97**. Genome scan using pcadapt k=3. **A.** Method: mahalanobis distances. Manhattan plot scanning the whole genome. **B** and **C.** Component-wise genome scans for component PC1 and component PC2, respectively.

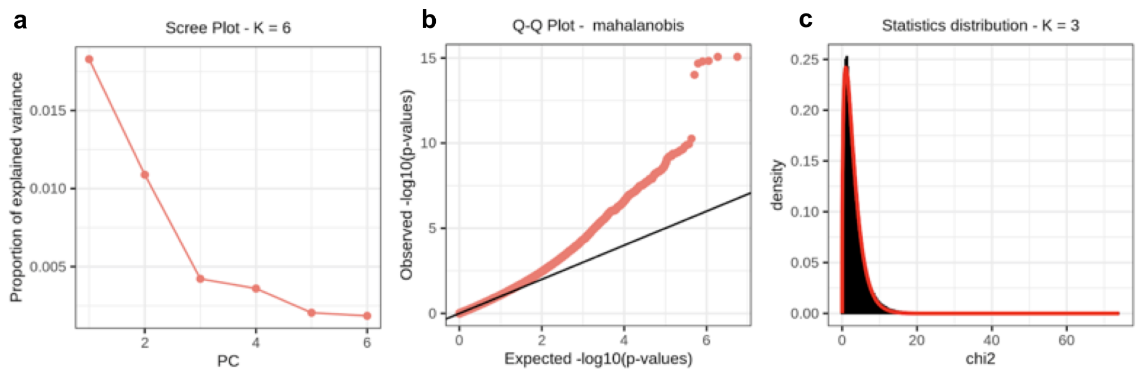


**Figure S98**. **A.** Scree plot showing proportion of explained variance of the first PCs in the pcadapt analysis. **B.** Q-Q plot using mahalanobis method for K=3. Distribution of pcadapt scores (k=3) compared to chi-squared distribution with one degree of freedom (red line).

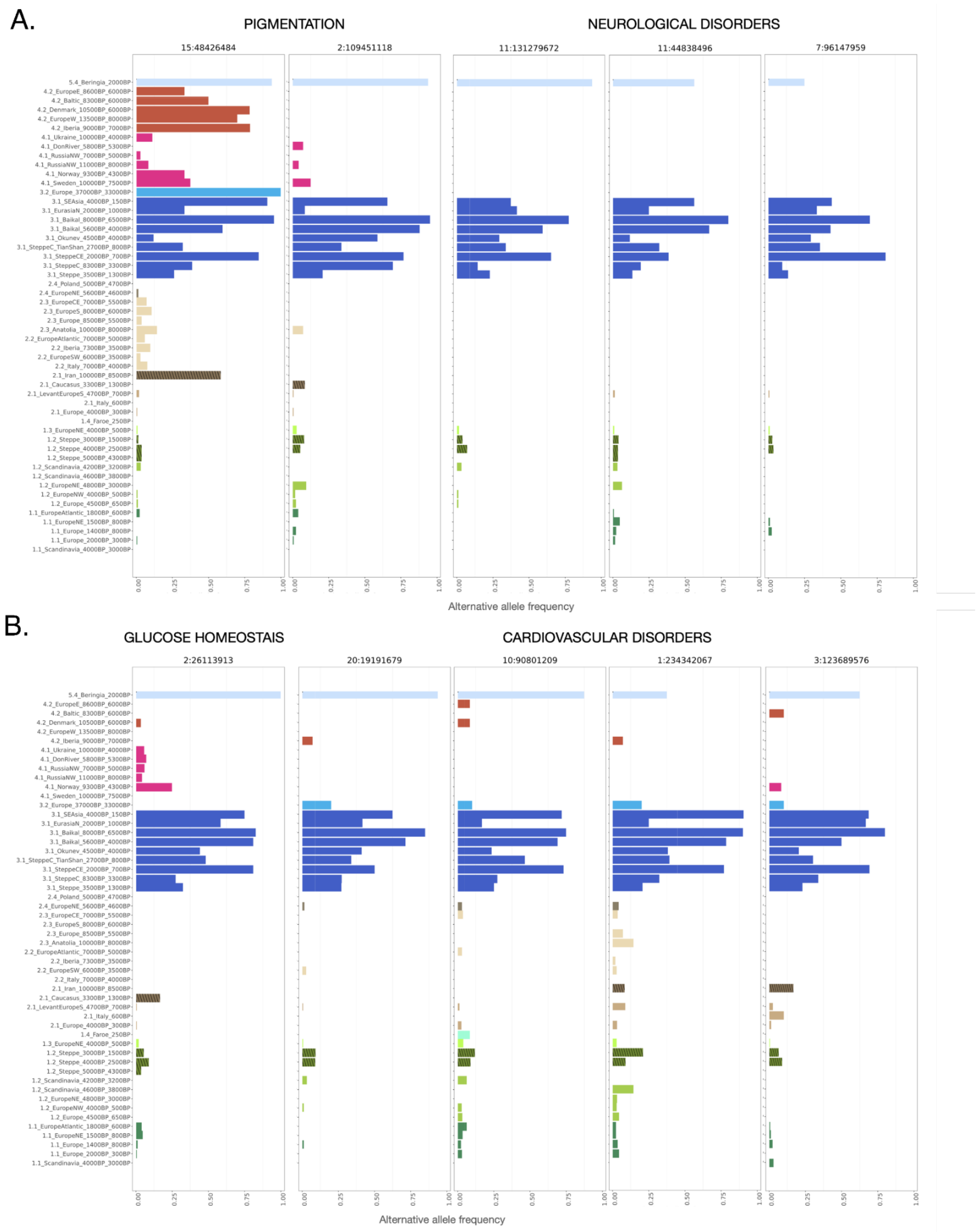


**Figure S99**. Alternative allele frequencies of the position with the lowest p-value in the top regions for the pcadapt eurasian scan.

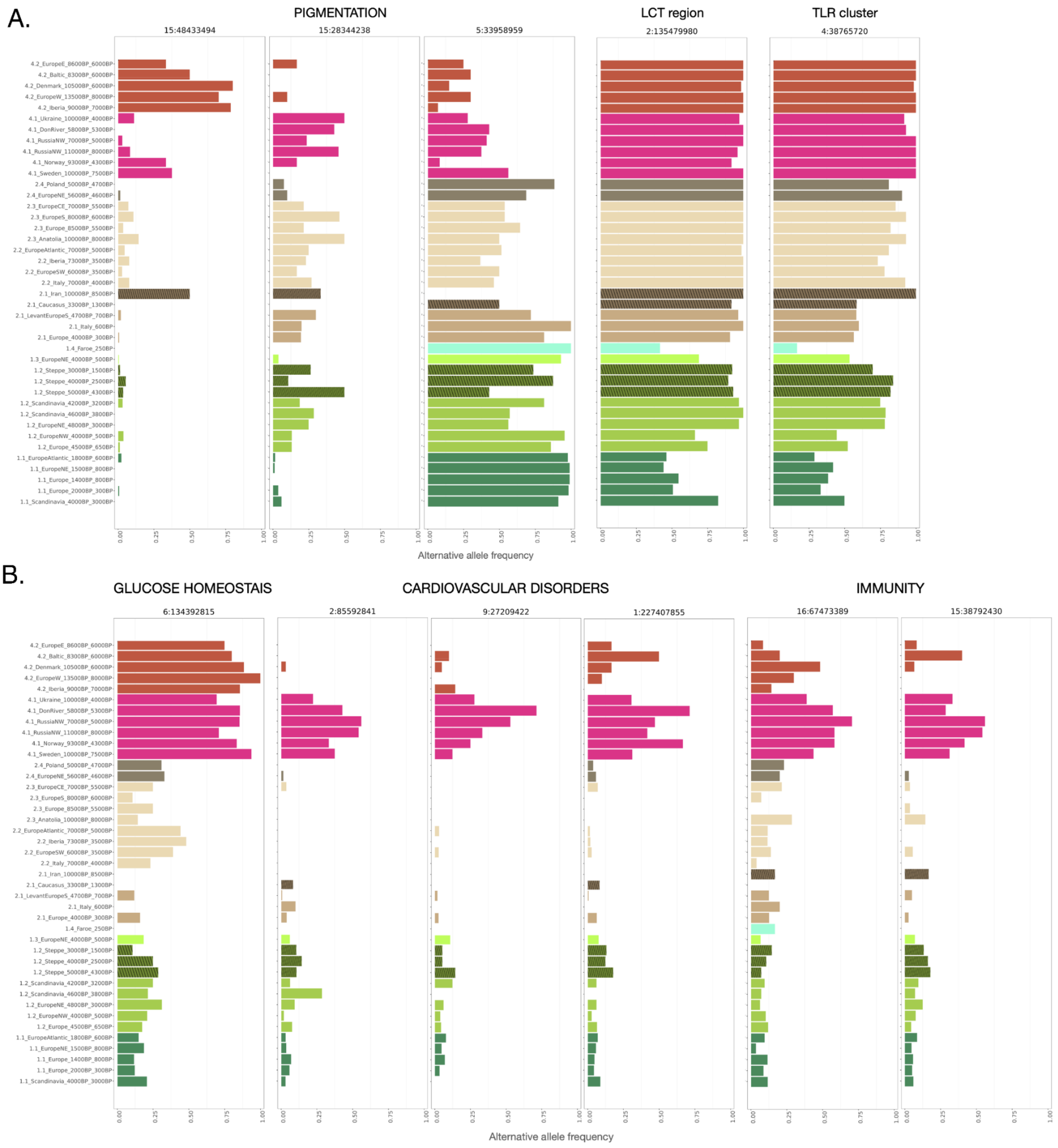


**Figure S100**. Alternative allele frequencies of the position with the lowest p-value in the top regions for the pcadapt West-eurasian scan.

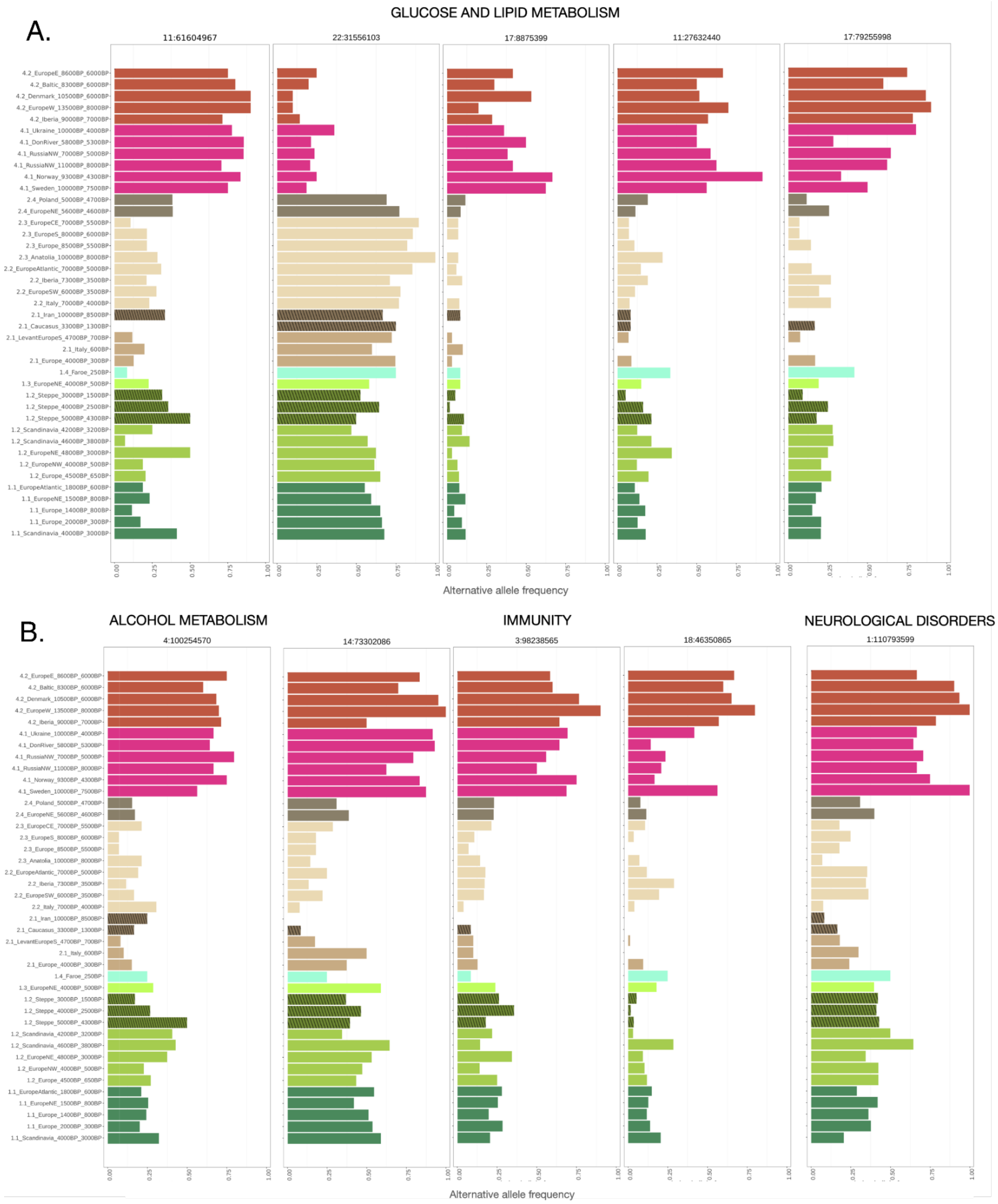


**Figure S101**. Alternative allele frequencies of the position with the lowest p-value in the top regions for the pcadapt HG West-eurasian scan

#### References


1. Luu, K., Bazin, E. & Blum, M. G. B. pcadapt: an R package to perform genome scans for selection based on principal component analysis. *Mol. Ecol. Resour.* **17**, 67–77 (2017).

2. Durinck, S., Spellman, P. T., Birney, E. & Huber, W. Mapping identifiers for the integration of genomic datasets with the R/Bioconductor package biomaRt. *Nat. Protoc.* **4**, 1184–1191 (2009).

3. Wilde, S. *et al.* Direct evidence for positive selection of skin, hair, and eye pigmentation in Europeans during the last 5,000 y. *Proc. Natl. Acad. Sci. U. S. A.* **111**, 4832–4837 (2014).

4. Pickrell, J. K. *et al.* Signals of recent positive selection in a worldwide sample of human populations. *Genome Research* vol. 19 826–837 (2009).

5. Lao, O., de Gruijter, J. M., van Duijn, K., Navarro, A. & Kayser, M. Signatures of positive selection in genes associated with human skin pigmentation as revealed from analyses of single nucleotide polymorphisms. *Ann. Hum. Genet.* **71**, 354–369 (2007).

6. Herchuelz, A. *Sodium-calcium exchange and the plasma membrane Ca2+-ATPase in cell function: fifth international conference*. (Wiley-Blackwell, 2007).

7. Bryk, J. *et al.* Positive Selection in East Asians for an EDAR Allele that Enhances NF-κB Activation. *PLoS ONE* vol. 3 e2209 (2008).

8. Hider, J. L. *et al.* Exploring signatures of positive selection in pigmentation candidate genes in populations of East Asian ancestry. *BMC Evol. Biol.* **13**, 150 (2013).

9. Duforet-Frebourg, N., Luu, K., Laval, G., Bazin, E. & Blum, M. G. B. Detecting Genomic Signatures of Natural Selection with Principal Component Analysis: Application to the 1000 Genomes Data. *Mol. Biol. Evol.* **33**, 1082–1093 (2016).

10. Prashanth, S. & Deshmukh, S. Ectodermal Dysplasia: A Genetic Review. *International Journal of Clinical Pediatric Dentistry* vol. 5 197–202 (2012).

11. Kimura, R. *et al.* A common variation in EDAR is a genetic determinant of shovel-shaped incisors. *Am. J. Hum. Genet.* **85**, 528–535 (2009).

12. Izawa, T. *et al.* ASXL2 Regulates Glucose, Lipid, and Skeletal Homeostasis. *Cell Rep.* **11**, 1625–1637 (2015).

13. Zou, W. *et al.* Myeloid-specific Asxl2 deletion limits diet-induced obesity by regulating energy expenditure. *J. Clin. Invest.* **130**, 2644–2656 (2020).

14. Park, U.-H., Yoon, S. K., Park, T., Kim, E.-J. & Um, S.-J. Additional Sex Comb-like (ASXL) Proteins 1 and 2 Play Opposite Roles in Adipogenesis via Reciprocal Regulation of Peroxisome Proliferator-activated Receptor γ. *Journal of Biological Chemistry* vol. 286 1354–1363 (2011).

15. Ponsuksili, S. *et al.* Epigenome-wide skeletal muscle DNA methylation profiles at the background of distinct metabolic types and ryanodine receptor variation in pigs. *BMC Genomics* **20**, 492 (2019).

16. Samad, M. B. *et al.* [6]-Gingerol, from Zingiber officinale, potentiates GLP-1 mediated glucose-stimulated insulin secretion pathway in pancreatic β-cells and increases RAB8/RAB10-regulated membrane presentation of GLUT4 transporters in skeletal muscle to improve hyperglycemia in Leprdb/db type 2 diabetic mice. *BMC Complementary and Alternative Medicine* vol. 17 (2017).

17. Vazirani, R. P. *et al.* Disruption of Adipose Rab10-Dependent Insulin Signaling Causes Hepatic Insulin Resistance. *Diabetes* **65**, 1577–1589 (2016).

18. Hsieh, P. *et al.* Exome Sequencing Provides Evidence of Polygenic Adaptation to a Fat-Rich Animal Diet in Indigenous Siberian Populations. *Mol. Biol. Evol.* **34**, 2913–2926 (2017).

19. Thapa, D. *et al.* The protein acetylase GCN5L1 modulates hepatic fatty acid oxidation activity via acetylation of the mitochondrial β-oxidation enzyme HADHA. *J. Biol. Chem.* **293**, 17676–17684 (2018).

20. Baloni, P. *et al.* Genome-scale metabolic model of the rat liver predicts effects of diet restriction. *Sci. Rep.* **9**, 9807 (2019).

21. Ong, H. S. & Yim, H. C. H. Microbial Factors in Inflammatory Diseases and Cancers. *Regulation of Inflammatory Signaling in Health and Disease* 153–174 (2017) doi:10.1007/978-981-10-5987-2_7.

22. Logsdon, B. A., Hoffman, G. E. & Mezey, J. G. Mouse obesity network reconstruction with a variational Bayes algorithm to employ aggressive false positive control. *BMC Bioinformatics* **13**, 53 (2012).

23. Pei, Y.-F. *et al.* Genomic variants at 20p11 associated with body fat mass in the European population. *Obesity* **25**, 757–764 (2017).

24. Sabir, J. S. M. *et al.* Unraveling the role of salt-sensitivity genes in obesity with integrated network biology and co-expression analysis. *PLoS One* **15**, e0228400 (2020).

25. Wu, H. *et al.* Transcriptome Sequencing to Detect the Potential Role of Long Noncoding RNAs in Salt-Sensitive Hypertensive Rats. *Biomed Res. Int.* **2019**, 2816959 (2019).

26. Wang, L. *et al.* Peakwide Mapping on Chromosome 3q13 Identifies the Kalirin Gene as a Novel Candidate Gene for Coronary Artery Disease. *The American Journal of Human Genetics* vol. 80 650–663 (2007).

27. Ikram, M. A., Seshadri, S. & Bis, J. C. Genomewide Association Studies of Stroke. *Journal of Vascular Surgery* vol. 50 467 (2009).

28. Krug, T. *et al.* Kalirin: a novel genetic risk factor for ischemic stroke. *Hum. Genet.* **127**, 513–523 (2010).

29. Zang, X.-L. *et al.* Association of a SNP in SLC35F3 Gene with the Risk of Hypertension in a Chinese Han Population. *Frontiers in Genetics* vol. 7 (2016).

30. Zhang, K. *et al.* Genetic implication of a novel thiamine transporter in human hypertension. *J. Am. Coll. Cardiol.* **63**, 1542–1555 (2014).

31. Pacheu-Grau, D. *et al.* COA6 Facilitates Cytochrome c Oxidase Biogenesis as Thiol-reductase for Copper Metallochaperones in Mitochondria. *J. Mol. Biol.* **432**, 2067–2079 (2020).

32. Russo, L. *et al.* Cholesterol 25-hydroxylase (CH25H) as a promoter of adipose tissue inflammation in obesity and diabetes. *Mol Metab* **39**, 100983 (2020).

33. Zhao, J., Chen, J., Li, M., Chen, M. & Sun, C. Multifaceted Functions of CH25H and 25HC to Modulate the Lipid Metabolism, Immune Responses, and Broadly

Antiviral Activities. *Viruses* **12**, (2020).

34. Demir, A., Kahraman, R., Candan, G. & Ergen, A. The role of FAS gene variants in inflammatory bowel disease. *Turk. J. Gastroenterol.* **31**, 356–361 (2020).

35. Rieux-Laucat, F., Magérus-Chatinet, A. & Neven, B. The Autoimmune Lymphoproliferative Syndrome with Defective FAS or FAS-Ligand Functions. *Journal of Clinical Immunology* vol. 38 558–568 (2018).

36. Karis, K. *et al.* Altered Expression Profile of IgLON Family of Neural Cell Adhesion Molecules in the Dorsolateral Prefrontal Cortex of Schizophrenic Patients. *Front. Mol. Neurosci.* **11**, 8 (2018).

37. Liu, J., Li, M. & Su, B. GWAS-identified schizophrenia risk SNPs atTSPAN18are highly diverged between Europeans and East Asians. *American Journal of Medical Genetics Part B: Neuropsychiatric Genetics* vol. 171 1032–1040 (2016).

38. Fu, H.-Y. *et al.* The mutation spectrum of the SLC25A13 gene in Chinese infants with intrahepatic cholestasis and aminoacidemia. *J. Gastroenterol.* **46**, 510–518 (2011).

39. Chen, J.-L., Zhang, Z.-H., Li, B.-X., Cai, Z. & Zhou, Q.-H. Bioinformatic and functional analysis of promoter region of human SLC25A13 gene. *Gene* **693**, 69–75 (2019).

40. Vitoria, I. *et al.* Citrin deficiency in a Romanian child living in Spain highlights the worldwide distribution of this defect and illustrates the value of nutritional therapy. *Molecular Genetics and Metabolism* vol. 110 181–183 (2013).

41. Fiermonte, G. *et al.* An adult with type 2 citrullinemia presenting in Europe. *N. Engl. J. Med.* **358**, 1408–1409 (2008).

42. Sarkar, A. & Nandineni, M. R. Association of common genetic variants with human skin color variation in Indian populations. *Am. J. Hum. Biol.* **30**, (2018).

43. Edwards, M. *et al.* Association of the OCA2 polymorphism His615Arg with melanin content in east Asian populations: further evidence of convergent evolution of skin pigmentation. *PLoS Genet.* **6**, e1000867 (2010).

44. Eiberg, H. *et al.* Blue eye color in humans may be caused by a perfectly

associated founder mutation in a regulatory element located within the HERC2 gene inhibiting OCA2 expression. *Human Genetics* vol. 123 177–187 (2008).

45. Sturm, R. A. *et al.* A Single SNP in an Evolutionary Conserved Region within Intron 86 of the HERC2 Gene Determines Human Blue-Brown Eye Color. *The American Journal of Human Genetics* vol. 82 424–431 (2008).

46. Allentoft, M. E. *et al.* Population genomics of Bronze Age Eurasia. *Nature* **522**, 167–172 (2015).

47. Mathieson, I. *et al.* Genome-wide patterns of selection in 230 ancient Eurasians. *Nature* **528**, 499–503 (2015).

48. Barreiro, L. B. *et al.* Evolutionary dynamics of human Toll-like receptors and their different contributions to host defense. *PLoS Genet.* **5**, e1000562 (2009).

49. Astiz, M. & Oster, H. GLUT12-A promising new target for the treatment of insulin resistance in obesity and type 2 diabetes. *Acta physiologica* vol. 226 e13329 (2019).

50. Waller, A. P. *et al.* GLUT12 functions as a basal and insulin-independent glucose transporter in the heart. *Biochim. Biophys. Acta* **1832**, 121–127 (2013).

51. Joehanes, R. *et al.* Integrated genome-wide analysis of expression quantitative trait loci aids interpretation of genomic association studies. *Genome Biol.* **18**, 16 (2017).

52. CARDIoGRAMplusC4D Consortium *et al.* Large-scale association analysis identifies new risk loci for coronary artery disease. *Nat. Genet.* **45**, 25–33 (2013).

53. Limaye, N. *et al.* Somatic mutations in angiopoietin receptor gene TEK cause solitary and multiple sporadic venous malformations. *Nat. Genet.* **41**, 118–124 (2009).

54. Jones, N., Iljin, K., Dumont, D. J. & Alitalo, K. Tie receptors: new modulators of angiogenic and lymphangiogenic responses. *Nature Reviews Molecular Cell Biology* vol. 2 257–267 (2001).

55. Dumont, D. J. *et al.* Dominant-negative and targeted null mutations in the endothelial receptor tyrosine kinase, tek, reveal a critical role in vasculogenesis of the embryo. *Genes Dev.* **8**, 1897–1909 (1994).

56. Gál, Z. *et al.* Investigation of the Possible Role of Tie2 Pathway and TEK Gene in Asthma and Allergic Conjunctivitis. *Frontiers in Genetics* vol. 11 (2020).

57. Cmejla, R. *et al.* Human MRCKalpha is regulated by cellular iron levels and interferes with transferrin iron uptake. *Biochem. Biophys. Res. Commun.* **395**, 163–167 (2010).

58. Richard, C. *et al.* Myotonic dystrophy kinase-related CDC42-binding kinase α, a new transferrin receptor type 2-binding partner, is a regulator of erythropoiesis. *Am. J. Hematol.* (2021) doi:10.1002/ajh.26104.

59. Molineros, J. E. *et al.* Mechanistic Characterization of RASGRP1 Variants Identifies an hnRNP-K-Regulated Transcriptional Enhancer Contributing to SLE Susceptibility. *Frontiers in Immunology* vol. 10 (2019).

60. Potier, M. L. *et al.* RasGRP1 and RasGRP3 expression in lymphocytes of rheumatoid arthritis patients. *Annals of the Rheumatic Diseases* vol. 71 A54.2–A54 (2012).

61. Somekh, I. *et al.* Correction to: Novel Mutations in RASGRP1 Are Associated with Immunodeficiency, Immune Dysregulation, and EBV-Induced Lymphoma. *J. Clin. Immunol.* **38**, 711 (2018).

62. Karlas, A. *et al.* Genome-wide RNAi screen identifies human host factors crucial for influenza virus replication. *Nature* **463**, 818–822 (2010).

63. Hao, L. *et al.* Drosophila RNAi screen identifies host genes important for influenza virus replication. *Nature* **454**, 890–893 (2008).

64. Mariniello, B. *et al.* Analysis of the 11β-Hydroxysteroid Dehydrogenase Type 2 Gene (HSD11B2) in Human Essential Hypertension*. *Am. J. Hypertens.* **18**, 1091–1098 (2005).

65. Zeller, T. *et al.* Transcriptome-Wide Analysis Identifies Novel Associations With Blood Pressure. *Hypertension* **70**, 743–750 (2017).

66. Gunaratnam, K., Vidal, C., Gimble, J. M. & Duque, G. Mechanisms of palmitate-induced lipotoxicity in human osteoblasts. *Endocrinology* **155**, 108–116 (2014).

67. Lumish, H. S., O'Reilly, M. & Reilly, M. P. Sex Differences in Genomic Drivers of Adipose Distribution and Related Cardiometabolic Disorders: Opportunities for Precision Medicine. *Arterioscler. Thromb. Vasc. Biol.* **40**, 45–60 (2020).

68. Ng, M. C. Y. *et al.* Discovery and fine-mapping of adiposity loci using high density imputation of genome-wide association studies in individuals of African ancestry: African Ancestry Anthropometry Genetics Consortium. *PLoS Genet.* **13**, e1006719 (2017).

69. Ueda, K. *et al.* Renal Dysfunction Induced by Kidney-Specific Gene Deletion of as a Primary Cause of Salt-Dependent Hypertension. *Hypertension* **70**, 111–118 (2017).

70. Agarwal, A. K. *et al.* CA-Repeat polymorphism in intron 1 of HSD11B2 : effects on gene expression and salt sensitivity. *Hypertension* **36**, 187–194 (2000).

71. Buckley, M. T. *et al.* Selection in Europeans on Fatty Acid Desaturases Associated with Dietary Changes. *Mol. Biol. Evol.* **34**, 1307–1318 (2017).

72. Ye, K., Gao, F., Wang, D., Bar-Yosef, O. & Keinan, A. Dietary adaptation of FADS genes in Europe varied across time and geography. *Nat Ecol Evol* **1**, 167 (2017).

73. Mathieson, S. & Mathieson, I. FADS1 and the Timing of Human Adaptation to Agriculture. *Mol. Biol. Evol.* **35**, 2957–2970 (2018).

74. Pan, G. *et al.* PATZ1 down-regulates FADS1 by binding to rs174557 and is opposed by SP1/SREBP1c. *Nucleic Acids Res.* **45**, 2408–2422 (2017).

75. Tabur, S. *et al.* Evidence for elevated (LIMK2 and CFL1) and suppressed (ICAM1, EZR, MAP2K2, and NOS3) gene expressions in metabolic syndrome. *Endocrine* **53**, 465–470 (2016).

76. Fairn, G. D. & McMaster, C. R. Emerging roles of the oxysterol-binding protein family in metabolism, transport, and signaling. *Cell. Mol. Life Sci.* **65**, 228–236 (2008).

77. Lehto, M. & Olkkonen, V. M. The OSBP-related proteins: a novel protein family involved in vesicle transport, cellular lipid metabolism, and cell signalling. *Biochim. Biophys. Acta* **1631**, 1–11 (2003).

78. Sánchez-Solana, B., Li, D.-Q. & Kumar, R. Cytosolic functions of MORC2 in

lipogenesis and adipogenesis. *Biochim. Biophys. Acta* **1843**, 316–326 (2014).

79. Sartipy, P., Camejo, G., Svensson, L. & Hurt-Camejo, E. Phospholipase A2 modification of lipoproteins: potential effects on atherogenesis. *Adv. Exp. Med. Biol.* **507**, 3–7 (2002).

80. Dennis, E. A., Cao, J., Hsu, Y.-H., Magrioti, V. & Kokotos, G. Phospholipase A2 Enzymes: Physical Structure, Biological Function, Disease Implication, Chemical Inhibition, and Therapeutic Intervention. *Chem. Rev.* **111**, 6130 (2011).

81. Wyles, J. P., Perry, R. J. & Ridgway, N. D. Characterization of the sterol-binding domain of oxysterol-binding protein (OSBP)-related protein 4 reveals a novel role in vimentin organization. *Experimental Cell Research* vol. 313 1426–1437 (2007).

82. Tintle, N. L. *et al.* A genome-wide association study of saturated, mono- and polyunsaturated red blood cell fatty acids in the Framingham Heart Offspring Study. *Prostaglandins Leukot. Essent. Fatty Acids* **94**, 65–72 (2015).

83. Thomas, C. *et al.* LPCAT3 deficiency in hematopoietic cells alters cholesterol and phospholipid homeostasis and promotes atherosclerosis. *Atherosclerosis* **275**, 409–418 (2018).

84. Liu, N. *et al.* Hyperuricemia induces lipid disturbances mediated by LPCAT3 upregulation in the liver. *FASEB J.* (2020) doi:10.1096/fj.202000950R.

85. Godahewa, G. I., Bathige, S. D. N. K., Herath, H. M. L. P. B., Noh, J. K. & Lee, J. Characterization of rock bream (Oplegnathus fasciatus) complement components C1r and C1s in terms of molecular aspects, genomic modulation, and immune responsive transcriptional profiles following bacterial and viral pathogen exposure. *Fish Shellfish Immunol.* **46**, 656–668 (2015).

86. Dai, D.-F. *et al.* Plasma concentration of SCUBE1, a novel platelet protein, is elevated in patients with acute coronary syndrome and ischemic stroke. *J. Am. Coll. Cardiol.* **51**, 2173–2180 (2008).

87. Huang, X., Liu, G., Guo, J. & Su, Z. The PI3K/AKT pathway in obesity and type 2 diabetes. *Int. J. Biol. Sci.* **14**, 1483–1496 (2018).

88. Zhong, X. *et al.* LNK deficiency decreases obesity-induced insulin resistance

by regulating GLUT4 through the PI3K-Akt-AS160 pathway in adipose tissue. *Aging* **12**, 17150–17166 (2020).

89. López-Gómez, C. *et al.* Oleic Acid Protects Against Insulin Resistance by Regulating the Genes Related to the PI3K Signaling Pathway. *J. Clin. Med. Res.* **9**, (2020).

90. Sandrini, L. *et al.* Association between Obesity and Circulating Brain-Derived Neurotrophic Factor (BDNF) Levels: Systematic Review of Literature and Meta-Analysis. *Int. J. Mol. Sci.* **19**, (2018).

91. Lommatzsch, M. *et al.* The impact of age, weight and gender on BDNF levels in human platelets and plasma. *Neurobiol. Aging* **26**, 115–123 (2005).

92. Tsuchida, A. *et al.* Brain-derived neurotrophic factor ameliorates lipid metabolism in diabetic mice. *Diabetes Obes. Metab.* **4**, 262–269 (2002).

93. Wu, A., Molteni, R., Ying, Z. & Gomez-Pinilla, F. A saturated-fat diet aggravates the outcome of traumatic brain injury on hippocampal plasticity and cognitive function by reducing brain-derived neurotrophic factor. *Neuroscience* **119**, 365–375 (2003).

94. Molteni, R., Barnard, R. J., Ying, Z., Roberts, C. K. & Gómez-Pinilla, F. A high-fat, refined sugar diet reduces hippocampal brain-derived neurotrophic factor, neuronal plasticity, and learning. *Neuroscience* **112**, 803–814 (2002).

95. Graber, T. G., Borack, M. S., Reidy, P. T., Volpi, E. & Rasmussen, B. B. Essential amino acid ingestion alters expression of genes associated with amino acid sensing, transport, and mTORC1 regulation in human skeletal muscle. *Nutr. Metab.* **14**, 35 (2017).

96. Hellsten, S. V., Hägglund, M. G., Eriksson, M. M. & Fredriksson, R. The neuronal and astrocytic protein SLC38A10 transports glutamine, glutamate, and aspartate, suggesting a role in neurotransmission. *FEBS Open Bio* **7**, 730–746 (2017).

97. Tripathi, R., Hosseini, K., Arapi, V., Fredriksson, R. & Bagchi, S. SLC38A10 (SNAT10) is Located in ER and Golgi Compartments and Has a Role in Regulating Nascent Protein Synthesis. *International Journal of Molecular Sciences* vol. 20 6265 (2019).

98. Schmidt, S. *et al.* Effect of omega-3 polyunsaturated fatty acids on the cytoskeleton: an open-label intervention study. *Lipids Health Dis.* **14**, 4 (2015).

99. Zulkafli, I. S., Waddell, B. J. & Mark, P. J. Postnatal Dietary Omega-3 Fatty Acid Supplementation Rescues Glucocorticoid-Programmed Adiposity, Hypertension, and Hyperlipidemia in Male Rat Offspring Raised on a High-Fat Diet. *Endocrinology* vol. 154 3110–3117 (2013).

100. Aqil, M., Mallik, S., Bandyopadhyay, S., Maulik, U. & Jameel, S. Transcriptomic Analysis of mRNAs in Human Monocytic Cells Expressing the HIV-1 Nef Protein and Their Exosomes. *Biomed Res. Int.* **2015**, (2015).

101. Kirpich, I. A. *et al.* Integrated hepatic transcriptome and proteome analysis of mice with high-fat diet-induced nonalcoholic fatty liver disease. *J. Nutr. Biochem.* **22**, 38–45 (2011).

102. Li, H. *et al.* Diversification of the ADH1B gene during expansion of modern humans. *Ann. Hum. Genet.* **75**, 497–507 (2011).

103. Polimanti, R. & Gelernter, J. ADH1B: From alcoholism, natural selection, and cancer to the human phenome. *Am. J. Med. Genet. B Neuropsychiatr. Genet.* **177**, 113–125 (2018).

104. Zhong, X. *et al.* The zinc-finger protein ZFYVE1 modulates TLR3-mediated signaling by facilitating TLR3 ligand binding. *Cell. Mol. Immunol.* **17**, 741–752 (2020).

105. Yang, Y. *et al.* The RNA-binding protein Mex3B is a coreceptor of Toll-like receptor 3 in innate antiviral response. *Cell Res.* **26**, 288–303 (2016).

106. Kim, S. V. *et al.* GPR15-mediated homing controls immune homeostasis in the large intestine mucosa. *Science* **340**, 1456–1459 (2013).

107. Nguyen, L. P. *et al.* Role and species-specific expression of colon T cell homing receptor GPR15 in colitis. *Nature Immunology* vol. 16 207–213 (2015).

108. Monteleone, G., Boirivant, M., Pallone, F. & MacDonald, T. T. TGF-β1 and Smad7 in the regulation of IBD. *Mucosal Immunology* vol. 1 S50–S53 (2008).

109. Kennedy, B. W. C. Mongersen, an Oral SMAD7 Antisense Oligonucleotide, and Crohn's Disease. *The New England journal of medicine* vol. 372 2461 (2015).

110. Garo, L. P. *et al.* Smad7 Controls Immunoregulatory PDL2/1-PD1 Signaling in Intestinal Inflammation and Autoimmunity. *Cell Rep.* **28**, 3353–3366.e5 (2019).

111. Iqbal, Z. *et al.* Homozygous SLC6A17 mutations cause autosomal-recessive intellectual disability with progressive tremor, speech impairment, and behavioral problems. *Am. J. Hum. Genet.* **96**, 386–396 (2015).

112. Park, J. *et al.* KCNC1-related disorders: new de novo variants expand the phenotypic spectrum. *Ann Clin Transl Neurol* **6**, 1319–1326 (2019).

### 8) Calling chr17q21.31 KANSL1 duplications in ancient genomes

Alma S. Halgren[1], Andrés Ingason[2,3], and Peter H. Sudmant[1]

[1] Department of Integrative Biology, University of California, Berkeley
[2] Institute of Biological Psychiatry, Mental Health Services, Copenhagen University Hospital
[3] Lundbeck Foundation GeoGenetics Centre, GLOBE Institute, University of Copenhagen, Copenhagen, Denmark

#### Introduction

The 17q21.31 locus harbours a large ~970 kb inversion that is segregating at high (~20-25%) frequency in European populations [1]. The inversion is present in two orientations with respect to the reference, H1, the direct orientation, and H2, the inverted orientation. The direct, H1 haplotype has been associated with neurodegeneration while the H2 haplotype predisposes to microdeletions [2]. The H2 inverted haplotype has also been reported to be under selection with increased fecundity observed in carriers [1]. However, the H1 and H2 haplotypes also exhibit additional structural complexity with both harbouring unique duplication polymorphisms [3,4]. Here, we explored the frequency of the duplications occurring in on the H1 and H2 inversion haplotypes.

#### Methods

To call the distinct H1 and H2 specific KANSL1 duplications at chr17q21.31 (H1D and H2D respectively), we first removed samples that were too noisy to be accurately genotyped. To do so, we calculated the standard deviation of genome-wide copy number (after removing the top and bottom fifth percentiles of copy number to exclude outliers). We chose standard deviation cut-offs based on a visual inspection of the data – 0.49 for the fastq samples and 0.804 for the bam samples. While bam and fastq samples exhibited genome-wide differences in coverage and noise resulting in these different cut-offs, the selected samples exhibit similar signatures at the KANSL1 locus. Together this resulted in a total of 1143 samples, 427 fastq samples and 716 bam samples. We then set a copy number cut-off of 10 at the KANSL1 locus (copy number signal above 10 is likely noise) and calculated the average copy number in the H1D and H2D coordinate blocks (Figure S102). We filtered for samples with an average copy number in both the H1D and H2D blocks between 1.5 and

5.25 (values outside of these bounds are likely noise). The samples clustered into duplication genotypes (coloured below) with the exception of a handful of samples on the border of genotype groups for which we could not accurately assign a genotype (marked with ‘x’). Figure S103 shows the SNP inversion calls for these samples, which align with the duplication calls.

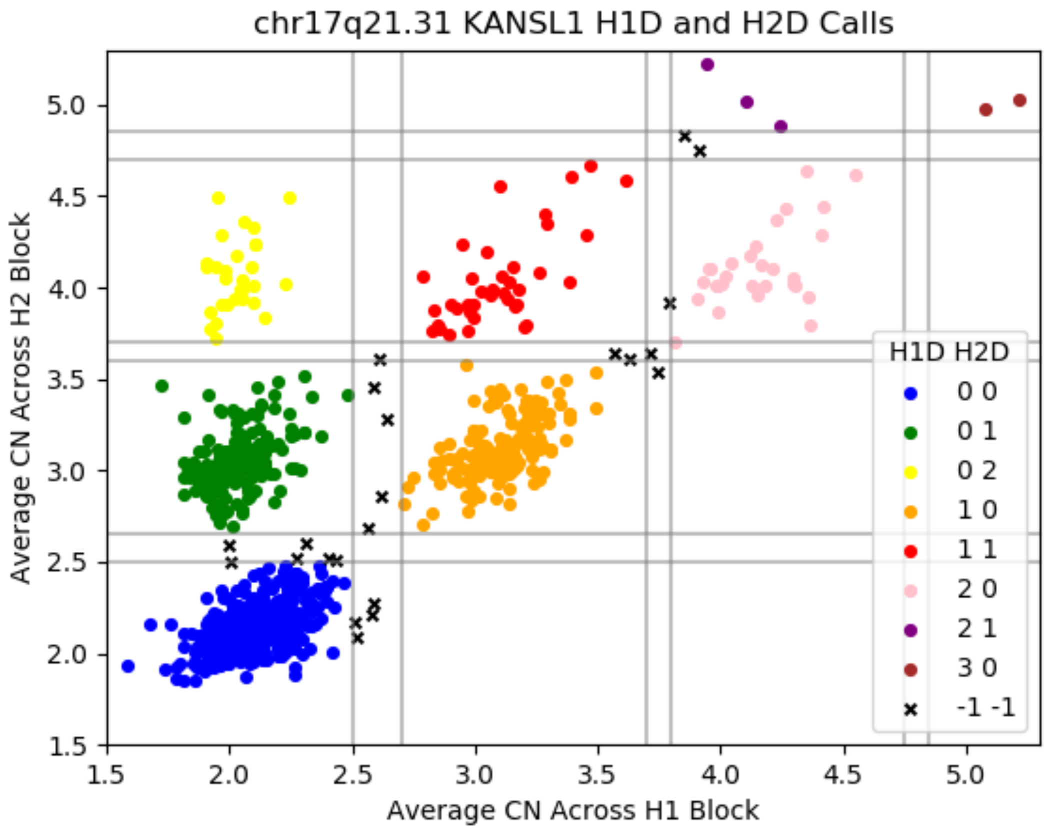


**Figure S102.** *KANSL1* H1D and H2D calls grouped by genotype. The samples marked with an ‘x’ are considered ambiguous as they are between groups.

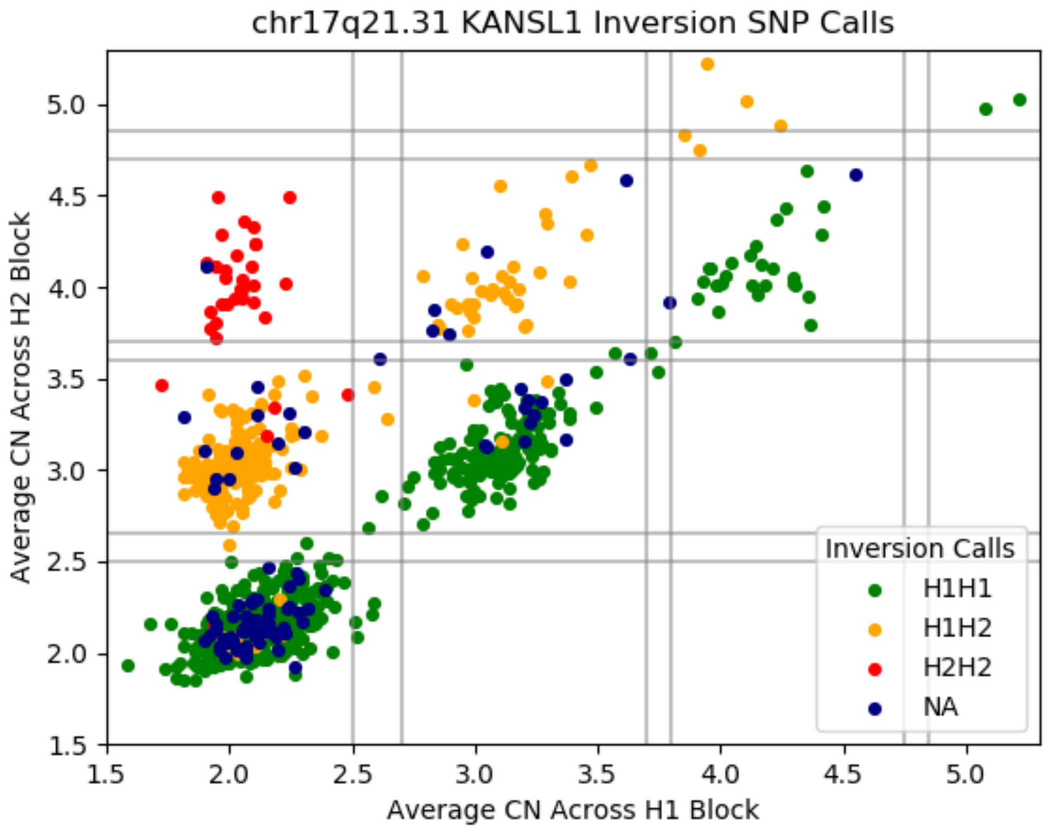


**Figure S103.** *KANSL1* H1 and H2 inversion calls mapped atop H1D and H2D average copy number values. The inversion and duplication calls align.

#### Results

Our read depth analyses were able to genotype the duplication status of both haplotypes across ancient samples. These results were able to identify H1 and H2 duplications in samples older than 8000 BP and trace the spread of duplicated and unduplicated haplotypes through time in Europe.

#### References

1. Stefansson, H. et al. A common inversion under selection in Europeans. Nature Genetics 37, 129–137 (2005).
2. Zody, M. C. et al. Evolutionary toggling of the MAPT 17q21.31 inversion region. Nat Genet 40, 1076–1083 (2008).

3. Boettger, L. M., Handsaker, R. E., Zody, M. C. & McCarroll, S. A. Structural haplotypes and recent evolution of the human 17q21.31 region. Nat Genet 44, 881–885 (2012).
4. Steinberg, K. M. et al. Structural diversity and African origin of the 17q21.31 inversion polymorphism. Nat Genet 44, 872–880 (2012).

### 9) Calculating ancestral contributions to modern complex phenotypes

William Barrie[1], Alba Refoyo Martínez[2], Astrid K. N. Iversen[3] and Dan Lawson[4]

[1] Zoology Department, University of Cambridge, UK.
[2] Lundbeck Foundation GeoGenetics Centre, GLOBE Institute, University of Copenhagen, Copenhagen, Denmark
[3] Oxford Centre for Neuroinflammation, Nuffield Department of Clinical Neurosciences, John Radcliffe Hospital, University of Oxford, Oxford, UK.
[4] School of Mathematics and Integrative Epidemiology Unit, University of Bristol, UK.

#### Introduction

Most studies that look at polygenic risk scores in ancient populations use genotypes of ancient individuals, combined with effect sizes from modern GWAS studies, to reconstruct risk scores for ancient individuals[1]. This involves exporting effect sizes across space and time, which is known to dramatically reduce the accuracy of the estimates[2]. Additionally, these scores are usually impossible to verify (except with specific phenotypes such as height where calibration is possible [3,4]), and don't necessarily measure what an ancient population contributed to phenotypic diversity in a modern population(s), especially when there has been selection or bottleneck events in between.

Here, we aim to use local ancestry information resulting from painting the UK Biobank (UKB) (Supplementary Note 2) to estimate ancestral contributions to modern complex phenotypes, by calculating polygenic risk scores for each ancestry based on local painting results. This is a well-powered approach due to the large modern sample size and is a more direct measure of the variants that a given ancestry contributed to the "white British" genetic landscape. Thus, we can draw conclusions about the differing contributions of each ancestry to modern genetic risk, whether due to drift or selection. We use bootstrapping to test whether some ancestries are significantly and systematically over-represented for a phenotype, indicating selection. Additionally, we look at the ancestral haplotypic background of two high effect variants of the *APOE* gene (ApoE2 and ApoE4), which modulate risk for Alzheimer's Disease [5,6].

#### Methods

##### Trait ascertainment

Instead of exhaustively calculating ancestry specific polygenic risk scores for all UKB traits, we firstly tested for evidence of overdispersion in polygenic scores between different ancient populations. We used effect size estimates from the UK Biobank Neale lab GWAS[7], and used 1,703 non-overlapping and approximately independent linkage disequilibrium (LD) blocks[8]. From the UKB case-control traits, we filtered out those where the N-case/N-control ratio is lower than 1%, and we exclude all traits that did not have any associated variants at the standard genome-wide significance threshold ($p < 5e-8$). We then filtered out variants with minor allele frequency lower than 5%, variants with INFO score < 50%, variants with a genotype probability lower than 0.8 in more than 10% of the individuals, triallelic variants and variants flagged as "low-confident" after imputation. From each LD block, we retrieved the variant with the lowest association p-value. We then selected only the variants with a p-value lower than the genome-wide significance threshold for downstream analyses.

Polygenic scores were calculated by summing over the allele frequency of the filtered trait-associated variants and weighting them by the effect size obtained from the summary statistics from the UK Biobank. The allele frequency was retrieved from the imputed ancient West-Eurasian individuals. Individuals were clustered into groups of closely related individuals using MDS. In our overdispersion test, we did not include genomes that were not successfully classified as belonging to any of the clusters and removed clusters that contained less than four individuals. This resulted in 1,119 individuals clustered into 41 groups.

To compute the $Q_X$ statistic, an empirical genome-wide covariance matrix is needed, which we constructed using a subset of SNPs with a trait-association p-value larger than 5e-8, and then we sampled every 20th "non-associated" SNPs across the genome. To further test the significance of the overdispersion and account for possible deviations from the assumptions the $Q_X$ statistic makes, we also computed p-values using two randomization schemes simulating a neutral scenario. The first method was based on the randomization of the signs of the effect size estimates of trait-associated SNPs while the other was based on sampling variants across the genome with frequencies matching those of trait-associated variants in the GBR panel from the 1000 Genomes Project.

To address mapping biases, we performed a one-tailed Wilcoxon rank-sum test for each trait. We evaluated if the candidate associated SNPs had higher values of the artefactual effect estimates than the non-associated SNPs used as our neutral baseline. None of the tests were significant (min-P value = 0.19; max-P Value = 0.99; Mean=0.56).

#### Calculating ancestral risk scores

To calculate ancestry specific polygenic risk scores, we used the same SNP ascertainment as for the $Q_X$ statistic and used these SNPs to calculate polygenic risk scores for each ancestry, using ancestry-specific 'effect allele frequencies' derived from the painting.

In order to calculate the effect allele frequency for a given ancestry $f_{\{anc,i\}}$ we used the formula:

$$f_{\{anc,i\}} = \frac{\sum_j^M \quad Painting\ certainty_{\{j,effect\}}}{\sum_j^M \quad Painting\ certainty_{\{j,effect\}} + \sum_j^M \quad Painting\ certainty_{\{j,alt\}}}$$

Where there are M individuals, and $\sum_j^M \quad Painting\ certainty_{\{j,effect\}}$ is the sum of the painting probabilities for that ancestry of all effect alleles. This calculates an effect allele frequency for an ancestry which is weighted by the painting probabilities: if a haplotype with the effect allele was painted with low probability for that ancestry, it will contribute little to the calculation, and vice versa. One benefit of this approach is that because it only matters how effect alleles are painted relative to alternate alleles for an ancestry group, and differences in genome-wide painting averages between ancestries will not cause bias.

To calculate an ancestry-specific PRS we used an additive model, including a transformation as in Berg & Coop[9], which converts scores to standard deviations from a pan-ancestry mean (i.e., z-scores). We derived standard deviations for each score by running a block bootstrap (1000 iterations) on (1) loci and (2) individuals. We calculated ancestry specific polygenic risk scores for 39 UKB traits shown to be significantly over-dispersed across ancient populations beyond what would be expected under a null model of genetic drift. For computational reasons, we used a random batch of 48,000 painted individuals to calculate the effect allele frequencies, which is sufficiently large to approximate the frequencies even for ancestries that are painted less.

Our calculations were limited to the 549,323 SNPs used in the painting of the UKB (Supplementary Note 2). This is expected to reduce predictive power compared to using the full set of imputed SNPs in the UKB, but only slightly[10]. There was a ~15% decrease in the number of SNPs included per phenotype in the PRS calculation compared with the imputed data.

To test the ancestral backgrounds of ApoE2 and ApoE4, we calculated the average painting score for each ancestry at all sites on the chromosome of haplotypes containing the effect allele. This makes it clear when there is an excess of a particular ancestry at the site of interest.

#### Results

Our results tell us about the ancestral contribution to modern phenotypes in the white British population (Figure S104, Figure S105), and we stress we are not making claims about the phenotypes of ancient populations.

We find that Yamnaya, CHG and EHG ancestral contributions (which together form a 'steppe' component) have relatively high scores for height, whereas Farmers and WHG ancestral contributions have relatively low scores. This accords with most previous studies [3,11,12] but not all[13]. EHG and Yamnaya both score highly for body mass and basal metabolic rate.

Hair and skin pigmentation show significant differences between the ancestral contributions, with risk scores for skin colour for the three hunter-gatherer ancestries being higher (i.e. darker) than Farmer and Steppe (as in [14]). On the other hand, traits related to malignant neoplasms of skin show higher scores for the Farmer ancestral contribution; while Farmer and Yamnaya ancestral contributions have higher scores for blonde and light brown hair, with the hunter-gatherer ancestries showing higher scores for dark brown. CHG is the only ancestral contribution which stands out as having a high risk score for black hair.

Intriguingly, the WHG ancestral component has strikingly high scores for traits related to cholesterol, blood pressure and diabetes, both when bootstrapping individuals and loci. In terms of psychiatric traits, the Farmer component scores highest for anxiety, guilty feelings, and irritability.

Our two bootstrapping methods mean slightly different things. Individuals in the UKB are related through shared genealogies, and so by bootstrapping over non-independent individuals (Figure S104) we are testing the consistency of the signal within the population. From this bootstrapping exercise we can conclude whether a difference in allele frequencies in ancient populations contributed to phenotypic variation today. Unsurprisingly, with a large enough sample size most phenotypes will show differences in ancestral contributions for this, usually due to drift or founder effects. However, this goes further than just reporting risk scores for ancient populations, because we are looking directly at coalescent tracts in the British population. We can conclude that “ancestry X contributes higher genetic risk for phenotype Y in the test population”. On the other hand, because we have used independent LD blocks to select SNPs to include in the PRS calculation, the requirement for independence is met when we bootstrap with loci (Figure S105). A positive result here is therefore much stronger, showing a systematic over/under-representation of an ancestry at loci affecting a given trait, beyond what is expected given the correlation among individuals. This points towards selection as an explanation.

The effect/risk allele (rs429358_C, n=127,760) of ApoE4 is preferentially painted as WHG/EHG, with a clear depletion of other ancestries (especially Farmer) at this locus compared to the genome-wide average (Figure S106). We replicated this result using another tag SNP for ApoE4 (rs429358_G, n=156,175, Figure S107). This indicates that the ApoE4 allele was contributed at least in part by hunter-gatherer ancestry into modern (British) populations, above what we would expect by chance. On the other hand, we found the ApoE2 allele (rs7412_T, n=66,362, which decreases risk for Alzheimer's disease) on a haplotypic background with affinities to Steppe pastoralists (Figure 2f.5).

#### Discussion

The methods here directly link genetic contributions from pre-defined ancestries to complex phenotypes in modern people. For most traits, each ancestry contributed differently to the modern genetic landscape, with some conveying enhanced or reduced risk either due to drift (including population bottlenecks/founder events) or selection. Because gradients exist in these ancestries across the British Isles and further afield (Supplementary Note 2), these differing risk scores indicate how geographically heterogeneous ancestry distributions may contribute to differing genetic risk profiles, in addition to other factors such as geography, socio-economic status etc.

A caveat for all studies involving polygenic risk calculation is that they rely on effect size estimates from an original GWAS which may be affected by population stratification in the GWAS panel, even when it has apparently been controlled for. This seems to be less of a problem in the UKB than in previous GWAS studies[15] but should be kept in mind. One benefit of our approach is that there is no requirement to export these risk scores across time and space: we are using effect sizes estimated from the modern population to calculate ancestral contributions to the same modern population.

ApoE4 is an isoform of the APOE gene, resulting from linkage disequilibrium between two SNPs, rs429358 and rs7412[16], and associated with increased risk for metabolic, vascular, and neurodegenerative diseases in adulthood[17]. It may provide some enhanced cognitive ability in children and young adults[18] and other health and immunity benefits, particularly in environments with high pathogen and parasite loads[19]. The ε4 and ε2 alleles are associated with increased risk of recurrent pregnancy loss relative to the ε3 allele[20]. There are several lines of evidence suggesting a link between the evolution of diet and the ApoE isoforms: ε2 and ε3 alleles are associated with lower levels of blood cholesterol [21,22], while ε4 is associated with higher levels, leading some to speculate that the derived ε3 allele is 'meat-adaptive' [23,24]. In a study of South Americans, there was a fivefold increase in the ApoE4 allele in hunter-gatherers versus horticulturalists[25], potentially because the immune benefits outweighed the advantages of low blood cholesterol[26]. Generally, ε4 prevalence is higher in indigenous foraging groups such as the Pygmies, Khoi San, Papuans and some Native Americans, while ε3 is most frequent in populations with a long-established agricultural economy[27]. Finally, ApoE4 is implicated in higher blood vitamin D levels[28].

The ε4 variant has been shown to be ancestral in humans[29]. There is a linear increasing trend in ε4 prevalence from South to North in Europe, with Sardinians showing the lowest prevalence [30–32], while there is a more than two-fold increase in Nordic versus Mediterranean countries[33]. Sardinians are an unusual population, having the highest level of neolithic farmer ancestry of all modern European populations[34]. In this light, differences in genome-wide ancestry proportions between northern (high WHG/EHG, low Farmer) and southern Europe (high Farmer, low WHG/EHG) (Supplementary Note 2) may explain at least part of the differences in frequency of the ε4 variant and subsequent AD genetic risk.

##### ApoE2 and its association with protection against severe malaria

*Plasmodium vivax*, an ancient and endemic malaria parasite, is believed to be the oldest among the four species that cause human malaria. It is thought to have evolved in Africa and spread globally through the Out of Africa migration about 60,000 YBP whereas *P. falciparum* appears to have originated in Africa about 10,000 YBP, although the timing of this event is under discussion [35,36]. Unlike the other three plasmodium parasites that infect humans (*P.*

*falciparum*, *P. malariae*, and *P. ovale*), *P. vivax* can survive at lower temperatures within mosquitoes, making it well-adapted to thrive in temperate, tropical, and subtropical regions. Therefore, it is likely that *P. vivax w*as the predominant plasmodium parasite in Europe and the Pontic-Caspian steppe during the Neolithic period and Bronze Age. The presence of malaria-like periodic fevers is evident in ancient writings, for example on clay tablets with cuneiform script from Mesopotamia (~5500-1800 YBP), in Indian writings of the Vedic period (~3500-2800 YBP), and in the Greek poet Homer's 'The Iliad" (~2750 YBP)[37]. Moreover, the plasmodium antigen has been detected in samples from Egyptian Mummies dating from 5200 and 3304 YBP[38] and *P. falciparum* DNA in an Egyptian Mummy dating ~4000 YBP[39]. *P. falciparum*, the most pathogenic form of malaria, is believed to have spread rapidly from Africa to other tropical and subtropical regions of the world only within the last 6,000 years, with evidence suggesting its arrival in Rome approximately 2000 years ago[40].

Malaria has exerted strong selective pressure throughout human history, leading to several independent adaptations in the human genome, particularly in regions where *P. falciparum* now predominates, such as Africa and Southeast Asia[41]. A unique characteristic of *P. vivax* is its requirement for the Duffy antigen on the surface of cells to invade red blood cells. As a result, its prevalence is currently much lower than *P. falciparum* in Africa where the population has low expression of the Duffy antigen, possibly due to earlier and long-term strong selection pressures induced by *P. vivax*. Various red blood cell polymorphisms protect against severe falciparum malaria, including thalassemia syndromes, anaemias caused by abnormalities in genes encoding haemoglobin, sickle cell haemoglobin, and glucose-6-phosphate dehydrogenase (G6PD) deficiency, and haemoglobin C in Africa and haemoglobin E  in Southeast Asia[42].

*P. vivax* malaria is widely regarded as a benign form of the disease with a low case-fatality ratio. However, recent evidence suggests that this view may only partially be accurate, particularly in the case of children. *P. vivax* malaria can cause severe and debilitating illness in children, with a similar spectrum of symptoms to that seen in *P. falciparum* malaria, such as cerebral malaria, with a fatality rate ranging from 16-40%[43]. In a recent study conducted in Bikaner, a region where both *P. vivax* and *P. falciparum* coexist, researchers observed a higher incidence of severe manifestations in *P. vivax* monoinfection compared to *P. falciparum* monoinfection among children aged 0-5 years[43]. The child mortality in severe *P. vivax* malaria was 6.15% (4/65), whereas the child mortality in severe *P. falciparum* was 7.59% (6/79). These findings are consistent with previous studies in India[44], Indonesia[45],

Papua New Guinea[45], and Vanuatu[46], which have also reported severe cases of *P. vivax* malaria. A longitudinal study from Papua New Guinean children reported that by nine years of age, children had acquired almost complete clinical immunity to *P. vivax*[47]. In contrast, the acquisition of immunity to symptomatic *P. falciparum* malaria remained incomplete. Similar differences in rates of immunity acquisition have also been described in Vanuatu where both species coexist[46] and the reason for this difference in immunity is not known. The underlying mechanisms that make *P. vivax* as serious as *P. falciparum* in infants and young children are not fully understood.

The climate on the Pontic steppe has undergone multiple changes over the past 8000 years, with shifts between moist to arid climates, particularly in the southern regions [48]. *Anopheles atroparvus is a P. vivax* malaria vector[37] and it is currently widespread in Europe and surrounding areas, including North-Eastern Europe and the Pontic Steppe (https://www.ecdc.europa.eu/en/disease-vectors/surveillance-and-disease-data/mosquito-maps). The *A. atroparvus* larvae can be found in various environments, including fresh and saline water, but prefer brackish, sunlit water with high amounts of filamentous algae or floating vegetation. This species rests and hibernates in animal sheds and stables, and adult females are often found indoors in such human-created habitats. Climate change on the Pontic steppe might have impacted *A. atroparvus* numbers and hence malaria spread, for example increasing the mosquito population during moist conditions which also favoured population growth of the local Neolithic populations and expanded the northern occupation zone into southern steppe regions about 6000 YBP[48]. It is tempting to speculate that this moist period might have increased the selection of ApoE2 in a pre-Yamnaya population in the already relatively moist Northern parts of the Pontic Steppe or in the Danube Delta wetlands before their later dispersal into Europe. The approximate timing of that moist period 6000 YPB is in line with our pan-ancestry analysis demonstrating positive selection of ApoE2 starting about 7000 YBP. A long period of climate aridity occurred in Eastern Europe starting about 3000 YBP[48], which approximately coincides with the timing of no additional positive selection of ApoE2 in our CLUES analyses (Supplementary Note 4) and a stable frequency of this isoform in the pan-ancestral population going forward.

Compared to other isoforms, ApoE2 exhibits lower efficacy in competing with sporozoites to bind to their receptor on hepatic cells, decreasing the host's ability to resist malaria infection[49]. The ApoE2 isoform is a risk factor for infection with malaria at an early age[50] while the ApoE3/ApoE4 genotype is linked to an increased likelihood of severe infection in children[49]. This seemingly contradictory observation is explained by epidemiological studies: a correlation exists between the age at which children experience their first malaria infection and the likelihood of developing severe symptoms[51]. Specifically, children in regions with high endemicity, who contract the disease during infancy while still having protection from clinical malaria and high parasitaemia likely conferred by innate defences and maternal antibodies, appear less susceptible to severe malaria in later life[51].

This suggests that exposure to malaria during this critical developmental window may prime the adaptive immune system, conferring long-term protective immunity against severe malaria. In contrast, after the waning of protective mechanisms, children who contract malaria for the first time at an older age are fully vulnerable and face a heightened risk of a severe or fatal disease. Consequently, ApoE2 carriers would have a selective advantage in a region with high endemicity as they would be more likely to be infected as infants and develop an immune response that protected them against severe disease later.

##### ApoE isoforms and hepatitis C infection

Conflicting results have arisen from studies investigating the relationship between hepatitis C virus (HCV) infection and Apolipoprotein E (ApoE) isoforms [52–54]. In one study, a protective effect against HCV infection was observed in Caucasian HCV patients homozygous for the ε2 allele, compared to healthy controls; this suggests that the ApoE ε2/ε2 genotype may provide some protection against HCV infection[53]. In addition, individuals with the ε3/ε3 genotype were found to have the highest risk of developing chronic HCV infection, with odds ratios of 0.39 and 0.59 for the ε2 and ε4 alleles, respectively, which supports the hypothesis that the ε2 and ε4 genotypes facilitate viral clearance. However, a separate study examining the association between ApoE genotype and the outcome of HCV infection among individuals with a chronic or cleared infection found no significant differences in ApoE allele frequencies between HCV-infected groups and controls[52]. Nevertheless, this study did find that individuals with the ε4 allele had less severe HCV-induced liver damage. Further investigations revealed that individuals with the ε4 allele had a lower risk of viral persistence[53] and chronic infection[54]. Taken together, these findings suggest that the ApoE4 isoform may protect HCV-infected patients from developing severe liver disease, while the protective effect of apoE2 is less conclusive.

#### ApoE isoforms and herpes simplex type 1 infection

The impact of HSV-1 on the host cell's protein synthesis has been widely studied, and while it typically results in an overall reduction of protein synthesis, it has also been found to trigger the production and accumulation of beta-amyloid, a hallmark of Alzheimer's disease (AD)[55]. The susceptibility to HSV-1 infection has not been found to be affected by the ApoE genotype[54]. However, studies of elderly individuals found that those carrying the ε4 allele of the ApoE gene had a higher frequency of HSV-1-positive AD patients than HSV-1-negative AD patients and non-AD patients, indicating a several-fold increase in AD risk for subjects with both the ε4 allele and HSV-1 infection compared to non-ε4 carriers [56,57]. These findings suggest that while HSV-1 infection alone does not necessarily lead to AD, the combination of HSV-1 infection and the ApoE4 genotype may substantially increase the risk of developing AD in older adults [56–58]. This may partly be due to age-related declines in immune or barrier function, which can facilitate viral entry into the central nervous system (CNS), as HSV-1 has not been detected in the brains of younger individuals[56]. Notably, the ApoE4 genotype has also been shown to increase the risk of HSV-1-induced cold sores in peripheral nervous tissue[56]. While the literature is not entirely consistent, likely due to small sample sizes and inadequate statistical power, current evidence suggests that ApoE4 may increase susceptibility to HSV-1-related herpes labialis and neuroinvasiveness compared to other ApoE forms, and that the combination of ApoE4 and HSV-1 may increase the risk of AD[59].

#### ApoE isoforms and coronavirus infection

The impacts of ApoE isoforms on severe acute respiratory virus 2 (SARS-CoV-2) infection risk, disease progression, and mortality are under investigation. One study has linked ApoE2 to a decrease in the risk of SARS-CoV-2 infection but not to the severity of the disease[60]. Several other papers have linked ApoE4 to an increased risk of infection and more severe disease [61–63]. In addition, ApoE4 has been linked to microvascular damage in the brain and increased neuroinflammation, with some pathways overlapping those activated in Alzheimer's[64], suggesting SARS-CoV-2 infection might work as a dementia disease accelerator, specifically in those suffering from - or predisposed to - Alzheimer's dementia. No studies appear to have investigated the link between ApoE isotypes and other human coronaviruses according to PubMed searches (HCoV-229E, HCoV-OC43, HCoV-HKU1, HCoV-NL63, MERS-CoV, and SARS-CoV-1). Taken together, these somewhat incomplete results suggest that it is possible ApoE2 might have reduced the risk of infection with a SARS-CoV-2-like coronavirus and might thus have been positively selected for in regions of high endemicity of this (and possibly several) coronaviruses. However, this suggestion is highly speculative due to the lack of data.

#### Figures

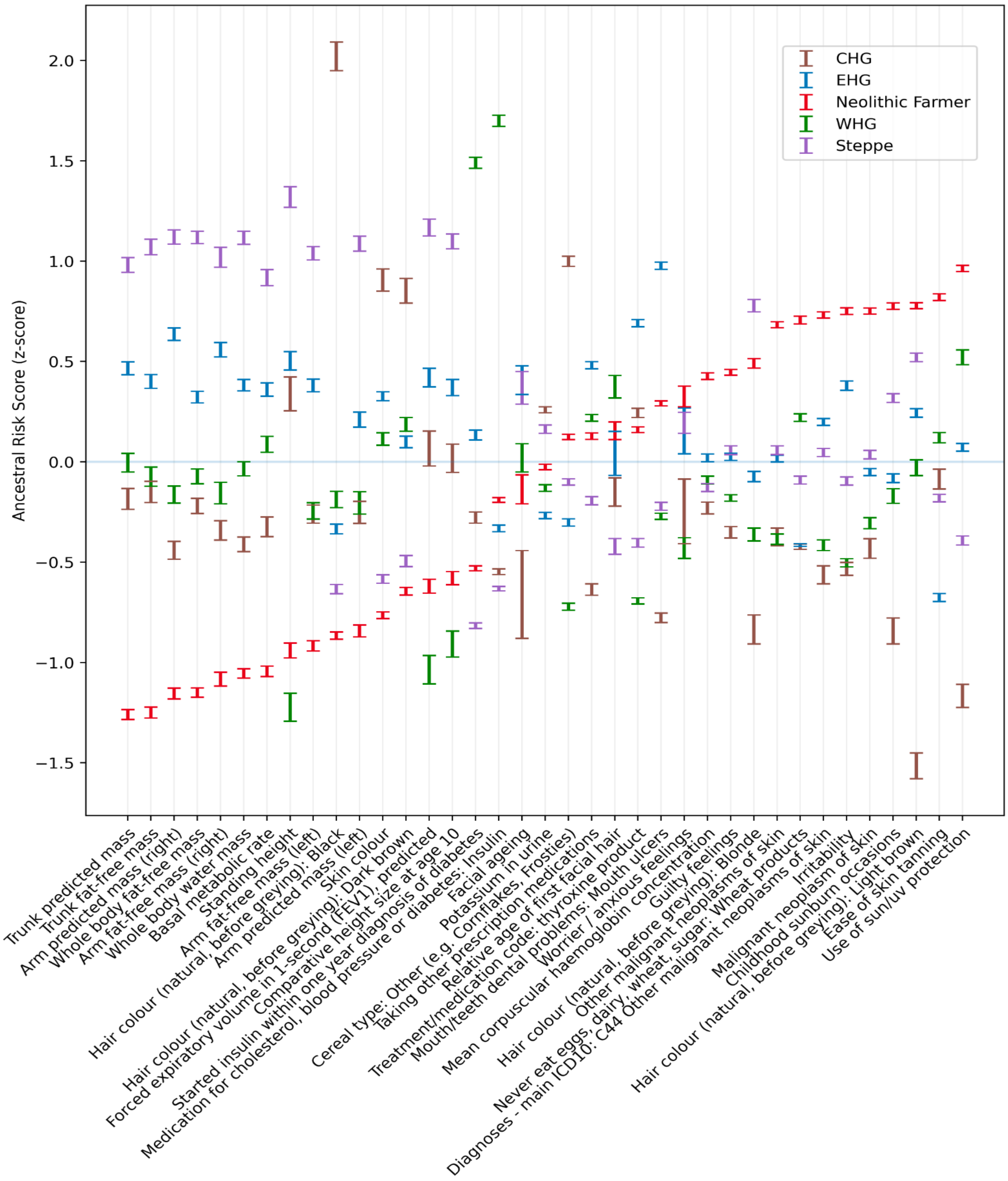


**Figure S104.** Ancestry-specific polygenic risk scores with 95% confidence intervals, centred on the mean, derived from bootstrapping individuals for phenotypes shown to be significantly over-dispersed between ancient populations. Confidence intervals were calculated by re-running PRS calculation on random batches of 48,000 individuals, with replacement (1000 iterations), while keeping all other annotations intact. Here we show 2 x standard deviation error bars, expected to represent ~95% confidence interval under a normal distribution. Bootstrapping individuals tests the extent to which ancestry X contributed higher genetic risk for phenotype Y in a given population, either due to drift or selection. Summary data used to make this figure can be found in Supplementary Table 12.

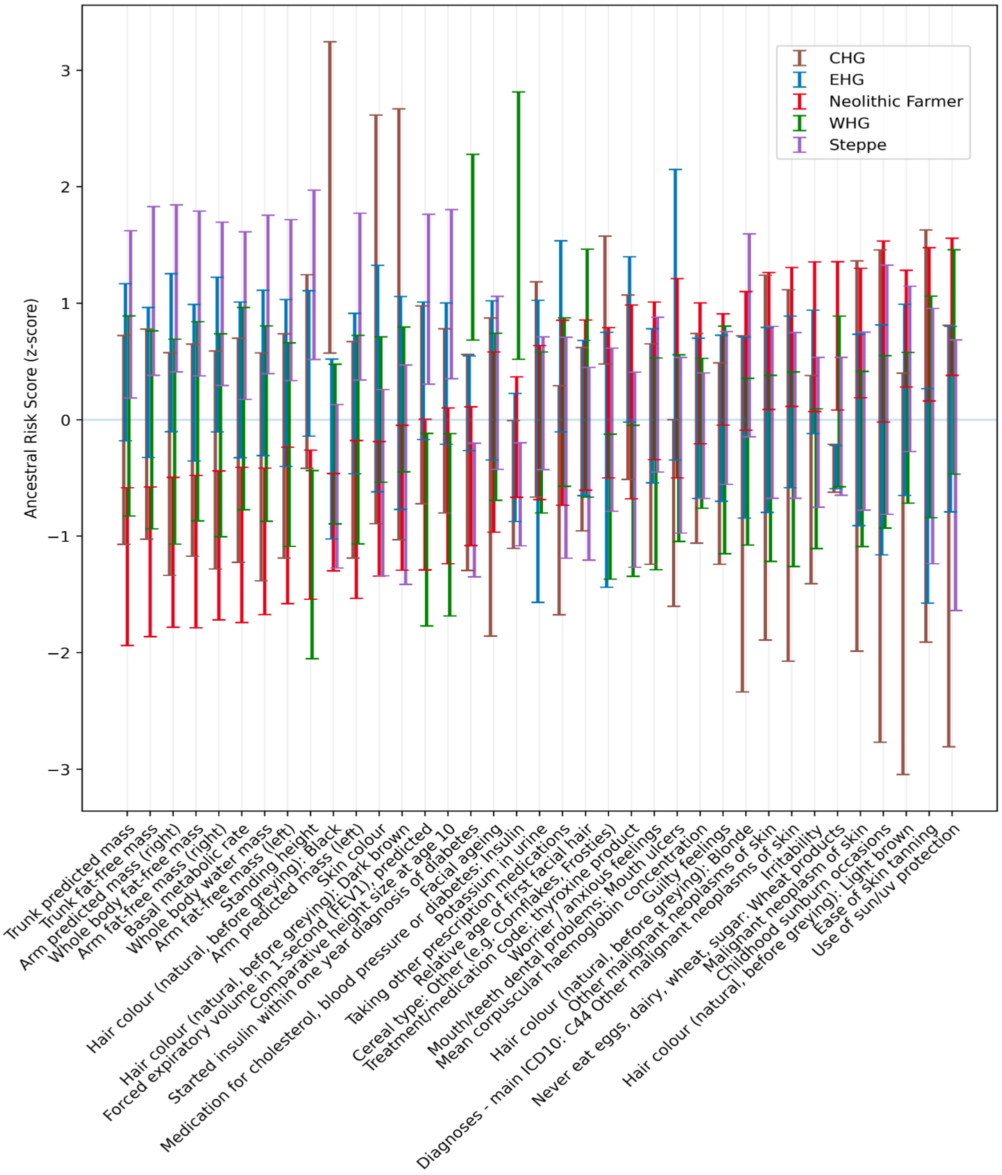


**Figure S105.** Ancestry-specific polygenic risk scores for present-day samples (n=408,884) with 95% confidence intervals, centred on the mean, derived from bootstrapping loci for phenotypes shown to be significantly over-dispersed between ancient populations. Confidence intervals were calculated by bootstrapping independent loci from separate LD blocks (1000 iterations), while keeping all other annotations intact. Here we show 2 x standard deviation error bars, expected to represent ~95% confidence interval under a normal distribution. Bootstrapping loci tests whether there is a systematic bias towards an ancestry for a given phenotype across all significant SNPs, possibly indicating selection. Summary data used to make this figure can be found in Supplementary Table 13.

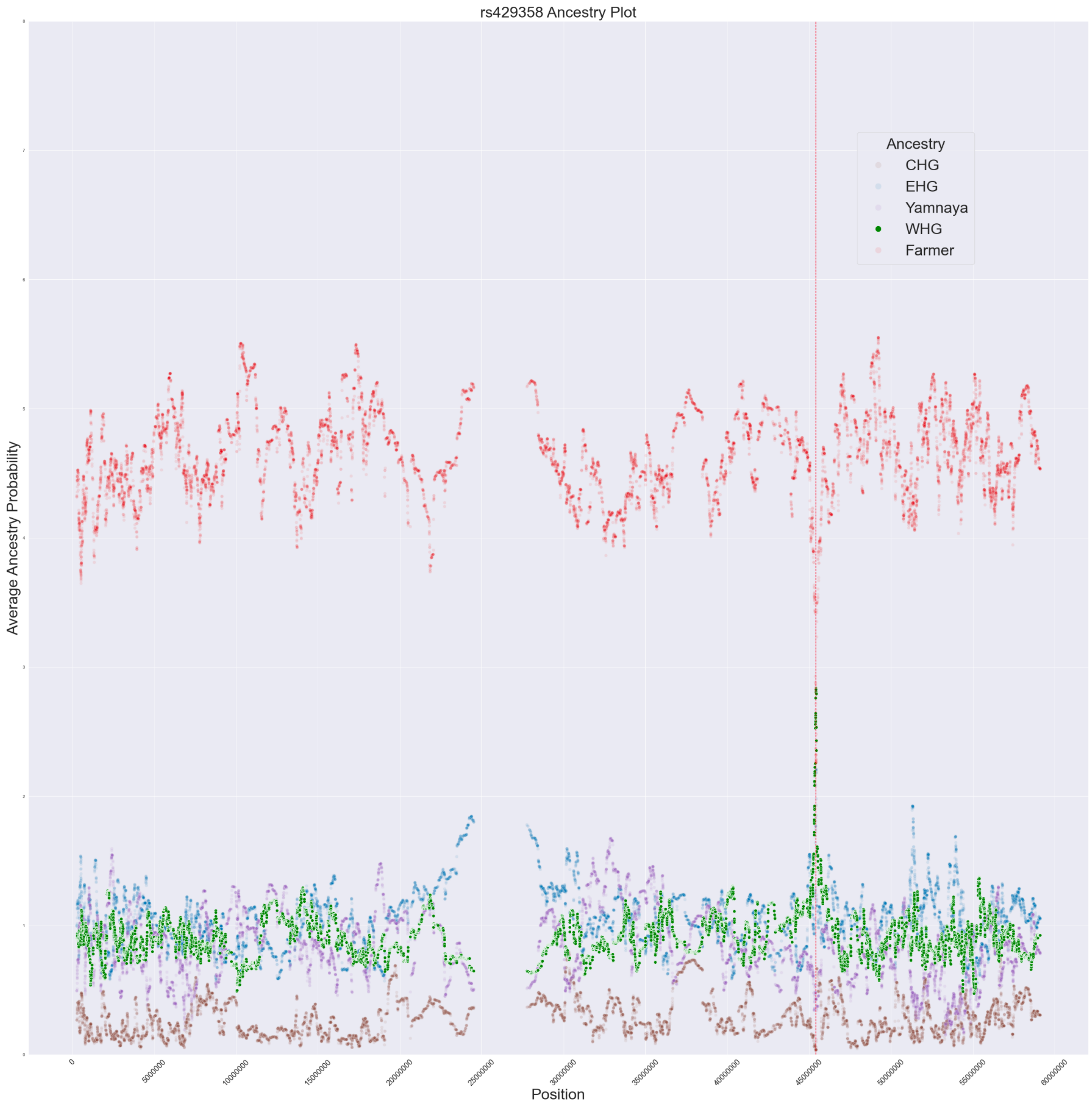


**Figure S106.** Average painting score for each ancestry at all sites on chromosome 19 of haplotypes containing the effect allele for ApoE4 (rs429358_C, n=127,760). The vertical red line indicates the position of the SNP of interest. There is an excess of WHG/EHG ancestry and a depletion of Farmer ancestry at this locus.

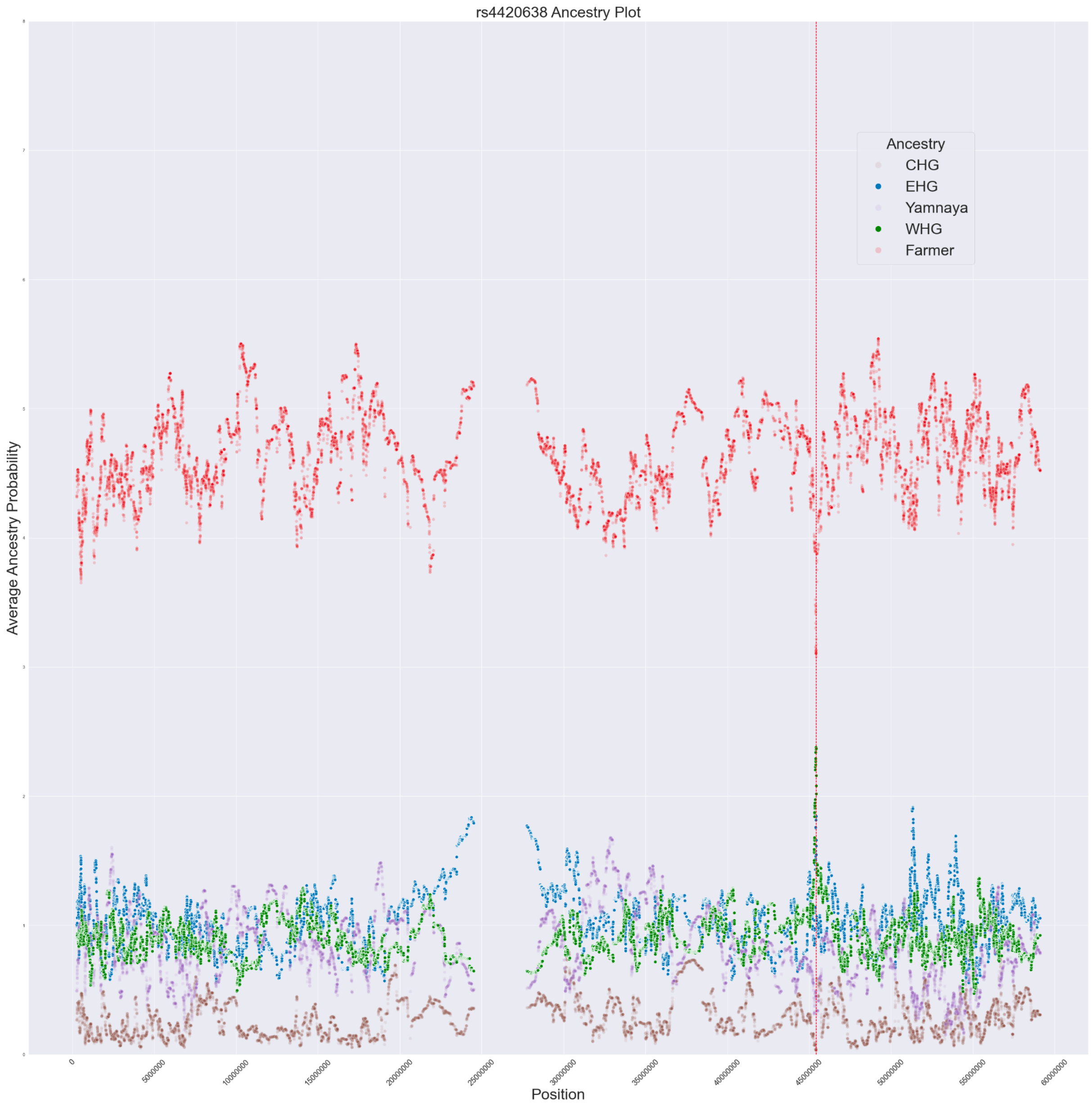


**Figure S107.** Average painting score for each ancestry at all sites on chromosome 19 of haplotypes containing the effect allele for ApoE4 using a tag SNP (rs4420638_G, n=156,175). The vertical red line indicates the position of the SNP of interest. There is an excess of WHG/EHG ancestry and a depletion of Farmer ancestry at this locus.

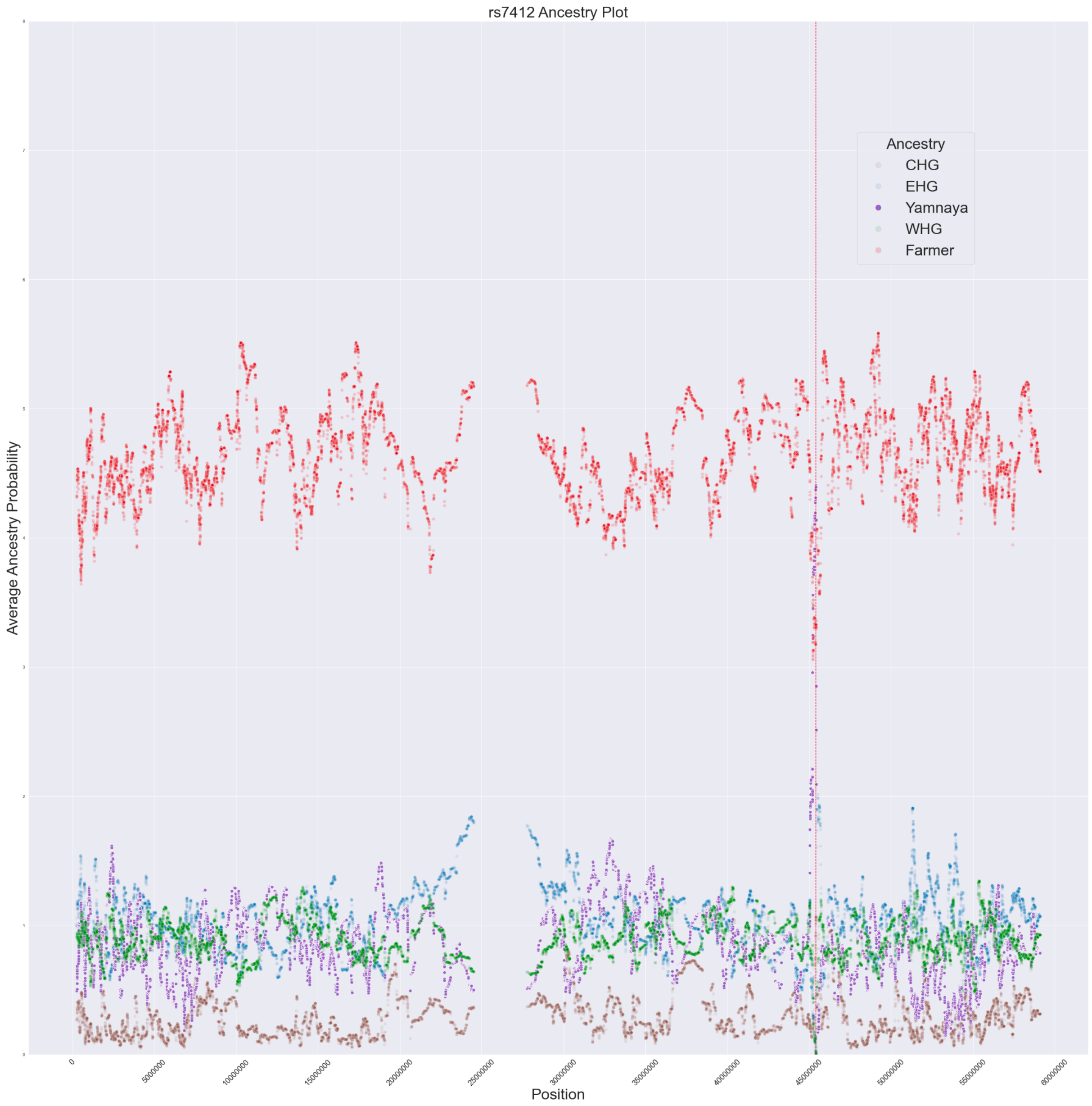


**Figure S108.** Average painting score for each ancestry at all sites on chromosome 19 of haplotypes containing the effect allele for ApoE2 using a tag SNP (rs7412_T, n=66,362). The vertical red line indicates the position of the SNP of interest. There is an excess of Yamnaya/Steppe ancestry and a depletion of Farmer ancestry at this locus.

#### References

1. Irving-Pease, E. K., Muktupavela, R., Dannemann, M. & Racimo, F. Quantitative Human Paleogenetics: What can Ancient DNA Tell us About Complex Trait Evolution? *Front. Genet.* **12**, 703541 (2021).

2. Duncan, L. *et al.* Analysis of polygenic risk score usage and performance in diverse human populations. *Nat. Commun.* **10**, 3328 (2019).

3. Cox, S. L., Ruff, C. B., Maier, R. M. & Mathieson, I. Genetic contributions to variation in human stature in prehistoric Europe. *Proc. Natl. Acad. Sci.* **116**, 21484–21492 (2019).

4. Cox, S. L., Moots, H., Stock, J. T., Shbat, A. & Bitarello, B. D. Predicting skeletal stature using ancient DNA. 20.

5. Corder, E. H. *et al.* Gene Dose of Apolipoprotein E Type 4 Allele and the Risk of Alzheimer's Disease in Late Onset Families. *Science* **261**, 921–923 (1993).

6. Strittmatter, W. J. *et al.* Apolipoprotein E: high-avidity binding to beta-amyloid and increased frequency of type 4 allele in late-onset familial Alzheimer disease. *Proc. Natl. Acad. Sci.* **90**, 1977–1981 (1993).

7. Bycroft, C. *et al.* The UK Biobank resource with deep phenotyping and genomic data. *Nature* **562**, 203–209 (2018).

8. Berisa, T. & Pickrell, J. K. Approximately independent linkage disequilibrium blocks in human populations. *Bioinformatics* btv546 (2015) doi:10.1093/bioinformatics/btv546.

9. Berg, J. J. & Coop, G. A Population Genetic Signal of Polygenic Adaptation. *PLoS Genet.* **10**, e1004412 (2014).

10. Choi, S. W. & O'Reilly, P. F. PRSice-2: Polygenic Risk Score software for biobank-scale data. *GigaScience* **8**, giz082 (2019).

11. Mathieson, I. *et al.* Genome-wide patterns of selection in 230 ancient Eurasians. *Nature* **528**, 499–503 (2015).

12. Martiniano, R. *et al.* The population genomics of archaeological transition in west

Iberia: Investigation of ancient substructure using imputation and haplotype-based methods. *PLOS Genet.* **13**, e1006852 (2017).

13. Marnetto, D. *et al. Ancestral contributions to contemporary European complex traits*. http://biorxiv.org/lookup/doi/10.1101/2021.08.03.454888 (2021) doi:10.1101/2021.08.03.454888.
14. Ju, D. & Mathieson, I. The evolution of skin pigmentation-associated variation in West Eurasia. *Proc. Natl. Acad. Sci.* **118**, e2009227118 (2021).
15. Berg, J. J. *et al.* Reduced signal for polygenic adaptation of height in UK Biobank. *eLife* **8**, e39725 (2019).
16. Rall, S. C., Weisgraber, K. H. & Mahley, R. W. Human apolipoprotein E. The complete amino acid sequence. *J. Biol. Chem.* **257**, 4171–4178 (1982).
17. de-Almada, B. V. P. *et al.* Protective effect of the APOE-e3 allele in Alzheimer's disease. *Braz. J. Med. Biol. Res.* **45**, 8–12 (2012).
18. Tuminello, E. R. & Han, S. D. The Apolipoprotein E Antagonistic Pleiotropy Hypothesis: Review and Recommendations. *Int. J. Alzheimers Dis.* **2011**, 1–12 (2011).
19. Oriá, R. B., Patrick, P. D., Blackman, J. A., Lima, A. A. M. & Guerrant, R. L. Role of apolipoprotein E4 in protecting children against early childhood diarrhea outcomes and implications for later development. *Med. Hypotheses* **68**, 1099–1107 (2007).
20. Lumsden, A. L., Mulugeta, A., Zhou, A. & Hyppönen, E. Apolipoprotein E (APOE) genotype-associated disease risks: a phenome-wide, registry-based, case-control study utilising the UK Biobank. *eBioMedicine* **59**, (2020).
21. Petkeviciene, J. *et al.* Associations between Apolipoprotein E Genotype, Diet, Body Mass Index, and Serum Lipids in Lithuanian Adult Population. *PLoS ONE* **7**, e41525 (2012).
22. Carvalho-Wells, A. L. *et al.* Interactions between age and apoE genotype on fasting and postprandial triglycerides levels. *Atherosclerosis* **212**, 481–487 (2010).

23. Finch, C. E. & Stanford, C. B. Meat-Adaptive Genes and the Evolution of Slower Aging in Humans. *Q. Rev. Biol.* **79**, 3–50 (2004).
24. Allen, J. S., Bruss, J. & Damasio, H. The aging brain: The cognitive reserve hypothesis and hominid evolution. *Am. J. Hum. Biol.* **17**, 673–689 (2005).
25. Reales, G. *et al.* A tale of agriculturalists and hunter-gatherers: Exploring the thrifty genotype hypothesis in native South Americans. *Am. J. Phys. Anthropol.* **163**, 591–601 (2017).
26. Trumble, B. C. *et al.* Apolipoprotein E4 is associated with improved cognitive function in Amazonian forager-horticulturalists with a high parasite burden. *FASEB J.* **31**, 1508–1515 (2017).
27. CORBO, R. M. & SCACCHI, R. Apolipoprotein E (APOE) allele distribution in the world. Is APOE*4 a 'thrifty' allele? *Ann. Hum. Genet.* **63**, 301–310 (1999).
28. Huebbe, P. *et al.* *APOE* ε *4* is associated with higher vitamin D levels in targeted replacement mice and humans. *FASEB J.* **25**, 3262–3270 (2011).
29. Fullerton, S. M. *et al.* Apolipoprotein E Variation at the Sequence Haplotype Level: Implications for the Origin and Maintenance of a Major Human Polymorphism. *Am. J. Hum. Genet.* **67**, 881–900 (2000).
30. CORBO, R. M., SCACCHI, R., MUREDDU, L., MULAS, G. & ALFANO, G. Apolipoprotein E polymorphism in Italy investigated in native plasma by a simple polyacrylamide gel isoelectric focusing technique. Comparison with frequency data of other European populations. *Ann. Hum. Genet.* **59**, 197–209 (1995).
31. Adler, G. *et al.* Bosnian study of *APOE* distribution (BOSAD): a comparison with other European populations. *Ann. Hum. Biol.* **44**, 568–573 (2017).
32. Lucotte, G., Loirat, F. & Hazout, S. Pattern of gradient of apolipoprotein E allele asterisk4 frequencies in Western Europe. *Hum. Biol.* **69**, 253–62 (1997).
33. Trumble, B. C. & Finch, C. E. The Exposome in Human Evolution: From Dust to

Diesel. *Q. Rev. Biol.* **94**, 333–394 (2019).

34. Chiang, C. W. K. *et al.* Genomic history of the Sardinian population. *Nat. Genet.* **50**, 1426–1434 (2018).

35. Prugnolle, F. *et al.* A Fresh Look at the Origin of Plasmodium falciparum, the Most Malignant Malaria Agent. *PLOS Pathog.* **7**, e1001283 (2011).

36. Carter, R. Speculations on the origins of Plasmodium vivax malaria. *Trends Parasitol.* **19**, 214–219 (2003).

37. Medicine, I. of. *Saving Lives, Buying Time: Economics of Malaria Drugs in an Age of Resistance*. (The National Academies Press, 2004). doi:10.17226/11017.

38. Miller, R. L. *et al.* Diagnosis of Plasmodium falciparum infections in mummies using the rapid manual ParaSight$^{TM}$-F test. *Trans. R. Soc. Trop. Med. Hyg.* **88**, 31–32 (1994).

39. Nerlich, A. G., Schraut, B., Dittrich, S., Jelinek, T. & Zink, A. R. Plasmodium falciparum in Ancient Egypt. *Emerg. Infect. Dis.* **14**, 1317–1319 (2008).

40. Sallares, R., Bouwman, A. & Anderung, C. The Spread of Malaria to Southern Europe in Antiquity: New Approaches to Old Problems. *Med. Hist.* **48**, 311–328 (2004).

41. Kwiatkowski, D. P. How Malaria Has Affected the Human Genome and What Human Genetics Can Teach Us about Malaria. *Am. J. Hum. Genet.* **77**, 171–192 (2005).

42. Ha, J., Martinson, R., Iwamoto, S. K. & Nishi, A. Hemoglobin E, malaria and natural selection. *Evol. Med. Public Health* **2019**, 232–241 (2019).

43. Kochar, D. K. *et al.* Clinical Features of Children Hospitalized with Malaria—A Study from Bikaner, Northwest India. *Am. Soc. Trop. Med. Hyg.* **83**, 981–989 (2010).

44. Kochar, D. K. *et al.* Plasmodium vivax Malaria. *Emerg. Infect. Dis. J.* **11**, 132 (2005).

45. Tjitra, E. *et al.* Multidrug-Resistant Plasmodium vivax Associated with Severe and Fatal Malaria: A Prospective Study in Papua, Indonesia. *PLOS Med.* **5**, e128 (2008).

46. Maitland, K. *et al.* The interaction between Plasmodium falciparum and P. vivax in children on Espiritu Santo island, Vanuatu. *Trans. R. Soc. Trop. Med. Hyg.* **90**, 614–620

(1996).
47. MICHON, P. *et al.* THE RISK OF MALARIAL INFECTIONS AND DISEASE IN PAPUA NEW GUINEAN CHILDREN. *Am. J. Trop. Med. Hyg. Am J Trop Med Hyg* **76**, 997–1008 (2007).
48. Kotova, N. & Makhortykh, S. Human adaptation to past climate changes in the northern Pontic steppe. *Clim. Dyn. Prehist. Occup. Eurasian Perspect. Environ. Archaeol.* **220**, 88–94 (2010).
49. C Aucan, A J Walley, & A V S Hill. Common apolipoprotein E polymorphisms and risk of clinical malaria in the Gambia. *J. Med. Genet.* **41**, 21 (2004).
50. M A Wozniak *et al.* Does apolipoprotein E polymorphism influence susceptibility to malaria? *J. Med. Genet.* **40**, 348 (2003).
51. Gupta, S., Snow, R. W., Donnelly, C. A., Marsh, K. & Newbold, C. Immunity to non-cerebral severe malaria is acquired after one or two infections. *Nat. Med.* **5**, 340–343 (1999).
52. Wozniak, M. A. *et al.* Apolipoprotein E-ϵ4 protects against severe liver disease caused by hepatitis C virus. *Hepatology* **36**, 456–463 (2002).
53. D A Price *et al.* Apolipoprotein ε3 allele is associated with persistent hepatitis C virus infection. *Gut* **55**, 715 (2006).
54. Kuhlmann, I., Minihane, A. M., Huebbe, P., Nebel, A. & Rimbach, G. Apolipoprotein E genotype and hepatitis C, HIV and herpes simplex disease risk: a literature review. *Lipids Health Dis.* **9**, 8 (2010).
55. Wozniak, M. A., Itzhaki, R. F., Shipley, S. J. & Dobson, C. B. Herpes simplex virus infection causes cellular β-amyloid accumulation and secretase upregulation. *Neurosci. Lett.* **429**, 95–100 (2007).
56. Itzhaki, R. *et al.* Herpes simplex virus type 1 in brain and risk of Alzheimer's disease. *Lancet* **349**, 241–244 (1997).

57. Linard, M. *et al.* Interaction between APOE4 and herpes simplex virus type 1 in Alzheimer’s disease. *Alzheimers Dement.* **16**, 200–208 (2020).
58. Itabashi, S., Arai, H., Matsui, T., Higuchi, S. & Sasaki, H. Herpes simplex virus and risk of Alzheimer’s disease. *The Lancet* **349**, 1102 (1997).
59. Protto, V. *et al.* Role of HSV-1 in Alzheimer’s disease pathogenesis: A challenge for novel preventive/therapeutic strategies. *Curr. Opin. Pharmacol.* **63**, 102200 (2022).
60. Espinosa-Salinas, I. *et al.* Potential protective effect against SARS-CoV-2 infection by APOE rs7412 polymorphism. *Sci. Rep.* **12**, 7247 (2022).
61. Zhang, H. *et al.* APOE interacts with ACE2 inhibiting SARS-CoV-2 cellular entry and inflammation in COVID-19 patients. *Signal Transduct. Target. Ther.* **7**, 261 (2022).
62. Chen, F. *et al.* ApoE4 associated with severe COVID-19 outcomes via downregulation of ACE2 and imbalanced RAS pathway. *J. Transl. Med.* **21**, 103 (2023).
63. Kuo, C.-L. *et al.* APOE e4 Genotype Predicts Severe COVID-19 in the UK Biobank Community Cohort. *J. Gerontol. Ser. A* **75**, 2231–2232 (2020).
64. Rahman, M. A., Islam, K., Rahman, S. & Alamin, M. Neurobiochemical Cross-talk Between COVID-19 and Alzheimer’s Disease. *Mol. Neurobiol.* **58**, 1017–1023 (2021).

### 10) Pathogenic structural variants in ancient vs. modern-day humans

Alma S. Halgren[1], Andrés Ingason[2,3], and Peter H. Sudmant[1]

[1] Department of Integrative Biology, University of California, Berkeley
[2] Institute of Biological Psychiatry, Mental Health Services, Copenhagen University Hospital
[3] Lundbeck Foundation GeoGenetics Centre, GLOBE Institute, University of Copenhagen, Copenhagen, Denmark

#### Introduction

Rare, recurrent copy-number variants (CNVs) are known to cause neurodevelopmental disorders and are associated with a range of psychiatric and physical traits with variable expressivity and incomplete penetrance [1,2]. We examined 50 regions susceptible to recurrent CNV known to be the most prevalent drivers of human developmental pathologies [3] in 1442 ancient Eurasians and 1093 modern human populations (for comparison) to understand the prevalence of pathogenic structural variants over time.

#### Methods

This analysis examines 1442 ancient humans from primarily West Eurasia and Central Asia as well as 1093 publicly-available high-coverage modern human genomes encompassing 136 populations worldwide (from the Human Genome Diversity Project [4] and the Simons Genome Diversity Project [5]). In modern humans and 690 ancient individuals with fastq files available, paired-end Illumina reads were mapped to the human reference genome GRCh38 with BWA-MEM [6]. In the remaining 984 samples, only BAM files that had been mapped to hg19 were available. Of note, in these 984 samples, the filtering of duplicate reads resulted in the absence of signal over segmental duplications. Nonetheless, we were able to characterise structural variants intersecting unique sequences in these samples. The large putatively pathogenic loci which we focused on in this analysis generally consist of unique sequences flanked by segmental duplications. 232 samples were removed due to low coverage and genotype yield out of 1674 total ancient samples, leaving 1442 samples for the final analysis (601 from the fastq hg38 dataset and 841 from the BAM hg19 dataset). In all samples, average read depth in 1kb sliding genomic windows was extracted from the subsequent BAM files with pysamstats [7]. All alternate haplotypes were removed prior to mapping. To approximate copy number from read depth, we masked tandem repeats (Tandem Repeat Finder [8]), corrected read-depth estimates for underlying GC content (similar method to Sudmant et al. 2013 [9]), and normalised by median read depth per individual. We implemented a Gaussian Hidden Markov Model [10] to call structural variants from read depth.

#### Results

We identified CNVs in ancient individuals at ten loci using digital Comparative Genomic Hybridization [11] (Table 26; Figures S109-S128). Although most of the observed CNVs (including duplications at 15q11.2 and *CHRNA7*, and CNVs spanning parts of the TAR locus and 22q11.2 distal) have not been unambiguously associated with disease in large studies, the identified CNVs include deletions and duplications that have been associated with developmental delay, dysmorphic features, and neuropsychiatric abnormalities such as autism (most notably at 1q21.1, 3q29, 16p12.1 and the DiGeorge/VCFS locus, but also deletions at 15q11.2 and duplications at 16p13.11). However, phenotypes and risk associated with these structural variants vary widely, and recent population-based studies [12,13] suggest that they may be more common in the general population than previously thought [1,2]. Overall, the carrier frequency in ancient samples is similar to that reported in the UK Biobank (1.25% vs 1.6% at 15q11.2 and *CHRNA7* combined, and 0.8% vs 1.1% across the remaining loci combined) [12]. These results suggest that large, recurrent CNVs that can lead to several pathologies were present in ancient populations at similar frequencies as modern populations.

| Region | Ancient Deletions | SGDP & HGDP Deletions | UK Biobank Deletions | Ancient Duplications | SGDP & HGDP Duplications | UK Biobank Duplications |
|---|---|---|---|---|---|---|
| **1q21.1** | 0 (0%) | 0 (0%) | 113 (0.027%) | 1 (0.069%) | 0 (0%) | 177 (0.042%) |
| **3q29** | 1 (0.069%) | 0 (0%) | 9 (0.002%) | 1 (0.069%) | 0 (0%) | 5 (0.001%) |
| **15q11.2** | 4 (0.28%) | 2 (0.18%) | 1664 (0.39%) | 10 (0.69%) | 9 (0.82%) | 2041 (0.48%) |
| **15q11q13 (BP3-BP4)** | 1 (0.069%) | 0 (0%) | 16 (0.004%) | 0 (0%) | 0 (0%) | 53 (0.013%) |
| **15q13.3 (*CHRNA7*)** | 0 (0%) | 1 (0.09%) | 10 (0.002%) | 4 (0.28%) | 8 (0.73%) | 3031 (0.72%) |
| **16p12.1** | 1 (0.069%) | 0 (0%) | 246 (0.058%) | 1 (0.069%) | 0 (0%) | 202 (0.048%) |
| **16p13.11** | 1 (0.069%) | 0 (0%) | 131 (0.031%) | 4 (0.28%) | 0 (0%) | 828 (0.2%) |
| **22q11.2 (distal)*** | 4 (0.28%) | 0 (0%) | N/A | 13 (0.90%) | 6 (0.55%) | N/A |
| **DiGeorge-VCFS** | 0 (0%) | 0 (0%) | 10 (0.0024%) | 1 (0.069%) | 0 (0%) | 280 (0.066%) |
| **TAR*** | 1 (0.069%) | 0 (0%) | N/A | 2 (0.14%) | 0 (0%) | N/A |

**Table 26.** Table of 10 pathogenic loci (out of 50 examined) with structural variants identified in the ancient dataset**.** For each dataset, we report the prevalence of each SV as both the number of individuals identified as well as the percentage in each population (ancient dataset: 1442 samples; modern human Simons Genome Diversity Project (SGDP) [5] and Human Genome Diversity Project (HGDP) [4] dataset: 1093 samples; UK Biobank dataset: 421268 samples).

*Ancient SVs do not span the entire locus, and therefore we cannot compare the UK Biobank frequencies for these loci (which span the entirety of the locus) to the ancient frequencies.

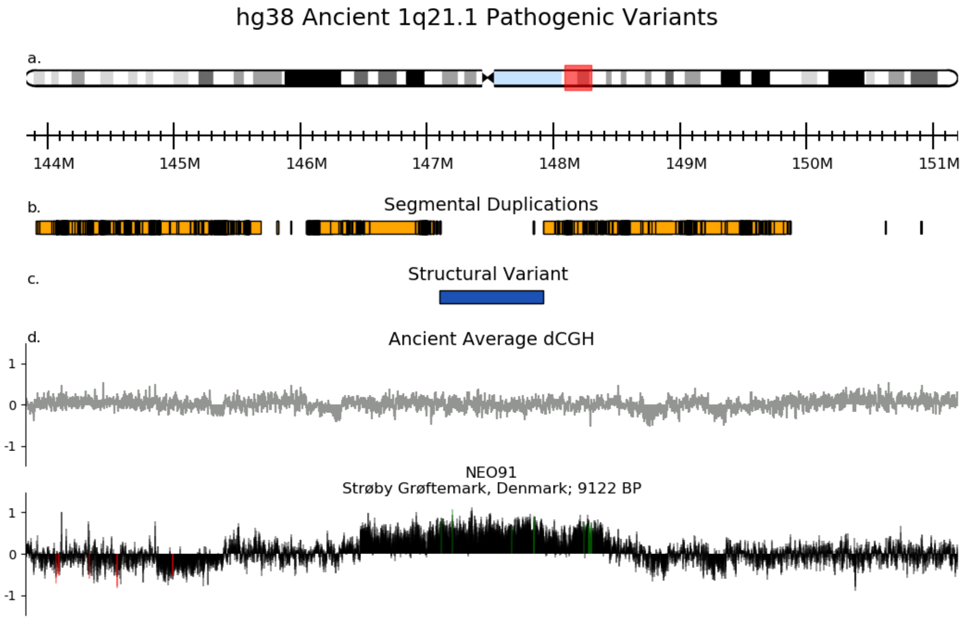


**Figure S109.** Microduplication of an ancient individual at the 1q21.1 locus from the hg38 fastq dataset. This 2 Mb microduplication is known to be associated with increased risk of neurological and psychiatric problems, delayed development, Tetralogy of Fallot, and micro/macrocephaly [14–16]. a. Chromosome 1 G-bands and axis bar (in Mbp). b. Segmental Duplications of >1000 bases of non-RepeatMasked [17] sequence from the UCSC Genome Browser [18]. c. Indices of the structural variant from the literature. d. The average dCGH (in grey) is computed as follows: first, a non-noisy individual for this locus is selected; then, for every other individual, the $\log_2$ ratio of its copy number values over the non-noisy individuals' copy number values is calculated; finally, the average of these ratios is depicted. The $\log_2$ ratio of NEO91's copy number values over those of the non-noisy individual is shown below (green = the ratio is at least 1.5 the standard deviation above the average dCGH; red = 1.5 std. dev. below). While there is little deviation from the "norm" copy number at this locus on average, the dCGH of NEO91 is significantly greater than the average and indicates a duplication.

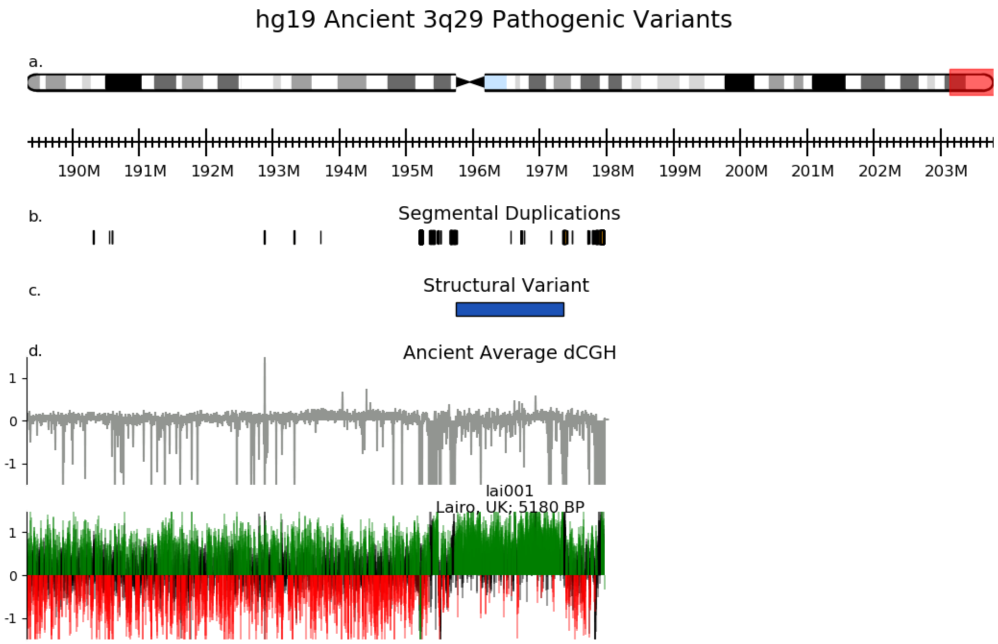


**Figure S110.** Microduplication of an ancient individual at the 3q29 locus from the hg19 BAM dataset. At 3q29, microdeletions and microduplications are associated with speech and developmental delay, cleft palate, microcephaly, and increased risk for psychiatric disorders [19,20]. No modern individuals in the SGDP or HGDP datasets present with a microdeletion or microduplication at 3q29.

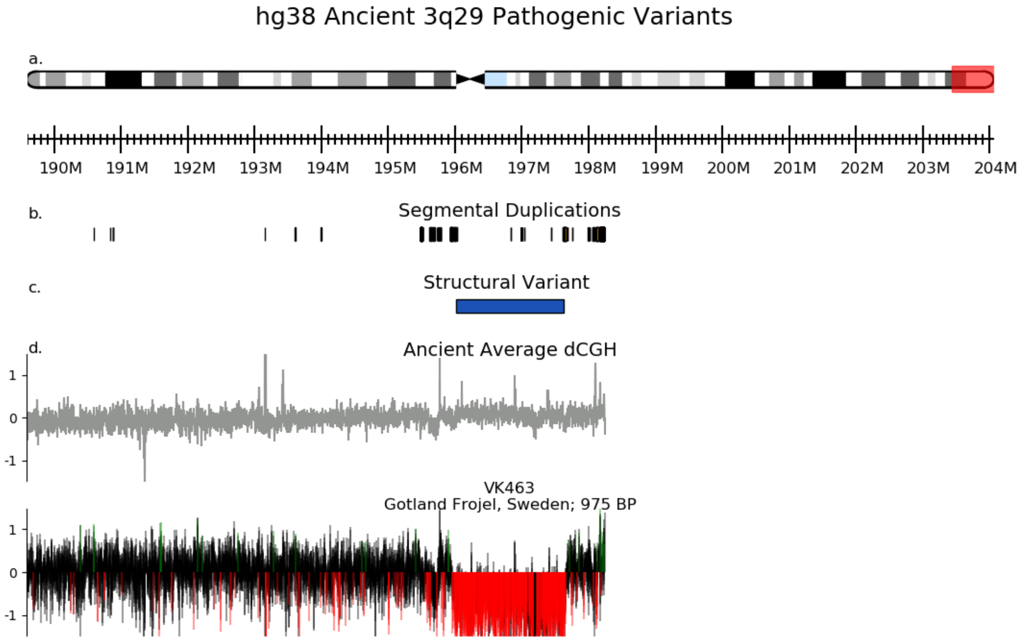


**Figure S111.** Microdeletion of an ancient individual at the 3q29 locus from the hg38 fastq dataset.

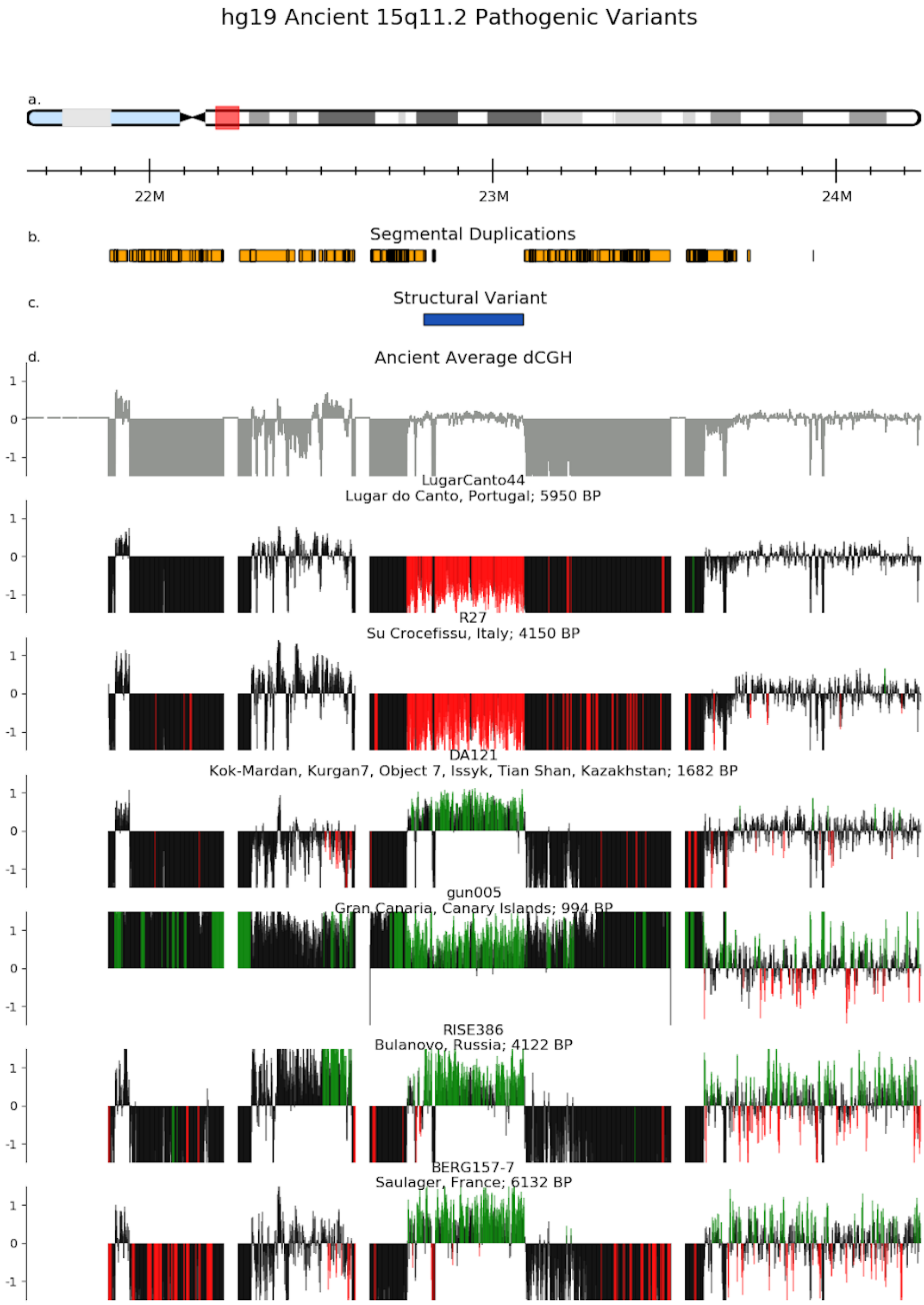


**Figure S112.** Two microdeletions and four microduplications are present at the 15q11.2 locus from the hg19 BAM dataset. Although the microdeletion is considered to confer modest risk of schizophrenia, epilepsy, learning problems, and ADHD, the microduplication is largely considered to be benign [21–24]. The blocks flanking the structural variant with copy number 0 are where duplicated reads have been filtered from the dataset.

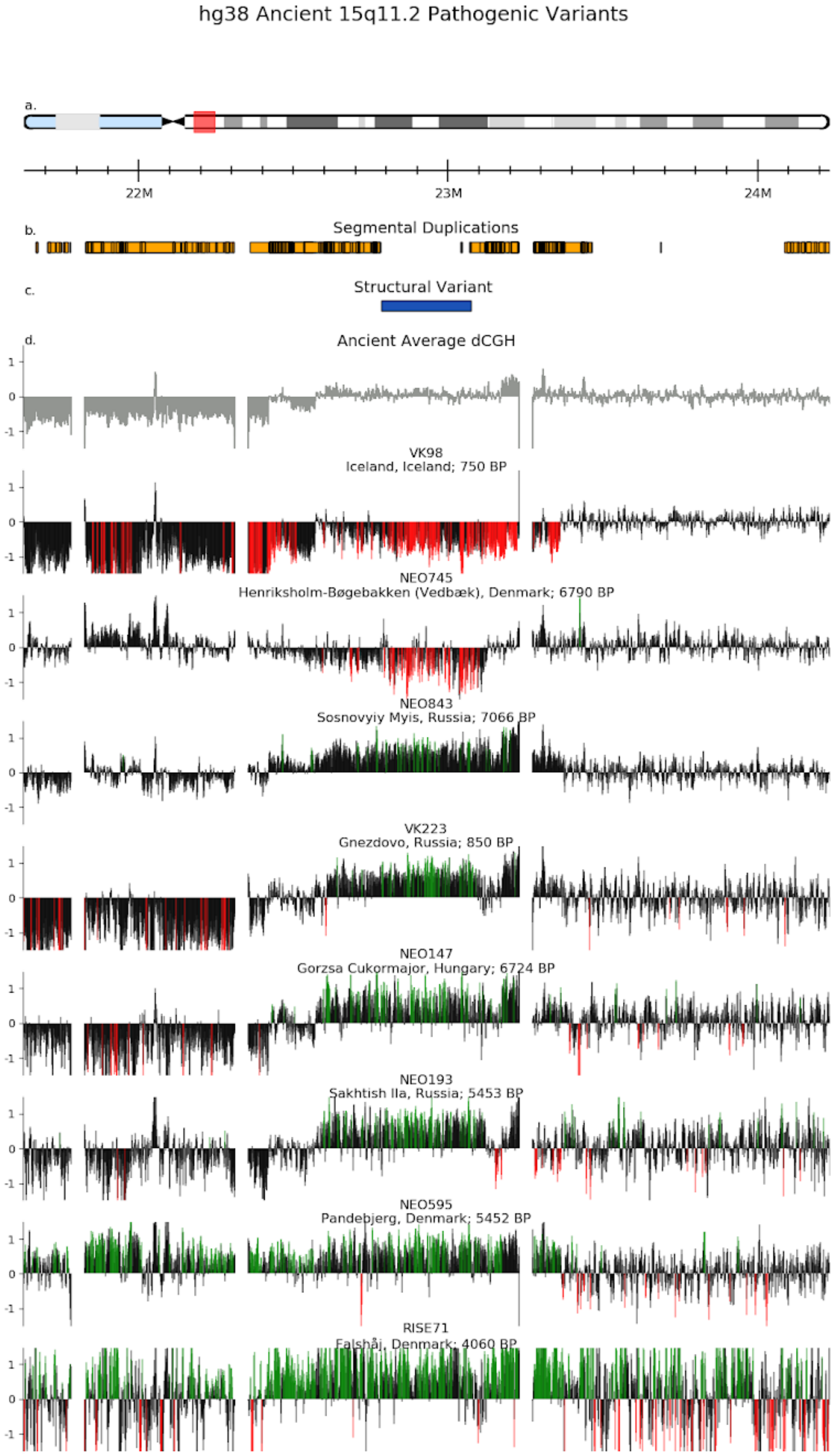


**Figure S113.** Two microdeletions and six microduplications are present at the 15q11.2 locus from the hg38 fastq dataset.

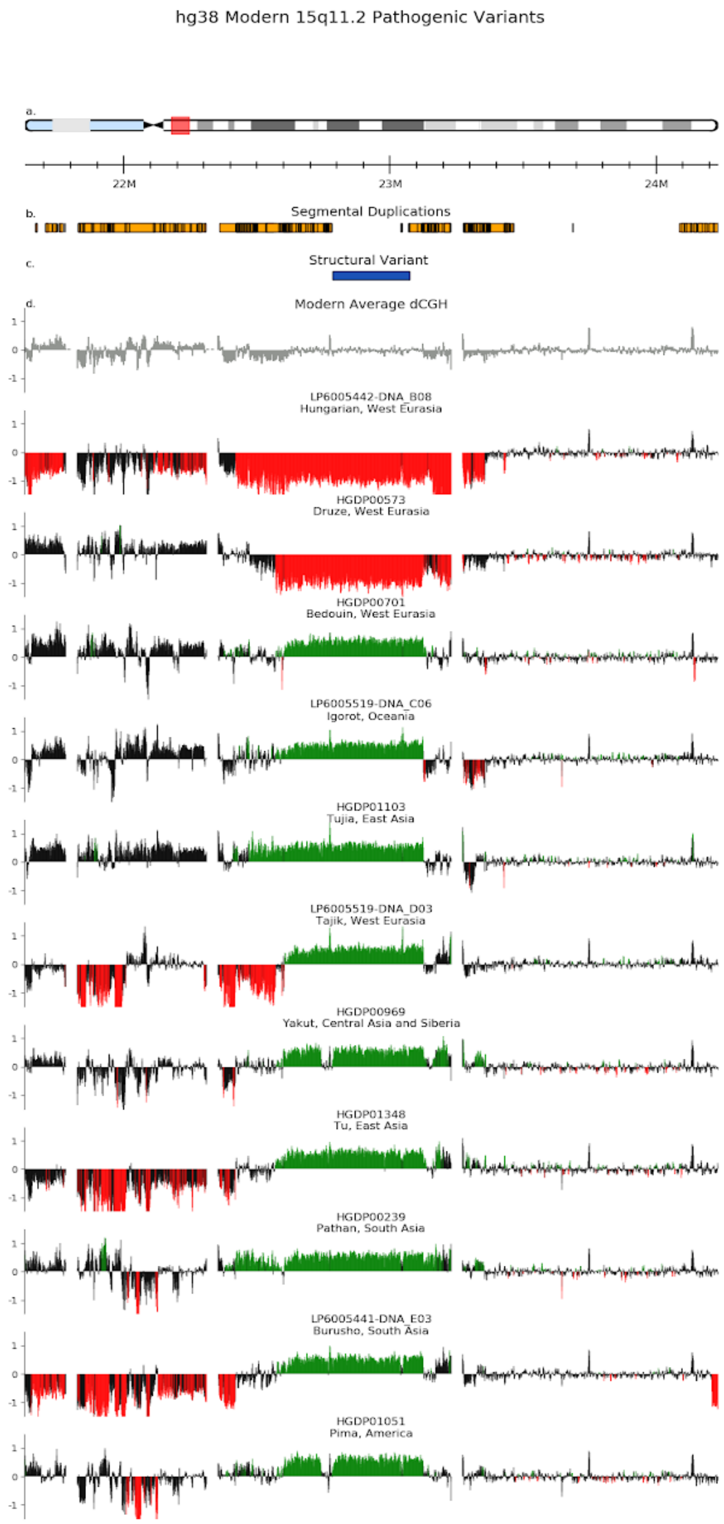


**Figure S114.** Two microdeletions and nine microduplications are present at the 15q11.2 locus from the modern human dataset. There are perhaps two duplications present – a long (HGDP01103 and HGDP00239) and a short (the other six humans).

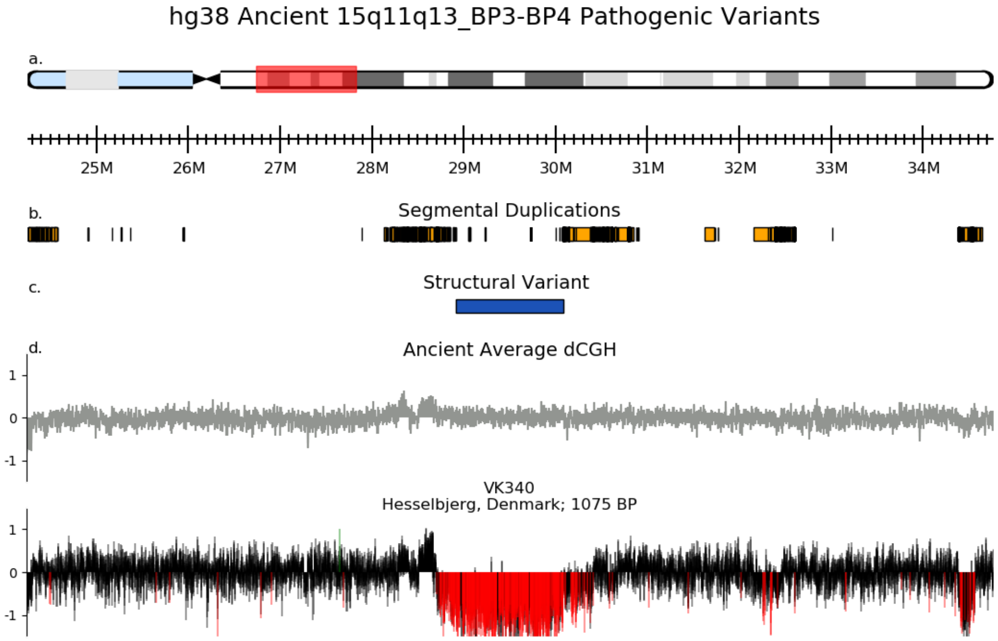


**Figure S115.** Microdeletion of an ancient individual at the 15q11q13 (breakpoints BP3-BP4) locus from the hg38 fastq dataset. Although the BP3-BP4 microdeletion at 15q11q13 has not been formally associated with disease, carriers have been reported with short stature, microcephaly, hypotonia, and facial dysmorphia [25]. None of the studied modern individuals have the deletion.

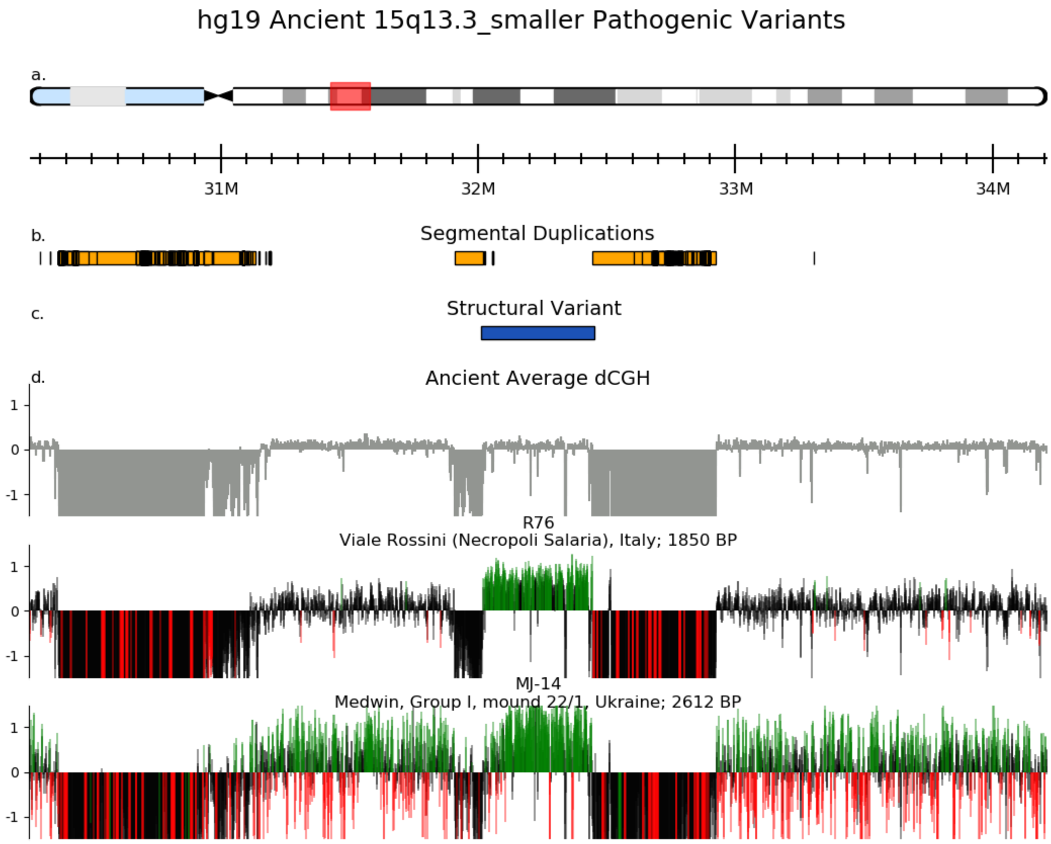


**Figure S116.** Two microduplications are present at the 15q13.3 (*CHRNA7*) locus from the hg19 BAM dataset. The *CHRNA7* microdeletion has been associated with developmental delay and psychiatric disorders, but it is less clear which (if any) phenotypes are associated with the *CHRNA7* microduplication and there is evidence that *CHRNA7* duplication carriers have just as good cognitive function as others [26].

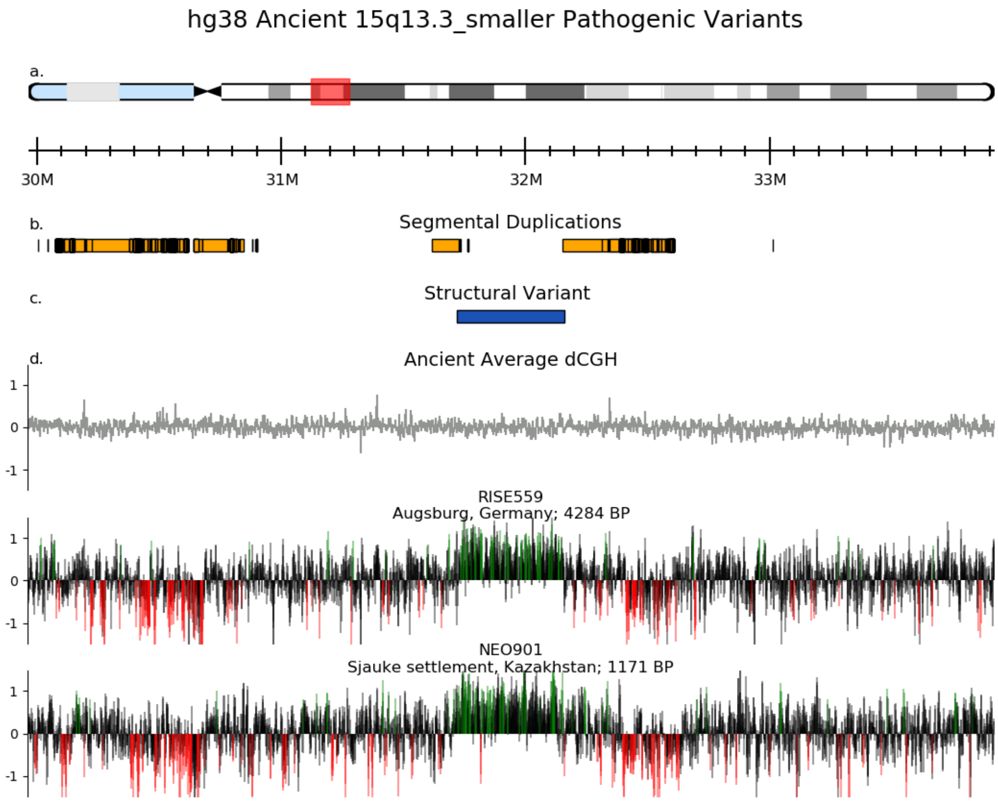


**Figure S117.** Two microduplications are present at the 15q13.3 (*CHRNA7*) locus from the hg38 fastq dataset.

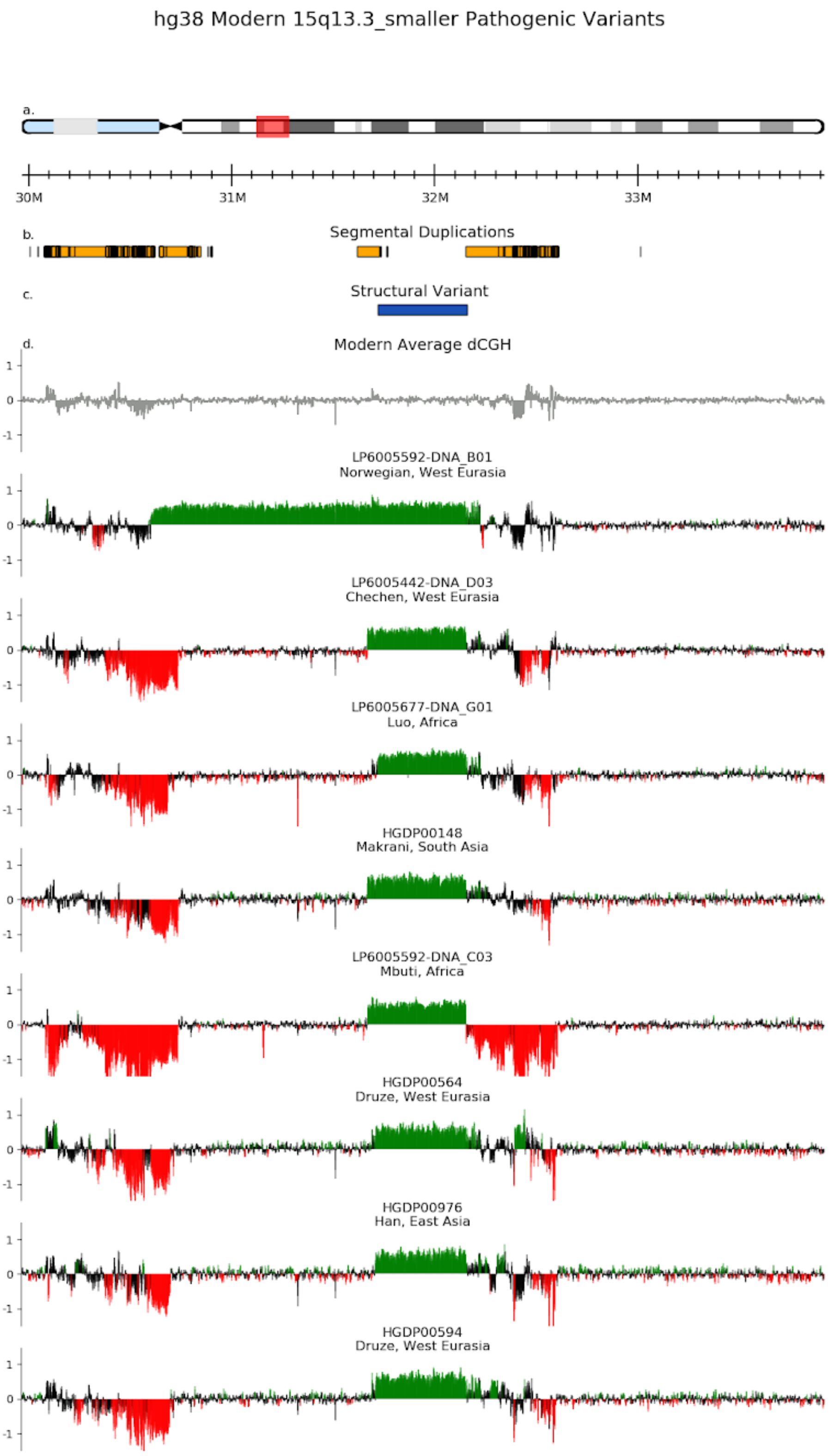


**Figure S118.** Two different microduplications are present at the 15q13.3 (*CHRNA7*) locus from the modern human dataset.

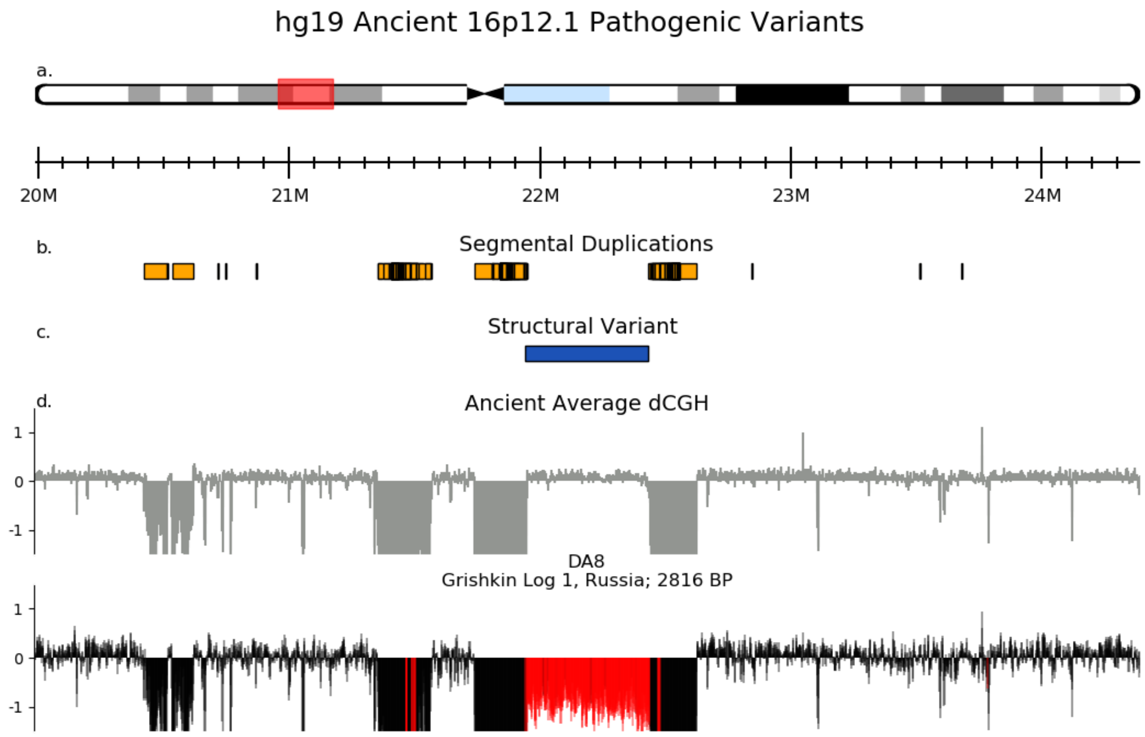


**Figure S119.** One ancient individual from the hg19 BAM dataset has a deletion at the 16p12.1 locus. At this locus, microdeletions and microduplications are associated with developmental delay, cognitive impairment, growth impairment, cardiac malformations, epilepsy, and psychiatric and behavioural problems [27,28].

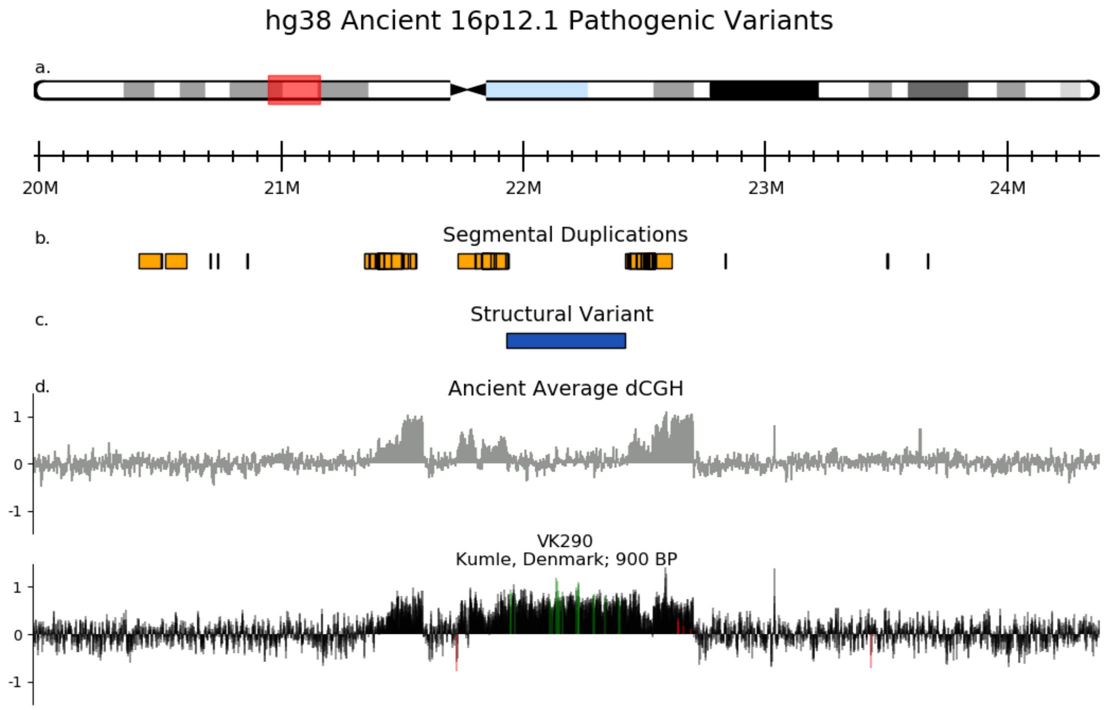


**Figure S120.** One ancient individual from the hg38 fastq dataset has a duplication at the 16p12.1 locus.

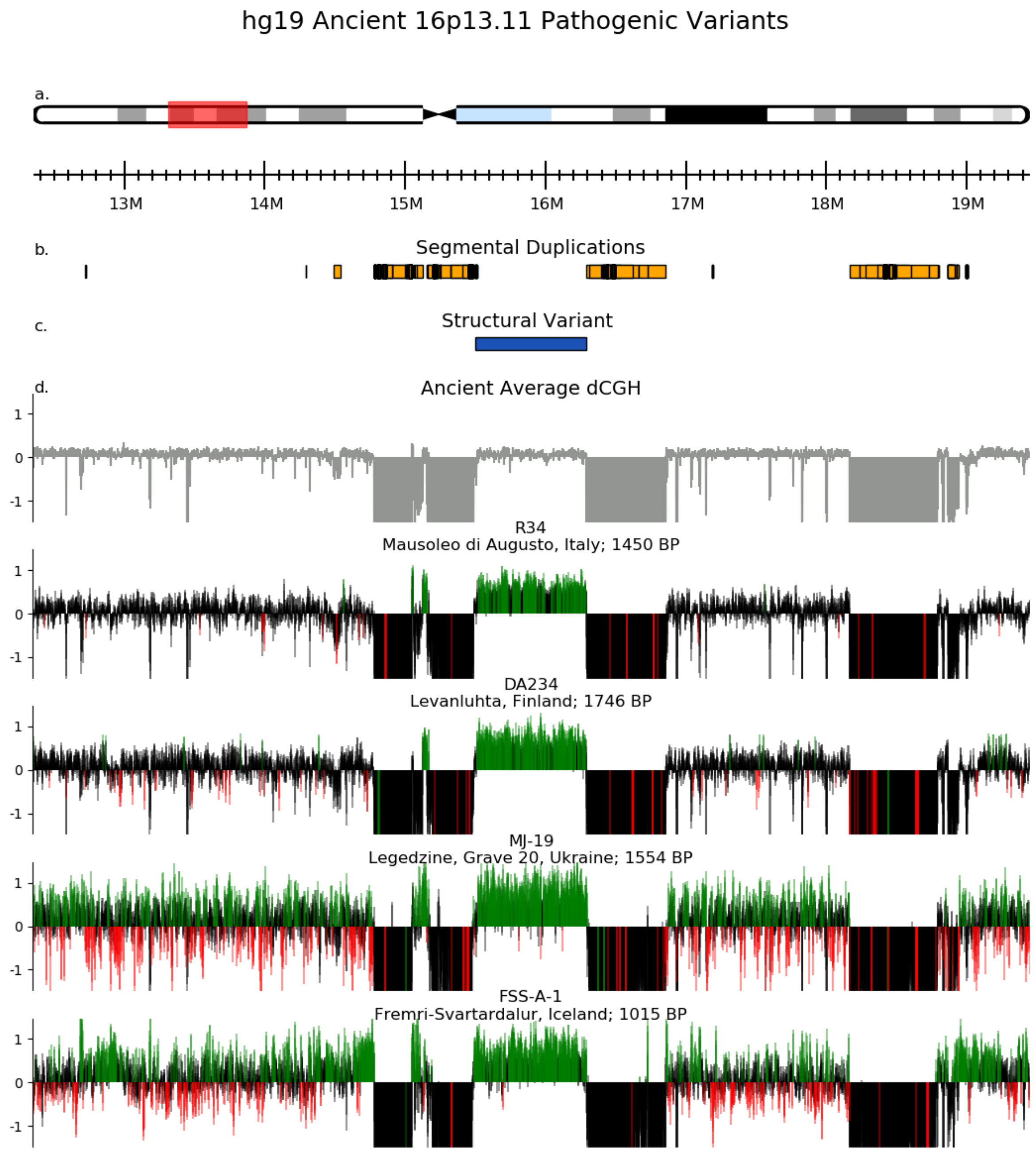


**Figure S121.** Four microduplications are present at the 16q13.11 locus from the hg19 BAM dataset. At the 16p13.11 locus, microduplications and microdeletions have been associated with several phenotypes including behavioural abnormalities, developmental delay, congenital heart defects, and skeletal abnormalities [29,30].

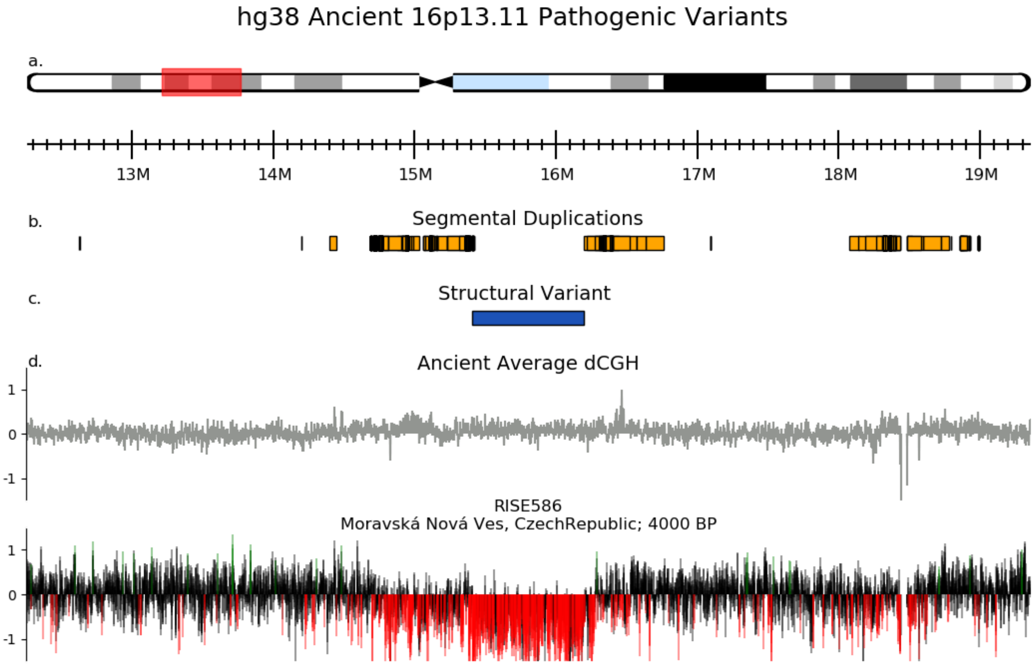


**Figure S122.** One microdeletion is present at the 16q13.11 locus from the hg38 fastq dataset.

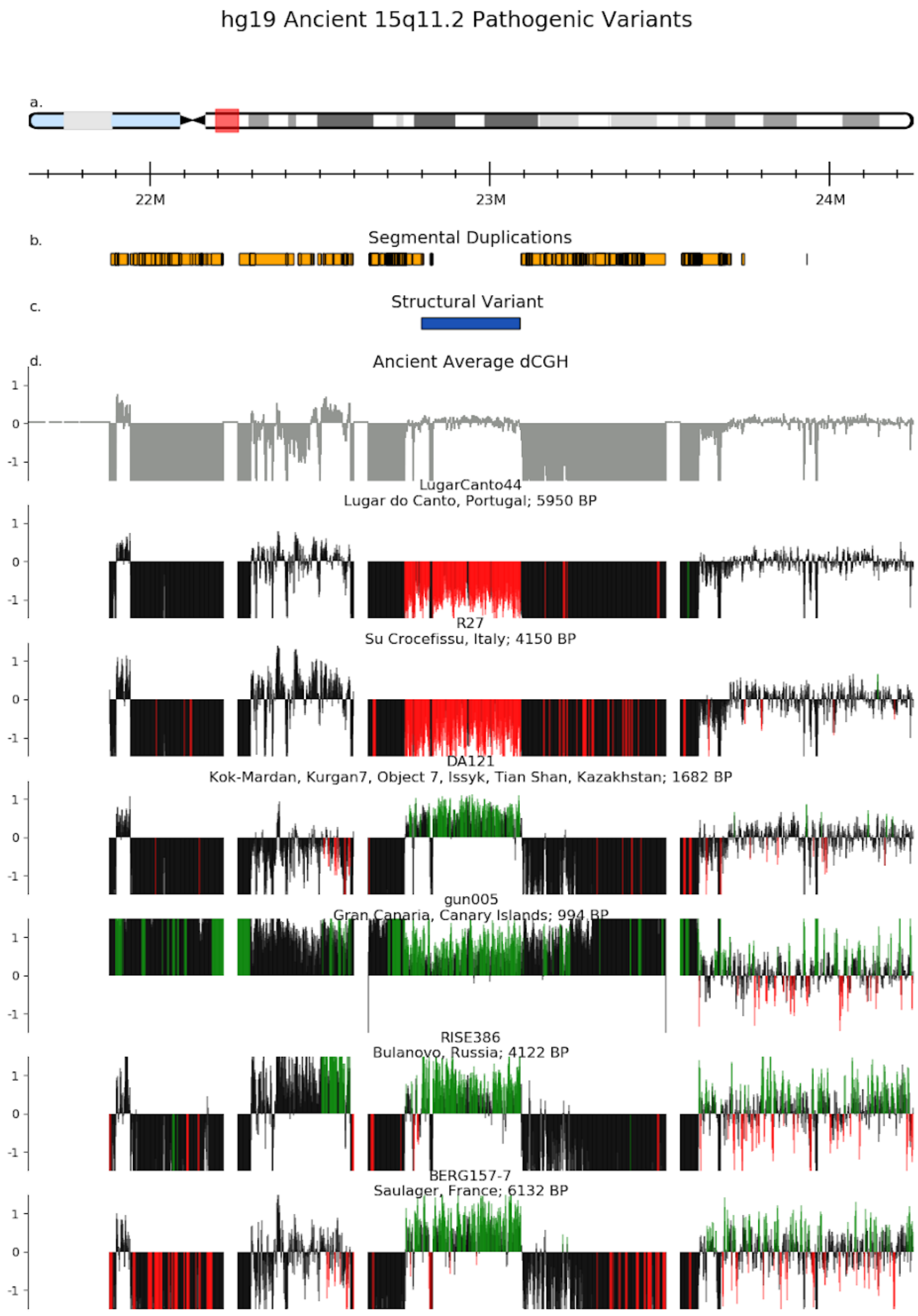

hg19 Ancient 15q11.2 Pathogenic Variants
a.
22M
23M
24M
b.
Segmental Duplications
c.
Structural Variant
d.
Ancient Average dCGH
LugarCanto44
Lugar do Canto, Portugal; 5950 BP
R27
Su Crocefissu, Italy; 4150 BP
DA121
Kok-Mardan, Kurgan7, Object 7, Issyk, Tian Shan, Kazakhstan; 1682 BP
gun005
Gran Canaria, Canary Islands; 994 BP
RISE386
Bulanovo, Russia; 4122 BP
BERG157-7
Saulager, France; 6132 BP

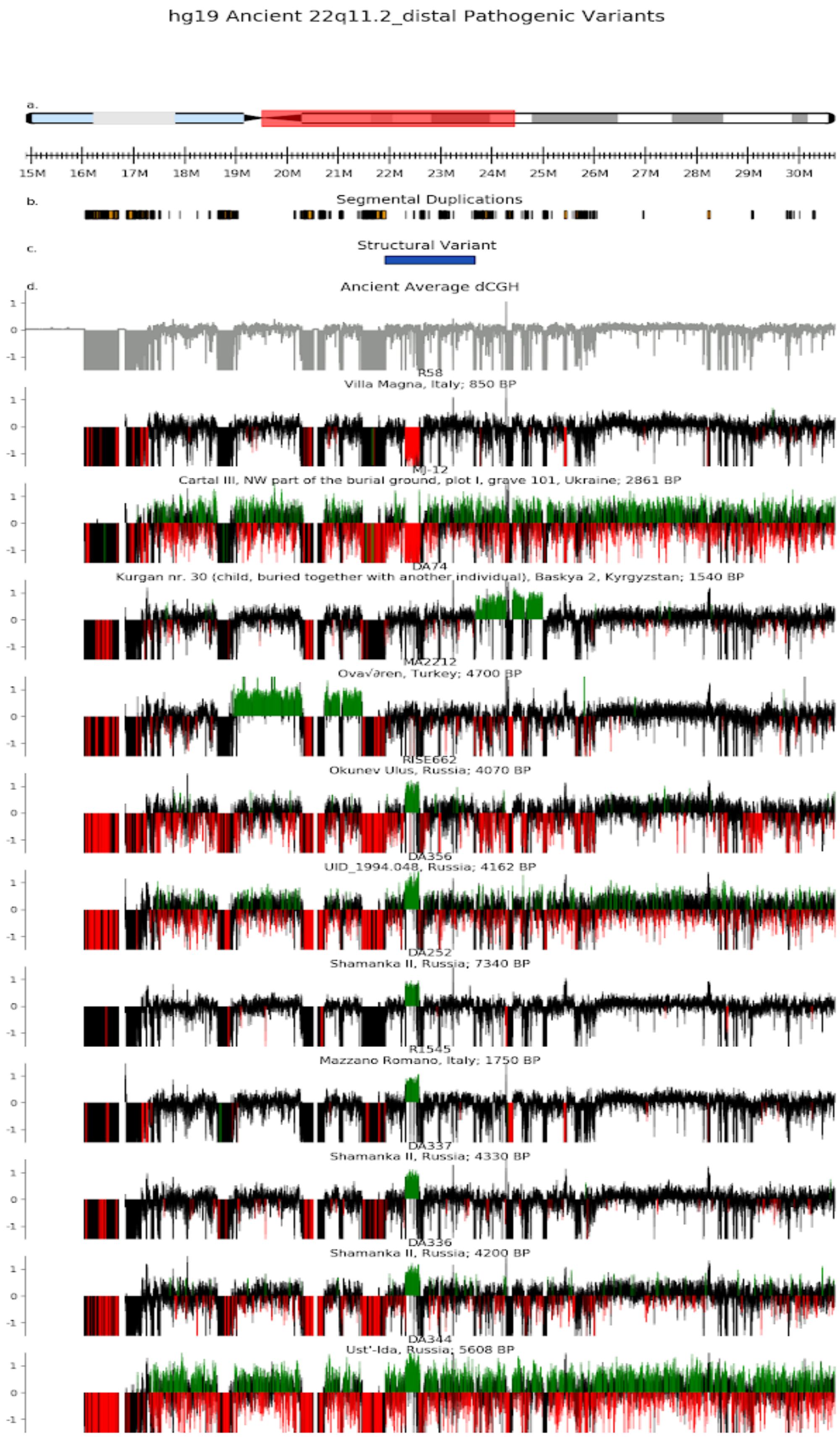
hg19 Ancient 22q11.2_distal Pathogenic Variants
a.
15M 16M 17M 18M 19M 20M 21M 22M 23M 24M 25M 26M 27M 28M 29M 30M
b.
Segmental Duplications
c.
Structural Variant
d.
Ancient Average dCGH
R58
Villa Magna, Italy; 850 BP
MJ-12
Cartal III, NW part of the burial ground, plot I, grave 101, Ukraine; 2861 BP
DA74
Kurgan nr. 30 (child, buried together with another individual), Baskya 2, Kyrgyzstan; 1540 BP
MA2212
Ova√ǝren, Turkey; 4700 BP
RISE662
Okunev Ulus, Russia; 4070 BP
DA356
UID_1994.048, Russia; 4162 BP
DA252
Shamanka II, Russia; 7340 BP
R1545
Mazzano Romano, Italy; 1750 BP
DA337
Shamanka II, Russia; 4330 BP
DA336
Shamanka II, Russia; 4200 BP
DA344
Ust'-Ida, Russia; 5608 BP

**Figure S123.** Two microdeletions and nine microduplications are present at the 22q11.2 (distal) locus from the hg19 BAM dataset. At the 22q11.2 locus, the *TOP3B* microdeletion has been reported to be associated with autism, learning disabilities, and dysmorphic features [31,32], and *TOP3B* knockout mice exhibit behaviour similar to psychiatric disorders and cognitive impairment [33], but risk estimates from large-scale population-based studies are lacking. All hg19 ancient individuals have this *TOP3B* structural variant with identical breakpoints as a 12-year-old patient with autism, cognitive impairment, and dysmorphic features [34], while two other ancient individuals have other duplication breakpoints (DA74 and MA2212).

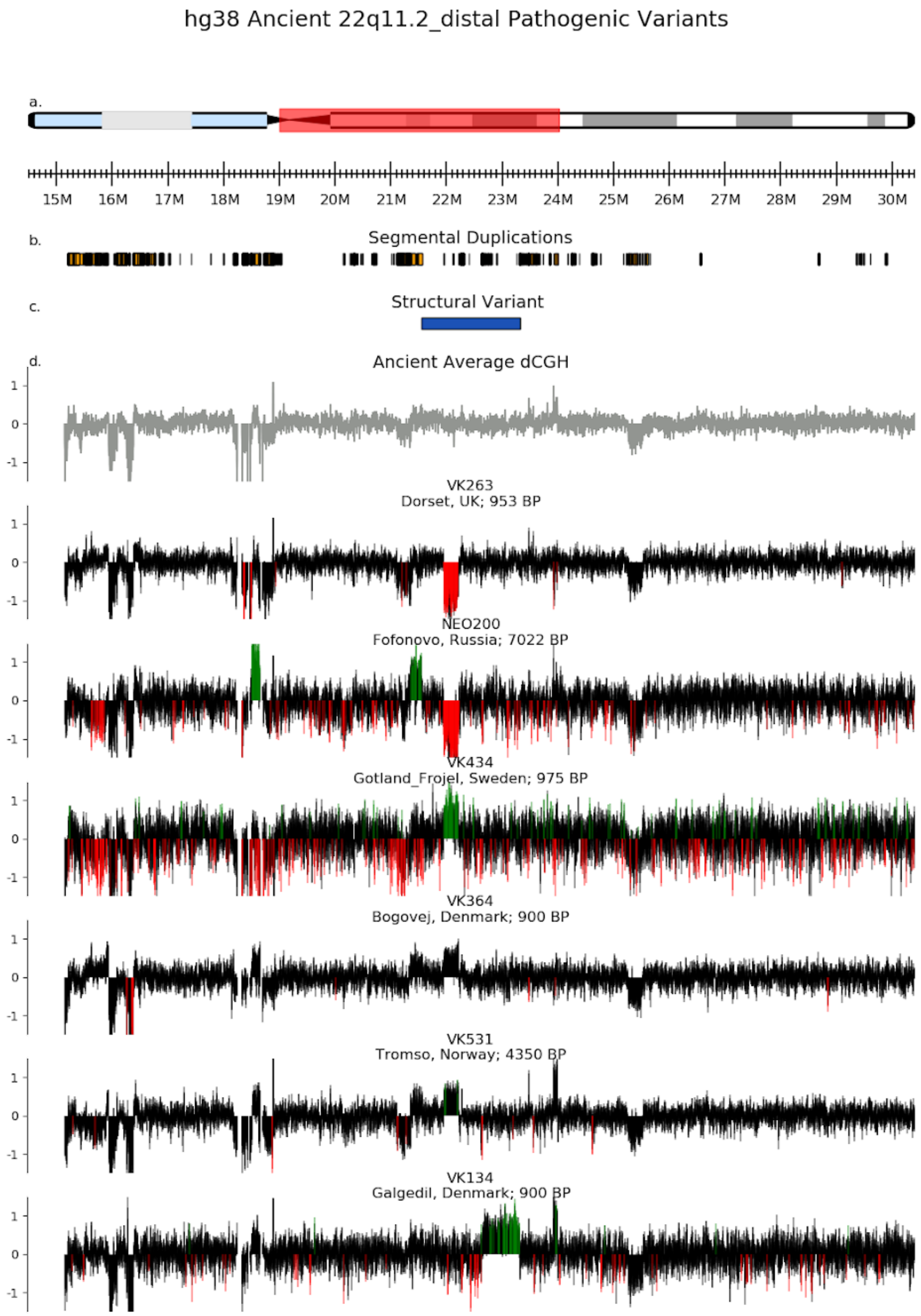


**Figure S124.** Two microdeletions and four microduplications are present at the 22q11.2 (distal) locus from the hg19 BAM dataset.

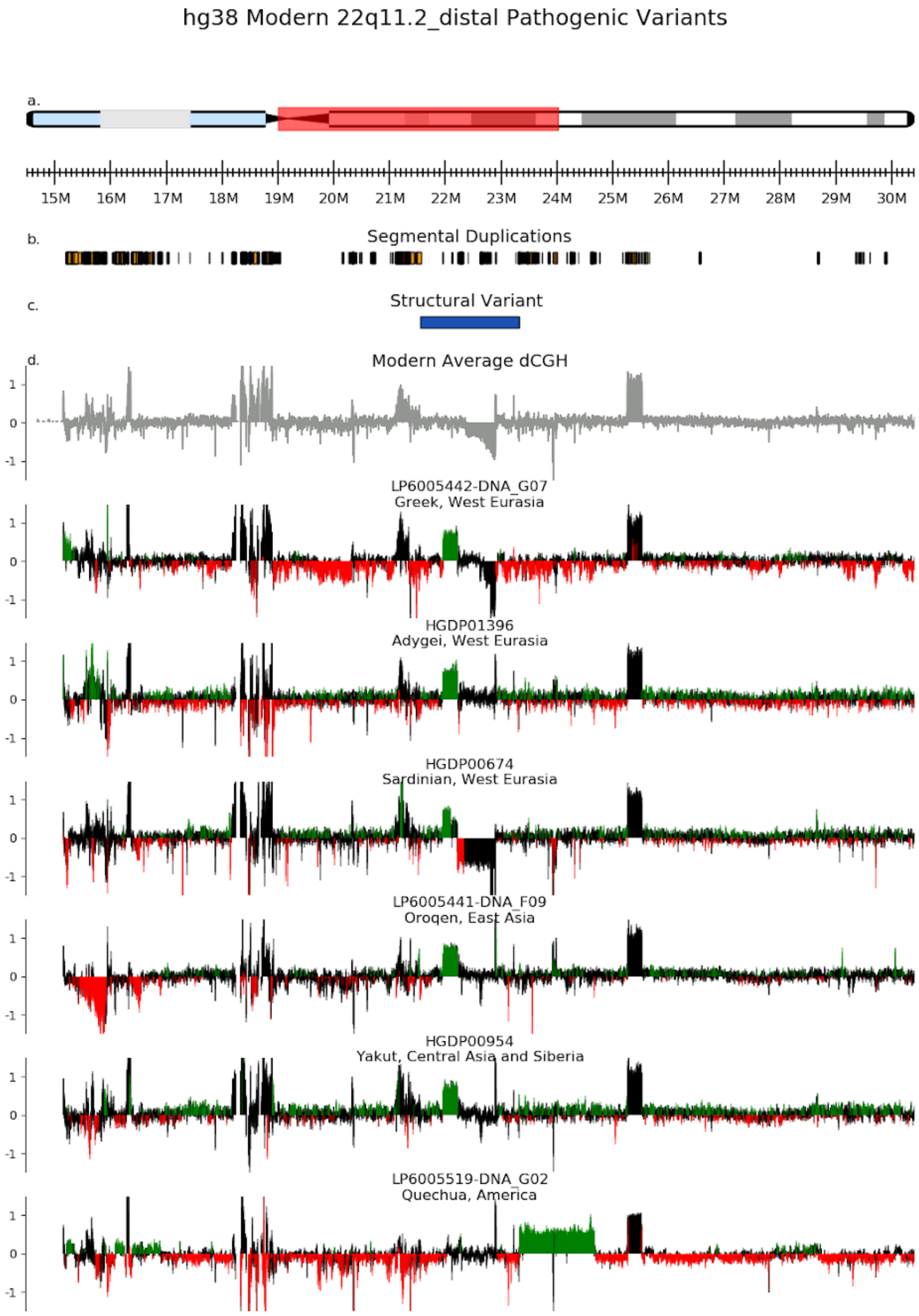


**Figure S125.** Six microduplications (five of which directly overlap with *TOP3B*) are present at the 22q11.2 (distal) locus from the modern human dataset.

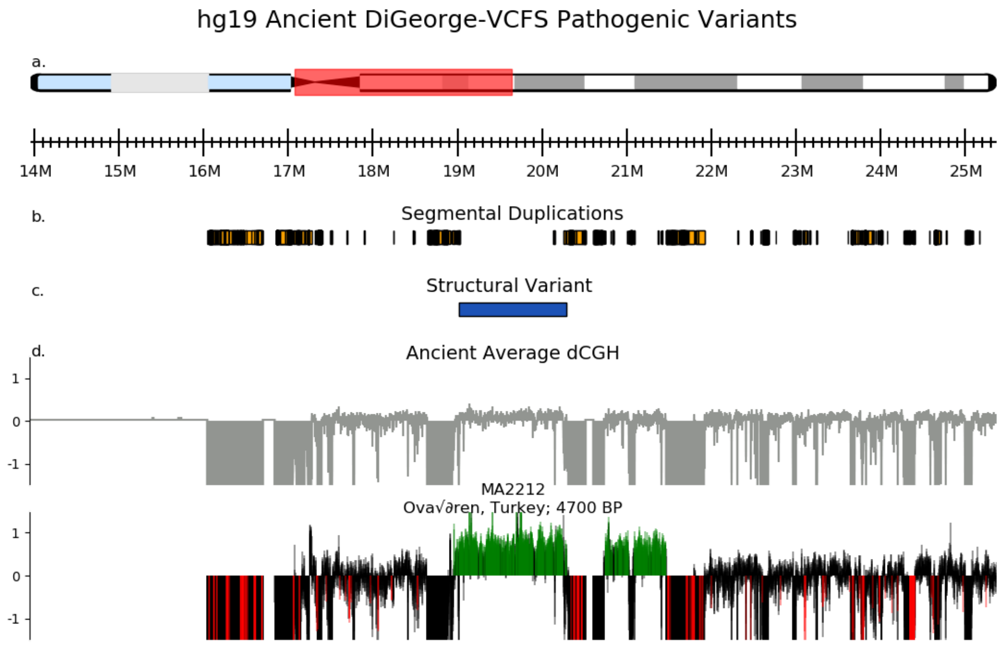


**Figure S126.** One ancient individual has a duplication at the DiGeorge-VCFS locus from the hg19 BAM dataset. Duplications at this locus are associated with schizophrenia as well as cardiovascular, parathyroid, thymic, and craniofacial abnormalities [35].

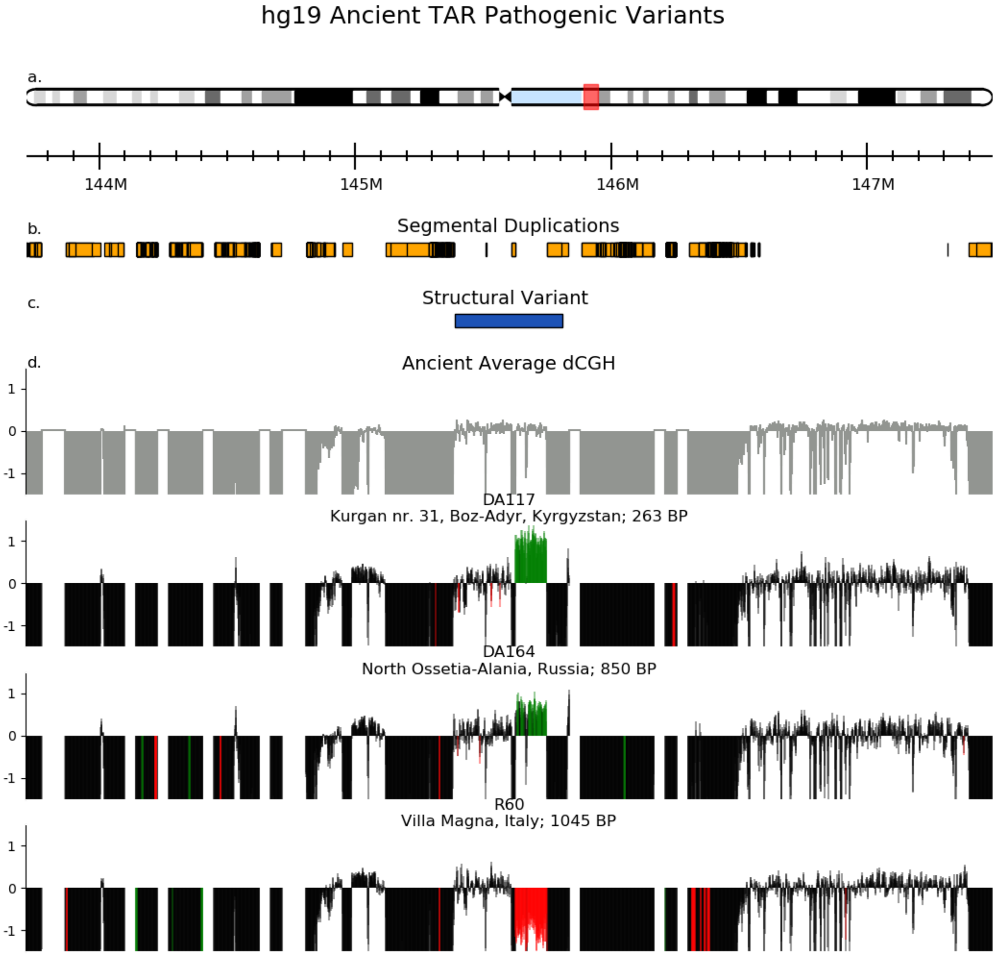


**Figure S127.** Ancient TAR pathogenic variants. TAR (thrombocytopenia with absent radius) syndrome is a rare genetic disorder featuring the absence of the radius bone in the forearm [36]; while no ancient individuals have a structural variant across the entire breakpoint of the syndrome, one ancient individual has a deletion across part of the locus and two ancient individuals have a duplication across the same breakpoints. The possible pathological implication of these CNVs is therefore unknown.

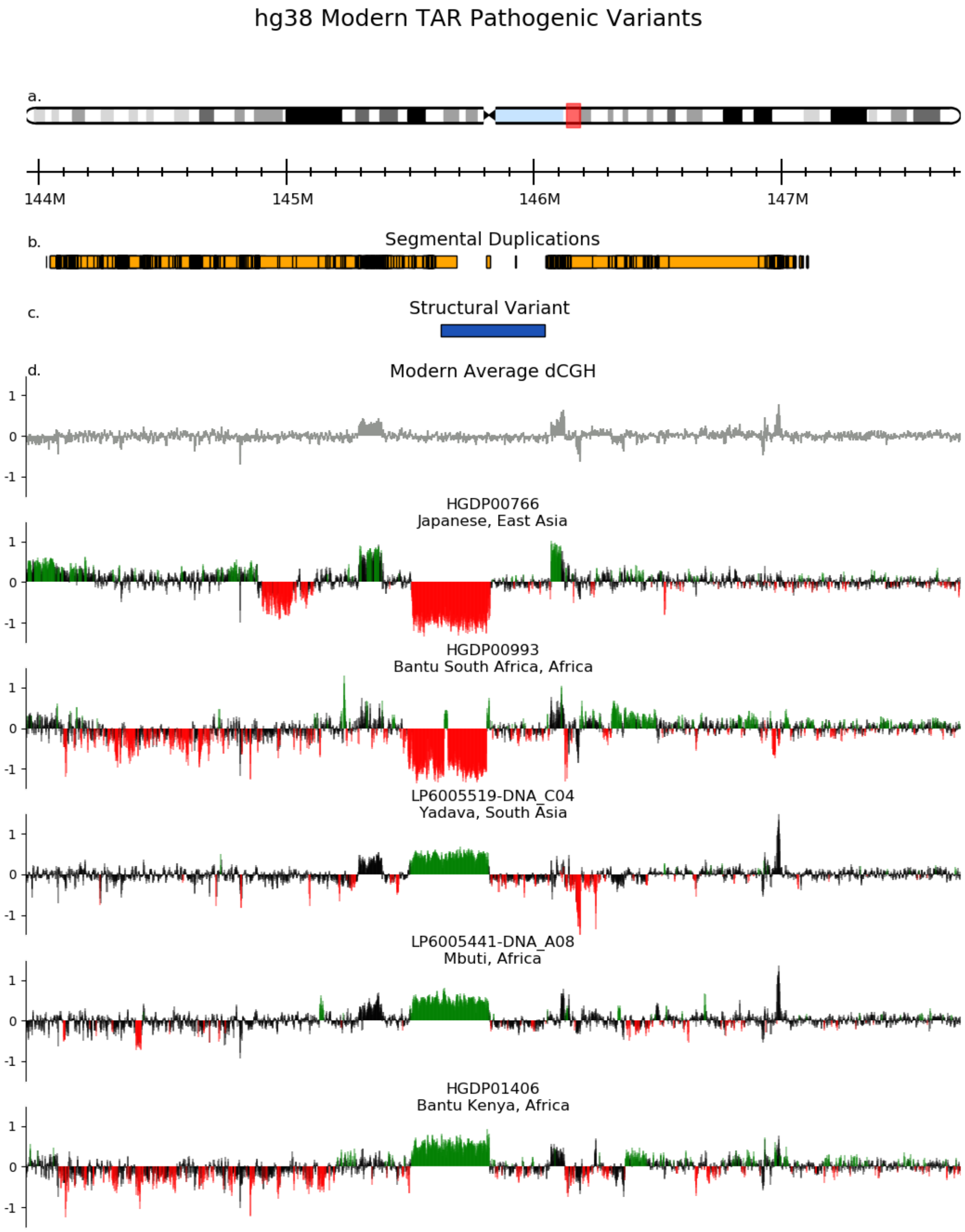


**Figure S128.** Two deletions and three duplications are present in the modern human dataset, but with different breakpoints than the ancient human structural variants.

#### References

1. Girirajan, S., Campbell, C. D. & Eichler, E. E. Human copy number variation and complex genetic disease. *Annu. Rev. Genet.* **45**, 203–226 (2011).

2. Weise, A. *et al.* Microdeletion and microduplication syndromes. *J. Histochem. Cytochem.* **60**, 346–358 (2012).

3. Girirajan, S. *et al.* Phenotypic heterogeneity of genomic disorders and rare copy-number variants. *N. Engl. J. Med.* **367**, 1321–1331 (2012).

4. Bergström, A. *et al.* Insights into human genetic variation and population history from 929 diverse genomes. *Science* **367**, (2020).

5. Mallick, S. *et al.* The Simons Genome Diversity Project: 300 genomes from 142 diverse populations. *Nature* **538**, 201–206 (2016).

6. Li, H. Burrows-Wheeler Aligner. http://bio-bwa.sourceforge.net/.

7. Miles, A. *pysamstats: A fast Python and command-line utility for extracting simple statistics against genome positions based on sequence alignments from a SAM or BAM file*. (Github).

8. Benson, G. Tandem repeats finder: a program to analyze DNA sequences. *Nucleic Acids Res.* **27**, 573–580 (1999).

9. Sudmant, P. H. *et al.* Evolution and diversity of copy number variation in the great ape lineage. *Genome Res.* **23**, 1373–1382 (2013).

10. Leffler, E. M. *et al.* Resistance to malaria through structural variation of red blood cell invasion receptors. *Science* **356**, (2017).

11. Sudmant, P. H. *et al.* Diversity of human copy number variation and multicopy genes. *Science* **330**, 641–646 (2010).

12. Crawford, K. *et al.* Medical consequences of pathogenic CNVs in adults: analysis of the UK Biobank. *J. Med. Genet.* **56**, 131–138 (2019).

13. Olsen, L. *et al.* Prevalence of rearrangements in the 22q11. 2 region and population-based risk of neuropsychiatric and developmental disorders in a Danish

population: a case-cohort study. *The Lancet Psychiatry* **5**, 573–580 (2018).

14. Dolcetti, A. *et al.* 1q21.1 Microduplication expression in adults. *Genet. Med.* **15**, 282–289 (2013).

15. Mefford, H. C. *et al.* Recurrent rearrangements of chromosome 1q21.1 and variable pediatric phenotypes. *N. Engl. J. Med.* **359**, 1685–1699 (2008).

16. Brunetti-Pierri, N. *et al.* Recurrent reciprocal 1q21.1 deletions and duplications associated with microcephaly or macrocephaly and developmental and behavioral abnormalities. *Nat. Genet.* **40**, 1466–1471 (2008).

17. Bailey, J. A. *et al.* Recent segmental duplications in the human genome. *Science* **297**, 1003–1007 (2002).

18. Kent, W. J. *et al.* The human genome browser at UCSC. *Genome Res.* **12**, 996–1006 (2002).

19. Coyan, A. G. & Dyer, L. M. 3q29 microduplication syndrome: Clinical and molecular description of eleven new cases. *Eur. J. Med. Genet.* **63**, 104083 (2020).

20. Glassford, M. R., Rosenfeld, J. A., Freedman, A. A., Zwick, M. E. & Mulle, J. G. Novel features of 3q29 deletion syndrome: Results from the 3q29 registry. *Am. J. Med. Genet. A* **170**, 999–1006 (2016).

21. Stefansson, H. *et al.* CNVs conferring risk of autism or schizophrenia affect cognition in controls. *Nature* **505**, 361–366 (2014).

22. Kirov, G. *et al.* The penetrance of copy number variations for schizophrenia and developmental delay. *Biol. Psychiatry* **75**, 378–385 (2014).

23. Gudmundsson, O. O. *et al.* Attention-deficit hyperactivity disorder shares copy number variant risk with schizophrenia and autism spectrum disorder. *Transl. Psychiatry* **9**, 258 (2019).

24. Jønch, A. E. *et al.* Estimating the effect size of the 15Q11.2 BP1–BP2 deletion and its contribution to neurodevelopmental symptoms: recommendations for practice. *J. Med. Genet.* **56**, 701–710 (2019).

25. Rosenfeld, J. A. *et al.* Deletions flanked by breakpoints 3 and 4 on 15q13 may

contribute to abnormal phenotypes. *Eur. J. Hum. Genet.* **19**, 547–554 (2011).

26. Kendall, K. M. *et al.* Cognitive performance and functional outcomes of carriers of pathogenic copy number variants: analysis of the UK Biobank. *Br. J. Psychiatry* **214**, 297–304 (2019).

27. Girirajan, S., Pizzo, L., Moeschler, J. & Rosenfeld, J. *16p12.2 Recurrent Deletion*. (University of Washington, 1993).

28. D'Angelo, D. *et al.* Defining the Effect of the 16p11.2 Duplication on Cognition, Behavior, and Medical Comorbidities. *JAMA Psychiatry* **73**, 20–30 (2016).

29. Nagamani, S. C. S. *et al.* Phenotypic manifestations of copy number variation in chromosome 16p13.11. *Eur. J. Hum. Genet.* **19**, 280–286 (2011).

30. Allach El Khattabi, L. *et al.* 16p13.11 microduplication in 45 new patients: refined clinical significance and genotype-phenotype correlations. *J. Med. Genet.* **57**, 301–307 (2020).

31. Stoll, G. *et al.* Deletion of TOP3β, a component of FMRP-containing mRNPs, contributes to neurodevelopmental disorders. *Nat. Neurosci.* **16**, 1228–1237 (2013).

32. Ahmad, M. *et al.* Topoisomerase 3β is the major topoisomerase for mRNAs and linked to neurodevelopment and mental dysfunction. *Nucleic Acids Res.* **45**, 2704–2713 (2017).

33. Joo, Y. *et al.* Topoisomerase 3β knockout mice show transcriptional and behavioural impairments associated with neurogenesis and synaptic plasticity. *Nat. Commun.* **11**, 3143 (2020).

34. Kaufman, C. S., Genovese, A. & Butler, M. G. Deletion of TOP3B Is Associated with Cognitive Impairment and Facial Dysmorphism. *Cytogenet. Genome Res.* **150**, 106–111 (2016).

35. Agatsuma, S. & Hiroi, N. Chromosome 22q11 and schizophrenia. *Nihon Shinkei Seishin Yakurigaku Zasshi* **25**, 79–84 (2005).

36. Boussion, S. *et al.* TAR syndrome: Clinical and molecular characterization of a cohort of 26 patients and description of novel noncoding variants of RBM8A. *Hum. Mutat.* **41**, 1220–1225 (2020)
